# Genomes of ancient asexual mites appear streamlined in their architecture

**DOI:** 10.64898/2025.12.01.691511

**Authors:** Nadège Guiglielmoni, Mohammed Errbii, Hüsna Öztoprak, Viktoria Bednarski, Karim Gueddach, Lea Borgschulte, Svenja Wulsch, Marshall Brickheimer, Isabella Di Stefano, Lucy Jimenez, Jakob Zelzner, Anja Schuster, Christian Becker, Kerstin Becker, Miklos Balint, Ricarda Lehmitz, Ina Schaefer, Leonie Schardt, Anna Seniczak, Philipp H. Schiffer, Jens Bast

## Abstract

The long-term persistence of obligate asexual lineages represents one of the most enduring and critical paradoxes in evolutionary biology. Sexual reproduction, through meiotic recombination and segregation, enables the efficient removal of deleterious mutations and facilitates rapid adaptation to shifting environmental pressures. Lineages that lose sex are therefore classically predicted to experience genomic decay and face rapid extinction. Oribatid mites (Acari, Sarcoptiformes) represent a unique system for testing these predictions, as they feature multiple, ancient, and independent transitions to asexuality, providing a natural experiment on the evolutionary fate of asexual genomes.

We compared four sets of sister sexual and asexual species using high-quality nuclear genome assemblies to investigate the genomic consequences of long-term asexuality. Our study revealed a profound, reproductive-mode-dependent dichotomy in the evolution of genome architecture. Contrary to their expected genomic decay, asexual species have mostly streamlined genomes with less novel genes than their sexual sister species. In contrast, sexual species have acquired genetic innovations, encompassing both gene and transposable element content. These results challenge classical expectations of genomic deterioration in asexual species and might explain the long-term evolutionary persistence of oribatid mites.

## 1 Introduction

Reproduction is the fundamental characteristic of life on earth, yet the mechanisms by which organisms generate offspring vary substantially among taxa. Reproductive modes have significant effects on the evolutionary trajectories of species. In metazoans, sex including meiosis and outcrossing, is by far the most frequent form of reproduction and the few asexual lineages occupy terminal positions in phylogenetic trees [Engelstädter, 2008, Schwander and Crespi, 2009]. It is thought that the ”two-fold cost of sex” [Smith, 1971, Bell, 1982, Lehtonen et al., 2012], due to the production of males who cannot bear offspring, is offset by a gain of evolutionary advantages in natural populations. These advantages are rooted in the process of recombination, through which sex decouples linked loci, thereby increasing the effectiveness of selection. Sex thus promotes the fixation of beneficial mutations, purges deleterious mutations, increases genetic diversity, and accelerates adaptation [McDonald et al., 2016, Otto, 2021].

Conversely, the absence of recombination in obligate parthenogenesis is expected to impose constraints on the efficacy of selection, as linked loci cannot be decoupled. This phenomenon is known as Hill-Robertson interference, which causes beneficial and deleterious alleles to hitchhike together [Hill and Robertson, 1966, Felsenstein, 1974, Keightley and Otto, 2006], thus reducing overall genetic diversity and potentially leading to irreversible accumulation of deleterious mutations across the genome (”Muller’s Ratchet”) [Muller, 1964]. Genomes of asexual species might also be more prone to accumulate large-scale structural variants, which are constrained in sexual species by segregation and recombination [McElroy et al., 2021]. While asexual reproduction might still offer short-term evolutionary advantages, for example by enabling organisms to quickly colonize marginal habitats [Villegas et al., 2025, Shatilovich et al., 2023], asexual species are expected to go extinct on evolutionary short timescales due to these detrimental mechanisms degrading their genomes [Maynard Smith, 1978].

Beyond the accumulation of slightly deleterious mutations [Villegas et al., 2024], due to reduced efficacy of selection, the consequences of prolonged asexuality for genome evolution are less predictable and empirical evidence for the genomic consequences of asexuality in natural lineages remains limited. In sexual species, the spread of transposable elements (TEs) is expected, acting as ”sexually transmitted nuclear parasites” [Hickey, 1982, Arkhipova and Meselson, 2000], and inefficient purging of TEs [Nuzhdin and Petrov, 2003] could promote genome expansion via TE accumulation in asexual species as well. However, empirical studies show that TE dynamics in asexual lineages are highly variable and depend on the balance between transposition activity and host suppression mechanisms. The evolution of TEs toward reduced activity in asexual species can lower TE load over time due to alignments of evolutionary trajectories of hosts and TEs [Bast et al., 2019]. Horizontal gene transfer (HGT) has been proposed as an important mechanism shaping the genomes of asexual species and could allow for the generation of genetic novelty in the absence of sex, but analyses of HGTs in bdelloid rotifers and tardigrades were inconclusive due to poor assembly quality [Koutsovoulos et al., 2016, Wilson et al., 2018]. Gene loss, such as that of sex-related genes, may occur over long evolutionary timescales in asexual species. As the overall number of these genes is small, the contribution to genome size evolution is expected to be minimal [van der Kooi and Schwander, 2014, Hiraki et al., 2017, Schurko and Logsdon Jr, 2008]. In general, insertions, duplications, deletions, and other structural variants (SVs) can produce a wide range of genomic changes, and result in large-scale rearrangements [Ho et al., 2020]. For example, among some sexual nematode species of the genus *Caenorhabditis*, genome size variations were found to be associated mainly with insertions and duplications [Adams et al., 2023]. Some short-term studies of arthropod asexual lineages report generally lower frequencies of SVs at the population level compared to sexual relatives [Bast et al., 2016, Jaron et al., 2022]. However, these mechanisms have not been studied in high-quality, chromosome-scale assemblies of asexual species. In summary, the long-term evolutionary trajectories of genome architecture in asexual lineages remain largely unknown.

The scarcity of empirical data from natural asexual populations underscores the need for study systems that provide both evolutionary replicates and long-term trait persistence. Oribatid mites (Acari, Sarcoptiformes) meet these criteria: they are a diverse and speciose group of soil-dwelling arthropods, feature multiple independent transitions to asexuality [Pachl et al., 2021] and approximately 10% of species reproduce asexually through thelytokous parthenogenesis [Palmer and Norton, 1991, von Saltzwedel et al., 2014], whereby females produce only female offspring from unfertilized eggs. Although precise age estimates are challenging, some parthenogenetic lineages have likely persisted for up to tens of millions of years [Hammer and Wallwork, 1979, Schaefer et al., 2010, Dabert et al., 2010, Lozano-Fernandez et al., 2020], with a fossil specimen from the genus *Hydrozetes* being dated to the early Jurassic [Sivhed and Wallwork, 1978]. The thus far best age estimate for an ancient asexual suggests a minimum age of at least 20 million years for *Platynothrus peltifer* [Öztoprak et al., 2025]. Oribatid mites provide an exceptional opportunity for comparative analyses, encompassing sexual-asexual species pairs within the same genus (i.e *Hydrozetes*, *Oppiella*), outside the same genus (*Atropacarus*, *Steganacarus*), and entire asexual clades (*Platynothrus*, *Nothrus*) with sexual sister lineages (*Hermannia*). The repeated and independent transitions to asexual reproduction provide evolutionary replicates to examine the genomic consequences of long-term asexuality across substantial evolutionary timescales.

In this study, we generated nine new high-quality genome assemblies of sexual and asexual oribatid mite species. Together with two previously published chromosome-level assemblies [Öztoprak et al., 2025, Wulsch et al., 2023], we analyzed a total of 11 genome assemblies representing multiple independent transitions to asexuality within Oribatida; they included four sets of sister sexual and asexual species, and two additional sexual and asexual species without counterpart. In addition, 10 assemblies were generated from wild individuals. By investigating chromosome conservation from a synteny standpoint, purifying selection, HGTs, TEs, and SVs, we uncovered a substantial effect of reproductive mode: asexual oribatid mite genomes are more ”streamlined,” i.e. featuring smaller genomes, fewer gene copies, and fewer TE content than their sexual relatives, suggesting a trajectory toward genome streamlining over evolutionary time.

## 2 Results

### 2.1 Comparison of 11 highly-contiguous assemblies of sexual and asexual oribatid mite species

As a backbone for all analyses to assess the impact of reproductive mode on genome architecture, we generated highly contiguous nuclear genome assemblies for nine species, including two chromosome-level assemblies for *Atropacarus striculus* and *Steganacarus magnus* (Table 1, Figure 1, Supplementary Table S1). In addition, we included two previously published chromosome-level assemblies of *Platynothrus peltifer* and *Hermannia gibba* [Öztoprak et al., 2025, Wulsch et al., 2023] in our comparison. *k* -mer analyses revealed that all species were diploid (Supplementary Figure S1-2) and collapsed assemblies were produced. Contaminants were removed from the final assemblies (Supplementary Figure S3-11). Long-read mitochondrial genome assemblies were generated for all species, with the exception of *Hypochthonius rufulus*. Nuclear genome assembly sizes range from 100.6 Mb (*H. lacustris*) to 377.0 Mb (*H. gibba*) (Table 1). Species have a mid-level heterozygosity (Supplementary Figure S2). A minimum of 2,753 single-copy orthologs was found in *A. striculus*, and a maximum of 2,839 orthologs in *N. palustris*, over a total of 2,934 orthologs in the Arachnida odb10 lineage, showing a high completeness in addition to *k* -mer comparisons (Supplementary Table 1).

**Figure 1:**
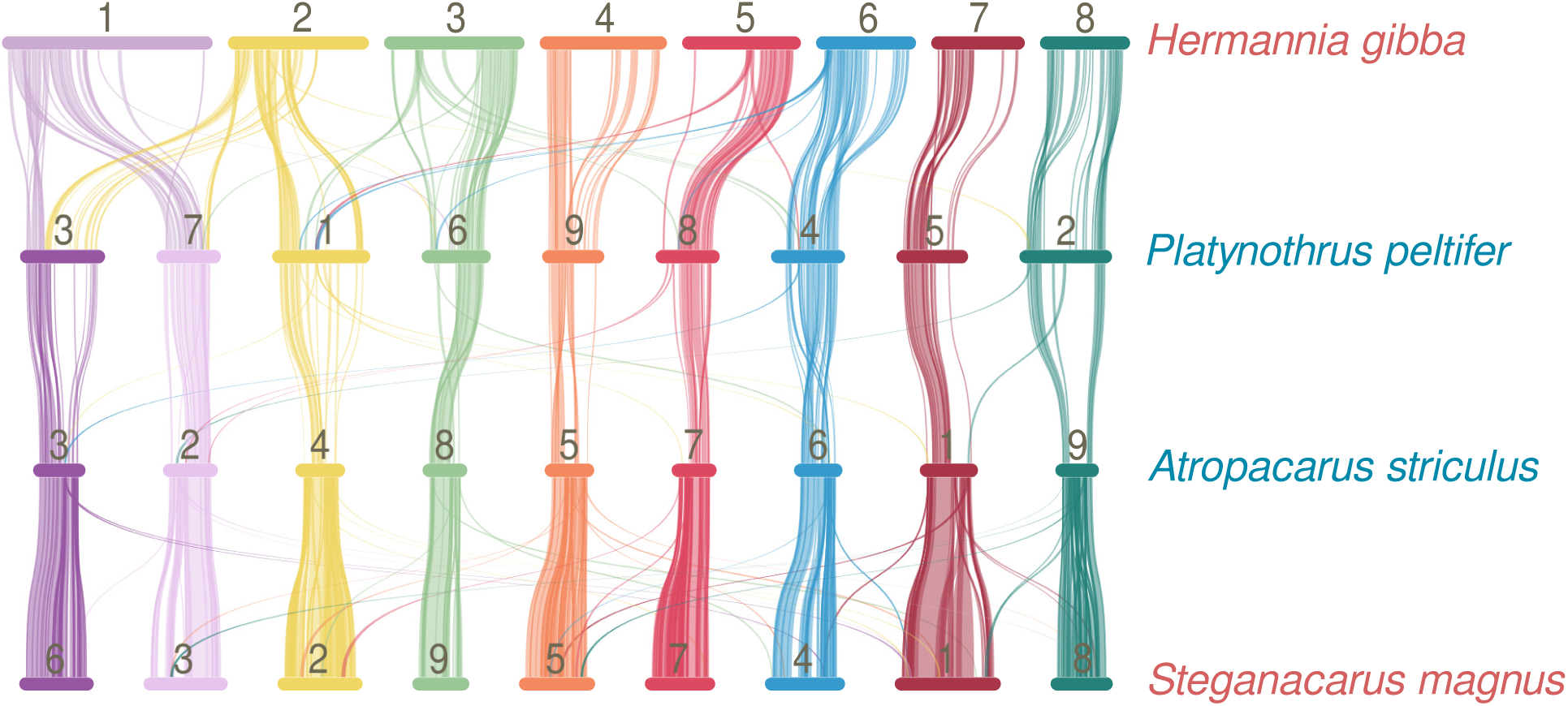
Conservation of nine chromosomes in oribatid mites. The macrosynteny of the chromosome-level assemblies of *Hermannia gibba* (Hga), *Platynothrus peltifer* (Ppr), *Atropacarus striculus* (Ass), *Steganacarus magnus* (Sms) (top to bottom) shows matching blocks of collinearity based on protein alignments. Chromosome colors highlight the main nine chromosomes found in *P. peltifer*, *A. striculus*, *S. magnus*, and partially rearranged in *H. gibba*. Names of sexual and asexual species are respectively in red and blue. For comparison arbitrary *H. gibba* chromosome numbers are given.

**Table 1:** Sets of sexual species and asexual relatives included in this study, and their assembly statistics. The divergence between *N. palustris* and *H. gibba* has been estimated to 385 millions years, and between *A. striculus* and *S. magnus* to 17 millions years.

| Species | Reproductive mode | Assembly size | N50 |
| --- | --- | --- | --- |
| *Hydrozetes lacustris* | asexual | 100.6 Mb | 5.0 Mb |
| *Hydrozetes thienemanni* | sexual | 272.0 Mb | 1.6 Mb |
| *Achipteria coleoptrata* | sexual | 145.3 Mb | 1.7 Mb |
| *Oppiella nova* | asexual | 123.7 Mb | 3.7 Mb |
| *Oppiella subpectinata* | sexual | 158.6 Mb | 1.5 Mb |
| *Nothrus palustris* | asexual | 254.7 Mb | 6.9 Mb |
| *Platynothrus peltifer* | asexual | 216.0 Mb | 23.4 Mb |
| *Hermannia gibba* | sexual | 352.2 Mb | 43.9 Mb |
| *Atropacarus striculus* | asexual | 138.7 Mb | 13.9 Mb |
| *Steganacarus magnus* | sexual | 261.0 Mb | 24.1 Mb |
| *Hypochthonius rufulus* | asexual | 184.7 Mb | 4.1 Mb |

Strikingly, sexual and asexual pairs show large variation in genome sizes, with larger genomes in sexual species, and smaller genomes in asexual species: from a ratio of 1.3 between *Oppiella subptectinata* (158.6 Mb) and *Oppiella nova* (123.7 Mb), to 2.7 between *Hydrozetes thienemanni* and *Hydrozetes lacustris* (Table 1). These differences were evaluated using a Phylogenetic Generalized Least Square test (PGLS), which accounts for the phylogenetic relation of the observed values, and were deemed significant (p-value = 0.014, multiple R^2^ = 0.510, phylogenetic signal = 0.804) with sexual genomes on average 97.2 Mb longer than asexual genomes. Nine chromosomes are conserved among *Platynothrus peltifer*, *Atropacarus striculus*, *Steganacarus magnus*. *H. gibba* has eight chromosomes due to the rearrangement of three chromosomes into two. The largely conserved chromosome numbers between these two pairs of asexual and sexual species, in addition to the smaller genome sizes of all parthenogens, suggest the absence of a polyploidization event in oribatid mites, consistent with predictions of diploidy (Supplementary Figure S1).

### 2.2 Non-categorical expansion of gene and TE content in sexual species

Differences in genome sizes suggest genome expansion in sexual species or genome compaction in asexual species, thus we searched for features associated with these variations by looking into genes and TEs. Gene numbers vary from 19,693 (*H. lacustris*) to 28,881 (*S. magnus*) (Figure 2), with significantly higher values in sexual species (PGLS test, p-value = 0.039, R^2^ = 0.394, phylogenetic signal = 0) which exhibit on average an extra 4,173 gene count. The highest difference is observed between *H. thienemanni* v. *H. lacustris* and *S. magnus* v. *A. striculus*, with a ratio of 1.4. For *H. lacustris*, *O. nova* and *A. striculus*, these reduced gene contents are associated with accrued ratios of gene loss over gene gain compared to their sexual counterpart (*H. lacustris* 2.4 v. *H. thienemanni* 0.6, *O. nova* 2.1 v. *O. subpectinata* 1.1, *A. striculus* 2.8 v. *S. magnus* 0.8). Discrepancies in gene content are in contrast with stable numbers of BUSCO orthologs across all species (Supplementary Table S1), both in single copy or duplicated, suggesting that core genes are not affected. Consequently, we investigated the conservation of Hierarchical Orthologous Groups (HOGs), which refer to genes classified along the phylogenetic tree based on their shared evolutionary history. From a total of 27,639 HOGs, most asexual species have retained less HOGs than their closest sexual sister species (*A. striculus*, *H. lacustris*, *O. nova*), with the exception of *N. palustris* and *P. peltifer* which retained more HOGs than *H. gibba* (PGLS test, p-value = 0.040, R^2^ = 0.391, phylogenetic signal = 0.979). We additionally searched whether reduced gene numbers in asexual species impacted preferentially categories associated with meiosis and sex determination; we identified several genes involved in these functions, but their copy numbers did not vary with reproductive modes (Supplementary Table S4). Functional enrichment analyses of gained and lost genes did not highlight specific categories (Supplementary Tables S5-6).

**Figure 2:**
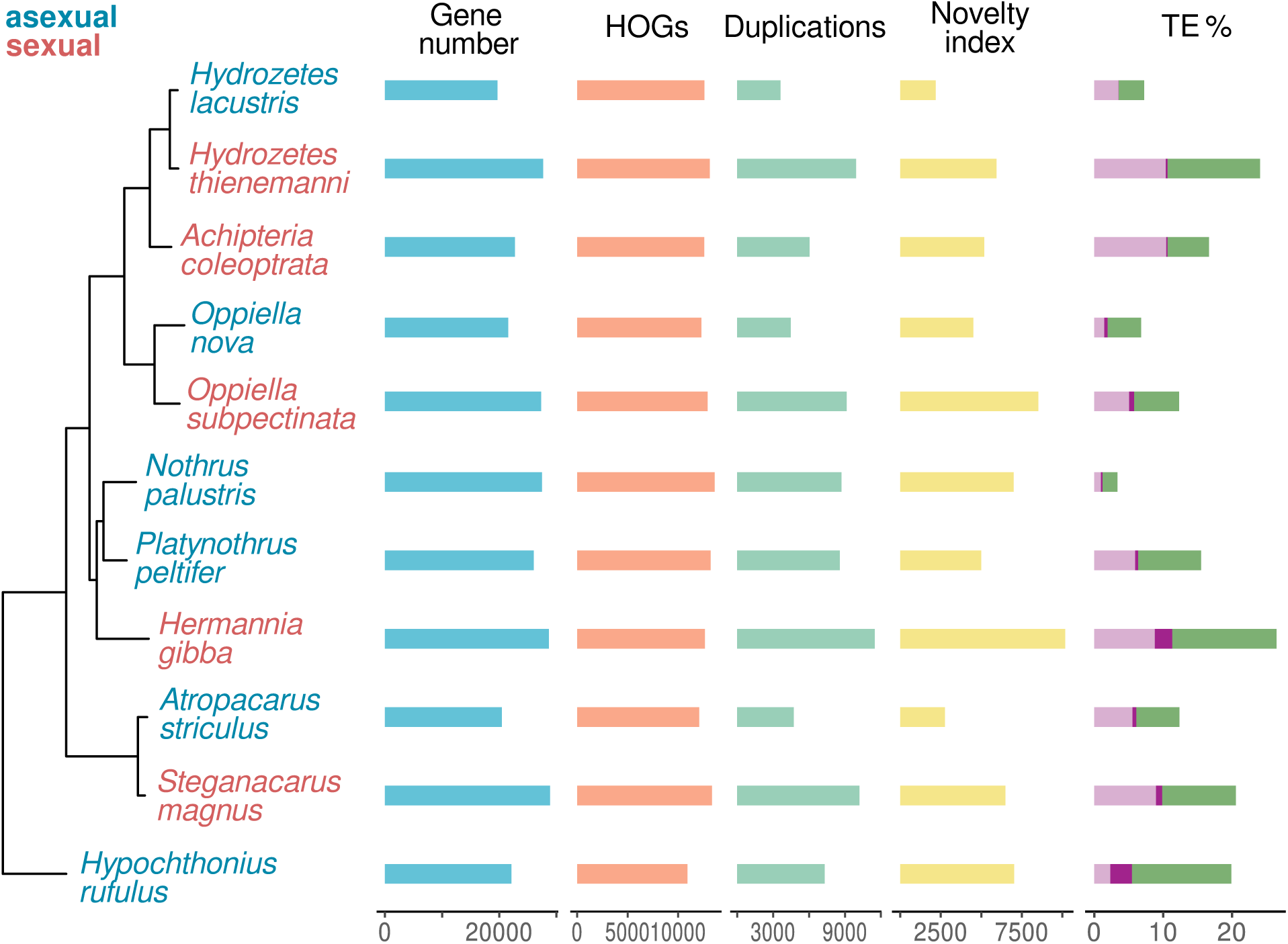
Large variations in genomic content between sexual and asexual species. Sexual species have increased numbers of overall genes, duplications, novelty index and TEs, compared to asexual species. From left to right: annotated phylogeny with gene gain and loss depicting asexual species in blue and sexual species in red, total number of genes (blue), number of retained HOGs (orange), number of duplication events (light green), the novelty index which includes unassigned genes, species-specific orthologs and TE genes (yellow), transposable elements classified as RNA elements (light purple), Helitrons (dark purple) and DNA elements (medium green).

Increased gene numbers are associated with more numerous duplications in sexual species (Figure 2; PGLS test, p-value = 0.004, R^2^ = 0.624, phylogenetic signal = 0.752) and an increased novelty index (PGLS test, p-value *<* 0.001, multiple R^2^ = 0.842, phylogenetic signal = 1). Sexual species have about 3,967 additional duplication events and a novelty index increased by 3,560 genes. Species-specific genes are the main contributors to these increased novelty indexes. We searched for HGTs, as a potential source of acquired innovation, and found moderate rates ranging from 1.19% (*H. thienemanni*) to 2.17% (*P. peltifer*) (Supplementary Table S1). The asexual species *P. peltifer*, *N. palustris*, *O. nova*, *H. lacustris* have slightly higher proportions of HGTs than their sexual sister species, but there is no significant difference in HGT content between sexual and asexual species (PGLS test, p-value = 0.414, multiple R^2^ = 0.075, phylogenetic signal = 0). We further looked into gene structure and found a maintained average exon length between 287 and 367 bp (Supplementary Table S1). Average intron length is systematically higher in sexual species, however these values are most likely driven by shared ancestry rather than reproductive mode (PGLS test, p-value = 0.141, R^2^ = 0.226, phylogenetic signal = 1; Supplementary Table S1).

Transposable element contents vary drastically within each set: proportions range between 2.2% (*N. palustris*) and 15.1% (*H. gibba*) for DNA elements, 1.1% (*N. palustris*) and 10.4% (*A. coleoptrata*) for RNA elements, and 0.0% (*H. lacustris*) to 3.1% (*H. rufulus*) for Helitrons. All asexual species have an overall lower TE load than their sexual counterpart (PGLS test, p-value = 0.011, R^2^ = 0.478, phylogenetic signal = 0.663), and this applies to DNA elements, RNA elements, and Helitrons (Figure 2). Detailed proportions of transposable elements per category are shown in Supplementary Table S3.

### 2.3 Selection remains as strong in asexual as in sexual mites

The differences in gene and TE content between asexual and sexual species may suggest variations in purifying selection between reproductive modes. Typically, as a consequence of meiotic recombination and segregation, the efficacy of natural selection is expected to be higher in sexual species compared to asexually reproducing organisms [Otto, 2021, Felsenstein, 1974]. Previously, asexual mites were found to feature more effective purifying selection than their sexual counterparts, which could factor into the reduction of parthenogen genomes. However, this inference was based on transcriptomic data from a limited set of six, sometimes relatively distant, species [Brandt et al., 2017]. Here we revisit these findings with the more robust comparative framework of 11 sexual and asexual oribatid mite species. The branch-wide rate of omega (*ω* ; the ratio of non-synonymous to synonymous substitutions (dN/dS)) was computed using 2,890 single-copy orthologous protein coding genes across the 11 mite species. Estimates of *ω* were on average reduced across the different species (median = 0.1), suggesting a strong overall purifying selection (Figure 2A). When compared to their sexual sister species, most asexual mite species exhibited lower *ω* values (e.g., *O. subpectinata* and *O. nova*; *H. gibba* and *P. peltifer*) or similar (e.g., *H. lacustris* and *H. thienemanni*) (Figure 2A; Kruskal–Wallis rank sum test, X2 = 2387.8, df = 10, p-value *<* 0.001; pairwise Wilcoxon rank sum *post hoc* tests are displayed in the figure). This suggests that purifying selection remains equally as effective in asexual species as in sexual species.

Similarly, evolutionary rate analyses along all the sexual and asexual mite lineages revealed no significant differences between the two sets in terms of positive selection (*ω* positive *≥* 1, Figure 2B top) and purifying selection (*ω* purifying *<* 1, Figure 2B bottom). The selection intensity parameter (k), which measures relaxation (k *<* 1) or intensification (k *>* 1) of natural selection between two sets of branches, showed that most genes had a k value of 1 (FDR *>* 0.05, Figure 2C). This indicates that, overall, natural selection was neither intensified nor relaxed along the asexual branches of oribatid mites. However, our analysis revealed a subset of 87 single-copy orthologs associated with k significantly different from 1 (at FDR *<* 0.05), 64 of them showing signatures of relaxed selection in the asexual branches. Gene Ontology (GO) enrichment analyzes of these genes revealed that terms putatively involved in recombination (e.g., reciprocal meiotic recombination (GO:0007131), Holliday junction resolution (GO:0000400), and double-strand break repair (GO:0045003)) were enriched (Supplementary Table S7; p-values *<* 0.05; FDR *>* 0.05).

### 2.4 No difference in the SV ”turnover” rate between sexual and asexual species

Asexuality is often associated with the loss of meiotic recombination, and it has been suggested that in parthenogenetic oribatid mites meiosis would only occur in telomeric regions [Taberly, 1987]. The relaxation of the constraints imposed by segregation and the alignment of chromatids is expected to facilitate the emergence of structural variants, which include insertions, duplications, deletions, transpositions and inversions. In addition, insertions and deletions could factor in genome size variability in asexual and sexual species, thus we searched for common SVs. Insertions were the most frequent mutations among the 11 species, followed by duplications and deletions, while transpositions and inversions occurred at lower frequencies (Figure 4A; Kruskal–Wallis rank sum test, X2 = 89.441, df = 4, p-value *<* 0.001; pairwise Wilcoxon rank sum *post hoc* tests are displayed in the figure). The number and size of each mutation type varied among the analyzed genomes and consisted largely of small mutation events (*<* 100 bp) (Supplementary Figure S14). Mutation frequencies, expressed as percent increase or decrease compared to the average rate, were not significantly enriched in asexual compared to sexual branches (Figure 4B). However, mutations were on average longer in the asexual mites compared to sexual sister species (e.g., *H. lacustris* v. *H. thienemanni* and *A. striculus* v. *S. magnus*) (Figure 4C; Kruskal–Wallis rank sum test, X2 = 390,890, df = 10, p-value *<* 0.001; pairwise Wilcoxon rank sum *post hoc* tests are displayed in the figure). When analyzed separately, insertions did not correlate with reproductive mode, but deletion events showed a subtle increase in asexual compared to sexual mites (Supplementary Figure S15).

**Figure 3:**
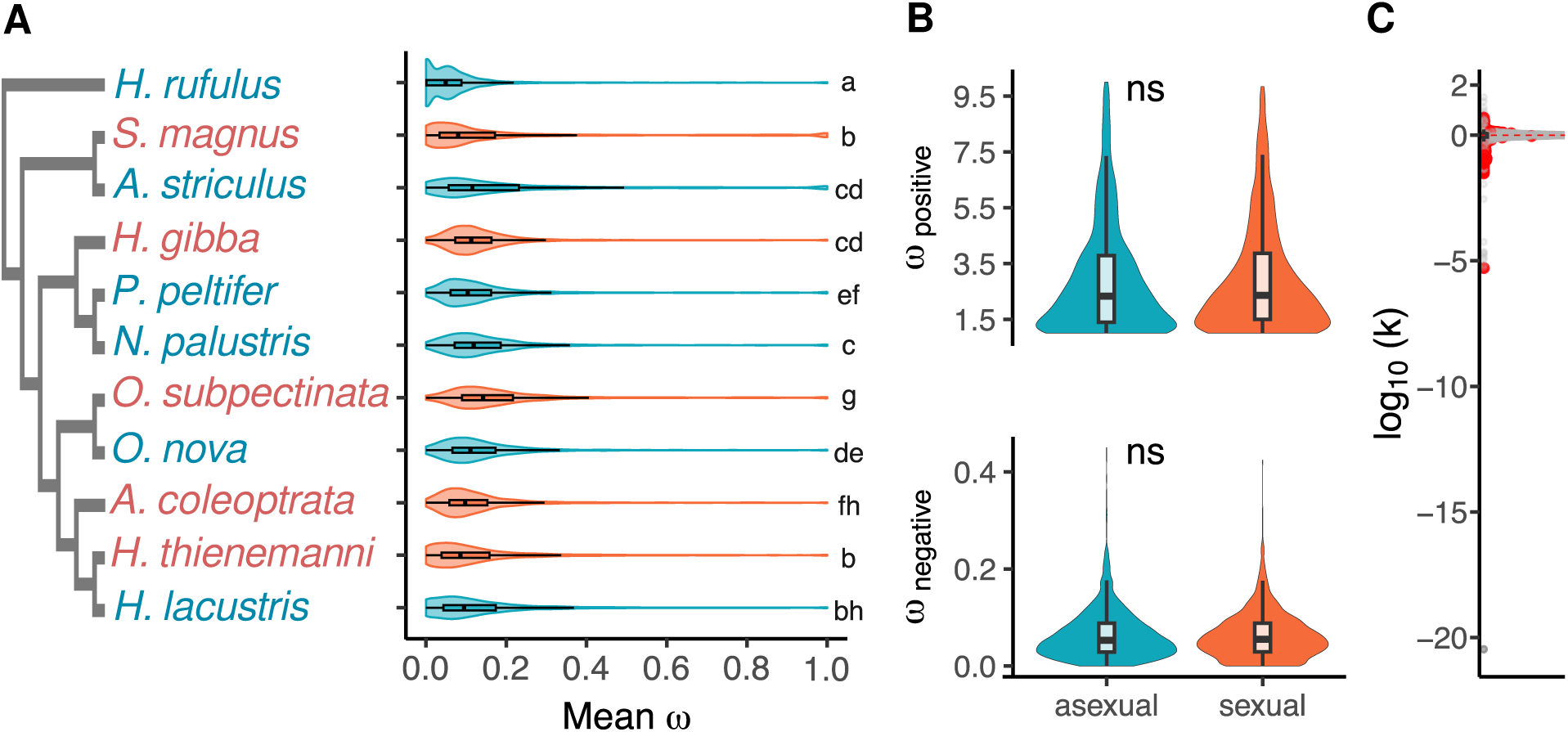
Similar levels of purifying selection across oribatid mites. (A) Distribution of mean *ω* values (ratios of non-synonymous to synonymous substitutions dN/dS) in terminal branches of eleven oribatid mites, using 2,890 single-copy orthologs. Different letters represent significant differences according to pairwise Wilcoxon rank sum *post hoc* tests. (B) Distribution of evolutionary rates in 1,740 single-copy orthologs under positive selection (dN/dS *≥* 1, top) and purifying selection (dN/dS *<* 1, bottom). (C) Distribution of the selection intensity parameter *k* for single-copy orthologs in asexual mites where positive values indicate an intensification of selection, while negative values suggest a relaxation of natural selection in asexual relative to sexual mite lineages. The *k* values were not significantly different from zero for the majority of the investigated genes (gray dots), whereas a small proportion showed *k* values significantly different from 1 (87 genes; red dots).

**Figure 4:**
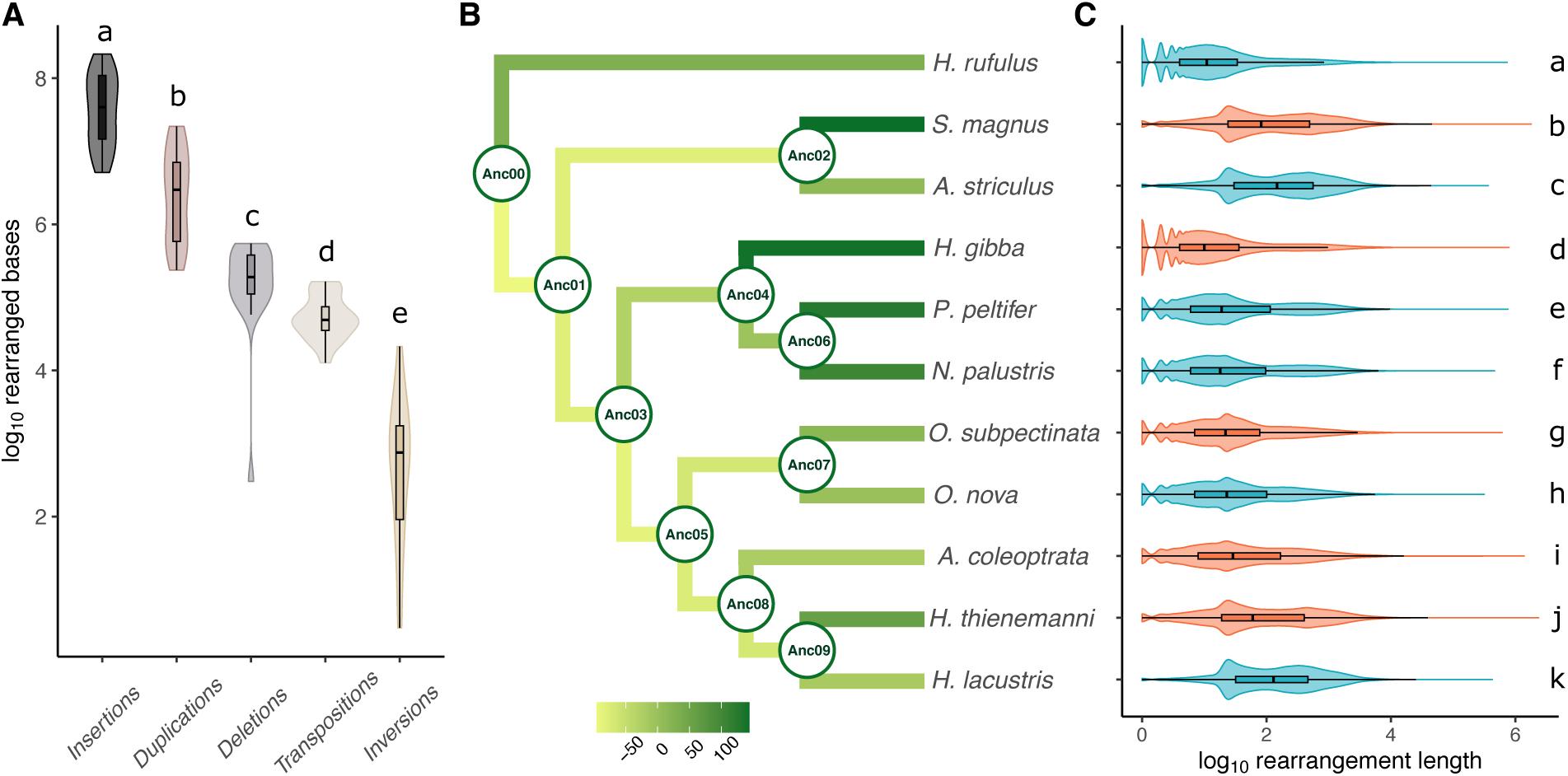
No significant enrichment of structural variants in asexual oribatid mites. (A) Distributions of different types of structural variations shown as proportion of affected base pairs (Y-axis). (B) Relative enrichment of mutational events shown as percent deviation from the mean across all branches in the phylogeny. (C) Rearrangement length distribution (in base pairs) in the different mite genomes. Different letters represent significant differences according to pairwise Wilcoxon rank sum *post hoc* tests.

## 3 Discussion

The persistence and diversification of ancient asexual lineages remain a central puzzle in evolutionary biology, given the theoretical challenges associated with obligate parthenogenesis. In this study, we investigated four sets of sexual and asexual oribatid mite species, with multiple transitions to asexuality estimated to have occurred tens of millions of years ago, and as far back as 200 million years ago [Hammer and Wallwork, 1979, Schaefer et al., 2010].

Pairs of sexual and asexual species compared in this study were assembled with comparable methods. Although not all genomes could be scaffolded to chromosome level (but at least to Megabase-level contiguity, in line with assembly standards for non-model species [Guiglielmoni et al., 2022]), our measures of variations in genomic content appear independent of contiguity. We found that sexual and asexual genomes follow distinct trajectories. Sexual species have larger genomes, due to higher gene content through duplications and emergence of novel genes, and higher TE content. In contrast, the comparatively smaller genomes of asexual species are characterized by low overall TE content and lower numbers of novel genes.

### 3.1 Large-scale chromosome structures are conserved across millions of years

Some soil-dwelling asexual species, such as the springtail *Folsomia candida*, have been found to exhibit extensive intragenomic collinearity blocks, palindromic regions and rearrangements, and this has been linked to genome evolution under parthenogenesis [Faddeeva-Vakhrusheva et al., 2017]. Our chromosome-level assemblies of *P. peltifer*, *H. gibba*, *A. striculus* and *S. magnus* show a remarkable conservation of chromosome structure across hundreds of millions of years of divergence in asexual oribatid mites. This conservation is also in contrast with multiple observations of loss of chromosome architecture over long evolutionary time [Lewin et al., 2024]. The differences between the springtails and oribatid mites, tiny soil organisms sharing similar habitats, might indicate that not ecological pressures, but evolutionary age has shaped genome architecture in these asexual species. It is possible that some asexual lineages maintain chromosomal integrity despite the absence of recombination and this allows them to persist in time.

### 3.2 No difference in effective selection and rate of SVs between reproduction modes

Contrary to the prediction that asexuality should lead to less effective selection, and consistent with previous results [Brandt et al., 2017], no difference in the effectiveness of purifying selection or positive selection could be detected. This suggests that asexual oribatid mites might avoid harmful mutations and adapt to changing conditions equally well as sexual relatives. Interestingly, gene enrichment terms that are under relaxed selection in asexual species include terms related to recombination. This is consistent with previous findings, that recombination likely only is maintained at the chromosome ends [Öztoprak et al., 2025]. We found no consistent increase in horizontally transferred genes in asexual lineages, which suggests that these previous reports of high HGT content [Flot et al., 2013, Debortoli et al., 2016] may rather be linked to the cryptobiotic ability of the studied bdelloid rotifers. It has been predicted that asexual genomes should be able to sustain more SVs compared to sexual relatives, as chromosomes do not need to align with the same precision. Here, no overall difference was found in the rate of SV turnover between reproductive modes. However, asexual species maintain longer SVs, consistent with predictions. By extension, HGTs were elevated in asexual species in some comparisons to sexual relatives, but not consistently and not by large effect sizes. This suggests that previous reports of massive HGT contents in asexual species are rather a consequence of other lineage specific dynamics than reproductive mode.

### 3.3 Asexual mites are not able to retain novel genes to the same degree as sexual species

Consistent with cytological observations of meiosis [Taberly, 1987], we find meiosis-related genes are retained in asexual mites. While TE identification can be impacted by assembly contiguity, our use of long reads delivered assemblies with a minimum N50 of 1.5 Mb, a contiguity far exceeding previous efforts [Collins et al., 2023]. Through this we are able to confirm a genomic dichotomy between sexual and asexual oribatid mites across all categories of TEs, with a low overall TE content in the asexual species. This is in stark contrast to recently evolved parthenogens such as *Timema* stick insects, which display no TE accumulation [Jaron et al., 2022].

Novel functions arise by duplication and divergence of genes and this creation of novelty is a driver of evolution. We found that the acquisition of species-specific novel genes is a major contributor to genome growth in sexual mite species. Arachnida-lineage orthologs were largely not affected by duplications, as would be expected for genes with core functions. The conservation of these *de novo* evolved genes among asexual species also does not support the hypothesis that lower gene numbers in parthenogens would result from gene loss. Specifically, the most contrasting overall gene contents in our dataset are observed between the youngest species pairs, *Hydrozetes thienemanni* v. *Hydrozetes lacustris*, and *S. magnus* v. *A. striculus*, with an estimated divergence 17.1 millions years ago [Dabert et al., 2010]. *N. palustris*, *P. peltifer* and *H. gibba*, with an estimated divergence time of 385 millions years [Schaefer et al., 2010, Lozano-Fernandez et al., 2020], have more similar gene content, but still sharp variations in TE content. In other words, diverging trajectories in genome architecture may be less pronounced over time between sexual and asexual species. This is logical, as evolutionary young species will create novelty by gene expansion, for example during adaptation to a new environment, before a more static phase of genome evolution is reached [Vargas-Chávez et al., 2025]. However, it appears that even relatively young asexual species in our dataset are not able to retain large number of novel genes. Again this appears logical under the assumption that most larger mutations, i.e. most newly evolved genes or duplicated genes, are deleterious. Asexual species, lacking outcrossing, will not be able to buffer against these comparatively large deleterious mutations, and individuals carrying these are thus quickly purged from the population. Notably, this process would be uncoupled from genomic decay through mutation accumulation (Muller’s ratchet), for which we do not find evidence in the ancient asexual lineages and is also in line with the lack of differences in purifying selection between reproductive modes.

### 3.4 Long-term asexuality appears linked to genome streamlining

Our findings in oribatid mites, an ancient asexual system with multiple independent transitions, provide a critical addition to the ongoing debate about the fate of genomes under parthenogenesis. In summary, we find that in oribatid mites, sex is associated with genome expansion and the evolution of novelty, whereas successful long-term asexuality in these species appears linked to a streamlined genome architecture over tens to hundreds of millions years. This is in contrast to evolutionary young asexual species, like *Timema* stick insects, which show clear signs of genomic decay, like extensive loss of heterozygosity and reduced adaptive potential [Jaron et al., 2022]. Our results highlight that the genomic consequences of asexuality are not universally degenerative but can follow distinct trajectories depending on evolutionary timescale and lineage-specific constraints. Coupled with previous evidence showing effective purifying selection in these mites [Brandt et al., 2017], our findings suggest that these asexual lineages have found a viable, long-term evolutionary pathway centered on a streamlined genome architecture. At the same time, this implies that asexual mites have traded off some of their ability to create evolutionary novelty through gene duplication. Thus, they might only be able to survive in relatively stable soil habitats, making them particularly vulnerable in times when global change negatively impacts these environments.

## 4 Material & Methods

### 4.1 Sample collection

Mite individuals were obtained from the following locations: *Achipteria coleoptrata* from Germany, Dahlem Moorpfad, 26/04/2023, November 2022; *Atropacarus striculus*, *Hypochthonius rufulus*, *Steganacarus magnus*, *Oppiella subpectinata* from Germany, Dahlem Moorpfad, 26/04/2023; *Hydrozetes lacustris* from Norway, Gimsøya, 68.314151, 14.155578, 28/07/2021; *Hydrozetes thienemanni* from Poland, Pruszcz Bagienica, 53.265750, 17.531062, 24/10/2021; *Oppiella nova* from a lab culture. Prior to library preparations, individuals were cleaned as described in [Öztoprak and Bast, 2023].

### 4.2 High-molecular-weight DNA sequencing

For *Atropacarus striculus*, *Achipteria coleoptrata*, *Hypochthonius rufulus*, *Nothrus palustris*, *Steganacarus magnus*, high-molecular-weight DNA was extracted from a single individual per species following [Öztoprak and Bast, 2023, Öztoprak et al., 2025]. Ultra-low input PacBio HiFi libraries were prepared at the Genomics & Transcriptomics lab (GTL; Düsseldorf, Germany) with the Express 2.0 Template kit (Pacific Biosciences, Menlo Park, CA, USA) and sequenced on a Sequel II/Sequel IIe instrument with 30 hours movie time. From the same extract, a TELL-seq library was prepared for *Nothrus palustris* using a TELL-Seq WGS Library Prep Kit (Universal Sequencing Technology, Carlsbad, CA) and sequenced on an Illumina NovaSeq 6000 at the Cologne Center for Genomics (CCG; Cologne, Germany). PacBio HiFi reads were generated using SMRT Link (v10, Pacific Biosciences, Menlo Park, CA, USA) with default parameters. For *Oppiella* and *Hydrozetes* species, single mite individuals were picked from ethanol and left to air dry on a sterile glass dish before being transferred to 55 µL ATL buffer (Qiagen, Germany), in which they were crushed completely with a pestle. DNA extractions and purifications followed the protocol for the MagAttract HMW DNA kit from Qiagen using quarter volumes; DNA was eluted in 80 µL AE buffer. DNA extracts were quantified using a Quantus Fluorometer and the corresponding QuantiFluor dsDNA System (Promega, USA). Fragment lengths were visualized using a Femto Pulse System and the Genomic DNA 165 kb Kit (Agilent, USA). PacBio HiFi library preparations followed a modified version of the protocol for preparing HiFi libraries from ultra-low DNA input [Schardt and Bálint, 2024] using the Express 2.0 Template kit (Pacific Biosciences, USA).

### 4.3 Hi-C sequencing

Pools of mites were crosslinked in 3% formaldehyde for 1 hour with low agitation, and quenched in 250 mM glycine for 30 minutes. The samples were flash frozen in liquid nitrogen and crushed with a pestle, then prepared with the Arima Hi-C+ kit following the animal protocol from the lysis step. The library was sequenced on an Illumina NovaSeq 6000 at the CCG.

### 4.4 RNA sequencing

RNA was extracted from pools of individuals using the Direct-zolTM RNA MicroPrep kit (Zymo Research, Irvine, CA) with DNase I-treated TRIzol. RNA-seq libraries were prepared using the TruSeq Stranded Total RNA with Ribo-Zero Globin protocol and sequenced on an Illumina NovaSeq 6000 at the CCG.

### 4.5 Nuclear genome assembly

Ploidy was estimated using KMC v3.2.1 [Kokot et al., 2017] and Smudgeplot v0.2.2 [Ranallo-Benavidez et al., 2020] with the PacBio HiFi reads. For all species, PacBio HiFi reads were first assembled using Flye v2.9 [Kolmogorov et al., 2019] with the parameter --pacbio-hifi. For *N. palustris*, *H. lacustris*, *H. thienemanni*, *O. nova*, *O. subpectinata*, *A. coleoptrata*, *H. rufulus*, a second assembly was produced using hifiasm v0.16 [Cheng et al., 2021] with parameter -l 2.

#### 4.5.1 Nothrus palustris

PacBio HiFi reads were mapped to the Flye and hifiasm assemblies using minimap2 v2.24 [Li, 2018] with parameter `-x map-hifi`, and the output was used to remove artefactual duplications using purge dups v2.1.5 [Guan et al., 2020] with thresholds `-l 5 -m 105 -u 350`. Both assemblies were scaffolded using Scaff10X v4.2 [High Performance Algorithms group, Wellcome Sanger Institute, 2018] with the modules scaff reads and scaff10X (parameters `-longread 1 -gap 10 -matrix 2000 -reads 10 -score 10 -edge 50000 -link 8 -block 50000`).

The post-processed Flye assembly was further scaffolded using RagTag v2.1.0 [Alonge et al., 2022] and the processed hifiasm assembly as reference. Gaps in the scaffolds were filled using TGS-GapCloser v1.2.1 [Xu et al., 2020] and the PacBio HiFi reads with parameters `--ne --tgstype pb --minmap arg ’ -x map-hifi’`. PacBio HiFi reads were mapped again against the assembly using minimap2 v2.24 with parameter `-ax map-hifi`; the output was sorted using SAMtools v1.6 and provided as input to HyPo v1.0.3 [Kundu et al., 2019], which was run with parameters `-s 250m -c 130 -k ccs`.

#### 4.5.2 Hydrozetes lacustris and Hydrozetes thienemanni

PacBio HiFi reads were mapped to the Flye and hifiasm assemblies using minimap2 v2.24 [Li, 2018] with parameter `-x map-hifi`, and the output was used to remove artefactual duplications using purge dups v2.1.5 [Guan et al., 2020] with thresholds `-l 50 -m 150 -u 500` for *H. lacustris* and `-l 10 -m 65 -u 220` for *H. thienemanni*. The purged Flye assembly was further scaffolded using RagTag v2.1.0 [Alonge et al., 2022] and the purged hifiasm assembly as reference. PacBio subreads were used to scaffold the assembly with LINKS v2.0.1 [Warren et al., 2015] and to fill gaps with TGS-GapCloser v1.2.1 [Xu et al., 2020] (parameter `--ne` *–tgstype pb*). PacBio HiFi reads were mapped to the final assemblies with minimap2 v2.24 with parameter -ax map-hifi, sorted using SAMtools v1.6, and used for polishing using HyPo v1.0.3 [Kundu et al., 2019] with parameters `-s 100m -c 200 -k` ccs for *H. lacustris* and `-s 250m -c 80 -k ccs` for *H. thienemanni*.

#### 4.5.3 Oppiella nova and Oppiella subpectinata

PacBio HiFi reads were mapped to the Flye and hifiasm assemblies using minimap2 v2.24 [Li, 2018] with parameter `-x map-hifi`, and the output was used to remove artefactual duplications using purge dups v2.1.5 [Guan et al., 2020] with thresholds `-l 5 -m 155 -u 500` for *O. nova* and `-l 5 -m 100 -u 700` for O. subpectinata. The purged Flye assembly was further scaffolded using RagTag v2.1.0 [Alonge et al., 2022] and the purged hifi-asm assembly as reference. Gaps were filled using TGS-GapCloser v1.2.1 [Xu et al., 2020] and the PacBio HiFi reads with parameters `--ne --tgstype pb --minmap arg ’ -x map-hifi’`. PacBio HiFi reads were mapped against the *O. nova* assembly using minimap2 v2.24 with parameter `-ax map-hifi`, sorted using SAMtools v1.6, and provided as input to HyPo v1.0.3 [Kundu et al., 2019] with PacBio HiFi reads and parameters `-s 120m -c 200 -k ccs`.

#### 4.5.4 Atropacarus striculus and Steganacarus magnus

PacBio HiFi reads were mapped to the Flye assembly using minimap2 v2.24 [Li, 2018] with parameter `-x map-hifi`, and the output was used to remove artefactual duplications using purge dups v2.1.5 [Guan et al., 2020] with thresholds `-l 5 -m 170 -u 900` for *A. striculus* and `-l 3 -m 130 -u 700` for *S. magnus*. Hi-C reads were trimmed using TrimGalore v0.6.10 [Krueger et al., 2023], which combines cutadapt v1.18 [Martin, 2011] and FastQC v0.12.1 [Babraham Bioinformatics, 2010], and pre-processed using BWA 0.7.17 [Li and Durbin, 2009] and hicstuff v3.1.6 [Matthey-Doret et al., 2020] with parameters `-e DpnII,HinfI -m iterative -a bwa`. The output and the purged assembly were provided as input to instaGRAAL v0.1.6 no-opengl [Baudry et al., 2020] for scaffolding with parameters `-l 4 -n 50 -c 1 -N 5` for *A. striculus* and `-l 5 -n 50 -c 1 -N 5` for *S. magnus*. The scaffolds were automatically curated using the module instagraal-polish with parameters `-m polishing -j NNNNNNNNNN`. Gap filling was performed using TGS-Gapcloser v1.2.1 [Xu et al., 2020] and the PacBio HiFi reads with parameters `--ne --tgstype pb --minmap arg ’ -x map-hifi’`. PacBio HiFi reads were mapped against the assembly again using minimap2 v2.24 with parameter `-ax map-hifi`, sorted using SAMtools v1.6, and provided as input to HyPo v1.0.3 [Kundu et al., 2019].

#### 4.5.5 Achipteria coleoptrata

PacBio HiFi reads were mapped to the Flye and hifiasm assemblies using minimap2 v2.24 [Li, 2018] with parameter `-x map-hifi`, and the output was used to remove artefactual duplications using purge dups v2.1.5 [Guan et al., 2020] with thresholds `-l 35 -m 220 -u 700`. The purged Flye assembly was further scaffolded using RagTag v2.1.0 [Alonge et al., 2022] and the purged hifiasm assembly as reference.

#### 4.5.6 Hypochthonis rufulus

PacBio HiFi reads were mapped to the Flye and hifiasm assemblies using minimap2 v2.24 [Li, 2018] with parameter `-x map-hifi`, and the output was used to remove artefactual duplications using purge dups v2.1.5 [Guan et al., 2020] with thresholds `-l 5 -m 150 -u 800`. Both assemblies were scaffolded using Scaff10X v4.2 [High Performance Algorithms group, Wellcome Sanger Institute, 2018] with the modules scaff reads and scaff10X (parameters `-longread 1 -gap 10 -matrix 2000 -reads 10 -score 10 -edge 50000 -link 8 -block 50000`). The post-processed Flye assembly was further scaffolded using RagTag v2.1.0 [Alonge et al., 2022] and the processed hifiasm assembly as reference. Gaps in the scaffolds were filled using TGS-GapCloser v1.2.1 [Xu et al., 2020] and the PacBio HiFi reads with parameters `--ne --tgstype pb --minmap arg ’ -x map-hifi’`. PacBio HiFi reads were mapped against the assembly again using minimap2 v2.24 with parameter `-ax map-hifi`; the output was sorted using SAMtools v1.6 and provided as input to HyPo v1.0.3 [Kundu et al., 2019], which was run with parameters `-s 180m -c 200 -k ccs`.

#### 4.5.7 Assembly evaluation

Assembly statistics were calculated using assembly-stats v1.0.0 [Pathogen Informatics, Wellcome Sanger Institute, 2014]. *k*-mer completeness was computed using KAT v2.4.2 [Mapleson et al., 2016] and the module kat comp with *k* =27 and the PacBio HiFi reads. Ortholog completeness was evaluated using the tool Benchmarking Universal Single-Copy Orthologs (BUSCO) v.5.4.7 [Manni et al., 2021] against the Metazoa odb10 lineage (954 orthologs) and the Arachnida odb10 lineage (2,934 orthologs). Assemblies were aligned against the nucleotide database using the Basic Local Alignment Search Tool (BLAST) v2.10.1 [Altschul et al., 1990] with parameters `-outfmt \6 qseqid staxids bitscore std sscinames scomnames” -max hsps 1 -evalue 1e-25`. PacBio HiFi reads were mapped using minimap2 v2.24 [Li, 2018] with parameters `-ax map-hifi` and the output was sorted using SAMtools v1.6 [Danecek et al., 2021]. The outputs of minimap2, BLAST, and BUSCO (against the Arachnida odb10 lineage) were provided as input to BlobTools2 v4.3.2 [Challis et al., 2020]. Sequences identified as contaminants were subsequently removed. For *Atropacarus striculus* and *Steganacarus magnus*, Hi-C reads were mapped to the assembly using BWA v0.7.17 [Li and Durbin, 2009] and hicstuff v3.1.6 [Matthey-Doret et al., 2020] as previously described. Contact maps were generated using the module hicstuff view with the parameter `-b 500`.

### 4.6 Mitochondrial genome assembly

The module findMitoReference.py from MitoHiFi v3.2.1 [Uliano-Silva et al., 2023] was used to search for closely related mitochondrial genomes to use as a reference for each species. Contigs generated using Flye were provided as input to MitoHiFi [Uliano-Silva et al., 2023].

### 4.7 Repeat and gene annotations

The Extensive *De novo* TE Annotator (EDTA) pipeline v2.0.1 [Ou et al., 2019] was run with parameters `--sensitive 1 --anno 1 --force 1`. This program combines predictions from LTRharvest [Gremme et al., 2013], LTR FINDER [Xu and Wang, 2007] LTR retriever [Ou and Jiang, 2018], HelitronScanner [Xiong et al., 2014], Generic Repeat Finder [Shi and Liang, 2019], TIR-learner [Su et al., 2019] to yield a combined transposable element library using RepeatModeler [Flynn et al., 2020]. The hardmasked assembly was converted into a softmasked assembly. For species for which RNA-seq reads were available, the reads were trimmed using TrimGalore v0.6.10 [Krueger et al., 2023] and mapped to the assemblies using hisat2 v2.2.1 [Kim et al., 2019]. After sorting the mapped reads using SAMTools v1.6 [Danecek et al., 2021], the output was provided as input to BRAKER v3.0.3 [Gabriel et al., 2023], with parameters `--gff3 --UTR off`, which combined predictions from Augustus v3.5.0 [Stanke et al., 2008] and GeneMark-ES v4.68 [Lomsadze et al., 2005]. For species for which no RNAseq data were available, gene annotations from chromosome-level assemblies were provided as protein hints. The longest isoform was selected using Another Gtf/Gff Analysis Toolkit (AGAT) v0.8.0 [Dainat, 2020] with the script agat sp keep longest isoform.pl. The protein predictions were evaluated using BUSCO v5.4.7 [Manni et al., 2021] against the Metazoa odb10 and Arachnida odb10 lineages with the parameter `-m protein`.

Functional annotations were generated for each species’ gene predictions using InterProScan v5.69-101.0 [Jones et al., 2014]. TE genes were identified using the module DeTEnGA from GAQET2 [Garcia-Carpintero et al., 2025], with the functional annotation step overridden by providing the already generated output of InterProScan. Genes selected as associated with TEs were retained from the following categories: PteMte (TE protein domains + TE mRNA support), PteM0 (TE protein domains without TE mRNA), and PchMte (Mixed TE/non-TE domains with TE mRNA). TEs were classified using the FasTE pipeline [Bell et al., 2022] to superfamily level using the convolutional neural network program DeepTE [Yan et al., 2020]. Phylogenetic Generalized Least Square tests were conducted on different features of genomic content using the package caper [Orme et al., 2011] in R v4.4.2.

### 4.8 Synteny

The Multiple Collinearity Scan toolkit X (MCScanX) (commit b1ca533) [Wang et al., 2012] was used to detect gene synteny and collinearity between the chromosome-level genome assemblies of *Hermannia gibba*, *Platynothrus peltifer*, *Atropacarus striculus* and *Steganacarus magnus*. Proteomes were aligned one-to-one using the Basic Local Alignment Search Tool (BLAST) with the module blastp v2.15.0 [Altschul et al., 1990] and the parameters-evalue 1e-10 `-outfmt 6`. Alignment files and proteomes in modified GFF format were provided as input to MCScanX which was run with parameters `-k 250 -e 1e-15`. The colinearity output was plotted as a parallel, multi-genome tree graph using SynVisio (commit 3415935) [Bandi and Gutwin, 2020].

### 4.9 Horizontal gene transfers

The potential HGT candidates including the putative donor species were inferred using the HGT pipeline v1.0 from [Nowell, 2017]. The protein files for the analyzed mite species were in a first step aligned to the Uniref90 database [Suzek et al., 2015] (downloaded on 05.08.2024) using DIAMOND BLASTp v2.0.15 [Buchfink et al., 2021]. The parameters `--sensitive --index-chunks 1 -k 500 -e 1e-5` were used for the alignments. Taxid information was added to the resulting DIAMOND tables, before using the output for the diamond_to_HTG_candidates.pl script (required taxdump database was downloaded on 05.08.2024). The script was used with default thresholds for the HGT index (hU *≥* 30) and the CHS (*≥* 90%). Metazoa was set as the ingroup and Acari was set as the taxid to skip to avoid bias using the parameter `-k 6933`. The latter two options were also used for the HGT_candidates_to_fasta.pl script, while the maximum number of sequences to extract was set to 15. The get_locations_of_HGT_candidates.pl script was used with the default parameters to test for possible contaminations (scaffolds with *≥* 95% HGT genes). The script further tests for intron presence in the HGT candidate genes as well as for linkage to unambiguously metazoan genes, both of which are indicators of a robust HGT assessment [Nowell et al., 2018, Jaron et al., 2021]. The HGT candidates were then filtered for expressed genes as an additional step to avoid including genes originating from contamination. Stacked histograms, depicting the proportions of HGT candidates relative to the total gene count by their respective inferred domains for each species, were produced using R v4.3.3. The R script takes the HGT candidate files produced by the diamond_to_HGT_candidates.pl script as input. The R packages ’dplyr’, ’tidyr’ and ’ggplot2’ were used for the script.

### 4.10 Phylogenetic tree construction and whole-genome alignment

To build our phylogenetic tree, we first retrieved and generated a concatenated multiple sequence alignments for 2,899 single-copy orthologous protein-coding sequences across eleven mite species using OrthoFinder v2.5.5 [Emms and Kelly, 2019]. The resulting multiple sequence alignment was used to perform a maximum likelihood analysis using IQ-TREE v2.3.0 COVID-edition) [Minh et al., 2020] and 1,000 ultrafast bootstrap replicates to compute branch support. The best-fit substitution model was selected automatically using the built-in ModelFinder option in IQ-TREE. For genomic feature analysis, the phylogenetic tree was represented using ggtree [Yu, 2022] v3.14.0 and treeio v1.30.0 [Wang et al., 2020] in R V4.4.2 with *Hypochtonius rufulus* set as outgroup. To examine the impact of asexuality on genome dynamics, we explored the mutational landscape of eleven oribatid mite species using the phylogeny-aware whole-genome aligner Progressive Cactus v2.9.0 [Armstrong et al., 2020]. For this, the tree produced using IQ-TREE was used as a guide for the Progressive Cactus analysis with default settings. Next, halSummarizeMutations [Hickey et al., 2013] was used to extract inferred mutations at each branch of the guide phylogeny. Following a pipeline described previously [Schrader et al., 2021], we normalized transpositions, insertions, deletions, inversions, and duplications using branch length estimates as proxy for divergence to calculate mutation rates per substitution per site. We then grouped these rearrangements into bins based on the number of affected bases per branch across the phylogeny (Supporting Figure S14). To test for increased mutation rates at specific branches/nodes, we calculated and visualized mutation rates across branches for all mutation types (Figure 3B) and for each type individually (Supporting Figure S15), as percent increase or decrease from the mean mutation rate across all branches.

### 4.11 Hierarchical Orthologous Groups

The output from OrthoFinder was used to retrieve HOGs. For each internal node N0–N9, OrthoFinder produces a table (N0.tsv . . . N9.tsv) listing all HOGs inferred to originate at that node, together with the gene IDs present for each extant species. To treat all taxa uniformly, extant species were considered leaf nodes equivalent to internal nodes (e.g., *Atropacarus striculus* corresponds to node ”Atropcarus striculus”, *Achipteria coleoptrata* to ”Achipteria coleoptrata”, etc.). In each HOG file, presence was defined as having *≥* 1 gene assigned to that species or node. The core (ancestral) HOG set was defined as all HOGs originating at node N0, representing the inferred ancestral genome content prior to diversification of the focal mite clade. For each internal node and species, we computed: core HOG retained (number of ancestral N0 HOGs still present, *≥* 1 gene), core HOG lost (number of ancestral HOGs absent from that node core HOG total – core HOG retained), and percent core HOG retained (proportion of ancestral HOGs retained). These values quantify genome erosion and evolutionary conservation. To characterize lineage-specific genomic content, we identified: species-specific HOGs, the HOGs present in exactly one species and absent from all others, with the number of species-specific HOGs and the number of genes contained in these HOGs; ORFans, the genes not assigned to any orthogroup (from Orthogroups UnassignedGenes.tsv); and TE-associated genes, previously identified. Overlapping genes from these categories were detected for calculations. For each species, we obtained three non-overlapping gene sets: non-TE ORFans, non-TE species-specific genes, TE-associated genes. To summarize lineage-specific genomic innovation, we defined a Novelty Index per species as a sum of non-TE ORFans, non-TE species-specific genes and TE-associated genes.

### 4.12 Evolutionary molecular rates analysis

To explore the relationship between reproductive mode and the efficiency of selection, we calculated molecular evolutionary rates for single-copy ortholog genes retrieved using OrthoFinder.

Briefly, following [Errbii et al., 2024], we used ClustalO v1.2.4 [Sievers et al., 2011] to generate amino acid alignments for each of the 2,899 single-copy ortholog groups. These alignments were then converted into nucleotide alignments using Pal2Nal v14 [Suyama et al., 2006]. Next, the alignments were filtered for poorly aligned positions using Gblocks v0.91b [Talavera and Castresana, 2007] and for sequences shorter than 100 bases, resulting in a final set of 2,890 single-copy orthologs. A phylogenetic tree was then generated for each of these genes using RAxML v.8.2.12 [Stamatakis, 2014].

Prior to analyzing protein sequence evolution, errors were further filtered out in the multiple sequence alignments following [Selberg et al., 2024]. For this, we first applied HyPhy’s BUSTED (Branch-site Unrestricted Statistical Test for Episodic Diversification) v.2.5.48 [Murrell et al., 2015] with the `--error-sink` Yes option. Next, we used BUSTED-E [Selberg et al., 2024], an extension of BUSTED, which improves alignment quality by identifying and removing misaligned and/or low-quality sequences that could introduce biases into evolutionary rate calculations. The resulting filtered alignments were then used for downstream analyses of selection efficiency. To estimate the branch-wide rate omega (*ω*), the ratio of non-synonymous to synonymous substitutions (dN/dS), we applied HyPhy’s aBSREL (adaptive Branch-Site Random Effects Likelihood) [Smith et al., 2015] to the 2,890 orthologous coding sequences. This model tests whether a proportion of sites have evolved under positive selection for each branch in the phylogeny. Similarly, we applied the RELAX model [Wertheim et al., 2015], implemented in the HyPhy package, to infer selection intensity differences between a test group (here the asexual species) and a reference group (here the sexual species). This model categorizes codon sites into three *ω* classes. The first two *ω* classes summarize regions under purifying selection (*ω <* 1) while the third category summarizes regions experiencing adaptive evolution (*ω >* 1). The RELAX model also estimates a selection intensity parameter (k), which infers whether natural selection is relaxed (k ¡ 1) or intensified (k *>* 1) in the test branches relative to the reference branches. To ensure data quality, we excluded genes with *ω* values exceeding 10, as extreme values may indicate errors or unreliable estimates. This filtering resulted in 2,890 single-copy orthologs for the aBSREL analysis and 1,740 single-copy orthologs for the RELAX analysis. Finally, GO (Gene Ontology) enrichment analyses were performed using topGO [Alexa and Rahnenführer, 2009] to identify functional categories enriched in genes showing either intensification (k *>* 1, FDR *<* 0.05) or relaxation (k *<* 1, FDR *<* 0.05) of selection in asexual relative to sexual lineages.

### 4.13 Gene family expansion and contraction analysis

The gene family expansion and contraction analysis was performed using the integrated CAFE5 [Mendes et al., 2020] function of a local docker-installation of Orthovenn3 v1.0.4 [Sun et al., 2023]. OrthoMCL [Li et al., 2003] was selected as the orthologous gene identification algorithm, and the default values were used for the e-value cutoff (1e-2) and the inflation value (1.50). The options for annotation, protein similarity and cluster relationship network were enabled. The mean divergence times for the pairs *A. coleoptrata* and *H. lacustris* (207 Mya) and *H. rufulus* and *S. magnus* (504 Mya) were obtained from the online tool Timetree 5 (timetree.org, accessed 30.10.2025) [Kumar et al., 2022] for manual input of the parameter in Orthovenn3. Information on GO enrichments for the expanded and contracted gene families was also obtained through the integrated functionality of Orthovenn3.

## Data availability

Assemblies and sequencing datasets were deposited in the European Nucleotide Archive under project number PRJEB104491.

## Acknowledgements

This work was supported by the DFG Research Infrastructure West German Genome Center (407493903) as part of the Next Generation Sequencing Competence Network (project 423957469). NGS analyses were carried out at the production site West German Genome Center Cologne. Jens Bast and Philipp Schiffer received the DFG grant BA 58004-1 as part of the DFG Sequencing call 2021. The project was further supported by a DFG Emmy Noether grants BA 5800/4-1 to JB and SCHI 1365/2-1 to PHS. KG was partly funded within the B08 sub-project of the SFB 1211 (DFG project 268236062) led by PHS, and LJ was funded through the UzK forum grant ”BioC2” to PHS. Computational support and infrastructure were provided by the “Centre for Information and Media Technology” (ZIM) at the University of Düsseldorf (Germany). NG’s position was funded by the European Union’s Horizon Europe research and innovation programme under the Marie Skl-odowska-Curie grant agreement No 101110569. We thank Jaroslaw Kowalski for his contribution to sampling *Hydrozetes thienemanni* individuals, and the Genome Technology Center (RGTC) at Radboudumc for the use of the Sequencing Core Facility (Nijmegen, The Netherlands), which provided the PacBio SMRT sequencing service on the Sequel IIe platform for the species *Oppiella nova*.

## Authors contributions

NG, HO, VB, LB, IDS, IS, ASe, RL, JB collected/cultured individuals. NG, HO, SW, MB, IDS, ASc, CB, KB, MB, LS generated sequencing data. NG, ME, VB, KG, LB, JZ analyzed the data. NG, PHS, JB acquired funding. NG, HO, JB wrote the initial manuscript, PHS wrote and edited the final manuscript and all co-authors edited and approved this. Specifically, ME analyzed molecular evolutionary rates and mutational landscapes, generated the two accompanying figures, and wrote the associated results and methods sections. HO collected and maintained oribatid mite cultures, and established and performed the DNA and RNA isolation protocols that enabled generation of molecular dataset. HO supervised and trained VB, LB, SW, and MB during laboratory work and data production. HO contributed to the original manuscript draft and provided organismal, biological, and evolutionary expertise that informed interpretation of the results. NG collected oribatid mites, performed DNA extractions, and prepared Hi-C libraries. NG validated and maintained sequencing data. NG generated all assemblies, repeat and gene annotations. NG generated mitochondrial genome assemblies along with LB and JZ. NG conducted analyses on gene content. NG supervised the overall data acquisition and completion of analyses for the project, coordination with sequencing centers, and supervised and trained in bioinformatics analyses VB, KG, LB, MB, IDS, and JZ. NG led the redaction of the original manuscript.

