## Supplementary_material for "Genomes of ancient asexual mites appear streamlined in their architecture"

### Supplementary Figures

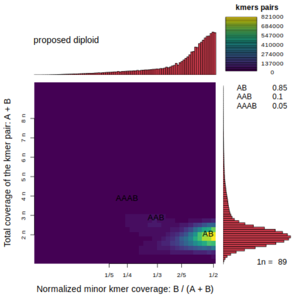

(a) *Platynothrus peltifer.*

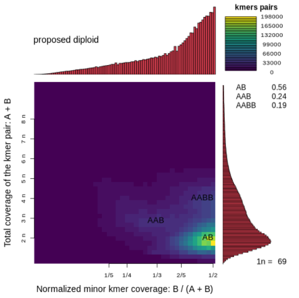

(b) *Nothrus palustris.*

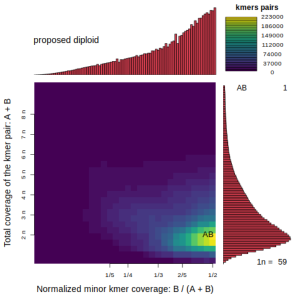

(c) *Hermannia gibba.*

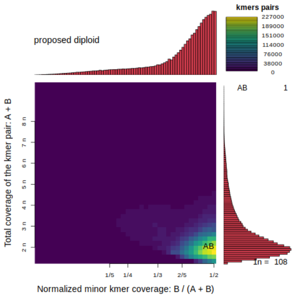

(d) *Hydrozetes lacustris.*

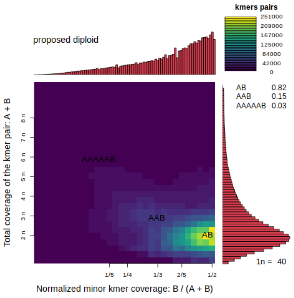

(e) *Hydrozetes thienemanni.*

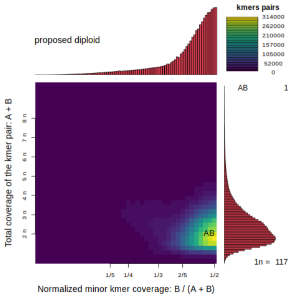

(f) *Oppiella nova.*

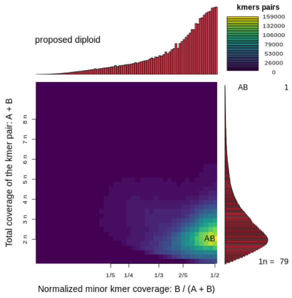

(g) *Oppiella subpectinata.*

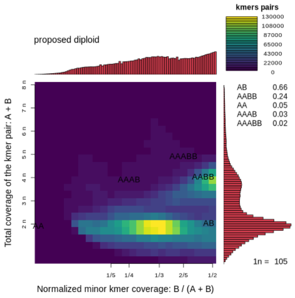

(h) *Atropacarus striculus.*

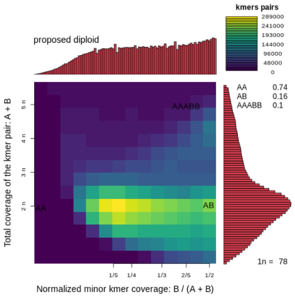

(i) *Steganacarus magnus.*

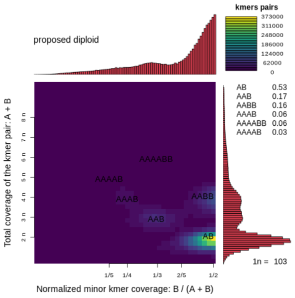

(j) *Hypochthonius rufulus.*

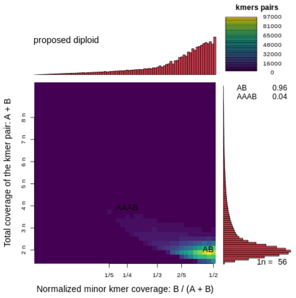

(k) *Achipteria coleoptrata.*

Figure 1: Smudgeplot ploidy prediction based on $k$-mers extracted from PacBio HiFi long reads, with $k$=27. All species are predicted as diploid.

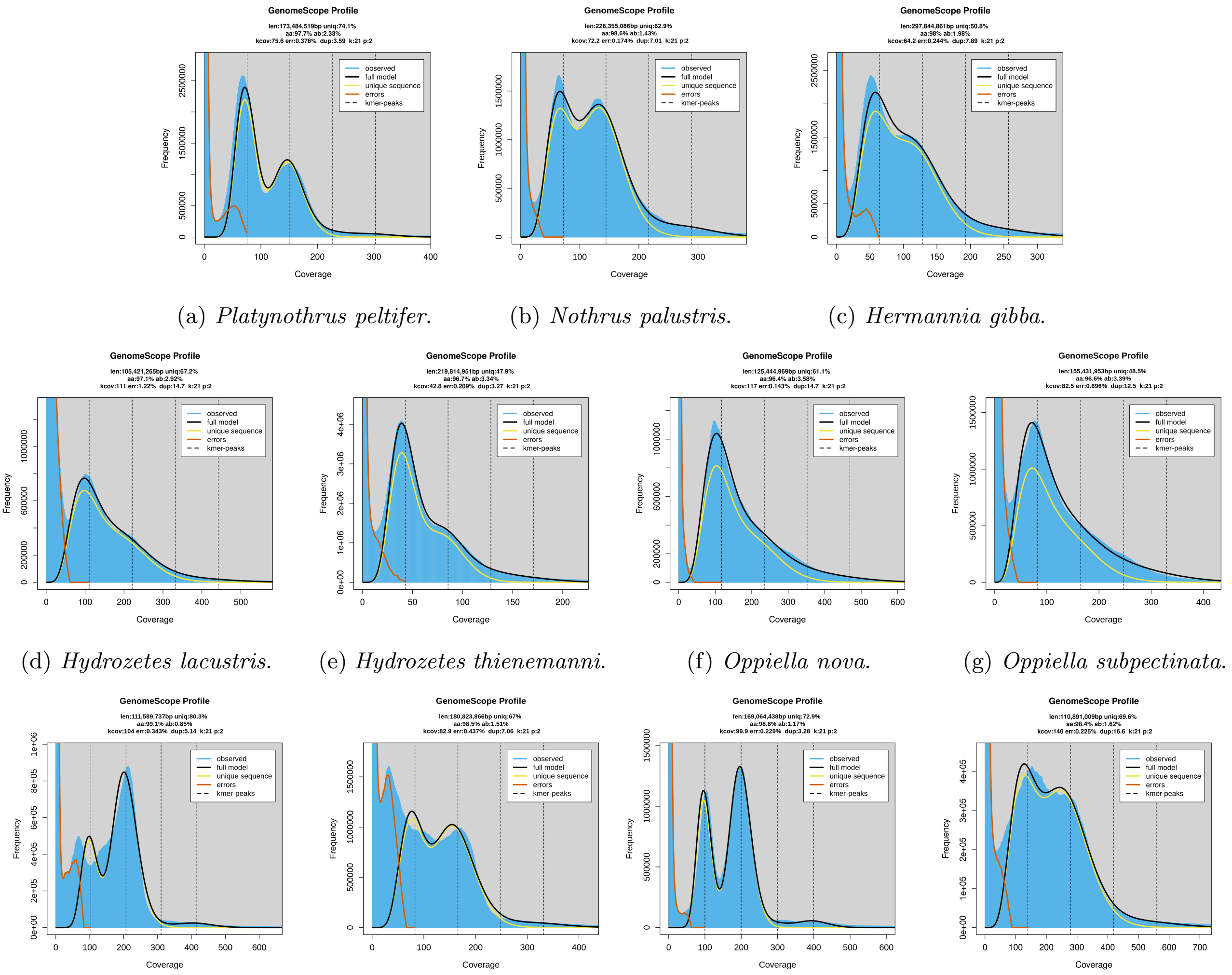

(a) *Platynothrus peltifer.* (b) *Nothrus palustris.* (c) *Hermannia gibba.*

(d) *Hydrozetes lacustris.* (e) *Hydrozetes thienemanni.* (f) *Oppiella nova.* (g) *Oppiella subpectinata.*

(h) *Atropacarus striculus.* (i) *Steganacarus magnus.* (j) *Hypochthonius rufulus.* (k) *Achipteria coleoptrata.*

Figure 2: GenomeScope analysis with $k$=21.

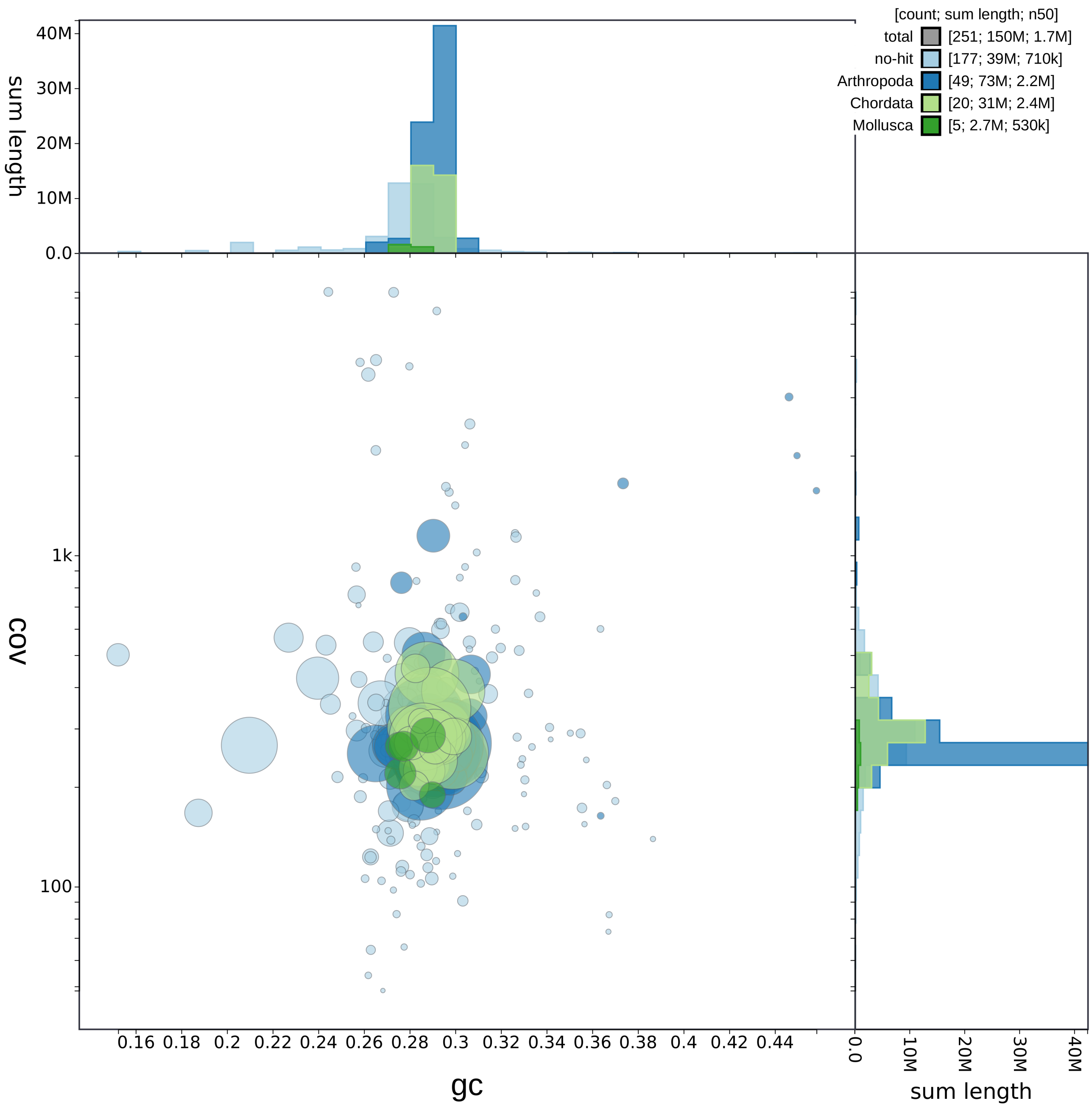

Figure 3: Blobplot of the genome assembly of *Achipteria coleoptrata* after decontamination.

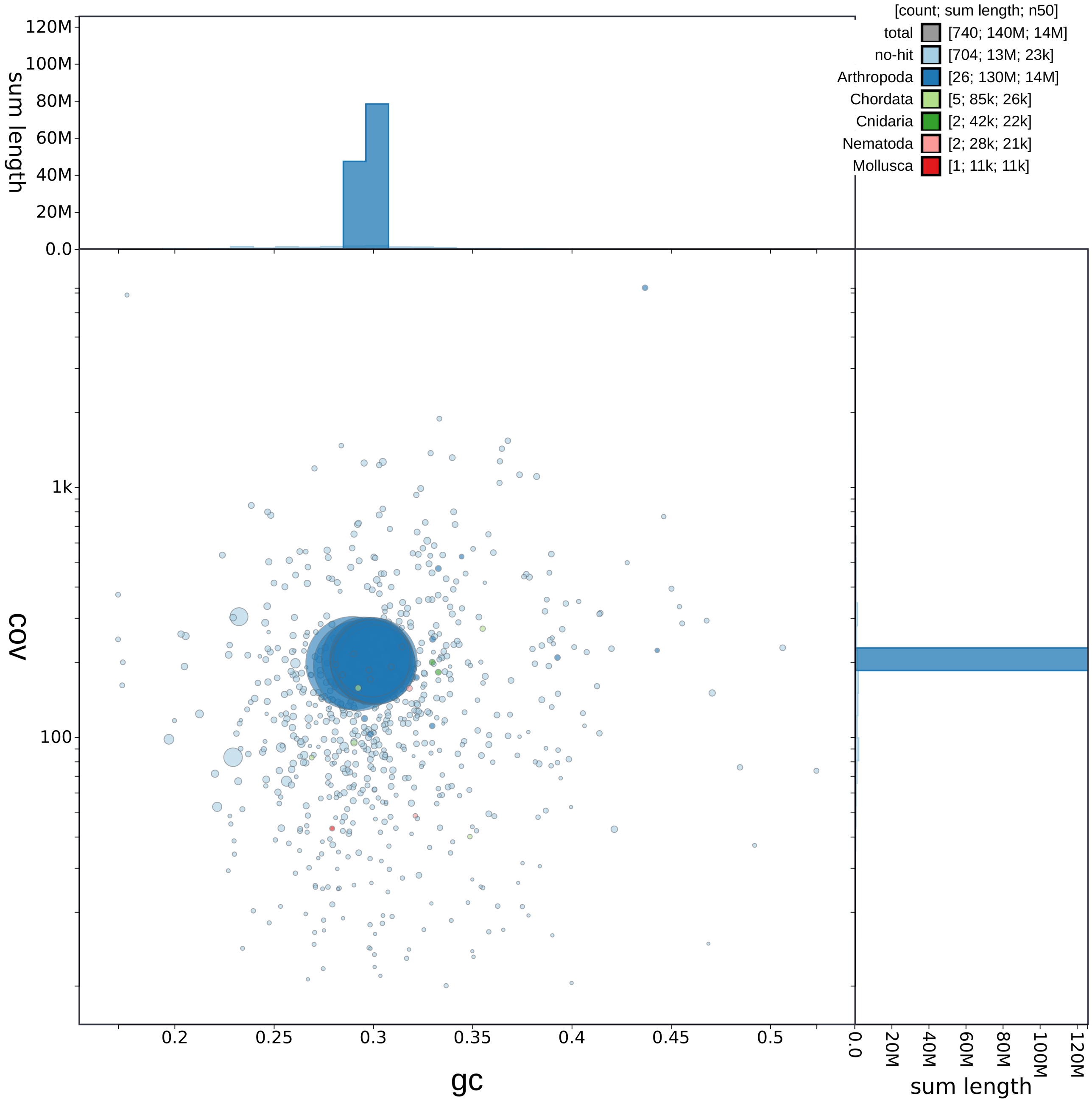

Figure 4: Blobplot of the genome assembly of *Atropacarus striculus* after decontamination.

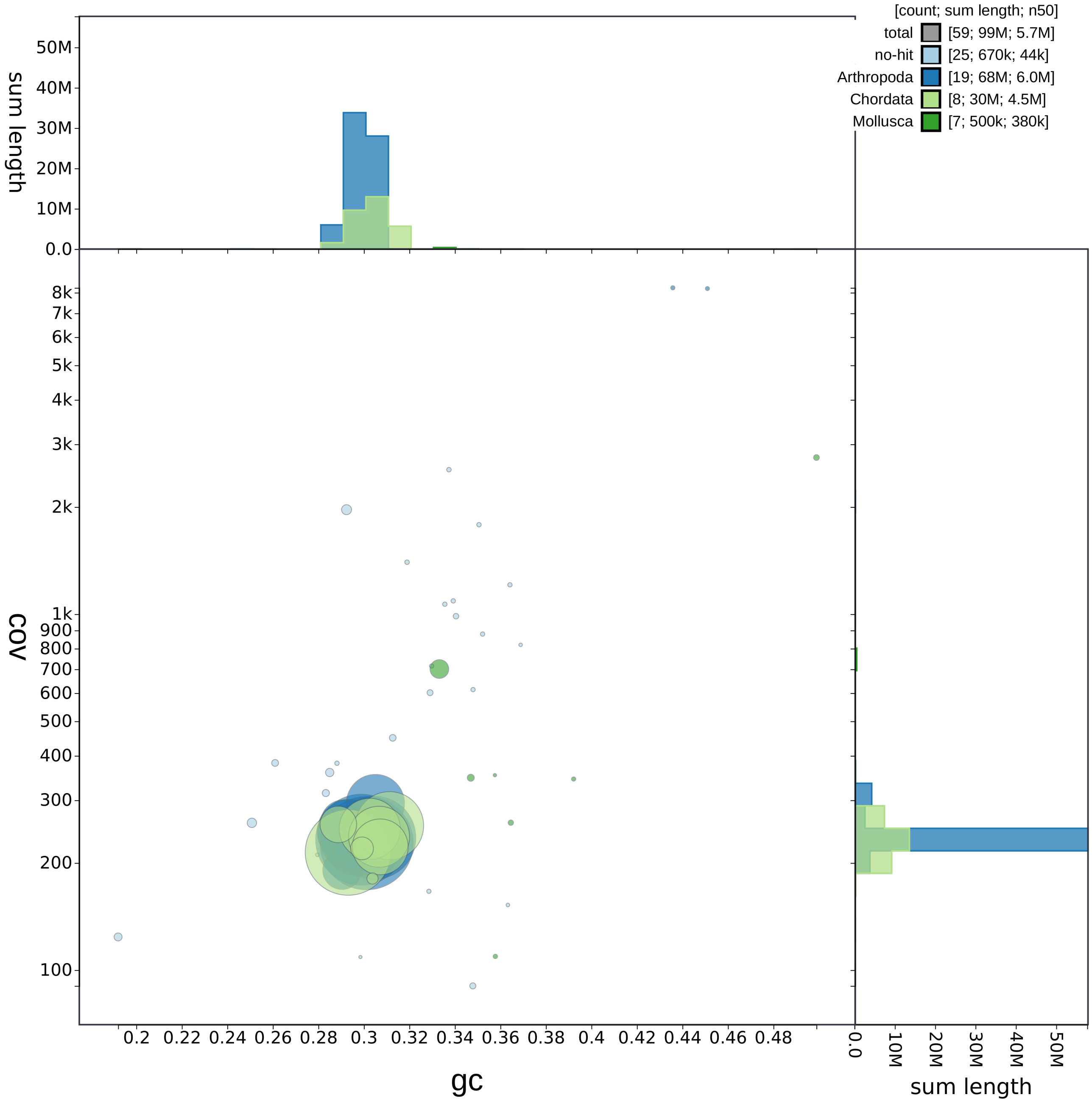

Figure 5: Blobplot of the genome assembly of *Hydrozetes lacustris* after decontamination.

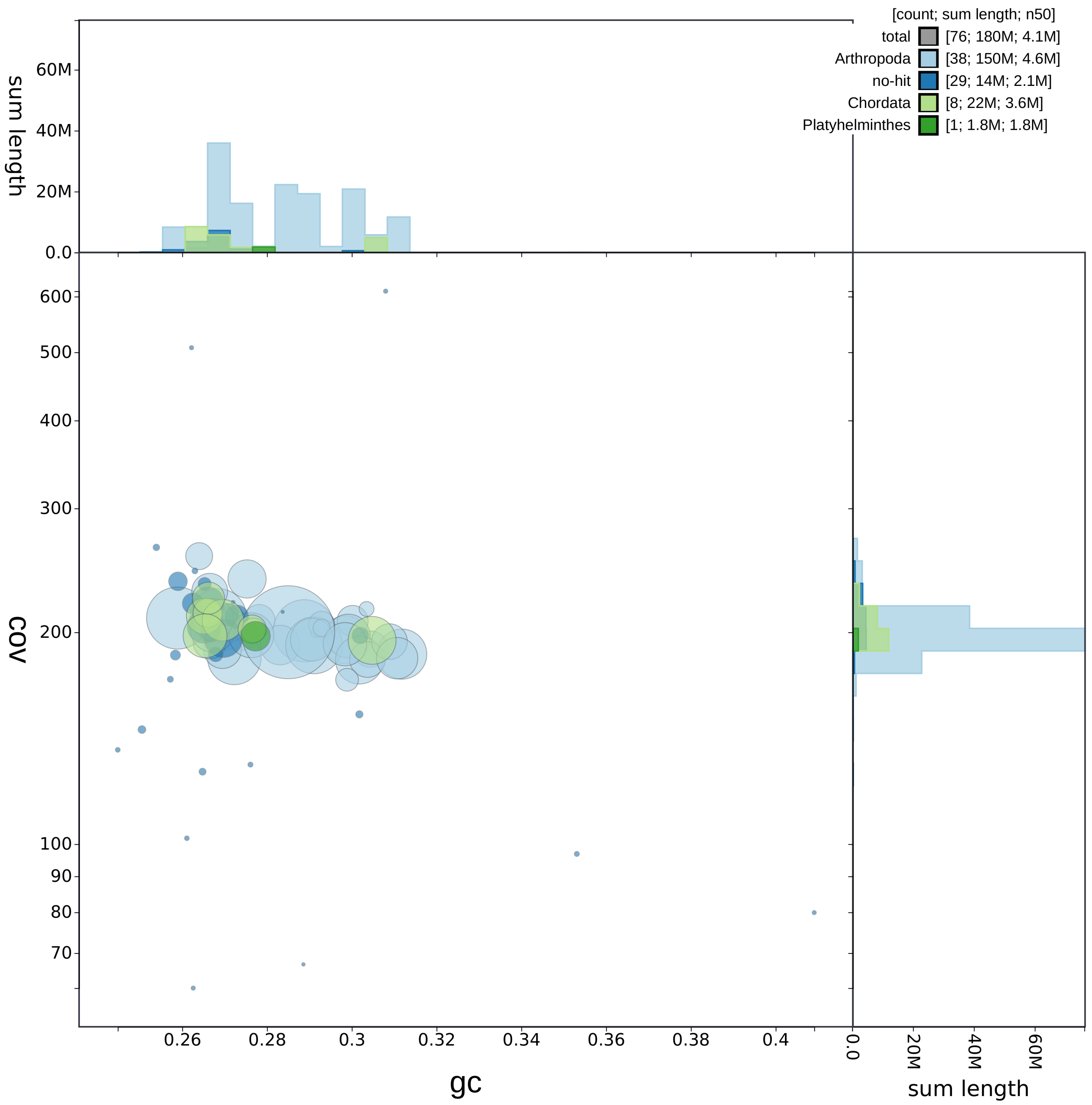

Figure 6: Blobplot of the genome assembly of *Hypochthonius rufulus* after decontamination.

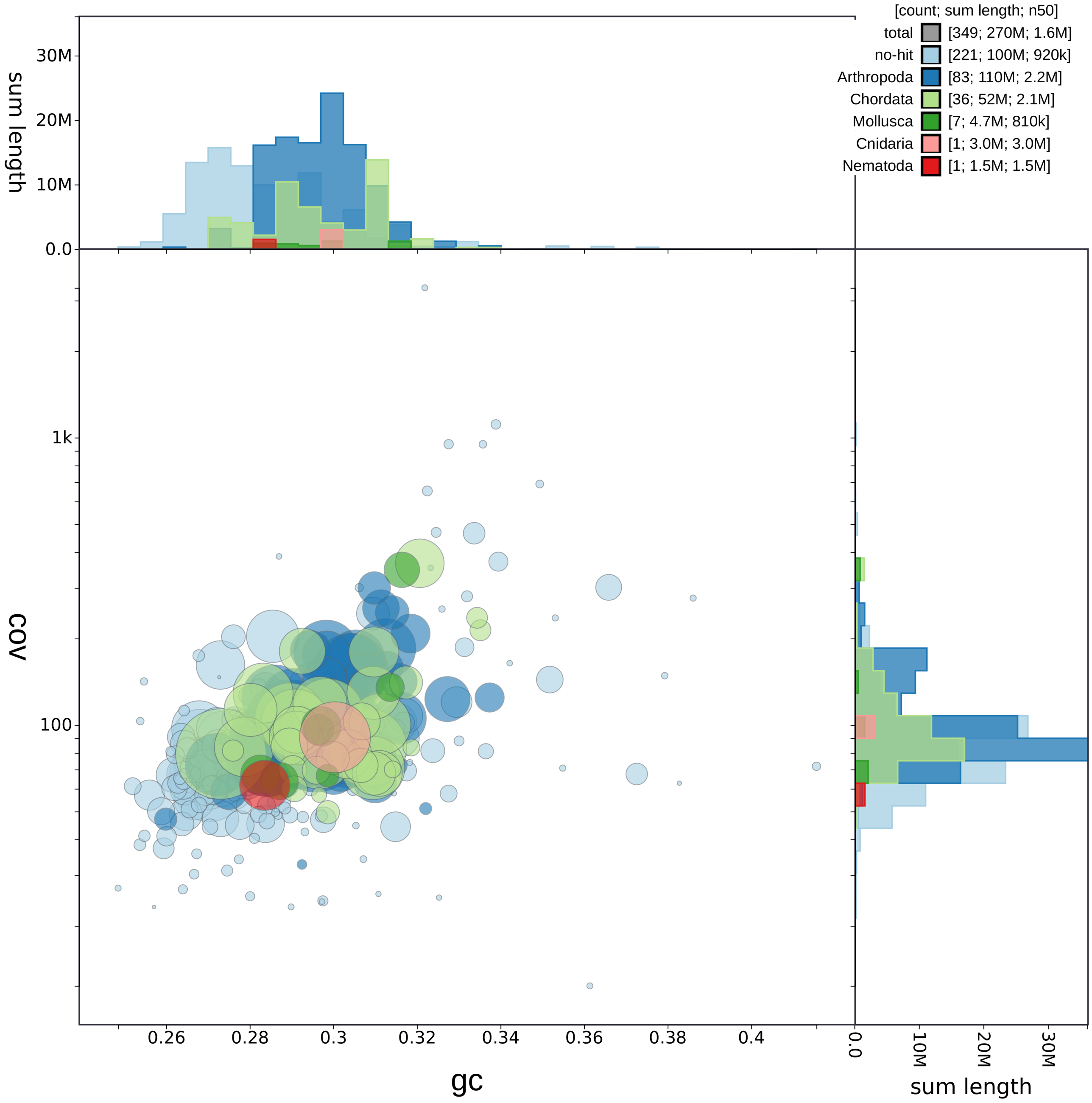

Figure 7: Blobplot of the genome assembly of *Hydrozetes thienemanni* after decontamination.

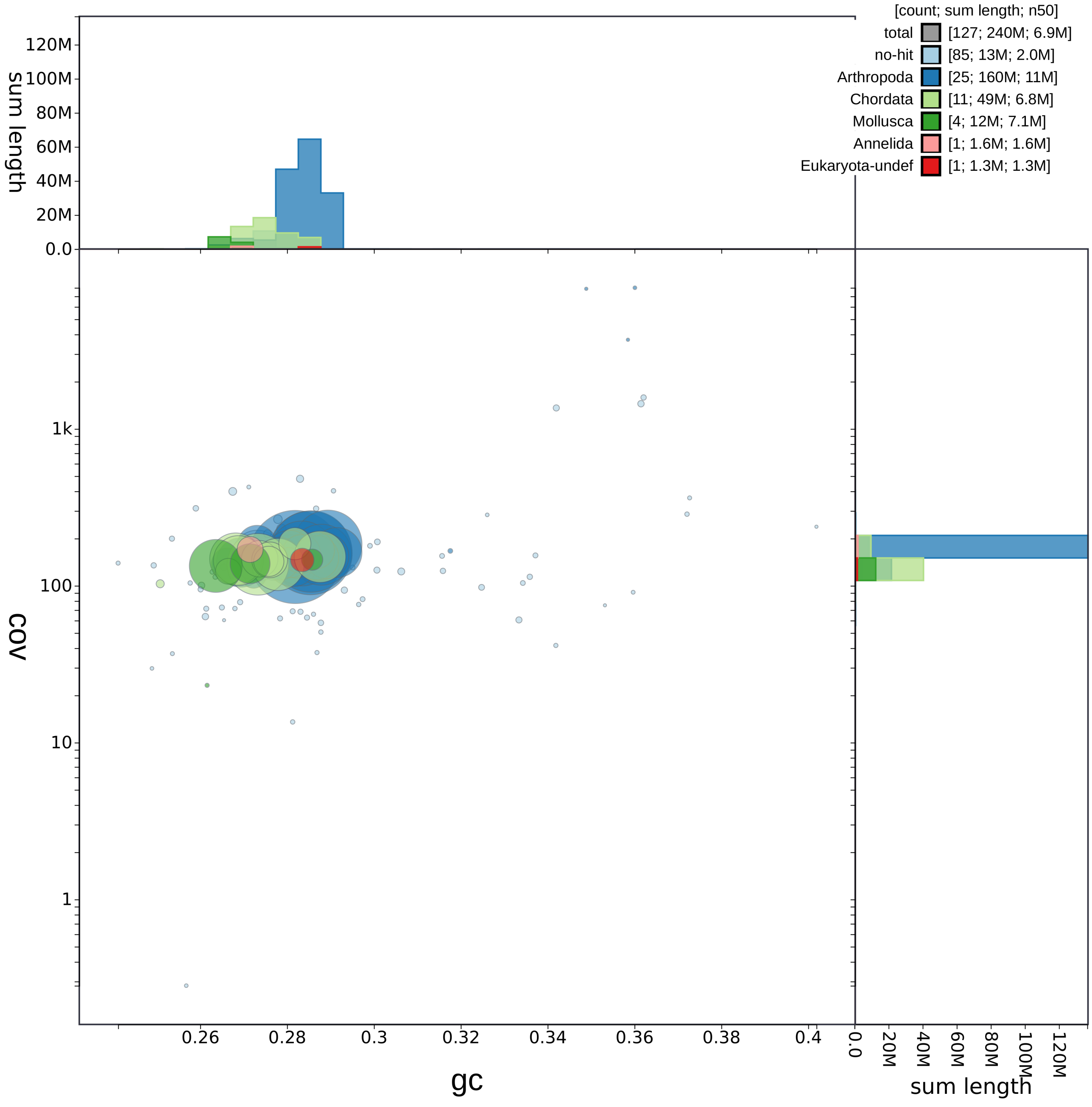

Figure 8: Blobplot of the genome assembly of *Nothrus palustris* after decontamination.

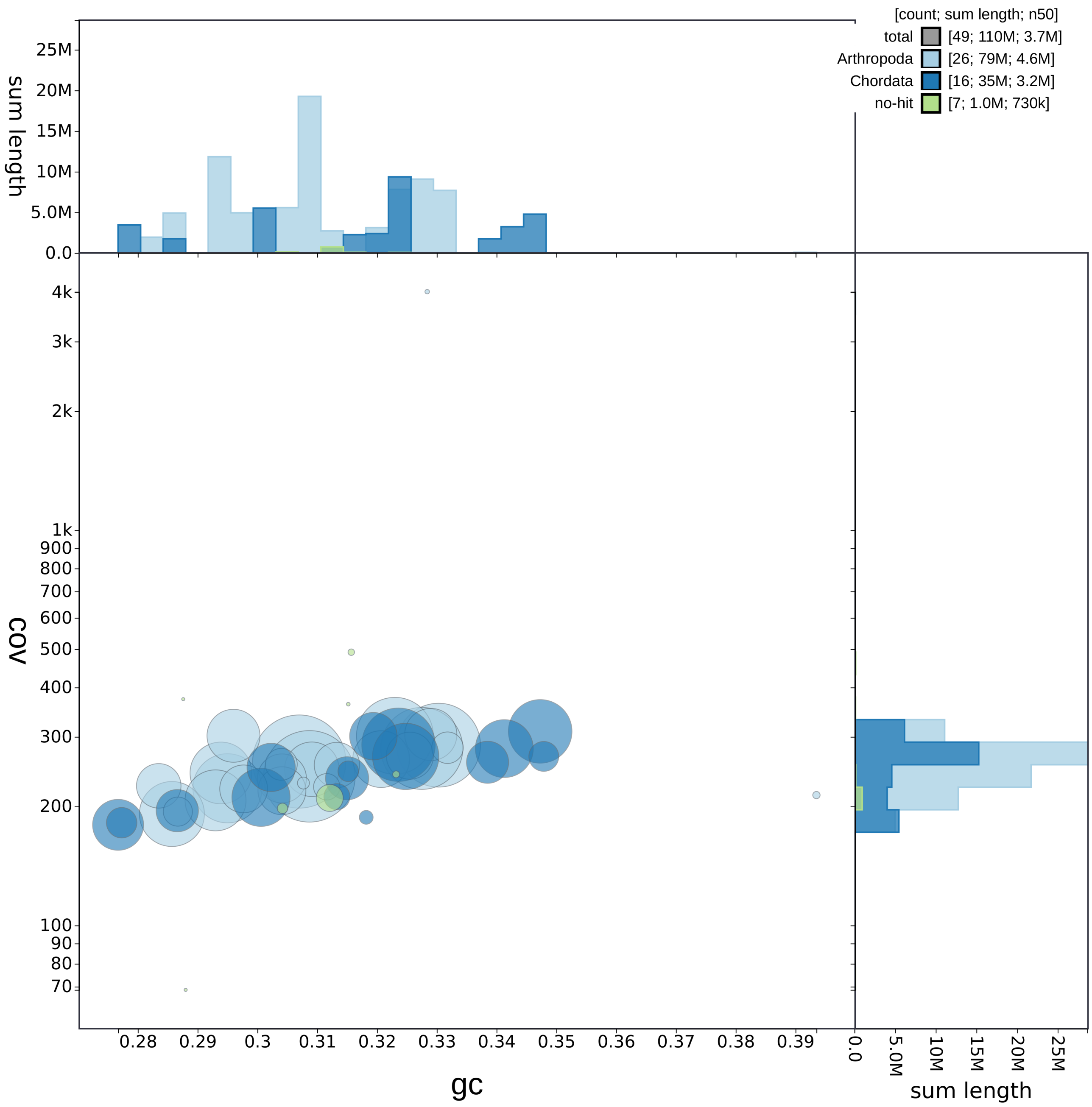

Figure 9: Blobplot of the genome assembly of *Oppiella nova* after decontamination.

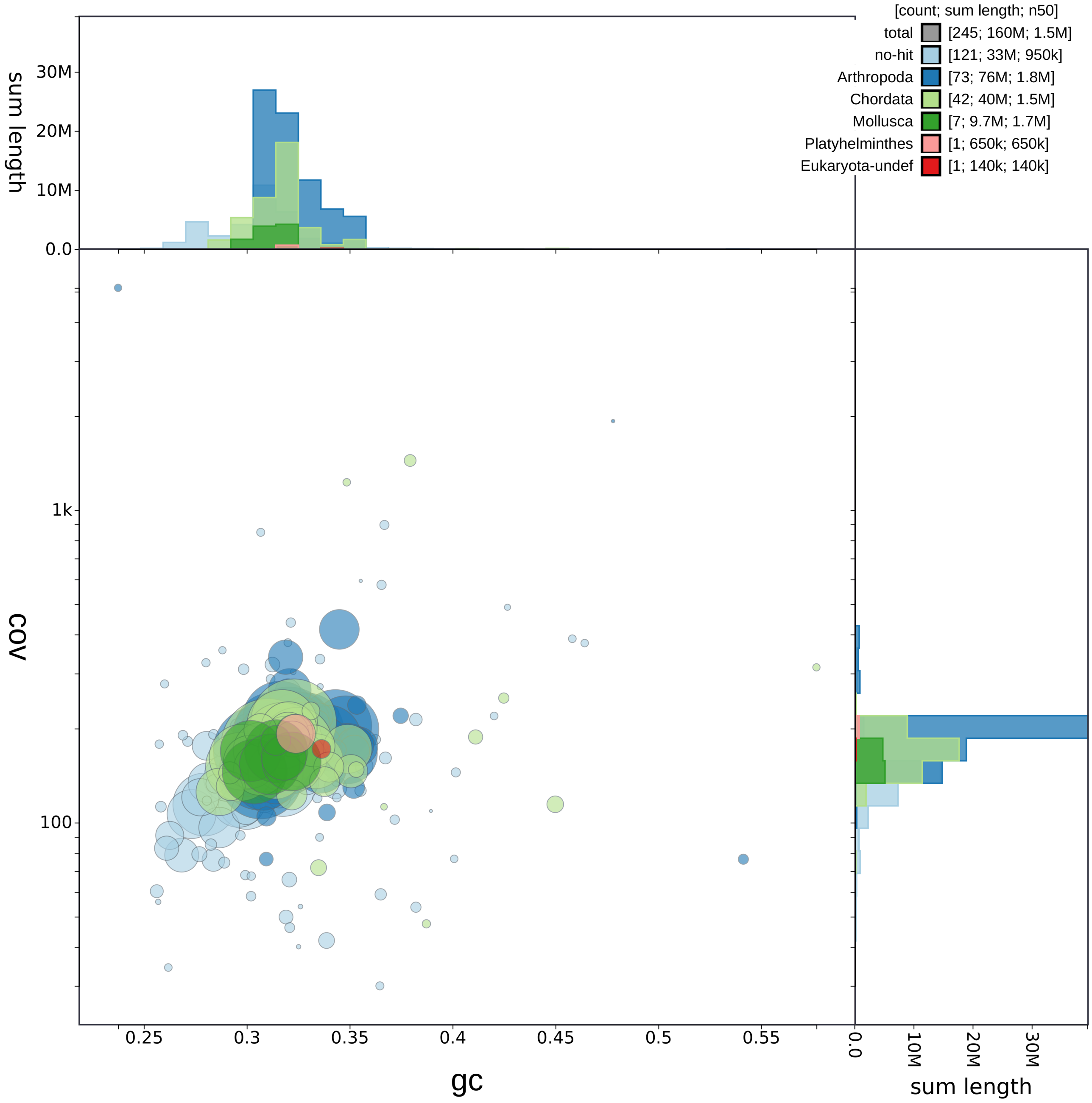

Figure 10: Blobplot of the genome assembly of *Oppiella subpectinata* after decontamination.

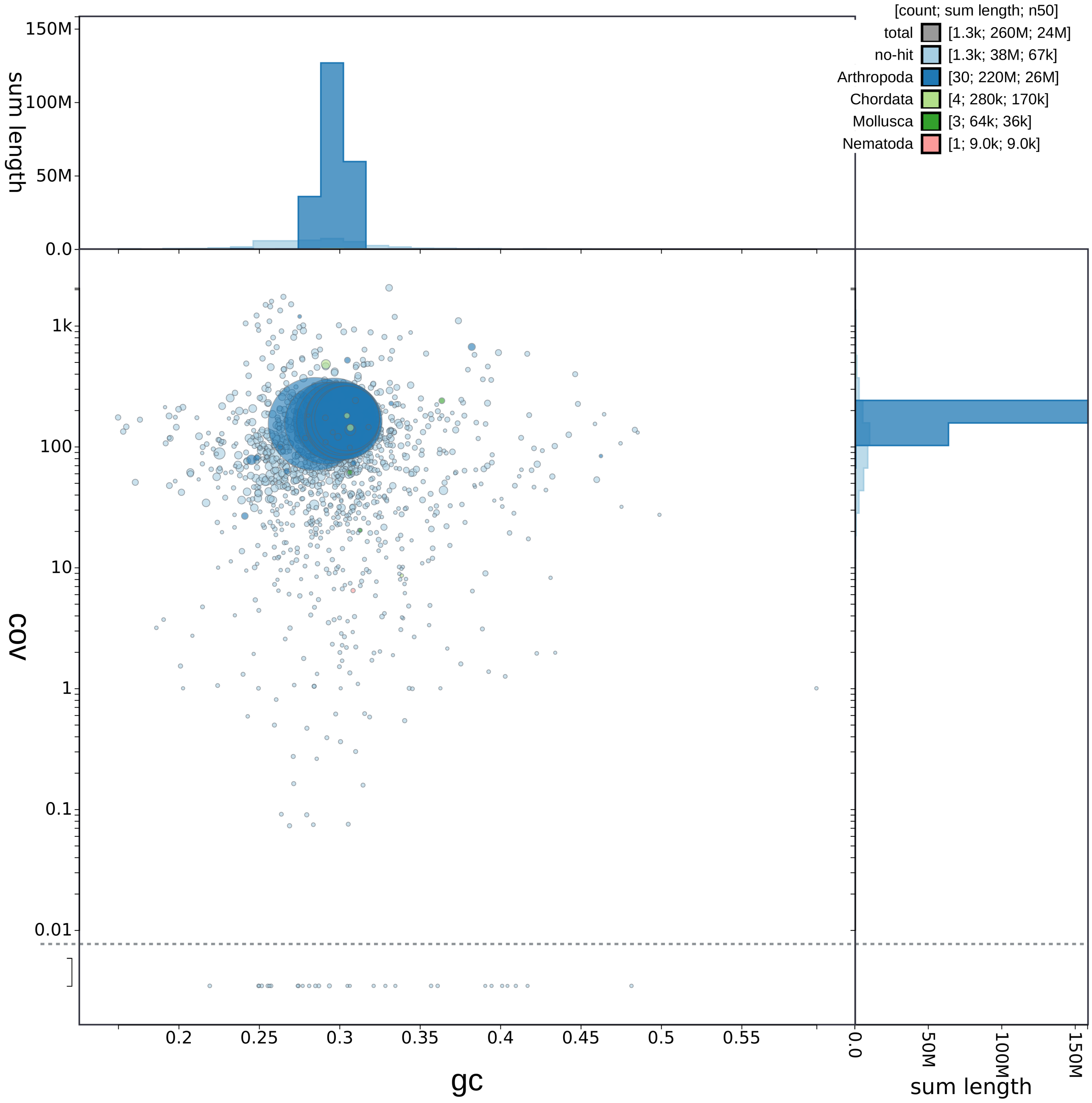

Figure 11: Blobplot of the genome assembly of *Steganacarus magnus* after decontamination.

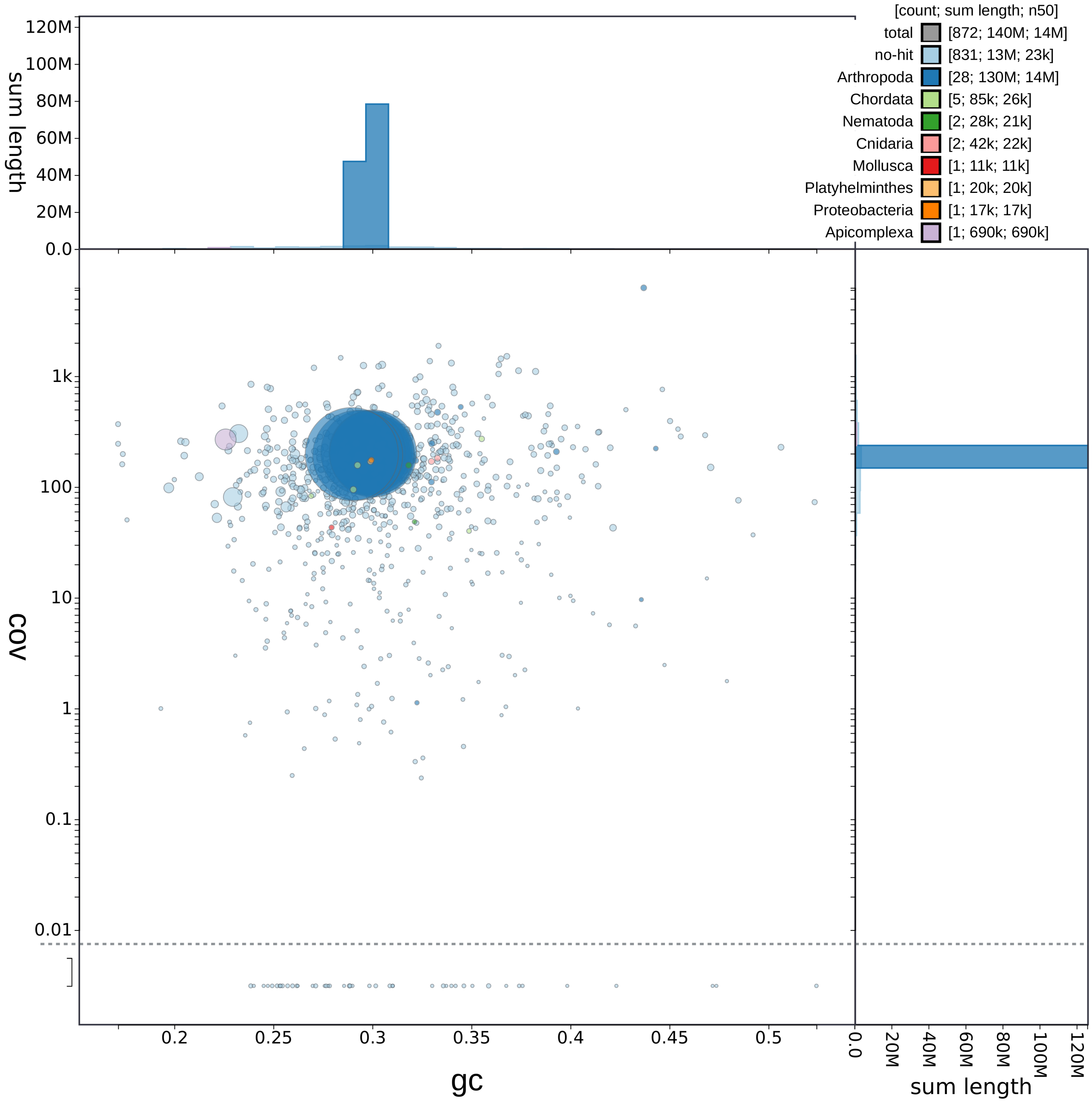

Figure 12: Blobplot of the genome assembly of *Atropacarus stricuIus* before decontamination.

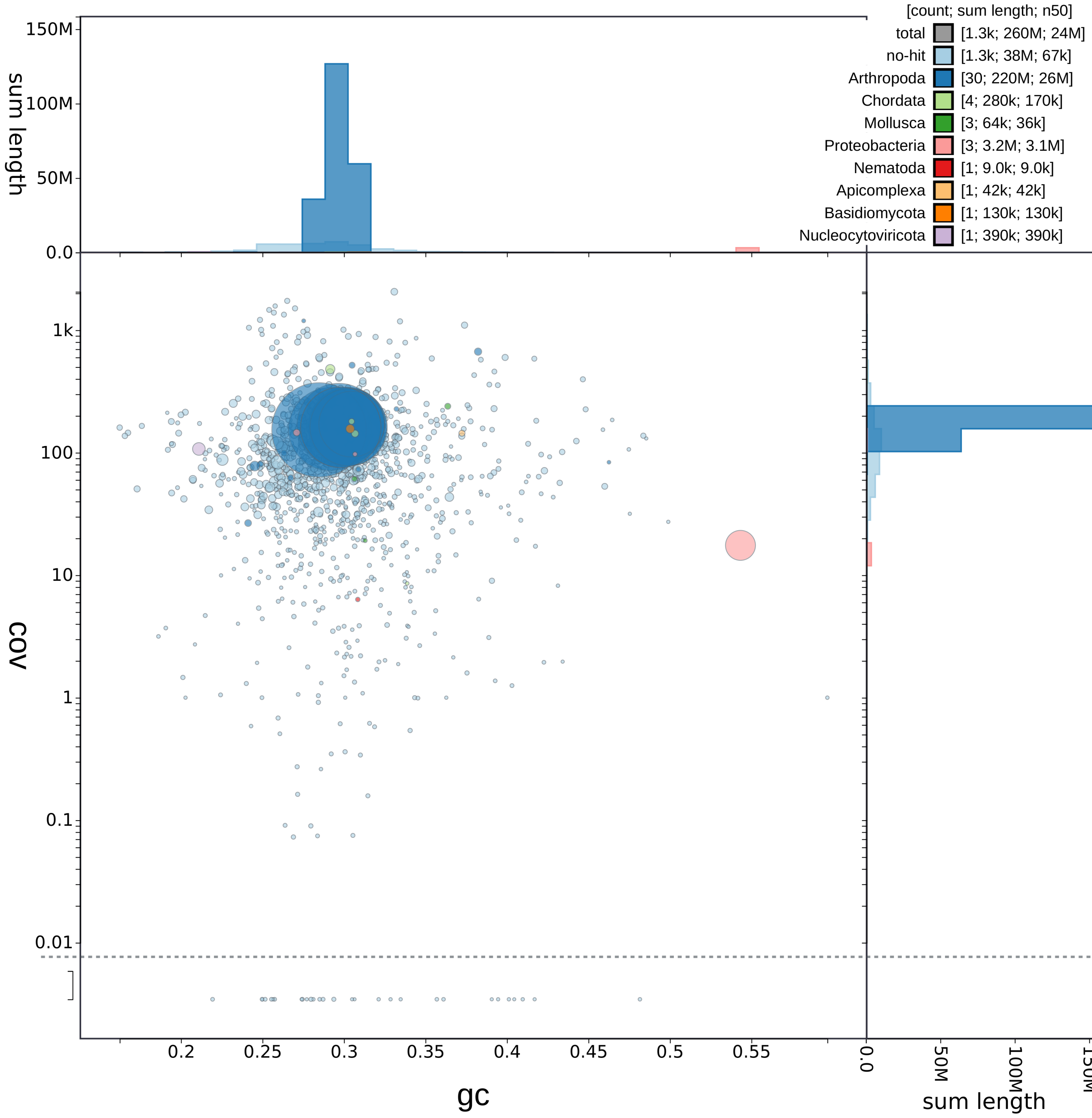

Figure 13: Blobplot of the genome assembly of *Steganacarus magnus* before decontamination.

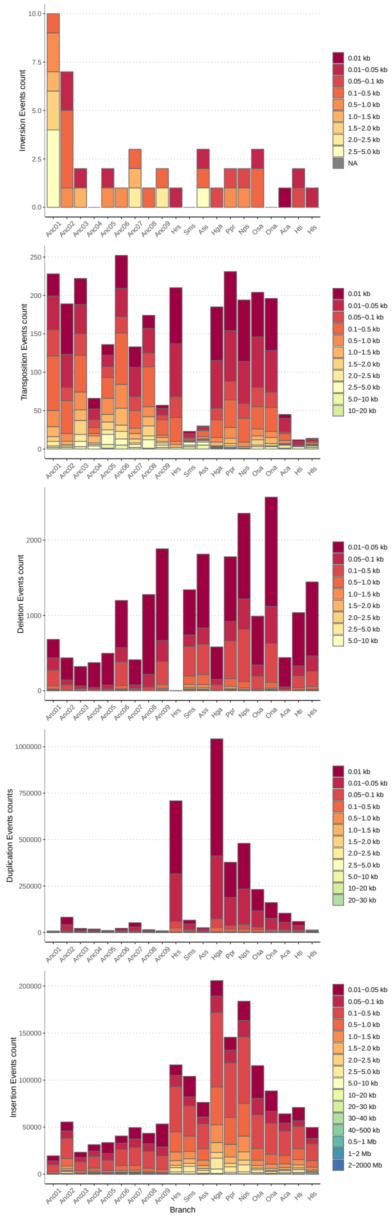

Figure 14: Mutational events across the phylogeny of eleven oribatid mites. The number and size of different mutational events are shown for each branch and node in the phylogeny. From top to bottom, the plots display inversions, transpositions, deletions, duplications, and insertions.

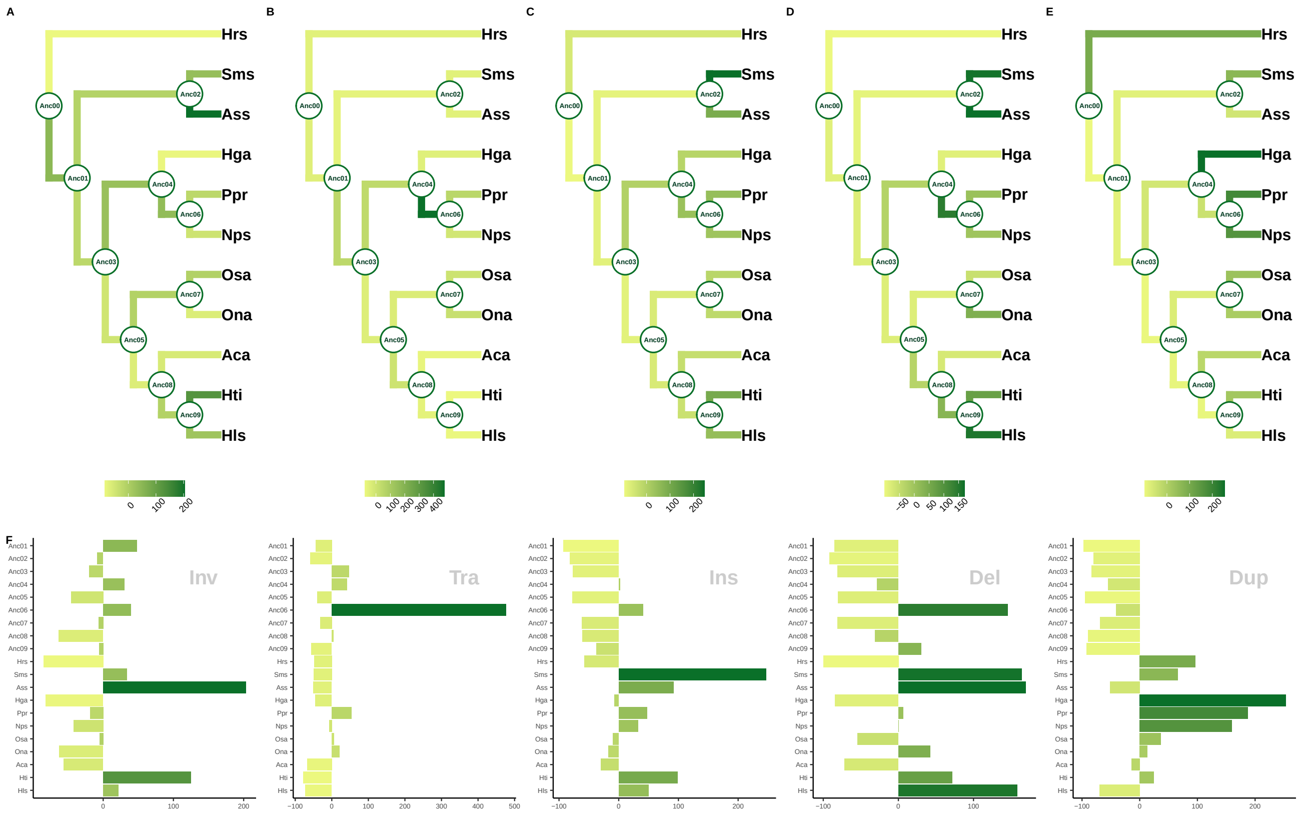

Figure 15: Relative enrichment of mutational events across the mite phylogeny. Shown are percent deviations from the mean number of events across all branches. From left to right, the trees and plots show the relative increase (positive values) or decrease (negative values) in inversions, transpositions, deletions, duplications, and insertions.

### Genomes of ancient asexual mites appear streamlined in their architecture

Nadège Guiglielmoni[1*], Mohammed Errbii[1], Hüsna Öztoprak[1], Viktoria Bednarski[1], Karim Gueddach[1], Lea Borgschulte[1], Svenja Wulsch[1], Marshall Brickheimer[1], Isabella Di Stefano[1], Lucy Jimenez[1], Jakob Zelzner[1], Anja Schuster[2,3], Christian Becker[4,5], Kerstin Becker[2,3,4,5], Miklos Balint[6,7,8], Ricarda Lehmitz[9], Ina Schaefer[6,7], Leonie Schardt[7], Anna Seniczak[10], Philipp H. Schiffer[1†], and Jens Bast[1*†]

[1]Institut für Zoologie, Universität zu Köln, Cologne, Germany

[2]Biological and Medical Research Center (BMFZ), Medical Faculty and University Hospital Düsseldorf, Heinrich Heine University Düsseldorf, Düsseldorf, Germany

[3]West German Genome Center (WGGC), Medical Faculty and University Hospital Düsseldorf, Heinrich Heine University Düsseldorf, Düsseldorf, Germany

[4]Cologne Center for Genomics (CCG), Medical Faculty, University of Cologne, Cologne, Germany

[5]West German Genome Center (WGGC), University of Cologne, Cologne, Germany

[6]Senckenberg Biodiversity Climate Research Center, Frankfurt Main, Germany

[7]Loewe Center for Translational Biodiversity Genomics (LOEWE-TBG), Frankfurt Main, Germany

[8]Department of Insect Biotechnology, Justus-Liebig University, Gießen, Germany

[9]Soil Zoology, Senckenberg Museum of Natural History, Görlitz, Germany

[10]Faculty of Applied Ecology, Agricultural Sciences and Biotechnology, University of Inland Norway, Elverum, Norway

[†]these authors contributed equally

### Supplementary Tables

Table 1: General statistics of the 11 genome assemblies, including assembly size, repeat content, N50, BUSCO scores for the genome assembly, number of genes, duplications events, ORFans, TE-associated genes, average exon length, average intron length, average intergenic space length, transposable elements including DNA elements, RNA elements, Helitrons.

Table 2: Mitochondrial genome assemblies.

| **Species** | Assembly size |
|---|---|
| *Hermannia gibba* | 14,340 |
| *Nothrus palustris* | 14,550 |
| *Steganacarus magnus* | 12,939 |
| *Atropacarus striculus* | 15,221 |
| *Hydrozetes thienemanni* | 14,114 |
| *Hydrozetes lacustris* | 17,730 |
| *Oppiella subpectinata* | 13,517 |
| *Oppiella nova* | 14,262 |
| *Achipteria coleoptrata* | 14,028 |

Table 3: Detailed transposable element content.

Table 4: Genes associated with meiosis and sex determination. Entries indicate for each species the number of representatives found for this gene in functionally annotated data.

| **Gene** | **Hls** | **Hti** | **Aca** | **Ona** | **Osa** | **Nps** | **Ppr** | **Hga** | **Ass** | **Sms** | **Hrs** |
|---|---|---|---|---|---|---|---|---|---|---|---|
| SPO11 | 3 | 2 | 0 | 1 | 1 | 1 | 1 | 1 | 1 | 1 | 1 |
| HOP2 | 1 | 1 | 1 | 1 | 1 | 1 | 1 | 1 | 1 | 1 | 1 |
| MND1 | 1 | 1 | 1 | 1 | 1 | 1 | 1 | 1 | 1 | 1 | 1 |
| REC8/RAD21 | 1 | 1 | 1 | 2 | 2 | 1 | 1 | 1 | 2 | 1 | 5 |
| RAD51 | 4 | 4 | 4 | 4 | 5 | 5 | 4 | 4 | 8 | 10 | 4 |
| MEIOC | 1 | 1 | 1 | 1 | 1 | 1 | 1 | 1 | 1 | 1 | 1 |
| HORMAD1 | 2 | 1 | 2 | 1 | 1 | 15 | 4 | 13 | 1 | 1 | 4 |
| tra-2 | 2 | 1 | 2 | 2 | 2 | 1 | 4 | 2 | 2 | 3 | 2 |
| DMRT | 10 | 7 | 8 | 11 | 12 | 10 | 12 | 19 | 13 | 11 | 11 |
| fem-3 | 1 | 1 | 1 | 1 | 1 | 1 | 0 | 1 | 1 | 1 | 1 |

Table 5: Functional enrichment of gained genes across species.

Table 6: Functional enrichment of lost genes across species.

Table 7: Functional enrichment of genes showing intensification or relaxation of selection.

| Species | Short name | Reproductive mode | Assembly_size | Repeat_content | N50 | BUSCO_single | BUSCO_dup | Gene_number | Core_HOG_retained | Core_HOG_lost | Core_HOG_retai | Total_genes_HO |
|---|---|---|---|---|---|---|---|---|---|---|---|---|
| Hypochtonius rufulus | Hrs | asexual | 184719097 | 78265590 | 4133765 | 2708 | 90 | 22129 | 10920 | 16719 | .5093889069793 | 19614 |
| Stegacanarus magnus | Sms | sexual | 260319967 | 117460954 | 24110726 | 2589 | 169 | 28881 | 13352 | 14287 | .3085495133688 | 26893 |
| Atropacarus striculus | Ass | asexual | 138684303 | 32198179 | 13898161 | 2581 | 172 | 20443 | 12091 | 15548 | .7461557943486 | 19428 |
| Hermannia gibba | Hga | sexual | 352241685 | 209544679 | 43860327 | 2721 | 101 | 28683 | 12665 | 14974 | 45.8229313651 | 26550 |
| Platynothrus peltifer | Ppr | asexual | 216037090 | 79131767 | 23397169 | 2748 | 87 | 26055 | 13219 | 14420 | .8273454177069 | 24316 |
| Nothrus palustris | Nps | asexual | 235400948 | 98738467 | 6875046 | 2734 | 105 | 27479 | 13628 | 14011 | 9.307138463765 | 25016 |
| Oppiella subpectinata | Osa | sexual | 158615823 | 37396294 | 1538901 | 2528 | 236 | 27350 | 12934 | 14705 | .7961937841456 | 24770 |
| Oppiella nova | Ona | asexual | 114951853 | 17794353 | 3670518 | 2708 | 90 | 21567 | 12311 | 15328 | .5421324939397 | 19022 |
| Achipteria coleoptrata | Aca | sexual | 145321555 | 46924607 | 1669686 | 2688 | 99 | 22739 | 12593 | 15046 | .5624298997793 | 20804 |
| Hydrozetes thienemanni | Hti | sexual | 271997005 | 137711482 | 1638458 | 2689 | 101 | 27676 | 13142 | 14497 | 47.54875357285 | 26075 |
| Hydrozetes lacustris | Hls | asexual | 98862563 | 15175683 | 5014443 | 2710 | 87 | 19693 | 12607 | 15032 | .6130829624806 | 18425 |

| Species | Mean_Genes_pe | Duplications | Unassigned gene | Species-specific | ORFans | Novelty index | TE_gene_candid | PteMte | PteM0 | PchMte | HGT | HGT% |
|---|---|---|---|---|---|---|---|---|---|---|---|---|
| Hypochtonius rufulus | '9615384615385 | 7302 | 2515 | 4354 | 6869 | 7023 | 201 | 137 | 62 | 2 | 302 | 1,36 |
| Stegacanarus magnus | )1415518274416 | 10208 | 1988 | 4375 | 6363 | 6487 | 147 | 89 | 51 | 7 | 585 | 2,03 |
| Atropacarus striculus | i0681498635349 | 4719 | 1015 | 1601 | 2616 | 2756 | 162 | 92 | 64 | 6 | 288 | 1,41 |
| Hermannia gibba | )9632846427161 | 11475 | 2133 | 7804 | 9937 | 10178 | 317 | 205 | 80 | 32 | 400 | 1,39 |
| Platynothrus peltifer | 33947348513503 | 8557 | 1739 | 2914 | 4653 | 4992 | 407 | 233 | 147 | 22 | 565 | 2,17 |
| Nothrus palustris | 33563252127972 | 8701 | 2463 | 4254 | 6717 | 6996 | 339 | 204 | 113 | 27 | 567 | 2,06 |
| Oppiella subpectinata | )1510746868718 | 9132 | 2580 | 5730 | 8310 | 8515 | 300 | 212 | 58 | 30 | 305 | 1,12 |
| Oppiella nova | i4512224839574 | 4472 | 2545 | 1894 | 4439 | 4502 | 73 | 35 | 32 | 6 | 370 | 1,72 |
| Achipteria coleoptrata | i5202890494719 | 6039 | 1935 | 3116 | 5051 | 5183 | 141 | 82 | 49 | 10 | 350 | 1,54 |
| Hydrozetes thienemanni | )8409678892102 | 9928 | 1601 | 4013 | 5614 | 5941 | 363 | 247 | 90 | 26 | 330 | 1,19 |
| Hydrozetes lacustris | I6148964860792 | 3610 | 1268 | 773 | 2041 | 2186 | 172 | 117 | 45 | 10 | 311 | 1,58 |

| Species | TE% | DNA_elements | RNA_elements | Helitron | Average_Exon | Average_Intron | Average_Intergenic_Space |
|---|---|---|---|---|---|---|---|
| Hypochtonius rufulus | 19,90 | 26681903 | 4269596 | 5805137 | 317,57 | 535,79 | 5578,56 |
| Stegacanarus magnus | 20,54 | 27828621 | 23318593 | 2333029 | 329,43 | 596,06 | 5729,73 |
| Atropacarus striculus | 12,36 | 8717305 | 7710494 | 717651 | 367,03 | 547,70 | 3583,59 |
| Hermannia gibba | 26,47 | 53292933 | 30917601 | 9013474 | 324,53 | 905,88 | 8387,16 |
| Platynothrus peltifer | 15,51 | 19763366 | 12768610 | 976379 | 317,08 | 637,28 | 5127,16 |
| Nothrus palustris | 3,35 | 5109814 | 2267756 | 513094 | 315,47 | 767,31 | 5707,53 |
| Oppiella subpectinata | 12,30 | 10314888 | 8013062 | 1185784 | 320,52 | 517,71 | 2846,84 |
| Oppiella nova | 6,79 | 5563028 | 1644534 | 598777 | 287,29 | 500,98 | 2696,76 |
| Achipteria coleoptrata | 16,65 | 8765698 | 15158293 | 267188 | 309,46 | 424,64 | 3575,08 |
| Hydrozetes thienemanni | 24,07 | 36552519 | 28254419 | 658817 | 344,98 | 462,00 | 6753,19 |
| Hydrozetes lacustris | 7,23 | 3665609 | 3477113 | 3785 | 337,38 | 371,60 | 2688,00 |

| Species | Assembly_size | DNA elements | RNA elements | DTA_hAT_all | DTB_Piggy_all | DTC_CACTA_al | DTH_Harbinger_ | DTE_Merli | DTM_Mutator_a | DTP_P | DTT_TC1 | DXX |
|---|---|---|---|---|---|---|---|---|---|---|---|---|
| Hrs | 184719097 | 26681903 | 4269596 | 15114108 | 207851 | 142380 | 467451 | 0 | 2048765 | 0 | 8701348 | 17194954 |
| Ass | 138684303 | 8717305 | 7710494 | 3576707 | 6517 | 1039288 | 377772 | 116264 | 905927 | 130893 | 2811094 | 4099690 |
| Sms | 260319967 | 27828621 | 23318593 | 12589272 | 11646 | 1337490 | 1806919 | 0 | 1044083 | 0 | 11039211 | 17450485 |
| Ppr | 216037090 | 19763366 | 12768610 | 7116114 | 0 | 1012274 | 531576 | 0 | 384343 | 1052191 | 10719059 | 12179501 |
| Nps | 235400948 | 5109814 | 2267756 | 2380526 | 17553 | 46374 | 160310 | 0 | 245643 | 103091 | 2259408 | 3162883 |
| Hga | 352241685 | 53292933 | 30917601 | 25585691 | 0 | 1300454 | 3112070 | 0 | 1518571 | 0 | 21776147 | 25472610 |
| Ona | 114951853 | 5563028 | 1644534 | 1911901 | 0 | 208381 | 118724 | 0 | 374675 | 0 | 2949347 | 4501877 |
| Osa | 158615823 | 10314888 | 8013062 | 3071607 | 44481 | 1212886 | 409171 | 0 | 1274749 | 0 | 4301994 | 6606788 |
| Aca | 145321555 | 8765698 | 15158293 | 3497399 | 0 | 62649 | 269186 | 0 | 153012 | 497342 | 4783452 | 8011813 |
| Hti | 271997005 | 36552519 | 28254419 | 8713985 | 0 | 3212084 | 2456701 | 0 | 2356889 | 493797 | 19812860 | 17985674 |
| Hls | 98862563 | 3665609 | 3477113 | 1198266 | 0 | 52785 | 169628 | 0 | 22079 | 197649 | 2222851 | 2802886 |

| Species | DHH_Helitron | DTA_hAT_MIT | DTA_hAT_unkno | DTB_Piggy | DTB_PiggyBac_MITE | DTC_CACTA | DTC_CACTA_MI | DTC_CACTA_unkno | DTH_Harbinger | DTH_Harbinger_MI | DTM_Mutator | DTM_Mutator_M | DTM_Mutator_ur | DTT_TC1_Marin | DTT_TC1_Marin | DTT_TC1_Marin |
|---|---|---|---|---|---|---|---|---|---|---|---|---|---|---|---|---|
| Hrs | 5805137 | 9180739 | 128232 | 207851 | 0 | 64704 | 77676 | 0 | 1358 | 466093 | 1303359 | 745406 | 0 | 3252559 | 5058361 | 390428 |
| Ass | 717651 | 2743540 | 115516 | 6517 | 0 | 743350 | 295938 | 0 | 47751 | 330021 | 650943 | 254984 | 0 | 1562383 | 1213617 | 35094 |
| Sms | 2333029 | 10156075 | 100168 | 11646 | 0 | 1018631 | 318859 | 0 | 927549 | 879370 | 618708 | 425375 | 0 | 7638627 | 3382309 | 18275 |
| Ppr | 976379 | 5983874 | 155861 | 0 | 0 | 848964 | 163310 | 0 | 0 | 531576 | 161452 | 213076 | 9815 | 4750768 | 5725255 | 243036 |
| Nps | 513094 | 1808776 | 58656 | 17553 | 0 | 34815 | 11559 | 0 | 4733 | 155577 | 130445 | 115198 | 0 | 1264699 | 969897 | 24812 |
| Hga | 9013474 | 16122451 | 449766 | 0 | 0 | 883506 | 416948 | 0 | 1190112 | 1921958 | 396946 | 1109034 | 12591 | 12219399 | 9121980 | 434768 |
| Ona | 598777 | 1313124 | 0 | 0 | 0 | 175320 | 33061 | 0 | 15446 | 103278 | 228612 | 146063 | 0 | 1946176 | 947749 | 55422 |
| Osa | 1185784 | 1766460 | 119363 | 42966 | 1515 | 1053870 | 159016 | 0 | 152563 | 256608 | 660310 | 614439 | 0 | 2702040 | 1596328 | 3626 |
| Aca | 267188 | 3169393 | 60818 | 0 | 0 | 42658 | 19991 | 0 | 15176 | 254010 | 0 | 153012 | 0 | 3152906 | 1610222 | 20324 |
| Hti | 658817 | 7808901 | 246267 | 0 | 0 | 3098786 | 81761 | 31537 | 66987 | 2389714 | 1731289 | 625600 | 0 | 8367052 | 11168504 | 277304 |
| Hls | 3785 | 1181506 | 12975 | 0 | 0 | 46480 | 6305 | 0 | 0 | 169628 | 1082 | 20997 | 0 | 1178879 | 1039752 | 4220 |

| Species | RLB_Bel_Pao | RLC_Copia | RLE_ERV | RLG_Gypsy | RLX | RII_I | RIJ_Jockey | RIL_L1 | RIR_R2 | RIT_RTE | RIX | RPP_Penelope | RST_tRNA | RXX_nLTR | RXX | XXX |
|---|---|---|---|---|---|---|---|---|---|---|---|---|---|---|---|---|
| Hrs | 241106 | 261122 | 0 | 2870741 | 375229 | 263541 | 0 | 45846 | 0 | 0 | 114653 | 63345 | 0 | 0 | 34013 | 10505001 |
| Ass | 488712 | 205861 | 2307 | 4136227 | 1779176 | 442365 | 54491 | 0 | 0 | 1934 | 19736 | 206778 | 9864 | 132615 | 230428 | 4509533 |
| Sms | 1672383 | 988480 | 0 | 13555779 | 2047315 | 196114 | 134068 | 329926 | 24256 | 2146 | 23931 | 2785647 | 18123 | 0 | 1540425 | 13457989 |
| Ppr | 332156 | 303106 | 72079 | 7300648 | 1544105 | 1309510 | 0 | 467593 | 0 | 56211 | 10845 | 1080604 | 33075 | 176075 | 82603 | 9951915 |
| Nps | 101172 | 82768 | 2876 | 1272196 | 239438 | 146988 | 18375 | 78027 | 0 | 15138 | 41075 | 93166 | 6889 | 0 | 169648 | 2430283 |
| Hga | 1273381 | 1032935 | 387339 | 18261731 | 1462066 | 4549894 | 0 | 54910 | 304110 | 297071 | 99708 | 1441387 | 24696 | 45455 | 1682918 | 22549749 |
| Ona | 92790 | 96226 | 6678 | 1169428 | 122584 | 35437 | 0 | 0 | 12600 | 0 | 90527 | 0 | 5331 | 0 | 12933 | 1249940 |
| Osa | 357143 | 205910 | 2655 | 5634042 | 467597 | 281969 | 6065 | 0 | 4438 | 43671 | 26771 | 410347 | 0 | 11480 | 560974 | 4157640 |
| Aca | 209403 | 380521 | 45362 | 8364862 | 858631 | 2527530 | 89495 | 179326 | 3932 | 26528 | 183297 | 979523 | 7130 | 161056 | 1141697 | 4826638 |
| Hti | 672491 | 1110165 | 24661 | 13375981 | 2223733 | 6657373 | 0 | 775284 | 0 | 84463 | 72745 | 2108855 | 0 | 47105 | 1101563 | 12372210 |
| Hls | 159344 | 83563 | 1218 | 1765209 | 185359 | 257552 | 44526 | 370410 | 0 | 0 | 35692 | 559057 | 7996 | 0 | 7187 | 1458328 |

| | | | | | |
|---|---|---|---|---|---|
| Atropacarus striculus | GO:0051384 | biological_process | response to glucocorticoid | 67 | 0,000000001098 |
| Atropacarus striculus | GO:0055085 | biological_process | transmembrane transport | 50 | 0,000000000115 |
| Atropacarus striculus | GO:0007165 | biological_process | signal transduction | 33 | 0 |
| Atropacarus striculus | GO:0046693 | biological_process | sperm storage | 33 | 0,000000046542 |
| Atropacarus striculus | GO:0030245 | biological_process | cellulose catabolic process | 33 | 0,000016531712 |
| Atropacarus striculus | GO:0015074 | biological_process | DNA integration | 27 | 0,000771226669 |
| Atropacarus striculus | GO:0006511 | biological_process | ubiquitin-dependent protein catabolic process | 25 | 0 |
| Atropacarus striculus | GO:0030162 | biological_process | regulation of proteolysis | 25 | 0,000000038120 |
| Atropacarus striculus | GO:0009263 | biological_process | deoxyribonucleotide biosynthetic process | 24 | 0 |
| Atropacarus striculus | GO:0045616 | biological_process | regulation of keratinocyte differentiation | 24 | 0,000000000220 |
| Atropacarus striculus | GO:0010383 | biological_process | cell wall polysaccharide metabolic process | 22 | 0 |
| Atropacarus striculus | GO:0005975 | biological_process | carbohydrate metabolic process | 22 | 0,000693358309 |
| Atropacarus striculus | GO:0006260 | biological_process | DNA replication | 21 | 0,004698021874 |
| Atropacarus striculus | GO:0035093 | biological_process | spermatogenesis, exchange of chromosomal proteins | 20 | 0,000000000939 |
| Atropacarus striculus | GO:0016055 | biological_process | Wnt signaling pathway | 19 | 0 |
| Atropacarus striculus | GO:0006062 | biological_process | sorbitol catabolic process | 19 | 0,000002084406 |
| Atropacarus striculus | GO:0030512 | biological_process | negative regulation of transforming growth factor beta receptor signaling pathway | 18 | 0 |
| Atropacarus striculus | GO:0003964 | molecular_function | RNA-directed DNA polymerase activity | 18 | 0,02026276207 |
| Atropacarus striculus | GO:0006468 | biological_process | protein phosphorylation | 17 | 0,000012152585 |
| Atropacarus striculus | GO:0002376 | biological_process | immune system process | 17 | 0,000037533618 |
| Atropacarus striculus | GO:0019882 | biological_process | antigen processing and presentation | 15 | 0,000000005832 |
| Atropacarus striculus | GO:0032434 | biological_process | regulation of proteasomal ubiquitin-dependent protein catabolic process | 15 | 0,000000013660 |
| Atropacarus striculus | GO:0016266 | biological_process | O-glycan processing | 15 | 0,002472262091 |
| Atropacarus striculus | GO:0046685 | biological_process | response to arsenic-containing substance | 14 | 0,000000000349 |
| Atropacarus striculus | GO:0016114 | biological_process | terpenoid biosynthetic process | 13 | 0,000027831851 |
| Atropacarus striculus | GO:0006355 | biological_process | regulation of transcription, DNA-templated | 12 | 0 |
| Atropacarus striculus | GO:0007275 | biological_process | multicellular organismal development | 12 | 0 |
| Atropacarus striculus | GO:0048512 | biological_process | circadian behavior | 12 | 0,000358706349 |
| Atropacarus striculus | GO:0031098 | biological_process | stress-activated protein kinase signaling cascade | 11 | 0,000398962572 |
| Atropacarus striculus | GO:0006470 | biological_process | protein dephosphorylation | 11 | 0,002839136703 |
| Atropacarus striculus | GO:0004721 | molecular_function | phosphoprotein phosphatase activity | 11 | 0,006938632907 |
| Atropacarus striculus | GO:0042761 | biological_process | very long-chain fatty acid biosynthetic process | 10 | 0,000047604203 |
| Atropacarus striculus | GO:0051260 | biological_process | protein homooligomerization | 10 | 0,000641097944 |
| Atropacarus striculus | GO:0007283 | biological_process | spermatogenesis | 9 | 0 |
| Atropacarus striculus | GO:0008380 | biological_process | RNA splicing | 9 | 0 |
| Atropacarus striculus | GO:0006633 | biological_process | fatty acid biosynthetic process | 7 | 0,000000161764 |
| Atropacarus striculus | GO:0006313 | biological_process | transposition, DNA-mediated | 7 | 0,000001826972 |

| | | | | | |
|---|---|---|---|---|---|
| Atropacarus striculus | GO:0007286 | biological_process | spermatid development | 7 | 0,000666964250 |
| Atropacarus striculus | GO:0001669 | cellular_component | acrosomal vesicle | 7 | 0,00125193187 |
| Atropacarus striculus | GO:0010193 | biological_process | response to ozone | 7 | 0,003467409466 |
| Atropacarus striculus | GO:0007179 | biological_process | transforming growth factor beta receptor signaling pathway | 7 | 0,01542198636 |
| Atropacarus striculus | GO:0009617 | biological_process | response to bacterium | 6 | 0 |
| Atropacarus striculus | GO:0006979 | biological_process | response to oxidative stress | 6 | 0 |
| Atropacarus striculus | GO:0006914 | biological_process | autophagy | 6 | 0,005901474595 |
| Atropacarus striculus | GO:0008218 | biological_process | bioluminescence | 6 | 0,01605897602 |
| Atropacarus striculus | GO:0006970 | biological_process | response to osmotic stress | 6 | 0,01958714713 |
| Atropacarus striculus | GO:0006412 | biological_process | translation | 5 | 0 |
| Atropacarus striculus | GO:0016021 | cellular_component | integral component of membrane | 5 | 0 |
| Atropacarus striculus | GO:0046872 | molecular_function | metal ion binding | 5 | 0 |
| Atropacarus striculus | GO:0045893 | biological_process | positive regulation of transcription, DNA-templated | 5 | 0,000001464509 |
| Atropacarus striculus | GO:0042060 | biological_process | wound healing | 5 | 0,000001658740 |
| Atropacarus striculus | GO:0051131 | biological_process | chaperone-mediated protein complex assembly | 5 | 0,000465898599 |
| Atropacarus striculus | GO:0031638 | biological_process | zymogen activation | 5 | 0,008954121717 |
| Atropacarus striculus | GO:0031149 | biological_process | sorocarp stalk cell differentiation | 4 | 0,000105021436 |
| Atropacarus striculus | GO:0009742 | biological_process | brassinosteroid mediated signaling pathway | 4 | 0,000105021436 |
| Atropacarus striculus | GO:0007586 | biological_process | digestion | 4 | 0,000389918641 |
| Atropacarus striculus | GO:0075732 | biological_process | viral penetration into host nucleus | 4 | 0,000638127585 |
| Atropacarus striculus | GO:0009213 | biological_process | pyrimidine deoxyribonucleoside triphosphate catabolic process | 4 | 0,00118902427 |
| Atropacarus striculus | GO:0007166 | biological_process | cell surface receptor signaling pathway | 4 | 0,002351869065 |
| Atropacarus striculus | GO:0007601 | biological_process | visual perception | 3 | 0 |
| Atropacarus striculus | GO:0006357 | biological_process | regulation of transcription from RNA polymerase II promoter | 3 | 0 |
| Atropacarus striculus | GO:0009636 | biological_process | response to toxic substance | 3 | 0,000000031027 |
| Atropacarus striculus | GO:0007605 | biological_process | sensory perception of sound | 3 | 0,000000865443 |
| Atropacarus striculus | GO:0000723 | biological_process | telomere maintenance | 3 | 0,000039280125 |
| Atropacarus striculus | GO:0055114 | biological_process | oxidation-reduction process | 3 | 0,000044508988 |
| Atropacarus striculus | GO:0004867 | molecular_function | serine-type endopeptidase inhibitor activity | 3 | 0,001422485727 |
| Atropacarus striculus | GO:0000011 | biological_process | vacuole inheritance | 3 | 0,002853685922 |
| Atropacarus striculus | GO:0072431 | biological_process | signal transduction involved in mitotic G1 DNA damage checkpoint | 3 | 0,005315983497 |
| Atropacarus striculus | GO:0001756 | biological_process | somitogenesis | 3 | 0,0148743593 |
| Atropacarus striculus | GO:0045087 | biological_process | innate immune response | 3 | 0,01809621118 |
| Atropacarus striculus | GO:0016192 | biological_process | vesicle-mediated transport | 2 | 0 |
| Atropacarus striculus | GO:0006805 | biological_process | xenobiotic metabolic process | 2 | 0 |
| Atropacarus striculus | GO:0010025 | biological_process | wax biosynthetic process | 2 | 0,000001282339 |
| Atropacarus striculus | GO:0006694 | biological_process | steroid biosynthetic process | 2 | 0,000058199881 |

| | | | | | |
|---|---|---|---|---|---|
| Atropacarus striculus | GO:0007155 | biological_process | cell adhesion | 2 | 0,000058199881 |
| Atropacarus striculus | GO:0048511 | biological_process | rhythmic process | 2 | 0,000307717516 |
| Atropacarus striculus | GO:0006417 | biological_process | regulation of translation | 2 | 0,000736116899 |
| Atropacarus striculus | GO:0071625 | biological_process | vocalization behavior | 2 | 0,002065291246 |
| Atropacarus striculus | GO:0051301 | biological_process | cell division | 2 | 0,002468968651 |
| Atropacarus striculus | GO:0007416 | biological_process | synapse assembly | 2 | 0,003951578101 |
| Atropacarus striculus | GO:2000377 | biological_process | regulation of reactive oxygen species metabolic process | 2 | 0,0136978678 |

| | | | | |
|---|---|---|---|---|
| Steganacarus magnus | GO:0055085 | biological_process | transmembrane transport | 329 |
| Steganacarus magnus | GO:0032100 | biological_process | positive regulation of appetite | 218 |
| Steganacarus magnus | GO:0042908 | biological_process | xenobiotic transport | 97 |
| Steganacarus magnus | GO:0016020 | cellular_component | membrane | 80 |
| Steganacarus magnus | GO:0016301 | molecular_function | kinase activity | 78 |
| Steganacarus magnus | GO:0005975 | biological_process | carbohydrate metabolic process | 72 |
| Steganacarus magnus | GO:0007283 | biological_process | spermatogenesis | 71 |
| Steganacarus magnus | GO:0006805 | biological_process | xenobiotic metabolic process | 71 |
| Steganacarus magnus | GO:0007601 | biological_process | visual perception | 68 |
| Steganacarus magnus | GO:0003081 | biological_process | regulation of systemic arterial blood pressure by renin-angiotensin | 62 |
| Steganacarus magnus | GO:0007165 | biological_process | signal transduction | 61 |
| Steganacarus magnus | GO:0030223 | biological_process | neutrophil differentiation | 58 |
| Steganacarus magnus | GO:0008270 | molecular_function | zinc ion binding | 55 |
| Steganacarus magnus | GO:0006508 | biological_process | proteolysis | 53 |
| Steganacarus magnus | GO:0000723 | biological_process | telomere maintenance | 52 |
| Steganacarus magnus | GO:0052697 | biological_process | xenobiotic glucuronidation | 46 |
| Steganacarus magnus | GO:0006486 | biological_process | protein glycosylation | 45 |
| Steganacarus magnus | GO:0007275 | biological_process | multicellular organismal development | 43 |
| Steganacarus magnus | GO:0019284 | biological_process | L-methionine biosynthetic process from S-adenosylmethionine | 42 |
| Steganacarus magnus | GO:0009617 | biological_process | response to bacterium | 42 |
| Steganacarus magnus | GO:0006706 | biological_process | steroid catabolic process | 42 |
| Steganacarus magnus | GO:0001507 | biological_process | acetylcholine catabolic process in synaptic cleft | 41 |
| Steganacarus magnus | GO:0004252 | molecular_function | serine-type endopeptidase activity | 37 |
| Steganacarus magnus | GO:0016705 | molecular_function | oxidoreductase activity, acting on paired donors, with incorporation or reduction of molecular oxygen | 37 |
| Steganacarus magnus | GO:0010224 | biological_process | response to UV-B | 36 |
| Steganacarus magnus | GO:0042135 | biological_process | neurotransmitter catabolic process | 36 |
| Steganacarus magnus | GO:0010842 | biological_process | retina layer formation | 34 |
| Steganacarus magnus | GO:0005506 | molecular_function | iron ion binding | 34 |
| Steganacarus magnus | GO:0004497 | molecular_function | monooxygenase activity | 33 |
| Steganacarus magnus | GO:0006814 | biological_process | sodium ion transport | 32 |
| Steganacarus magnus | GO:0046872 | molecular_function | metal ion binding | 29 |
| Steganacarus magnus | GO:0043252 | biological_process | sodium-independent organic anion transport | 28 |
| Steganacarus magnus | GO:0009888 | biological_process | tissue development | 28 |
| Steganacarus magnus | GO:0042302 | molecular_function | structural constituent of cuticle | 28 |
| Steganacarus magnus | GO:0009636 | biological_process | response to toxic substance | 26 |
| Steganacarus magnus | GO:0033609 | biological_process | oxalate metabolic process | 25 |
| Steganacarus magnus | GO:0014832 | biological_process | urinary bladder smooth muscle contraction | 25 |

| | | | | |
|---|---|---|---|---|
| Steganacarus magnus | GO:0016266 | biological_process | O-glycan processing | 25 |
| Steganacarus magnus | GO:0044245 | biological_process | polysaccharide digestion | 24 |
| Steganacarus magnus | GO:0071555 | biological_process | cell wall organization | 24 |
| Steganacarus magnus | GO:0015020 | molecular_function | glucuronosyltransferase activity | 23 |
| Steganacarus magnus | GO:0045493 | biological_process | xylan catabolic process | 23 |
| Steganacarus magnus | GO:0006511 | biological_process | ubiquitin-dependent protein catabolic process | 22 |
| Steganacarus magnus | GO:0006412 | biological_process | translation | 21 |
| Steganacarus magnus | GO:0016567 | biological_process | protein ubiquitination | 21 |
| Steganacarus magnus | GO:0003677 | molecular_function | DNA binding | 20 |
| Steganacarus magnus | GO:0016192 | biological_process | vesicle-mediated transport | 19 |
| Steganacarus magnus | GO:0006468 | biological_process | protein phosphorylation | 19 |
| Steganacarus magnus | GO:0008210 | biological_process | estrogen metabolic process | 19 |
| Steganacarus magnus | GO:0006069 | biological_process | ethanol oxidation | 19 |
| Steganacarus magnus | GO:0006355 | biological_process | regulation of transcription, DNA-templated | 18 |
| Steganacarus magnus | GO:0016042 | biological_process | lipid catabolic process | 18 |
| Steganacarus magnus | GO:0008146 | molecular_function | sulfotransferase activity | 17 |
| Steganacarus magnus | GO:0031098 | biological_process | stress-activated protein kinase signaling cascade | 17 |
| Steganacarus magnus | GO:0051923 | biological_process | sulfation | 17 |
| Steganacarus magnus | GO:0031030 | biological_process | negative regulation of septation initiation signaling | 16 |
| Steganacarus magnus | GO:0021591 | biological_process | ventricular system development | 16 |
| Steganacarus magnus | GO:0008194 | molecular_function | UDP-glycosyltransferase activity | 16 |
| Steganacarus magnus | GO:0006090 | biological_process | pyruvate metabolic process | 15 |
| Steganacarus magnus | GO:0006810 | biological_process | transport | 15 |
| Steganacarus magnus | GO:0042742 | biological_process | defense response to bacterium | 15 |
| Steganacarus magnus | GO:0005983 | biological_process | starch catabolic process | 14 |
| Steganacarus magnus | GO:0001963 | biological_process | synaptic transmission, dopaminergic | 14 |
| Steganacarus magnus | GO:0046355 | biological_process | mannan catabolic process | 14 |
| Steganacarus magnus | GO:0004970 | molecular_function | ionotropic glutamate receptor activity | 14 |
| Steganacarus magnus | GO:0015074 | biological_process | DNA integration | 14 |
| Steganacarus magnus | GO:0006694 | biological_process | steroid biosynthetic process | 14 |
| Steganacarus magnus | GO:0045944 | biological_process | positive regulation of transcription from RNA polymerase II promoter | 13 |
| Steganacarus magnus | GO:1901639 | biological_process | XDP catabolic process | 13 |
| Steganacarus magnus | GO:0035092 | biological_process | sperm chromatin condensation | 13 |
| Steganacarus magnus | GO:0001649 | biological_process | osteoblast differentiation | 13 |
| Steganacarus magnus | GO:0036324 | biological_process | vascular endothelial growth factor receptor-2 signaling pathway | 12 |
| Steganacarus magnus | GO:0007419 | biological_process | ventral cord development | 12 |
| Steganacarus magnus | GO:0006941 | biological_process | striated muscle contraction | 12 |

| | | | | |
|---|---|---|---|---|
| Steganacarus magnus | GO:0051603 | biological_process | proteolysis involved in cellular protein catabolic process | 12 |
| Steganacarus magnus | GO:0035249 | biological_process | synaptic transmission, glutamatergic | 12 |
| Steganacarus magnus | GO:0042572 | biological_process | retinol metabolic process | 12 |
| Steganacarus magnus | GO:0006749 | biological_process | glutathione metabolic process | 12 |
| Steganacarus magnus | GO:0006470 | biological_process | protein dephosphorylation | 11 |
| Steganacarus magnus | GO:0045490 | biological_process | pectin catabolic process | 11 |
| Steganacarus magnus | GO:0034976 | biological_process | response to endoplasmic reticulum stress | 11 |
| Steganacarus magnus | GO:0008277 | biological_process | regulation of G-protein coupled receptor protein signaling pathway | 11 |
| Steganacarus magnus | GO:0019233 | biological_process | sensory perception of pain | 11 |
| Steganacarus magnus | GO:0003309 | biological_process | type B pancreatic cell differentiation | 11 |
| Steganacarus magnus | GO:0005634 | cellular_component | nucleus | 10 |
| Steganacarus magnus | GO:0006030 | biological_process | chitin metabolic process | 10 |
| Steganacarus magnus | GO:0010569 | biological_process | regulation of double-strand break repair via homologous recombination | 10 |
| Steganacarus magnus | GO:0008361 | biological_process | regulation of cell size | 10 |
| Steganacarus magnus | GO:0047935 | molecular_function | glucose 1-dehydrogenase (NADP+) activity | 10 |
| Steganacarus magnus | GO:0070212 | biological_process | protein poly-ADP-ribosylation | 10 |
| Steganacarus magnus | GO:0007608 | biological_process | sensory perception of smell | 10 |
| Steganacarus magnus | GO:0016021 | cellular_component | integral component of membrane | 9 |
| Steganacarus magnus | GO:0008380 | biological_process | RNA splicing | 9 |
| Steganacarus magnus | GO:0019062 | biological_process | virion attachment to host cell | 9 |
| Steganacarus magnus | GO:0016488 | biological_process | farnesol catabolic process | 9 |
| Steganacarus magnus | GO:0006644 | biological_process | phospholipid metabolic process | 9 |
| Steganacarus magnus | GO:0009405 | biological_process | pathogenesis | 9 |
| Steganacarus magnus | GO:0042626 | molecular_function | ATPase activity, coupled to transmembrane movement of substances | 8 |
| Steganacarus magnus | GO:0008241 | molecular_function | peptidyl-dipeptidase activity | 8 |
| Steganacarus magnus | GO:0016117 | biological_process | carotenoid biosynthetic process | 8 |
| Steganacarus magnus | GO:0009408 | biological_process | response to heat | 8 |
| Steganacarus magnus | GO:0048010 | biological_process | vascular endothelial growth factor receptor signaling pathway | 7 |
| Steganacarus magnus | GO:0045892 | biological_process | negative regulation of transcription, DNA-templated | 6 |
| Steganacarus magnus | GO:0055114 | biological_process | oxidation-reduction process | 6 |
| Steganacarus magnus | GO:0007399 | biological_process | nervous system development | 6 |
| Steganacarus magnus | GO:0007584 | biological_process | response to nutrient | 6 |
| Steganacarus magnus | GO:0001501 | biological_process | skeletal system development | 6 |
| Steganacarus magnus | GO:0001894 | biological_process | tissue homeostasis | 6 |
| Steganacarus magnus | GO:0030162 | biological_process | regulation of proteolysis | 6 |
| Steganacarus magnus | GO:0007605 | biological_process | sensory perception of sound | 5 |
| Steganacarus magnus | GO:0007155 | biological_process | cell adhesion | 5 |

| | | | | |
|---|---|---|---|---|
| Steganacarus magnus | GO:0031647 | biological_process | regulation of protein stability | 5 |
| Steganacarus magnus | GO:0016887 | molecular_function | ATPase activity | 5 |
| Steganacarus magnus | GO:0032197 | biological_process | transposition, RNA-mediated | 5 |
| Steganacarus magnus | GO:0036323 | biological_process | vascular endothelial growth factor receptor-1 signaling pathway | 5 |
| Steganacarus magnus | GO:0001659 | biological_process | temperature homeostasis | 5 |
| Steganacarus magnus | GO:0043129 | biological_process | surfactant homeostasis | 5 |
| Steganacarus magnus | GO:0035556 | biological_process | intracellular signal transduction | 5 |
| Steganacarus magnus | GO:0016055 | biological_process | Wnt signaling pathway | 4 |
| Steganacarus magnus | GO:0015031 | biological_process | protein transport | 4 |
| Steganacarus magnus | GO:0007179 | biological_process | transforming growth factor beta receptor signaling pathway | 4 |
| Steganacarus magnus | GO:0006641 | biological_process | triglyceride metabolic process | 4 |
| Steganacarus magnus | GO:0008080 | molecular_function | N-acetyltransferase activity | 4 |
| Steganacarus magnus | GO:0005412 | molecular_function | glucose:sodium symporter activity | 4 |
| Steganacarus magnus | GO:0008240 | molecular_function | tripeptidyl-peptidase activity | 4 |
| Steganacarus magnus | GO:0060074 | biological_process | synapse maturation | 4 |
| Steganacarus magnus | GO:0007416 | biological_process | synapse assembly | 4 |
| Steganacarus magnus | GO:0009845 | biological_process | seed germination | 4 |
| Steganacarus magnus | GO:0016032 | biological_process | viral process | 3 |
| Steganacarus magnus | GO:0051384 | biological_process | response to glucocorticoid | 3 |
| Steganacarus magnus | GO:0006313 | biological_process | transposition, DNA-mediated | 3 |
| Steganacarus magnus | GO:0006099 | biological_process | tricarboxylic acid cycle | 3 |
| Steganacarus magnus | GO:0006457 | biological_process | protein folding | 3 |
| Steganacarus magnus | GO:0035220 | biological_process | wing disc development | 3 |
| Steganacarus magnus | GO:0018279 | biological_process | protein N-linked glycosylation via asparagine | 3 |
| Steganacarus magnus | GO:0023021 | biological_process | termination of signal transduction | 3 |
| Steganacarus magnus | GO:0004712 | molecular_function | protein serine/threonine/tyrosine kinase activity | 3 |
| Steganacarus magnus | GO:0072164 | biological_process | mesonephric tubule development | 3 |
| Steganacarus magnus | GO:0042981 | biological_process | regulation of apoptotic process | 3 |
| Steganacarus magnus | GO:0010951 | biological_process | negative regulation of endopeptidase activity | 3 |
| Steganacarus magnus | GO:0050661 | molecular_function | NADP binding | 3 |
| Steganacarus magnus | GO:0031175 | biological_process | neuron projection development | 3 |
| Steganacarus magnus | GO:0006357 | biological_process | regulation of transcription from RNA polymerase II promoter | 2 |
| Steganacarus magnus | GO:0006979 | biological_process | response to oxidative stress | 2 |
| Steganacarus magnus | GO:0042060 | biological_process | wound healing | 2 |
| Steganacarus magnus | GO:0005576 | cellular_component | extracellular region | 2 |
| Steganacarus magnus | GO:0010025 | biological_process | wax biosynthetic process | 2 |
| Steganacarus magnus | GO:0007264 | biological_process | small GTPase mediated signal transduction | 2 |

| | | | | |
|---|---|---|---|---|
| Steganacarus magnus | GO:0007186 | biological_process | G-protein coupled receptor signaling pathway | 2 |
| Steganacarus magnus | GO:0006413 | biological_process | translational initiation | 2 |
| Steganacarus magnus | GO:0031638 | biological_process | zymogen activation | 2 |
| Steganacarus magnus | GO:0006260 | biological_process | DNA replication | 2 |
| Steganacarus magnus | GO:0006680 | biological_process | glucosylceramide catabolic process | 2 |
| Steganacarus magnus | GO:0006915 | biological_process | apoptotic process | 2 |
| Steganacarus magnus | GO:0051301 | biological_process | cell division | 2 |
| Steganacarus magnus | GO:0008168 | molecular_function | methyltransferase activity | 2 |
| Steganacarus magnus | GO:0016485 | biological_process | protein processing | 2 |
| Steganacarus magnus | GO:0032465 | biological_process | regulation of cytokinesis | 2 |
| Steganacarus magnus | GO:0019216 | biological_process | regulation of lipid metabolic process | 2 |
| Steganacarus magnus | GO:0001505 | biological_process | regulation of neurotransmitter levels | 2 |
| Steganacarus magnus | GO:0008039 | biological_process | synaptic target recognition | 2 |
| Steganacarus magnus | GO:0015693 | biological_process | magnesium ion transport | 2 |
| Steganacarus magnus | GO:0042770 | biological_process | signal transduction in response to DNA damage | 2 |
| Steganacarus magnus | GO:0019285 | biological_process | glycine betaine biosynthetic process from choline | 2 |
| Steganacarus magnus | GO:0032880 | biological_process | regulation of protein localization | 2 |
| Steganacarus magnus | GO:0051290 | biological_process | protein heterotetramerization | 2 |
| Steganacarus magnus | GO:0001516 | biological_process | prostaglandin biosynthetic process | 2 |
| Steganacarus magnus | GO:0097264 | biological_process | self proteolysis | 2 |
| Steganacarus magnus | GO:0071357 | biological_process | cellular response to type I interferon | 2 |
| Steganacarus magnus | GO:0042802 | molecular_function | identical protein binding | 2 |
| Steganacarus magnus | GO:0070445 | biological_process | regulation of oligodendrocyte progenitor proliferation | 2 |
| Steganacarus magnus | GO:0007338 | biological_process | single fertilization | 2 |
| Steganacarus magnus | GO:0043434 | biological_process | response to peptide hormone | 2 |
| Steganacarus magnus | GO:0051438 | biological_process | regulation of ubiquitin-protein transferase activity | 2 |

| |
|---|
| 0 |
| 0 |
| 0 |
| 0 |
| 0 |
| 0,01437208568 |
| 0 |
| 0,000148951092 |
| 0,000384620661 |
| 0 |
| 0 |
| 0 |
| 0,001503670081 |
| 0 |
| 0,000000086156 |
| 0 |
| 0,001275609278 |
| 0,000000004561 |
| 0,000000245124 |
| 0,000160976472 |
| 0,000387187567 |
| 0,00183790742 |
| 0,000018380603 |
| 0,01290864965 |
| 0,000045451305 |
| 0,01764613371 |
| 0,00000000114[illegible] |
| 0,01317518675 |
| 0,001032734524 |
| 0,00011693307[illegible] |
| 0 |
| 0,000008407710 |
| 0,00017924918[illegible] |
| 0,007318529168 |
| 0,008471158982 |
| 0 |
| 0,000000000268 |

| |
|---|
| 0,000047766956 |
| 0 |
| 0,000665076125 |
| 0,000000001871 |
| 0,000022500054 |
| 0 |
| 0 |
| 0 |
| 0,000027285675 |
| 0 |
| 0 |
| 0 |
| 0,001426612632 |
| 0 |
| 0,000000074758 |
| 0,000000023306 |
| 0,000004822796 |
| 0,001627784154 |
| 0,000000062737 |
| 0,002289332617 |
| 0,007647494059 |
| 0,00001591144[illegible] |
| 0,000068631002 |
| 0,004198705759 |
| 0,000000009462 |
| 0,000618512522 |
| 0,002548365554 |
| 0,00749524369 |
| 0,007893388153 |
| 0,01816415418 |
| 0 |
| 0,000000001454 |
| 0,00000011008[illegible] |
| 0,000036238554 |
| 0,000000577838 |
| 0,000015204777 |
| 0,000090387590 |

| |
|---|
| 0,000121191302[illegible] |
| 0,000543542270 |
| 0,009907269798 |
| 0,02114915949 |
| 0,000000295421 |
| 0,000001558985 |
| 0,000002017880 |
| 0,000303689445 |
| 0,006298085924 |
| 0,01429623726 |
| 0,000000003491 |
| 0,000024084763 |
| 0,000289684127 |
| 0,002016200759 |
| 0,002016200759 |
| 0,005501253023 |
| 0,01548840374 |
| 0 |
| 0 |
| 0,000000019807 |
| 0,000364753251 |
| 0,004975207392 |
| 0,007716044486 |
| 0,000007414958[illegible] |
| 0,000650902296 |
| 0,01147794698 |
| 0,01158997083 |
| 0,01061676763 |
| 0,000000127078 |
| 0,000001092198 |
| 0,000011483718[illegible] |
| 0,007722251579 |
| 0,01722969887 |
| 0,02081759982 |
| 0,02081759982 |
| 0,000000000379 |
| 0,000004787649 |

| |
|---|
| 0,000079315428 |
| 0,000357739617 |
| 0,001268956434 |
| 0,005613825037 |
| 0,00951551458 |
| 0,01504285011 |
| 0,0155786115 |
| 0 |
| 0 |
| 0,000000702321 |
| 0,000072108305 |
| 0,000145647761 |
| 0,000145647761 |
| 0,000145647761 |
| 0,000648662409 |
| 0,000648849409 |
| 0,0213667219 |
| 0 |
| 0 |
| 0 |
| 0 |
| 0,000000000252 |
| 0,000002612644 |
| 0,000066973045 |
| 0,000120573440 |
| 0,001330039987 |
| 0,001330039987 |
| 0,002008091104 |
| 0,003128896697 |
| 0,003727010499 |
| 0,0102937665 |
| 0 |
| 0 |
| 0 |
| 0 |
| 0 |
| 0,000000000366 |

0,000000000581
0,000000000731
0,000000185494
0,000000185494
0,000001200535
0,000010347008
0,000012796909
0,000024125177
0,000084309097
0,000172920980
0,000259583554
0,000287185629
0,000706043868
0,000779348205
0,001046585094
0,001154064187
0,001402134305
0,001873771534
0,002497406162
0,006979534331
0,01096452402
0,01198515549
0,01211990983
0,01429979679
0,01560811153
0,0239863613

| | | | | | |
|---|---|---|---|---|---|
| Hypochthonius rufulus | GO:0055085 | biological_process | transmembrane transport | 242 | 0 |
| Hypochthonius rufulus | GO:0016301 | molecular_function | kinase activity | 106 | 0 |
| Hypochthonius rufulus | GO:0006313 | biological_process | transposition, DNA-mediated | 88 | 0 |
| Hypochthonius rufulus | GO:0007283 | biological_process | spermatogenesis | 76 | 0 |
| Hypochthonius rufulus | GO:0008218 | biological_process | bioluminescence | 69 | 0 |
| Hypochthonius rufulus | GO:0006805 | biological_process | xenobiotic metabolic process | 66 | 0,000464177876 |
| Hypochthonius rufulus | GO:0051384 | biological_process | response to glucocorticoid | 65 | 0,003588415132 |
| Hypochthonius rufulus | GO:0016266 | biological_process | O-glycan processing | 59 | 0 |
| Hypochthonius rufulus | GO:0008194 | molecular_function | UDP-glycosyltransferase activity | 58 | 0 |
| Hypochthonius rufulus | GO:0003677 | molecular_function | DNA binding | 57 | 0,000736301771 |
| Hypochthonius rufulus | GO:0006646 | biological_process | phosphatidylethanolamine biosynthetic process | 56 | 0,000000036665 |
| Hypochthonius rufulus | GO:0052697 | biological_process | xenobiotic glucuronidation | 55 | 0 |
| Hypochthonius rufulus | GO:0004364 | molecular_function | glutathione transferase activity | 54 | 0 |
| Hypochthonius rufulus | GO:0003964 | molecular_function | RNA-directed DNA polymerase activity | 46 | 0,000000000280 |
| Hypochthonius rufulus | GO:0009617 | biological_process | response to bacterium | 46 | 0,003456368132 |
| Hypochthonius rufulus | GO:0055129 | biological_process | L-proline biosynthetic process | 44 | 0 |
| Hypochthonius rufulus | GO:0006706 | biological_process | steroid catabolic process | 44 | 0,000034712304 |
| Hypochthonius rufulus | GO:0030435 | biological_process | sporulation resulting in formation of a cellular spore | 40 | 0,000030385630 |
| Hypochthonius rufulus | GO:0006589 | biological_process | octopamine biosynthetic process | 37 | 0 |
| Hypochthonius rufulus | GO:0004497 | molecular_function | monooxygenase activity | 37 | 0,000021443420 |
| Hypochthonius rufulus | GO:0009311 | biological_process | oligosaccharide metabolic process | 36 | 0 |
| Hypochthonius rufulus | GO:0006517 | biological_process | protein deglycosylation | 34 | 0,000010143027 |
| Hypochthonius rufulus | GO:0042178 | biological_process | xenobiotic catabolic process | 32 | 0,000000000325 |
| Hypochthonius rufulus | GO:0004222 | molecular_function | metalloendopeptidase activity | 31 | 0,000059537141 |
| Hypochthonius rufulus | GO:0016491 | molecular_function | oxidoreductase activity | 31 | 0,000547783128 |
| Hypochthonius rufulus | GO:0015020 | molecular_function | glucuronosyltransferase activity | 30 | 0 |
| Hypochthonius rufulus | GO:0004518 | molecular_function | nuclease activity | 30 | 0 |
| Hypochthonius rufulus | GO:0006013 | biological_process | mannose metabolic process | 30 | 0,000000024739 |
| Hypochthonius rufulus | GO:0006633 | biological_process | fatty acid biosynthetic process | 30 | 0,003670020339 |
| Hypochthonius rufulus | GO:0006511 | biological_process | ubiquitin-dependent protein catabolic process | 29 | 0 |
| Hypochthonius rufulus | GO:0005525 | molecular_function | GTP binding | 29 | 0,000003924289 |
| Hypochthonius rufulus | GO:0046548 | biological_process | retinal rod cell development | 28 | 0 |
| Hypochthonius rufulus | GO:0046872 | molecular_function | metal ion binding | 27 | 0 |
| Hypochthonius rufulus | GO:0071357 | biological_process | cellular response to type I interferon | 27 | 0,000000000230 |
| Hypochthonius rufulus | GO:0001516 | biological_process | prostaglandin biosynthetic process | 27 | 0,000000042122 |
| Hypochthonius rufulus | GO:0048864 | biological_process | stem cell development | 25 | 0 |
| Hypochthonius rufulus | GO:0008270 | molecular_function | zinc ion binding | 25 | 0 |

| | | | | | |
|---|---|---|---|---|---|
| Hypochthonius rufulus | GO:0006979 | biological_process | response to oxidative stress | 25 | 0,000000015859 |
| Hypochthonius rufulus | GO:0006508 | biological_process | proteolysis | 24 | 0 |
| Hypochthonius rufulus | GO:0010811 | biological_process | positive regulation of cell-substrate adhesion | 24 | 0,000000009059 |
| Hypochthonius rufulus | GO:0009058 | biological_process | biosynthetic process | 24 | 0,000229603856 |
| Hypochthonius rufulus | GO:0003081 | biological_process | regulation of systemic arterial blood pressure by renin-angiotensin | 23 | 0,000001766657 |
| Hypochthonius rufulus | GO:0009268 | biological_process | response to pH | 23 | 0,000006824788 |
| Hypochthonius rufulus | GO:0045599 | biological_process | negative regulation of fat cell differentiation | 23 | 0,000006824788 |
| Hypochthonius rufulus | GO:0006468 | biological_process | protein phosphorylation | 22 | 0,000000001204 |
| Hypochthonius rufulus | GO:0051260 | biological_process | protein homooligomerization | 21 | 0,002024151605 |
| Hypochthonius rufulus | GO:0000272 | biological_process | polysaccharide catabolic process | 20 | 0,000103150740 |
| Hypochthonius rufulus | GO:0006814 | biological_process | sodium ion transport | 19 | 0,000000008477 |
| Hypochthonius rufulus | GO:0006062 | biological_process | sorbitol catabolic process | 19 | 0,000541034528 |
| Hypochthonius rufulus | GO:0030512 | biological_process | negative regulation of transforming growth factor beta receptor signaling pathway | 18 | 0,000000001254 |
| Hypochthonius rufulus | GO:0009636 | biological_process | response to toxic substance | 18 | 0,000196289125 |
| Hypochthonius rufulus | GO:0001507 | biological_process | acetylcholine catabolic process in synaptic cleft | 18 | 0,0195214201 |
| Hypochthonius rufulus | GO:0007165 | biological_process | signal transduction | 17 | 0 |
| Hypochthonius rufulus | GO:0016117 | biological_process | carotenoid biosynthetic process | 17 | 0,000000000996 |
| Hypochthonius rufulus | GO:0010025 | biological_process | wax biosynthetic process | 17 | 0,009713238381 |
| Hypochthonius rufulus | GO:0016021 | cellular_component | integral component of membrane | 16 | 0 |
| Hypochthonius rufulus | GO:0016567 | biological_process | protein ubiquitination | 16 | 0 |
| Hypochthonius rufulus | GO:0006090 | biological_process | pyruvate metabolic process | 16 | 0,000001700891 |
| Hypochthonius rufulus | GO:0006486 | biological_process | protein glycosylation | 15 | 0 |
| Hypochthonius rufulus | GO:0009168 | biological_process | purine ribonucleoside monophosphate biosynthetic process | 15 | 0,000000000812 |
| Hypochthonius rufulus | GO:0042060 | biological_process | wound healing | 15 | 0,000023051898 |
| Hypochthonius rufulus | GO:1902477 | biological_process | regulation of defense response to bacterium, incompatible interaction | 14 | 0,000000000108 |
| Hypochthonius rufulus | GO:0009698 | biological_process | phenylpropanoid metabolic process | 14 | 0,003970629664 |
| Hypochthonius rufulus | GO:0007021 | biological_process | tubulin complex assembly | 14 | 0,01617171822 |
| Hypochthonius rufulus | GO:0006355 | biological_process | regulation of transcription, DNA-templated | 13 | 0 |
| Hypochthonius rufulus | GO:0033089 | biological_process | positive regulation of T cell differentiation in thymus | 13 | 0,000000002237 |
| Hypochthonius rufulus | GO:0006152 | biological_process | purine nucleoside catabolic process | 13 | 0,000000005741 |
| Hypochthonius rufulus | GO:0051603 | biological_process | proteolysis involved in cellular protein catabolic process | 13 | 0,000681961230 |
| Hypochthonius rufulus | GO:0009099 | biological_process | valine biosynthetic process | 12 | 0,000001140635 |
| Hypochthonius rufulus | GO:0007601 | biological_process | visual perception | 11 | 0 |
| Hypochthonius rufulus | GO:0072089 | biological_process | stem cell proliferation | 11 | 0,000000000162 |
| Hypochthonius rufulus | GO:0042110 | biological_process | T cell activation | 11 | 0,000074142566 |
| Hypochthonius rufulus | GO:0010383 | biological_process | cell wall polysaccharide metabolic process | 11 | 0,000356221314 |
| Hypochthonius rufulus | GO:0035249 | biological_process | synaptic transmission, glutamatergic | 11 | 0,000511626636 |

| | | | | | |
|---|---|---|---|---|---|
| Hypochthonius rufulus | GO:0006470 | biological_process | protein dephosphorylation | 10 | 0,000000321562 |
| Hypochthonius rufulus | GO:0000723 | biological_process | telomere maintenance | 10 | 0,000292252174 |
| Hypochthonius rufulus | GO:0031647 | biological_process | regulation of protein stability | 10 | 0,01471875101 |
| Hypochthonius rufulus | GO:0007608 | biological_process | sensory perception of smell | 10 | 0,02302210035 |
| Hypochthonius rufulus | GO:0010869 | biological_process | regulation of receptor biosynthetic process | 9 | 0,000044422495 |
| Hypochthonius rufulus | GO:0016999 | biological_process | antibiotic metabolic process | 9 | 0,003269770047 |
| Hypochthonius rufulus | GO:0043171 | biological_process | peptide catabolic process | 9 | 0,004914491298 |
| Hypochthonius rufulus | GO:0044179 | biological_process | hemolysis in other organism | 9 | 0,01329301167 |
| Hypochthonius rufulus | GO:0006367 | biological_process | transcription initiation from RNA polymerase II promoter | 8 | 0,000004122165 |
| Hypochthonius rufulus | GO:0016032 | biological_process | viral process | 7 | 0 |
| Hypochthonius rufulus | GO:0002175 | biological_process | protein localization to paranode region of axon | 7 | 0,000119339400 |
| Hypochthonius rufulus | GO:1901639 | biological_process | XDP catabolic process | 7 | 0,001281445154 |
| Hypochthonius rufulus | GO:0016887 | molecular_function | ATPase activity | 7 | 0,004848775091 |
| Hypochthonius rufulus | GO:0030321 | biological_process | transepithelial chloride transport | 7 | 0,01334628392 |
| Hypochthonius rufulus | GO:0009699 | biological_process | phenylpropanoid biosynthetic process | 6 | 0,000008111484 |
| Hypochthonius rufulus | GO:0007586 | biological_process | digestion | 6 | 0,000009121360 |
| Hypochthonius rufulus | GO:0004674 | molecular_function | protein serine/threonine kinase activity | 6 | 0,000163412876 |
| Hypochthonius rufulus | GO:0051725 | biological_process | protein de-ADP-ribosylation | 6 | 0,000175276472 |
| Hypochthonius rufulus | GO:0015939 | biological_process | pantothenate metabolic process | 6 | 0,000175276472 |
| Hypochthonius rufulus | GO:0006487 | biological_process | protein N-linked glycosylation | 6 | 0,000225975808 |
| Hypochthonius rufulus | GO:0008354 | biological_process | germ cell migration | 6 | 0,000345637271 |
| Hypochthonius rufulus | GO:0048010 | biological_process | vascular endothelial growth factor receptor signaling pathway | 6 | 0,007389547292 |
| Hypochthonius rufulus | GO:0016055 | biological_process | Wnt signaling pathway | 5 | 0 |
| Hypochthonius rufulus | GO:0004252 | molecular_function | serine-type endopeptidase activity | 5 | 0 |
| Hypochthonius rufulus | GO:0031098 | biological_process | stress-activated protein kinase signaling cascade | 5 | 0 |
| Hypochthonius rufulus | GO:0016705 | molecular_function | oxidoreductase activity, acting on paired donors, with incorporation or reduction of molecular oxygen | 5 | 0,000000074715 |
| Hypochthonius rufulus | GO:0007399 | biological_process | nervous system development | 5 | 0,000006955119 |
| Hypochthonius rufulus | GO:0015893 | biological_process | drug transport | 5 | 0,000012455091 |
| Hypochthonius rufulus | GO:0031638 | biological_process | zymogen activation | 5 | 0,000076400396 |
| Hypochthonius rufulus | GO:0006641 | biological_process | triglyceride metabolic process | 5 | 0,000525139943 |
| Hypochthonius rufulus | GO:0036323 | biological_process | vascular endothelial growth factor receptor-1 signaling pathway | 5 | 0,004578095432 |
| Hypochthonius rufulus | GO:0048364 | biological_process | root development | 5 | 0,01627210269 |
| Hypochthonius rufulus | GO:0006366 | biological_process | transcription from RNA polymerase II promoter | 4 | 0 |
| Hypochthonius rufulus | GO:0009405 | biological_process | pathogenesis | 4 | 0,000085247376 |
| Hypochthonius rufulus | GO:0034405 | biological_process | response to fluid shear stress | 4 | 0,00144181761 |
| Hypochthonius rufulus | GO:0051958 | biological_process | methotrexate transport | 4 | 0,00144181761 |

| | | | | | |
|---|---|---|---|---|---|
| Hypochthonius rufulus | GO:0007584 | biological_process | response to nutrient | 4 | 0,001696867555 |
| Hypochthonius rufulus | GO:2000504 | biological_process | positive regulation of blood vessel remodeling | 4 | 0,003014112452 |
| Hypochthonius rufulus | GO:0006401 | biological_process | RNA catabolic process | 4 | 0,004055575491 |
| Hypochthonius rufulus | GO:0008654 | biological_process | phospholipid biosynthetic process | 4 | 0,006409088443 |
| Hypochthonius rufulus | GO:0001932 | biological_process | regulation of protein phosphorylation | 4 | 0,00870897087 |
| Hypochthonius rufulus | GO:0000070 | biological_process | mitotic sister chromatid segregation | 4 | 0,01300313641 |
| Hypochthonius rufulus | GO:0006364 | biological_process | rRNA processing | 3 | 0 |
| Hypochthonius rufulus | GO:0007605 | biological_process | sensory perception of sound | 3 | 0 |
| Hypochthonius rufulus | GO:0007169 | biological_process | transmembrane receptor protein tyrosine kinase signaling pathway | 3 | 0 |
| Hypochthonius rufulus | GO:0055114 | biological_process | oxidation-reduction process | 3 | 0,000000017479 |
| Hypochthonius rufulus | GO:0001570 | biological_process | vasculogenesis | 3 | 0,000508494965 |
| Hypochthonius rufulus | GO:0003676 | molecular_function | nucleic acid binding | 3 | 0,000607621856 |
| Hypochthonius rufulus | GO:0015739 | biological_process | sialic acid transport | 3 | 0,001589312543 |
| Hypochthonius rufulus | GO:0001894 | biological_process | tissue homeostasis | 3 | 0,001731670881 |
| Hypochthonius rufulus | GO:0046680 | biological_process | response to DDT | 3 | 0,004115151543 |
| Hypochthonius rufulus | GO:0042770 | biological_process | signal transduction in response to DNA damage | 3 | 0,006038348585 |
| Hypochthonius rufulus | GO:0006904 | biological_process | vesicle docking involved in exocytosis | 3 | 0,007678442178 |
| Hypochthonius rufulus | GO:0008360 | biological_process | regulation of cell shape | 3 | 0,01051833203 |
| Hypochthonius rufulus | GO:0051225 | biological_process | spindle assembly | 3 | 0,01931423317 |
| Hypochthonius rufulus | GO:0032225 | biological_process | regulation of synaptic transmission, dopaminergic | 3 | 0,01931423317 |
| Hypochthonius rufulus | GO:0001558 | biological_process | regulation of cell growth | 3 | 0,02079198336 |
| Hypochthonius rufulus | GO:0006412 | biological_process | translation | 2 | 0 |
| Hypochthonius rufulus | GO:0015031 | biological_process | protein transport | 2 | 0 |
| Hypochthonius rufulus | GO:0006357 | biological_process | regulation of transcription from RNA polymerase II promoter | 2 | 0 |
| Hypochthonius rufulus | GO:0001669 | cellular_component | acrosomal vesicle | 2 | 0 |
| Hypochthonius rufulus | GO:0000122 | biological_process | negative regulation of transcription from RNA polymerase II promoter | 2 | 0,000000000288 |
| Hypochthonius rufulus | GO:0006413 | biological_process | translational initiation | 2 | 0,000000002574 |
| Hypochthonius rufulus | GO:0005886 | cellular_component | plasma membrane | 2 | 0,000000006828 |
| Hypochthonius rufulus | GO:0000209 | biological_process | protein polyubiquitination | 2 | 0,000000030745 |
| Hypochthonius rufulus | GO:0035220 | biological_process | wing disc development | 2 | 0,000000898740 |
| Hypochthonius rufulus | GO:0008542 | biological_process | visual learning | 2 | 0,000003104566 |
| Hypochthonius rufulus | GO:1902600 | biological_process | hydrogen ion transmembrane transport | 2 | 0,000017468370 |
| Hypochthonius rufulus | GO:0006915 | biological_process | apoptotic process | 2 | 0,000021353127 |
| Hypochthonius rufulus | GO:0051301 | biological_process | cell division | 2 | 0,000026087470 |
| Hypochthonius rufulus | GO:0006383 | biological_process | transcription from RNA polymerase III promoter | 2 | 0,000077670333 |
| Hypochthonius rufulus | GO:0032465 | biological_process | regulation of cytokinesis | 2 | 0,000302656794 |
| Hypochthonius rufulus | GO:0001659 | biological_process | temperature homeostasis | 2 | 0,000403172798 |

| | | | | | |
|---|---|---|---|---|---|
| Hypochthonius rufulus | GO:0001505 | biological_process | regulation of neurotransmitter levels | 2 | 0,000487619415 |
| Hypochthonius rufulus | GO:0010842 | biological_process | retina layer formation | 2 | 0,000647562454 |
| Hypochthonius rufulus | GO:0019285 | biological_process | glycine betaine biosynthetic process from choline | 2 | 0,001800271039 |
| Hypochthonius rufulus | GO:0008289 | molecular_function | lipid binding | 2 | 0,009710144402 |
| Hypochthonius rufulus | GO:0071622 | biological_process | regulation of granulocyte chemotaxis | 2 | 0,01099180367 |
| Hypochthonius rufulus | GO:0042381 | biological_process | hemolymph coagulation | 2 | 0,01480317708 |
| Hypochthonius rufulus | GO:0016925 | biological_process | protein sumoylation | 2 | 0,01608438098 |
| Hypochthonius rufulus | GO:0042802 | molecular_function | identical protein binding | 2 | 0,01608438098 |

| | | | | |
|---|---|---|---|---|
| Oppiella subpectinata | GO:0032100 | biological_process | positive regulation of appetite | 218 |
| Oppiella subpectinata | GO:0055085 | biological_process | transmembrane transport | 199 |
| Oppiella subpectinata | GO:0006486 | biological_process | protein glycosylation | 95 |
| Oppiella subpectinata | GO:0007283 | biological_process | spermatogenesis | 88 |
| Oppiella subpectinata | GO:0032197 | biological_process | transposition, RNA-mediated | 62 |
| Oppiella subpectinata | GO:0030223 | biological_process | neutrophil differentiation | 59 |
| Oppiella subpectinata | GO:0005975 | biological_process | carbohydrate metabolic process | 56 |
| Oppiella subpectinata | GO:0015074 | biological_process | DNA integration | 52 |
| Oppiella subpectinata | GO:0005524 | molecular_function | ATP binding | 51 |
| Oppiella subpectinata | GO:0051384 | biological_process | response to glucocorticoid | 48 |
| Oppiella subpectinata | GO:0004364 | molecular_function | glutathione transferase activity | 47 |
| Oppiella subpectinata | GO:0006633 | biological_process | fatty acid biosynthetic process | 46 |
| Oppiella subpectinata | GO:0019542 | biological_process | propionate biosynthetic process | 41 |
| Oppiella subpectinata | GO:0003964 | molecular_function | RNA-directed DNA polymerase activity | 40 |
| Oppiella subpectinata | GO:0042178 | biological_process | xenobiotic catabolic process | 38 |
| Oppiella subpectinata | GO:0046872 | molecular_function | metal ion binding | 36 |
| Oppiella subpectinata | GO:0006511 | biological_process | ubiquitin-dependent protein catabolic process | 34 |
| Oppiella subpectinata | GO:0006517 | biological_process | protein deglycosylation | 34 |
| Oppiella subpectinata | GO:0008270 | molecular_function | zinc ion binding | 34 |
| Oppiella subpectinata | GO:0055129 | biological_process | L-proline biosynthetic process | 31 |
| Oppiella subpectinata | GO:0045214 | biological_process | sarcomere organization | 31 |
| Oppiella subpectinata | GO:0000723 | biological_process | telomere maintenance | 31 |
| Oppiella subpectinata | GO:0006013 | biological_process | mannose metabolic process | 30 |
| Oppiella subpectinata | GO:0006706 | biological_process | steroid catabolic process | 30 |
| Oppiella subpectinata | GO:0009888 | biological_process | tissue development | 29 |
| Oppiella subpectinata | GO:0019284 | biological_process | L-methionine biosynthetic process from S-adenosylmethionine | 29 |
| Oppiella subpectinata | GO:0006915 | biological_process | apoptotic process | 26 |
| Oppiella subpectinata | GO:0010992 | biological_process | ubiquitin homeostasis | 25 |
| Oppiella subpectinata | GO:0031048 | biological_process | chromatin silencing by small RNA | 25 |
| Oppiella subpectinata | GO:0016192 | biological_process | vesicle-mediated transport | 25 |
| Oppiella subpectinata | GO:0004252 | molecular_function | serine-type endopeptidase activity | 25 |
| Oppiella subpectinata | GO:0044245 | biological_process | polysaccharide digestion | 24 |
| Oppiella subpectinata | GO:0006805 | biological_process | xenobiotic metabolic process | 24 |
| Oppiella subpectinata | GO:0009617 | biological_process | response to bacterium | 23 |
| Oppiella subpectinata | GO:0046693 | biological_process | sperm storage | 23 |
| Oppiella subpectinata | GO:0016491 | molecular_function | oxidoreductase activity | 21 |
| Oppiella subpectinata | GO:0006811 | biological_process | ion transport | 21 |

| | | | | |
|---|---|---|---|---|
| Oppiella subpectinata | GO:0090502 | biological_process | RNA phosphodiester bond hydrolysis, endonucleolytic | 20 |
| Oppiella subpectinata | GO:0006069 | biological_process | ethanol oxidation | 19 |
| Oppiella subpectinata | GO:0033609 | biological_process | oxalate metabolic process | 18 |
| Oppiella subpectinata | GO:0006310 | biological_process | DNA recombination | 18 |
| Oppiella subpectinata | GO:0010842 | biological_process | retina layer formation | 18 |
| Oppiella subpectinata | GO:0023052 | biological_process | signaling | 17 |
| Oppiella subpectinata | GO:0043252 | biological_process | sodium-independent organic anion transport | 17 |
| Oppiella subpectinata | GO:0006728 | biological_process | pteridine biosynthetic process | 16 |
| Oppiella subpectinata | GO:0006656 | biological_process | phosphatidylcholine biosynthetic process | 16 |
| Oppiella subpectinata | GO:0016042 | biological_process | lipid catabolic process | 16 |
| Oppiella subpectinata | GO:0031098 | biological_process | stress-activated protein kinase signaling cascade | 16 |
| Oppiella subpectinata | GO:0006470 | biological_process | protein dephosphorylation | 16 |
| Oppiella subpectinata | GO:0006979 | biological_process | response to oxidative stress | 15 |
| Oppiella subpectinata | GO:1902477 | biological_process | regulation of defense response to bacterium, incompatible interaction | 14 |
| Oppiella subpectinata | GO:0007016 | biological_process | cytoskeletal anchoring at plasma membrane | 14 |
| Oppiella subpectinata | GO:0006412 | biological_process | translation | 13 |
| Oppiella subpectinata | GO:0033089 | biological_process | positive regulation of T cell differentiation in thymus | 13 |
| Oppiella subpectinata | GO:0006152 | biological_process | purine nucleoside catabolic process | 13 |
| Oppiella subpectinata | GO:0007601 | biological_process | visual perception | 12 |
| Oppiella subpectinata | GO:0006468 | biological_process | protein phosphorylation | 12 |
| Oppiella subpectinata | GO:0005983 | biological_process | starch catabolic process | 12 |
| Oppiella subpectinata | GO:0007605 | biological_process | sensory perception of sound | 12 |
| Oppiella subpectinata | GO:0000272 | biological_process | polysaccharide catabolic process | 11 |
| Oppiella subpectinata | GO:0016567 | biological_process | protein ubiquitination | 10 |
| Oppiella subpectinata | GO:0016790 | molecular_function | thiolester hydrolase activity | 10 |
| Oppiella subpectinata | GO:0055069 | biological_process | zinc ion homeostasis | 10 |
| Oppiella subpectinata | GO:0010569 | biological_process | regulation of double-strand break repair via homologous recombination | 10 |
| Oppiella subpectinata | GO:0008380 | biological_process | RNA splicing | 9 |
| Oppiella subpectinata | GO:0003677 | molecular_function | DNA binding | 9 |
| Oppiella subpectinata | GO:0097066 | biological_process | response to thyroid hormone | 9 |
| Oppiella subpectinata | GO:0009636 | biological_process | response to toxic substance | 9 |
| Oppiella subpectinata | GO:0034765 | biological_process | regulation of ion transmembrane transport | 9 |
| Oppiella subpectinata | GO:0006099 | biological_process | tricarboxylic acid cycle | 8 |
| Oppiella subpectinata | GO:0005506 | molecular_function | iron ion binding | 8 |
| Oppiella subpectinata | GO:0004559 | molecular_function | alpha-mannosidase activity | 7 |
| Oppiella subpectinata | GO:0007586 | biological_process | digestion | 7 |
| Oppiella subpectinata | GO:0007179 | biological_process | transforming growth factor beta receptor signaling pathway | 7 |

| | | | | |
|---|---|---|---|---|
| Oppiella subpectinata | GO:0008202 | biological_process | steroid metabolic process | 7 |
| Oppiella subpectinata | GO:0007165 | biological_process | signal transduction | 6 |
| Oppiella subpectinata | GO:0016614 | molecular_function | oxidoreductase activity, acting on CH-OH group of donors | 6 |
| Oppiella subpectinata | GO:0015939 | biological_process | pantothenate metabolic process | 6 |
| Oppiella subpectinata | GO:0006694 | biological_process | steroid biosynthetic process | 6 |
| Oppiella subpectinata | GO:0004867 | molecular_function | serine-type endopeptidase inhibitor activity | 6 |
| Oppiella subpectinata | GO:0033384 | biological_process | geranyl diphosphate biosynthetic process | 6 |
| Oppiella subpectinata | GO:0050951 | biological_process | sensory perception of temperature stimulus | 5 |
| Oppiella subpectinata | GO:0030449 | biological_process | regulation of complement activation | 5 |
| Oppiella subpectinata | GO:0071928 | biological_process | tyramine signaling pathway | 5 |
| Oppiella subpectinata | GO:0016032 | biological_process | viral process | 4 |
| Oppiella subpectinata | GO:0005576 | cellular_component | extracellular region | 4 |
| Oppiella subpectinata | GO:0055114 | biological_process | oxidation-reduction process | 4 |
| Oppiella subpectinata | GO:0048670 | biological_process | regulation of collateral sprouting | 4 |
| Oppiella subpectinata | GO:0008240 | molecular_function | tripeptidyl-peptidase activity | 4 |
| Oppiella subpectinata | GO:0007155 | biological_process | cell adhesion | 4 |
| Oppiella subpectinata | GO:0042572 | biological_process | retinol metabolic process | 4 |
| Oppiella subpectinata | GO:0031638 | biological_process | zymogen activation | 4 |
| Oppiella subpectinata | GO:0006749 | biological_process | glutathione metabolic process | 4 |
| Oppiella subpectinata | GO:0006629 | biological_process | lipid metabolic process | 4 |
| Oppiella subpectinata | GO:0007608 | biological_process | sensory perception of smell | 4 |
| Oppiella subpectinata | GO:0030497 | biological_process | fatty acid elongation | 4 |
| Oppiella subpectinata | GO:0006357 | biological_process | regulation of transcription from RNA polymerase II promoter | 3 |
| Oppiella subpectinata | GO:0016705 | molecular_function | oxidoreductase activity, acting on paired donors, with incorporation or reduction of molecular oxygen | 3 |
| Oppiella subpectinata | GO:0045892 | biological_process | negative regulation of transcription, DNA-templated | 3 |
| Oppiella subpectinata | GO:0008542 | biological_process | visual learning | 3 |
| Oppiella subpectinata | GO:0006641 | biological_process | triglyceride metabolic process | 3 |
| Oppiella subpectinata | GO:0010758 | biological_process | regulation of macrophage chemotaxis | 3 |
| Oppiella subpectinata | GO:0010468 | biological_process | regulation of gene expression | 3 |
| Oppiella subpectinata | GO:0004930 | molecular_function | G-protein coupled receptor activity | 3 |
| Oppiella subpectinata | GO:0072431 | biological_process | signal transduction involved in mitotic G1 DNA damage checkpoint | 3 |
| Oppiella subpectinata | GO:0009107 | biological_process | lipoate biosynthetic process | 3 |
| Oppiella subpectinata | GO:0010506 | biological_process | regulation of autophagy | 3 |
| Oppiella subpectinata | GO:0050661 | molecular_function | NADP binding | 3 |
| Oppiella subpectinata | GO:0006355 | biological_process | regulation of transcription, DNA-templated | 2 |
| Oppiella subpectinata | GO:0016055 | biological_process | Wnt signaling pathway | 2 |
| Oppiella subpectinata | GO:0007275 | biological_process | multicellular organismal development | 2 |

| | | | | |
|---|---|---|---|---|
| Oppiella subpectinata | GO:0006814 | biological_process | sodium ion transport | 2 |
| Oppiella subpectinata | GO:0006351 | biological_process | transcription, DNA-templated | 2 |
| Oppiella subpectinata | GO:0030433 | biological_process | ER-associated ubiquitin-dependent protein catabolic process | 2 |
| Oppiella subpectinata | GO:0042060 | biological_process | wound healing | 2 |
| Oppiella subpectinata | GO:0007286 | biological_process | spermatid development | 2 |
| Oppiella subpectinata | GO:0016787 | molecular_function | hydrolase activity | 2 |
| Oppiella subpectinata | GO:0035249 | biological_process | synaptic transmission, glutamatergic | 2 |
| Oppiella subpectinata | GO:0007399 | biological_process | nervous system development | 2 |
| Oppiella subpectinata | GO:0010224 | biological_process | response to UV-B | 2 |
| Oppiella subpectinata | GO:0006417 | biological_process | regulation of translation | 2 |
| Oppiella subpectinata | GO:0007584 | biological_process | response to nutrient | 2 |
| Oppiella subpectinata | GO:0031146 | biological_process | SCF-dependent proteasomal ubiquitin-dependent protein catabolic process | 2 |
| Oppiella subpectinata | GO:0001659 | biological_process | temperature homeostasis | 2 |
| Oppiella subpectinata | GO:0030163 | biological_process | protein catabolic process | 2 |
| Oppiella subpectinata | GO:0042981 | biological_process | regulation of apoptotic process | 2 |
| Oppiella subpectinata | GO:0005154 | molecular_function | epidermal growth factor receptor binding | 2 |
| Oppiella subpectinata | GO:0010951 | biological_process | negative regulation of endopeptidase activity | 2 |
| Oppiella subpectinata | GO:0030246 | molecular_function | carbohydrate binding | 2 |
| Oppiella subpectinata | GO:0032880 | biological_process | regulation of protein localization | 2 |
| Oppiella subpectinata | GO:0010977 | biological_process | negative regulation of neuron projection development | 2 |

| |
|---|
| 0 |
| 0 |
| 0 |
| 0,000664283285 |
| 0 |
| 0 |
| 0,0173853986 |
| 0 |
| 0 |
| 0,02411287018 |
| 0 |
| 0,01008482811 |
| 0 |
| 0,000000000448 |
| 0 |
| 0,000000017915 |
| 0 |
| 0,000000061459 |
| 0,000086784557 |
| 0 |
| 0,002150960647 |
| 0,002285458678 |
| 0,000000000115[illegible] |
| 0,00525036936 |
| 0,000000403741 |
| 0,000122626450 |
| 0,000040643623[illegible] |
| 0 |
| 0 |
| 0 |
| 0,000018237478 |
| 0 |
| 0,003716380801 |
| 0,000001513381 |
| 0,009358219422 |
| 0,000160227650 |
| 0,01742552594 |

| |
|---|
| 0,000022133192 |
| 0,000052478303 |
| 0 |
| 0,000089199191 |
| 0,001338001728 |
| 0,000017608638 |
| 0,005238707425 |
| 0 |
| 0,000000002103 |
| 0,000029122190 |
| 0,000840567082 |
| 0,006150483373 |
| 0,000000001649 |
| 0 |
| 0,000000584332 |
| 0 |
| 0,000000000134 |
| 0,000000000353 |
| 0 |
| 0 |
| 0,000000054041[illegible] |
| 0,001508547447 |
| 0,000004752087 |
| 0 |
| 0 |
| 0,000002113461[illegible] |
| 0,000030283591 |
| 0 |
| 0,000000348802 |
| 0,000001176479 |
| 0,000004442736 |
| 0,004212183134 |
| 0,000007355677 |
| 0,001415165014 |
| 0,000002568944 |
| 0,001031454445 |
| 0,002555271125 |

| |
|---|
| 0,006651960993 |
| 0 |
| 0,000001599744 |
| 0,000049938971 |
| 0,001160441497 |
| 0,00545335481 |
| 0,02186222192 |
| 0,000022319873 |
| 0,002613351271 |
| 0,02065667721 |
| 0 |
| 0,000000165755 |
| 0,000010720852 |
| 0,000048860196 |
| 0,000048860196 |
| 0,000102750652 |
| 0,000193655733 |
| 0,000442803128 |
| 0,000542634594 |
| 0,000811298133 |
| 0,001465845854 |
| 0,01643905256 |
| 0 |
| 0,000000309489 |
| 0,000000335672 |
| 0,000398211264 |
| 0,000568712850 |
| 0,000586938887 |
| 0,005355873427 |
| 0,007855671994 |
| 0,009043791152 |
| 0,01450809493 |
| 0,01742064837 |
| 0,02471457027 |
| 0 |
| 0 |
| 0 |

| |
|---|
| 0 |
| 0 |
| 0 |
| 0 |
| 0,000000006788 |
| 0,000000089210 |
| 0,000000148363 |
| 0,000002147623 |
| 0,000026551496 |
| 0,000086879863 |
| 0,001075991106 |
| 0,001554550683 |
| 0,003203197728 |
| 0,003964429852 |
| 0,005252780101 |
| 0,007030346607 |
| 0,007430042889 |
| 0,009763523956 |
| 0,01117500945 |
| 0,01933022316 |

| | | | | | |
|---|---|---|---|---|---|
| Oppiella nova | GO:0055085 | biological_process | transmembrane transport | 315 | 0 |
| Oppiella nova | GO:0042908 | biological_process | xenobiotic transport | 96 | 0 |
| Oppiella nova | GO:0003677 | molecular_function | DNA binding | 74 | 0 |
| Oppiella nova | GO:0006517 | biological_process | protein deglycosylation | 63 | 0 |
| Oppiella nova | GO:0005975 | biological_process | carbohydrate metabolic process | 59 | 0,000002294505 |
| Oppiella nova | GO:0051384 | biological_process | response to glucocorticoid | 48 | 0,000093560940 |
| Oppiella nova | GO:0009313 | biological_process | oligosaccharide catabolic process | 47 | 0 |
| Oppiella nova | GO:0003964 | molecular_function | RNA-directed DNA polymerase activity | 41 | 0 |
| Oppiella nova | GO:0006313 | biological_process | transposition, DNA-mediated | 40 | 0,000155361265 |
| Oppiella nova | GO:0006680 | biological_process | glucosylceramide catabolic process | 39 | 0 |
| Oppiella nova | GO:0007601 | biological_process | visual perception | 38 | 0,005866232295 |
| Oppiella nova | GO:0009311 | biological_process | oligosaccharide metabolic process | 36 | 0 |
| Oppiella nova | GO:0052697 | biological_process | xenobiotic glucuronidation | 32 | 0 |
| Oppiella nova | GO:0007283 | biological_process | spermatogenesis | 31 | 0 |
| Oppiella nova | GO:0000270 | biological_process | peptidoglycan metabolic process | 30 | 0 |
| Oppiella nova | GO:0007286 | biological_process | spermatid development | 30 | 0,000457483959 |
| Oppiella nova | GO:0071357 | biological_process | cellular response to type I interferon | 25 | 0 |
| Oppiella nova | GO:0001516 | biological_process | prostaglandin biosynthetic process | 24 | 0 |
| Oppiella nova | GO:1901998 | biological_process | toxin transport | 23 | 0,000006306416 |
| Oppiella nova | GO:0010383 | biological_process | cell wall polysaccharide metabolic process | 22 | 0 |
| Oppiella nova | GO:0006706 | biological_process | steroid catabolic process | 21 | 0,02025918526 |
| Oppiella nova | GO:0010951 | biological_process | negative regulation of endopeptidase activity | 20 | 0,000000732316 |
| Oppiella nova | GO:0008210 | biological_process | estrogen metabolic process | 19 | 0 |
| Oppiella nova | GO:0007165 | biological_process | signal transduction | 17 | 0 |
| Oppiella nova | GO:0008146 | molecular_function | sulfotransferase activity | 17 | 0 |
| Oppiella nova | GO:0051923 | biological_process | sulfation | 17 | 0,000001280954 |
| Oppiella nova | GO:0004252 | molecular_function | serine-type endopeptidase activity | 17 | 0,000092152242 |
| Oppiella nova | GO:0045332 | biological_process | phospholipid translocation | 16 | 0,000000051702 |
| Oppiella nova | GO:0002376 | biological_process | immune system process | 16 | 0,000030263708 |
| Oppiella nova | GO:0010842 | biological_process | retina layer formation | 16 | 0,000325500578 |
| Oppiella nova | GO:0071555 | biological_process | cell wall organization | 16 | 0,000864634950 |
| Oppiella nova | GO:0032434 | biological_process | regulation of proteasomal ubiquitin-dependent protein catabolic process | 15 | 0,000000002448 |
| Oppiella nova | GO:0009617 | biological_process | response to bacterium | 14 | 0,000002834548 |
| Oppiella nova | GO:0007021 | biological_process | tubulin complex assembly | 14 | 0,000155067092 |
| Oppiella nova | GO:0016266 | biological_process | O-glycan processing | 14 | 0,002100817823 |
| Oppiella nova | GO:0045944 | biological_process | positive regulation of transcription from RNA polymerase II promoter | 12 | 0,000000668949 |
| Oppiella nova | GO:0004869 | molecular_function | cysteine-type endopeptidase inhibitor activity | 12 | 0,000001604099 |

| | | | | | |
|---|---|---|---|---|---|
| Oppiella nova | GO:0004518 | molecular_function | nuclease activity | 12 | 0,000959144421 |
| Oppiella nova | GO:0006508 | biological_process | proteolysis | 11 | 0 |
| Oppiella nova | GO:0014003 | biological_process | oligodendrocyte development | 11 | 0,000006332728 |
| Oppiella nova | GO:0046355 | biological_process | mannan catabolic process | 11 | 0,000544105624 |
| Oppiella nova | GO:0006013 | biological_process | mannose metabolic process | 11 | 0,02108746546 |
| Oppiella nova | GO:0006355 | biological_process | regulation of transcription, DNA-templated | 10 | 0 |
| Oppiella nova | GO:0019882 | biological_process | antigen processing and presentation | 9 | 0,000232665760 |
| Oppiella nova | GO:0042135 | biological_process | neurotransmitter catabolic process | 8 | 0,01384184796 |
| Oppiella nova | GO:1901639 | biological_process | XDP catabolic process | 7 | 0,000047517901 |
| Oppiella nova | GO:0044179 | biological_process | hemolysis in other organism | 7 | 0,007177857707 |
| Oppiella nova | GO:0006511 | biological_process | ubiquitin-dependent protein catabolic process | 6 | 0 |
| Oppiella nova | GO:0006468 | biological_process | protein phosphorylation | 6 | 0,000000000273 |
| Oppiella nova | GO:0006470 | biological_process | protein dephosphorylation | 6 | 0,000145381603 |
| Oppiella nova | GO:0000103 | biological_process | sulfate assimilation | 6 | 0,003982995238 |
| Oppiella nova | GO:0007155 | biological_process | cell adhesion | 5 | 0,009901683158 |
| Oppiella nova | GO:0016491 | molecular_function | oxidoreductase activity | 4 | 0,000000000918 |
| Oppiella nova | GO:0001932 | biological_process | regulation of protein phosphorylation | 4 | 0,001259785645 |
| Oppiella nova | GO:0006351 | biological_process | transcription, DNA-templated | 3 | 0,000000048577 |
| Oppiella nova | GO:0006325 | biological_process | chromatin organization | 3 | 0,00011187382 |
| Oppiella nova | GO:0060675 | biological_process | ureteric bud morphogenesis | 3 | 0,000848563557 |
| Oppiella nova | GO:0008202 | biological_process | steroid metabolic process | 3 | 0,002425032343 |
| Oppiella nova | GO:0031638 | biological_process | zymogen activation | 3 | 0,003245189087 |
| Oppiella nova | GO:0032197 | biological_process | transposition, RNA-mediated | 3 | 0,02091294315 |
| Oppiella nova | GO:0016055 | biological_process | Wnt signaling pathway | 2 | 0 |
| Oppiella nova | GO:0016192 | biological_process | vesicle-mediated transport | 2 | 0 |
| Oppiella nova | GO:0046872 | molecular_function | metal ion binding | 2 | 0 |
| Oppiella nova | GO:0016567 | biological_process | protein ubiquitination | 2 | 0 |
| Oppiella nova | GO:0008270 | molecular_function | zinc ion binding | 2 | 0 |
| Oppiella nova | GO:0006979 | biological_process | response to oxidative stress | 2 | 0 |
| Oppiella nova | GO:0007605 | biological_process | sensory perception of sound | 2 | 0,00000118426 |
| Oppiella nova | GO:0042626 | molecular_function | ATPase activity, coupled to transmembrane movement of substances | 2 | 0,000026363944 |
| Oppiella nova | GO:0005506 | molecular_function | iron ion binding | 2 | 0,00004711998 |
| Oppiella nova | GO:0006694 | biological_process | steroid biosynthetic process | 2 | 0,000259398409 |
| Oppiella nova | GO:0004674 | molecular_function | protein serine/threonine kinase activity | 2 | 0,000594710312 |
| Oppiella nova | GO:0006644 | biological_process | phospholipid metabolic process | 2 | 0,001412739411 |
| Oppiella nova | GO:0006417 | biological_process | regulation of translation | 2 | 0,00227529802 |
| Oppiella nova | GO:0006641 | biological_process | triglyceride metabolic process | 2 | 0,00280488779 |

| | | | | | |
|---|---|---|---|---|---|
| Oppiella nova | GO:0030497 | biological_process | fatty acid elongation | 2 | 0,01614269236 |
| Oppiella nova | GO:0046601 | biological_process | positive regulation of centriole replication | 2 | 0,01970981718 |
| Oppiella nova | GO:0061077 | biological_process | chaperone-mediated protein folding | 2 | 0,01970981718 |
| Oppiella nova | GO:0001501 | biological_process | skeletal system development | 2 | 0,02140907902 |

| | | | | |
|---|---|---|---|---|
| Hydrozetes thienemanni | GO:0032100 | biological_process | positive regulation of appetite | 218 |
| Hydrozetes thienemanni | GO:0042908 | biological_process | xenobiotic transport | 96 |
| Hydrozetes thienemanni | GO:0007283 | biological_process | spermatogenesis | 93 |
| Hydrozetes thienemanni | GO:0006633 | biological_process | fatty acid biosynthetic process | 89 |
| Hydrozetes thienemanni | GO:0016020 | cellular_component | membrane | 80 |
| Hydrozetes thienemanni | GO:0015074 | biological_process | DNA integration | 77 |
| Hydrozetes thienemanni | GO:0055129 | biological_process | L-proline biosynthetic process | 75 |
| Hydrozetes thienemanni | GO:0032197 | biological_process | transposition, RNA-mediated | 71 |
| Hydrozetes thienemanni | GO:0016567 | biological_process | protein ubiquitination | 68 |
| Hydrozetes thienemanni | GO:0006508 | biological_process | proteolysis | 65 |
| Hydrozetes thienemanni | GO:0003677 | molecular_function | DNA binding | 65 |
| Hydrozetes thienemanni | GO:0008218 | biological_process | bioluminescence | 63 |
| Hydrozetes thienemanni | GO:0030245 | biological_process | cellulose catabolic process | 62 |
| Hydrozetes thienemanni | GO:0030223 | biological_process | neutrophil differentiation | 58 |
| Hydrozetes thienemanni | GO:0004252 | molecular_function | serine-type endopeptidase activity | 57 |
| Hydrozetes thienemanni | GO:0006260 | biological_process | DNA replication | 52 |
| Hydrozetes thienemanni | GO:0008194 | molecular_function | UDP-glycosyltransferase activity | 50 |
| Hydrozetes thienemanni | GO:0006511 | biological_process | ubiquitin-dependent protein catabolic process | 49 |
| Hydrozetes thienemanni | GO:0007601 | biological_process | visual perception | 46 |
| Hydrozetes thienemanni | GO:0046982 | molecular_function | protein heterodimerization activity | 42 |
| Hydrozetes thienemanni | GO:0003964 | molecular_function | RNA-directed DNA polymerase activity | 42 |
| Hydrozetes thienemanni | GO:0010224 | biological_process | response to UV-B | 40 |
| Hydrozetes thienemanni | GO:0009617 | biological_process | response to bacterium | 39 |
| Hydrozetes thienemanni | GO:0005506 | molecular_function | iron ion binding | 39 |
| Hydrozetes thienemanni | GO:0004867 | molecular_function | serine-type endopeptidase inhibitor activity | 38 |
| Hydrozetes thienemanni | GO:0016705 | molecular_function | oxidoreductase activity, acting on paired donors, with incorporation or reduction of molecular oxygen | 38 |
| Hydrozetes thienemanni | GO:0007165 | biological_process | signal transduction | 37 |
| Hydrozetes thienemanni | GO:0016192 | biological_process | vesicle-mediated transport | 36 |
| Hydrozetes thienemanni | GO:0052697 | biological_process | xenobiotic glucuronidation | 36 |
| Hydrozetes thienemanni | GO:0006364 | biological_process | rRNA processing | 34 |
| Hydrozetes thienemanni | GO:0006517 | biological_process | protein deglycosylation | 34 |
| Hydrozetes thienemanni | GO:0006979 | biological_process | response to oxidative stress | 31 |
| Hydrozetes thienemanni | GO:0010951 | biological_process | negative regulation of endopeptidase activity | 30 |
| Hydrozetes thienemanni | GO:0006013 | biological_process | mannose metabolic process | 30 |
| Hydrozetes thienemanni | GO:0046548 | biological_process | retinal rod cell development | 28 |
| Hydrozetes thienemanni | GO:0009888 | biological_process | tissue development | 28 |
| Hydrozetes thienemanni | GO:0009636 | biological_process | response to toxic substance | 28 |

| | | | | |
|---|---|---|---|---|
| Hydrozetes thienemanni | GO:1902358 | biological_process | sulfate transmembrane transport | 27 |
| Hydrozetes thienemanni | GO:0004518 | molecular_function | nuclease activity | 26 |
| Hydrozetes thienemanni | GO:0005975 | biological_process | carbohydrate metabolic process | 25 |
| Hydrozetes thienemanni | GO:0014832 | biological_process | urinary bladder smooth muscle contraction | 25 |
| Hydrozetes thienemanni | GO:0051260 | biological_process | protein homooligomerization | 25 |
| Hydrozetes thienemanni | GO:0006355 | biological_process | regulation of transcription, DNA-templated | 24 |
| Hydrozetes thienemanni | GO:0010811 | biological_process | positive regulation of cell-substrate adhesion | 24 |
| Hydrozetes thienemanni | GO:0071357 | biological_process | cellular response to type I interferon | 24 |
| Hydrozetes thienemanni | GO:0016139 | biological_process | glycoside catabolic process | 24 |
| Hydrozetes thienemanni | GO:0045616 | biological_process | regulation of keratinocyte differentiation | 24 |
| Hydrozetes thienemanni | GO:0001516 | biological_process | prostaglandin biosynthetic process | 24 |
| Hydrozetes thienemanni | GO:0006470 | biological_process | protein dephosphorylation | 24 |
| Hydrozetes thienemanni | GO:0045493 | biological_process | xylan catabolic process | 23 |
| Hydrozetes thienemanni | GO:0008270 | molecular_function | zinc ion binding | 22 |
| Hydrozetes thienemanni | GO:0006310 | biological_process | DNA recombination | 22 |
| Hydrozetes thienemanni | GO:0043171 | biological_process | peptide catabolic process | 20 |
| Hydrozetes thienemanni | GO:0016042 | biological_process | lipid catabolic process | 20 |
| Hydrozetes thienemanni | GO:0043097 | biological_process | pyrimidine nucleoside salvage | 20 |
| Hydrozetes thienemanni | GO:0046872 | molecular_function | metal ion binding | 19 |
| Hydrozetes thienemanni | GO:0006062 | biological_process | sorbitol catabolic process | 19 |
| Hydrozetes thienemanni | GO:0007605 | biological_process | sensory perception of sound | 18 |
| Hydrozetes thienemanni | GO:0006805 | biological_process | xenobiotic metabolic process | 17 |
| Hydrozetes thienemanni | GO:0070189 | biological_process | kynurenine metabolic process | 17 |
| Hydrozetes thienemanni | GO:0016021 | cellular_component | integral component of membrane | 16 |
| Hydrozetes thienemanni | GO:0006486 | biological_process | protein glycosylation | 16 |
| Hydrozetes thienemanni | GO:0097310 | biological_process | cap2 mRNA methylation | 16 |
| Hydrozetes thienemanni | GO:0031030 | biological_process | negative regulation of septation initiation signaling | 16 |
| Hydrozetes thienemanni | GO:0034976 | biological_process | response to endoplasmic reticulum stress | 16 |
| Hydrozetes thienemanni | GO:0004722 | molecular_function | protein serine/threonine phosphatase activity | 16 |
| Hydrozetes thienemanni | GO:0006605 | biological_process | protein targeting | 16 |
| Hydrozetes thienemanni | GO:0006457 | biological_process | protein folding | 16 |
| Hydrozetes thienemanni | GO:0007129 | biological_process | synapsis | 16 |
| Hydrozetes thienemanni | GO:0000209 | biological_process | protein polyubiquitination | 16 |
| Hydrozetes thienemanni | GO:0006412 | biological_process | translation | 15 |
| Hydrozetes thienemanni | GO:0006468 | biological_process | protein phosphorylation | 15 |
| Hydrozetes thienemanni | GO:0009168 | biological_process | purine ribonucleoside monophosphate biosynthetic process | 15 |
| Hydrozetes thienemanni | GO:0042761 | biological_process | very long-chain fatty acid biosynthetic process | 15 |

| | | | | |
|---|---|---|---|---|
| Hydrozetes thienemanni | GO:0009263 | biological_process | deoxyribonucleotide biosynthetic process | 15 |
| Hydrozetes thienemanni | GO:0032434 | biological_process | regulation of proteasomal ubiquitin-dependent protein catabolic process | 15 |
| Hydrozetes thienemanni | GO:0006810 | biological_process | transport | 15 |
| Hydrozetes thienemanni | GO:0010139 | biological_process | pyrimidine deoxyribonucleotide salvage | 14 |
| Hydrozetes thienemanni | GO:0016117 | biological_process | carotenoid biosynthetic process | 14 |
| Hydrozetes thienemanni | GO:0001963 | biological_process | synaptic transmission, dopaminergic | 14 |
| Hydrozetes thienemanni | GO:0015020 | molecular_function | glucuronosyltransferase activity | 14 |
| Hydrozetes thienemanni | GO:0033089 | biological_process | positive regulation of T cell differentiation in thymus | 13 |
| Hydrozetes thienemanni | GO:0031098 | biological_process | stress-activated protein kinase signaling cascade | 12 |
| Hydrozetes thienemanni | GO:0035092 | biological_process | sperm chromatin condensation | 12 |
| Hydrozetes thienemanni | GO:0016055 | biological_process | Wnt signaling pathway | 11 |
| Hydrozetes thienemanni | GO:0042060 | biological_process | wound healing | 11 |
| Hydrozetes thienemanni | GO:0016031 | biological_process | tRNA import into mitochondrion | 11 |
| Hydrozetes thienemanni | GO:0042110 | biological_process | T cell activation | 11 |
| Hydrozetes thienemanni | GO:0005509 | molecular_function | calcium ion binding | 11 |
| Hydrozetes thienemanni | GO:0004674 | molecular_function | protein serine/threonine kinase activity | 11 |
| Hydrozetes thienemanni | GO:0008380 | biological_process | RNA splicing | 10 |
| Hydrozetes thienemanni | GO:0051384 | biological_process | response to glucocorticoid | 10 |
| Hydrozetes thienemanni | GO:0016790 | molecular_function | thiolester hydrolase activity | 10 |
| Hydrozetes thienemanni | GO:0072089 | biological_process | stem cell proliferation | 10 |
| Hydrozetes thienemanni | GO:0097066 | biological_process | response to thyroid hormone | 10 |
| Hydrozetes thienemanni | GO:0051603 | biological_process | proteolysis involved in cellular protein catabolic process | 10 |
| Hydrozetes thienemanni | GO:0014883 | biological_process | transition between fast and slow fiber | 10 |
| Hydrozetes thienemanni | GO:0007399 | biological_process | nervous system development | 10 |
| Hydrozetes thienemanni | GO:0010569 | biological_process | regulation of double-strand break repair via homologous recombination | 10 |
| Hydrozetes thienemanni | GO:0042572 | biological_process | retinol metabolic process | 10 |
| Hydrozetes thienemanni | GO:0007275 | biological_process | multicellular organismal development | 9 |
| Hydrozetes thienemanni | GO:0016261 | biological_process | selenocysteine catabolic process | 9 |
| Hydrozetes thienemanni | GO:1902531 | biological_process | regulation of intracellular signal transduction | 9 |
| Hydrozetes thienemanni | GO:0055114 | biological_process | oxidation-reduction process | 9 |
| Hydrozetes thienemanni | GO:0016491 | molecular_function | oxidoreductase activity | 8 |
| Hydrozetes thienemanni | GO:0009116 | biological_process | nucleoside metabolic process | 8 |
| Hydrozetes thienemanni | GO:0007224 | biological_process | smoothened signaling pathway | 8 |
| Hydrozetes thienemanni | GO:0034418 | biological_process | urate biosynthetic process | 8 |
| Hydrozetes thienemanni | GO:0039648 | biological_process | modulation by virus of host protein ubiquitination | 8 |
| Hydrozetes thienemanni | GO:0000272 | biological_process | polysaccharide catabolic process | 7 |
| Hydrozetes thienemanni | GO:0007179 | biological_process | transforming growth factor beta receptor signaling pathway | 7 |

| | | | | |
|---|---|---|---|---|
| Hydrozetes thienemanni | GO:0001944 | biological_process | vasculature development | 7 |
| Hydrozetes thienemanni | GO:0000165 | biological_process | MAPK cascade | 6 |
| Hydrozetes thienemanni | GO:0015939 | biological_process | pantothenate metabolic process | 6 |
| Hydrozetes thienemanni | GO:0005739 | cellular_component | mitochondrion | 6 |
| Hydrozetes thienemanni | GO:0016887 | molecular_function | ATPase activity | 6 |
| Hydrozetes thienemanni | GO:0045087 | biological_process | innate immune response | 6 |
| Hydrozetes thienemanni | GO:0016485 | biological_process | protein processing | 6 |
| Hydrozetes thienemanni | GO:1901639 | biological_process | XDP catabolic process | 6 |
| Hydrozetes thienemanni | GO:0007155 | biological_process | cell adhesion | 5 |
| Hydrozetes thienemanni | GO:0006629 | biological_process | lipid metabolic process | 5 |
| Hydrozetes thienemanni | GO:0006114 | biological_process | glycerol biosynthetic process | 5 |
| Hydrozetes thienemanni | GO:0007223 | biological_process | Wnt signaling pathway, calcium modulating pathway | 5 |
| Hydrozetes thienemanni | GO:0006351 | biological_process | transcription, DNA-templated | 4 |
| Hydrozetes thienemanni | GO:0005576 | cellular_component | extracellular region | 4 |
| Hydrozetes thienemanni | GO:0008202 | biological_process | steroid metabolic process | 4 |
| Hydrozetes thienemanni | GO:0006641 | biological_process | triglyceride metabolic process | 4 |
| Hydrozetes thienemanni | GO:0006267 | biological_process | pre-replicative complex assembly involved in nuclear cell cycle DNA replication | 4 |
| Hydrozetes thienemanni | GO:0009213 | biological_process | pyrimidine deoxyribonucleoside triphosphate catabolic process | 4 |
| Hydrozetes thienemanni | GO:0008360 | biological_process | regulation of cell shape | 4 |
| Hydrozetes thienemanni | GO:0048741 | biological_process | skeletal muscle fiber development | 4 |
| Hydrozetes thienemanni | GO:0015031 | biological_process | protein transport | 3 |
| Hydrozetes thienemanni | GO:0016032 | biological_process | viral process | 3 |
| Hydrozetes thienemanni | GO:0005634 | cellular_component | nucleus | 3 |
| Hydrozetes thienemanni | GO:0042135 | biological_process | neurotransmitter catabolic process | 3 |
| Hydrozetes thienemanni | GO:0006694 | biological_process | steroid biosynthetic process | 3 |
| Hydrozetes thienemanni | GO:0031638 | biological_process | zymogen activation | 3 |
| Hydrozetes thienemanni | GO:0030162 | biological_process | regulation of proteolysis | 3 |
| Hydrozetes thienemanni | GO:0001505 | biological_process | regulation of neurotransmitter levels | 3 |
| Hydrozetes thienemanni | GO:0042770 | biological_process | signal transduction in response to DNA damage | 3 |
| Hydrozetes thienemanni | GO:0060675 | biological_process | ureteric bud morphogenesis | 3 |
| Hydrozetes thienemanni | GO:0042381 | biological_process | hemolymph coagulation | 3 |
| Hydrozetes thienemanni | GO:0042742 | biological_process | defense response to bacterium | 3 |
| Hydrozetes thienemanni | GO:0030433 | biological_process | ER-associated ubiquitin-dependent protein catabolic process | 2 |
| Hydrozetes thienemanni | GO:0006636 | biological_process | unsaturated fatty acid biosynthetic process | 2 |
| Hydrozetes thienemanni | GO:0006890 | biological_process | retrograde vesicle-mediated transport, Golgi to ER | 2 |
| Hydrozetes thienemanni | GO:0016787 | molecular_function | hydrolase activity | 2 |
| Hydrozetes thienemanni | GO:0008168 | molecular_function | methyltransferase activity | 2 |

| | | | | |
|---|---|---|---|---|
| Hydrozetes thienemanni | GO:0007584 | biological_process | response to nutrient | 2 |
| Hydrozetes thienemanni | GO:0030497 | biological_process | fatty acid elongation | 2 |
| Hydrozetes thienemanni | GO:0030163 | biological_process | protein catabolic process | 2 |
| Hydrozetes thienemanni | GO:0010506 | biological_process | regulation of autophagy | 2 |
| Hydrozetes thienemanni | GO:0032496 | biological_process | response to lipopolysaccharide | 2 |
| Hydrozetes thienemanni | GO:0007472 | biological_process | wing disc morphogenesis | 2 |
| Hydrozetes thienemanni | GO:0030148 | biological_process | sphingolipid biosynthetic process | 2 |
| Hydrozetes thienemanni | GO:0004104 | molecular_function | cholinesterase activity | 2 |
| Hydrozetes thienemanni | GO:0032880 | biological_process | regulation of protein localization | 2 |
| Hydrozetes thienemanni | GO:0002376 | biological_process | immune system process | 2 |
| Hydrozetes thienemanni | GO:0042311 | biological_process | vasodilation | 2 |
| Hydrozetes thienemanni | GO:0010718 | biological_process | positive regulation of epithelial to mesenchymal transition | 2 |
| Hydrozetes thienemanni | GO:0004970 | molecular_function | ionotropic glutamate receptor activity | 2 |
| Hydrozetes thienemanni | GO:0007585 | biological_process | respiratory gaseous exchange | 2 |
| Hydrozetes thienemanni | GO:0005154 | molecular_function | epidermal growth factor receptor binding | 2 |
| Hydrozetes thienemanni | GO:0097090 | biological_process | presynaptic membrane organization | 2 |

| |
|---|
| 0 |
| 0 |
| 0 |
| 0,000000001782 |
| 0 |
| 0 |
| 0 |
| 0 |
| 0,000351279994 |
| 0 |
| 0,000238659310 |
| 0 |
| 0 |
| 0 |
| 0,01722311261 |
| 0 |
| 0 |
| 0 |
| 0 |
| 0,000000000491 |
| 0,000000819415[illegible] |
| 0,000007022162 |
| 0,000002185621 |
| 0,003606801366 |
| 0,000029151032 |
| 0,01921450287 |
| 0 |
| 0 |
| 0,000000000314 |
| 0 |
| 0,000106350978[illegible] |
| 0,000000028665 |
| 0,000000211223[illegible] |
| 0,000000343553 |
| 0 |
| 0,000524788289 |
| 0,007251739088 |

| |
|---|
| 0 |
| 0,000000060960 |
| 0,000000000180 |
| 0,00000000117 |
| 0,002400293699 |
| 0 |
| 0,000000090124 |
| 0,000000322571 |
| 0,000002909316 |
| 0,000005994967 |
| 0,000018022163 |
| 0,003283628736 |
| 0,000065164327 |
| 0 |
| 0,000105928166 |
| 0 |
| 0,000000051692 |
| 0,00039118961 |
| 0 |
| 0,002020803445 |
| 0,00042013447 |
| 0,000000000422 |
| 0,000004709514 |
| 0 |
| 0 |
| 0 |
| 0,000000165825 |
| 0,000060369152 |
| 0,00035445975 |
| 0,000435829740 |
| 0,000957242756 |
| 0,009737519923 |
| 0,01895733129 |
| 0 |
| 0 |
| 0,000000004288 |
| 0,00000000711 |

0,000001794221
0,000013177497
0,000147674217
0
0,000002838520
0,001184894466
0,00205503152
0,000000009680
0,000000001442
0,000002261192
0
0,000000007502
0,000028163781
0,000210748081
0,002777941043
0,003399730321
0
0
0,000000000482
0,000000024877
0,000001844391
0,000003252927
0,000248026189
0,000317043830
0,000499966971
0,001011620866
0
0,000000004156
0,000009922489
0,000012935209
0
0,000001256866
0,000174540483
0,000285581733
0,01920445914
0
0,000015089169

| |
|---|
| 0,01310902466 |
| 0,000016271502 |
| 0,000336356586 |
| 0,000346976367 |
| 0,000453232733 |
| 0,004712346377 |
| 0,005942329345 |
| 0,01099654482 |
| 0,000001169128 |
| 0,000025660422 |
| 0,004928764078 |
| 0,01808288977 |
| 0 |
| 0 |
| 0,000001122535 |
| 0,000024560369 |
| 0,000192119989 |
| 0,008168122652 |
| 0,01222539259 |
| 0,01816890297 |
| 0 |
| 0 |
| 0 |
| 0 |
| 0,000000031448 |
| 0,000000354961 |
| 0,000503177744 |
| 0,00061530277 |
| 0,002204143436 |
| 0,00577821285 |
| 0,02095215306 |
| 0,02095215306 |
| 0 |
| 0 |
| 0 |
| 0 |
| 0,000008608644 |

0,000019108420
0,000037600818
0,000142671276
0,000246475313
0,000246475313
0,000380096790
0,000583800719
0,000722358904
0,000722358904
0,001358970243
0,002281180361
0,00252756729
0,008391242944
0,01121190616
0,01391537551
0,01391537551

| | | | | | |
|---|---|---|---|---|---|
| Hydrozetes lacustris | GO:0006334 | biological_process | nucleosome assembly | 92 | 0 |
| Hydrozetes lacustris | GO:0032197 | biological_process | transposition, RNA-mediated | 63 | 0 |
| Hydrozetes lacustris | GO:0005975 | biological_process | carbohydrate metabolic process | 61 | 0,000000065257 |
| Hydrozetes lacustris | GO:0006313 | biological_process | transposition, DNA-mediated | 60 | 0 |
| Hydrozetes lacustris | GO:0015074 | biological_process | DNA integration | 52 | 0 |
| Hydrozetes lacustris | GO:0009313 | biological_process | oligosaccharide catabolic process | 47 | 0 |
| Hydrozetes lacustris | GO:0055085 | biological_process | transmembrane transport | 42 | 0,000000004589 |
| Hydrozetes lacustris | GO:0003964 | molecular_function | RNA-directed DNA polymerase activity | 40 | 0 |
| Hydrozetes lacustris | GO:0004222 | molecular_function | metalloendopeptidase activity | 37 | 0 |
| Hydrozetes lacustris | GO:0001944 | biological_process | vasculature development | 33 | 0 |
| Hydrozetes lacustris | GO:0007275 | biological_process | multicellular organismal development | 29 | 0,001573988178 |
| Hydrozetes lacustris | GO:0007283 | biological_process | spermatogenesis | 26 | 0 |
| Hydrozetes lacustris | GO:0048864 | biological_process | stem cell development | 25 | 0 |
| Hydrozetes lacustris | GO:0030245 | biological_process | cellulose catabolic process | 23 | 0,002988491481 |
| Hydrozetes lacustris | GO:0006623 | biological_process | protein targeting to vacuole | 20 | 0 |
| Hydrozetes lacustris | GO:0006511 | biological_process | ubiquitin-dependent protein catabolic process | 19 | 0 |
| Hydrozetes lacustris | GO:0006310 | biological_process | DNA recombination | 19 | 0,000000112286 |
| Hydrozetes lacustris | GO:0007601 | biological_process | visual perception | 17 | 0,000000003040 |
| Hydrozetes lacustris | GO:0031647 | biological_process | regulation of protein stability | 17 | 0,008173101144 |
| Hydrozetes lacustris | GO:0051384 | biological_process | response to glucocorticoid | 15 | 0,001989519454 |
| Hydrozetes lacustris | GO:0009617 | biological_process | response to bacterium | 14 | 0,000010866177 |
| Hydrozetes lacustris | GO:0006508 | biological_process | proteolysis | 13 | 0 |
| Hydrozetes lacustris | GO:0046685 | biological_process | response to arsenic-containing substance | 13 | 0,000000000543 |
| Hydrozetes lacustris | GO:0001756 | biological_process | somitogenesis | 13 | 0,01591226671 |
| Hydrozetes lacustris | GO:0036324 | biological_process | vascular endothelial growth factor receptor-2 signaling pathway | 12 | 0,000000000398 |
| Hydrozetes lacustris | GO:0006941 | biological_process | striated muscle contraction | 12 | 0,000000117028 |
| Hydrozetes lacustris | GO:0016491 | molecular_function | oxidoreductase activity | 12 | 0,000204229924 |
| Hydrozetes lacustris | GO:0070633 | biological_process | transepithelial transport | 11 | 0,000043777665 |
| Hydrozetes lacustris | GO:0046355 | biological_process | mannan catabolic process | 11 | 0,000352856240 |
| Hydrozetes lacustris | GO:0006412 | biological_process | translation | 10 | 0 |
| Hydrozetes lacustris | GO:0044179 | biological_process | hemolysis in other organism | 10 | 0,000053718280 |
| Hydrozetes lacustris | GO:0009698 | biological_process | phenylpropanoid metabolic process | 10 | 0,002676494408 |
| Hydrozetes lacustris | GO:0005634 | cellular_component | nucleus | 10 | 0,003487910287 |
| Hydrozetes lacustris | GO:0090023 | biological_process | positive regulation of neutrophil chemotaxis | 10 | 0,008171201107 |
| Hydrozetes lacustris | GO:0043890 | molecular_function | N-acetylgalactosamine-6-sulfatase activity | 9 | 0 |
| Hydrozetes lacustris | GO:0008484 | molecular_function | sulfuric ester hydrolase activity | 9 | 0,000027131653 |
| Hydrozetes lacustris | GO:0042311 | biological_process | vasodilation | 9 | 0,02408707539 |

| | | | | | |
|---|---|---|---|---|---|
| Hydrozetes lacustris | GO:0006486 | biological_process | protein glycosylation | 8 | 0,000000010674 |
| Hydrozetes lacustris | GO:0000272 | biological_process | polysaccharide catabolic process | 8 | 0,000234943348 |
| Hydrozetes lacustris | GO:0016567 | biological_process | protein ubiquitination | 7 | 0 |
| Hydrozetes lacustris | GO:0031098 | biological_process | stress-activated protein kinase signaling cascade | 7 | 0,000126882407 |
| Hydrozetes lacustris | GO:0006470 | biological_process | protein dephosphorylation | 7 | 0,000906324858 |
| Hydrozetes lacustris | GO:0016747 | molecular_function | transferase activity, transferring acyl groups other than amino-acyl groups | 7 | 0,01363416516 |
| Hydrozetes lacustris | GO:0008354 | biological_process | germ cell migration | 6 | 0,000012313401 |
| Hydrozetes lacustris | GO:0039648 | biological_process | modulation by virus of host protein ubiquitination | 6 | 0,007084813425 |
| Hydrozetes lacustris | GO:0046470 | biological_process | phosphatidylcholine metabolic process | 6 | 0,007084813425 |
| Hydrozetes lacustris | GO:0043066 | biological_process | negative regulation of apoptotic process | 6 | 0,008166564509 |
| Hydrozetes lacustris | GO:0015893 | biological_process | drug transport | 5 | 0,000000628232 |
| Hydrozetes lacustris | GO:0036372 | biological_process | opsin transport | 5 | 0,000000628232 |
| Hydrozetes lacustris | GO:0051260 | biological_process | protein homooligomerization | 5 | 0,000042174021 |
| Hydrozetes lacustris | GO:0036323 | biological_process | vascular endothelial growth factor receptor-1 signaling pathway | 5 | 0,000330431218 |
| Hydrozetes lacustris | GO:0047936 | molecular_function | glucose 1-dehydrogenase [NAD(P)] activity | 5 | 0,01827618205 |
| Hydrozetes lacustris | GO:0016055 | biological_process | Wnt signaling pathway | 4 | 0 |
| Hydrozetes lacustris | GO:0046872 | molecular_function | metal ion binding | 4 | 0 |
| Hydrozetes lacustris | GO:0042060 | biological_process | wound healing | 4 | 0,000009667151 |
| Hydrozetes lacustris | GO:0031149 | biological_process | sorocarp stalk cell differentiation | 4 | 0,000051966719 |
| Hydrozetes lacustris | GO:0009742 | biological_process | brassinosteroid mediated signaling pathway | 4 | 0,000051966719 |
| Hydrozetes lacustris | GO:0034405 | biological_process | response to fluid shear stress | 4 | 0,000147011473 |
| Hydrozetes lacustris | GO:2000504 | biological_process | positive regulation of blood vessel remodeling | 4 | 0,000323463878 |
| Hydrozetes lacustris | GO:0055114 | biological_process | oxidation-reduction process | 4 | 0,001560912099 |
| Hydrozetes lacustris | GO:0006809 | biological_process | nitric oxide biosynthetic process | 4 | 0,006130313579 |
| Hydrozetes lacustris | GO:0010960 | biological_process | magnesium ion homeostasis | 4 | 0,006130313579 |
| Hydrozetes lacustris | GO:1902236 | biological_process | negative regulation of endoplasmic reticulum stress-induced intrinsic apoptotic signaling pathway | 4 | 0,01480805889 |
| Hydrozetes lacustris | GO:0006325 | biological_process | chromatin organization | 3 | 0,000025316639 |
| Hydrozetes lacustris | GO:0001669 | cellular_component | acrosomal vesicle | 3 | 0,000093831719 |
| Hydrozetes lacustris | GO:0042135 | biological_process | neurotransmitter catabolic process | 3 | 0,000141532344 |
| Hydrozetes lacustris | GO:0090073 | biological_process | positive regulation of protein homodimerization activity | 3 | 0,000726008444 |
| Hydrozetes lacustris | GO:0007179 | biological_process | transforming growth factor beta receptor signaling pathway | 3 | 0,001661692236 |
| Hydrozetes lacustris | GO:0006260 | biological_process | DNA replication | 3 | 0,005010048742 |
| Hydrozetes lacustris | GO:0010224 | biological_process | response to UV-B | 3 | 0,005985177732 |
| Hydrozetes lacustris | GO:0032482 | biological_process | Rab protein signal transduction | 3 | 0,01194849516 |
| Hydrozetes lacustris | GO:0070292 | biological_process | N-acylphosphatidylethanolamine metabolic process | 3 | 0,02490202891 |
| Hydrozetes lacustris | GO:0016021 | cellular_component | integral component of membrane | 2 | 0 |

| | | | | | |
|---|---|---|---|---|---|
| Hydrozetes lacustris | GO:0006357 | biological_process | regulation of transcription from RNA polymerase II promoter | 2 | 0 |
| Hydrozetes lacustris | GO:0045892 | biological_process | negative regulation of transcription, DNA-templated | 2 | 0,000026134701 |
| Hydrozetes lacustris | GO:0030435 | biological_process | sporulation resulting in formation of a cellular spore | 2 | 0,000299717026 |
| Hydrozetes lacustris | GO:0007399 | biological_process | nervous system development | 2 | 0,00033309836 |
| Hydrozetes lacustris | GO:0007217 | biological_process | tachykinin receptor signaling pathway | 2 | 0,003152983288 |
| Hydrozetes lacustris | GO:0070445 | biological_process | regulation of oligodendrocyte progenitor proliferation | 2 | 0,003348505604 |
| Hydrozetes lacustris | GO:0006641 | biological_process | triglyceride metabolic process | 2 | 0,004228488498 |
| Hydrozetes lacustris | GO:0051301 | biological_process | cell division | 2 | 0,009075466584 |
| Hydrozetes lacustris | GO:1901557 | biological_process | response to fenofibrate | 2 | 0,009467945747 |

| | | | | | |
|---|---|---|---|---|---|
| Achipteria coleoptrata | GO:0006517 | biological_process | protein deglycosylation | 63 | 0 |
| Achipteria coleoptrata | GO:0010025 | biological_process | wax biosynthetic process | 58 | 0 |
| Achipteria coleoptrata | GO:0006680 | biological_process | glucosylceramide catabolic process | 56 | 0 |
| Achipteria coleoptrata | GO:0000272 | biological_process | polysaccharide catabolic process | 54 | 0,000000738047 |
| Achipteria coleoptrata | GO:0030245 | biological_process | cellulose catabolic process | 42 | 0,000000019818 |
| Achipteria coleoptrata | GO:0006260 | biological_process | DNA replication | 33 | 0,000000464031 |
| Achipteria coleoptrata | GO:0050661 | molecular_function | NADP binding | 32 | 0 |
| Achipteria coleoptrata | GO:0000270 | biological_process | peptidoglycan metabolic process | 30 | 0 |
| Achipteria coleoptrata | GO:0007165 | biological_process | signal transduction | 29 | 0 |
| Achipteria coleoptrata | GO:0006508 | biological_process | proteolysis | 28 | 0 |
| Achipteria coleoptrata | GO:0019284 | biological_process | L-methionine biosynthetic process from S-adenosylmethionine | 28 | 0,000059137386 |
| Achipteria coleoptrata | GO:0009617 | biological_process | response to bacterium | 27 | 0,000426815864 |
| Achipteria coleoptrata | GO:0030162 | biological_process | regulation of proteolysis | 25 | 0,000000254087 |
| Achipteria coleoptrata | GO:0005975 | biological_process | carbohydrate metabolic process | 25 | 0,000619731233 |
| Achipteria coleoptrata | GO:0016139 | biological_process | glycoside catabolic process | 24 | 0,000000000691 |
| Achipteria coleoptrata | GO:0045214 | biological_process | sarcomere organization | 24 | 0,02507853578 |
| Achipteria coleoptrata | GO:0045599 | biological_process | negative regulation of fat cell differentiation | 23 | 0,000000024499 |
| Achipteria coleoptrata | GO:1901998 | biological_process | toxin transport | 23 | 0,000188648277 |
| Achipteria coleoptrata | GO:0010383 | biological_process | cell wall polysaccharide metabolic process | 22 | 0 |
| Achipteria coleoptrata | GO:0009268 | biological_process | response to pH | 22 | 0,000000116864 |
| Achipteria coleoptrata | GO:0008270 | molecular_function | zinc ion binding | 22 | 0,000000135931 |
| Achipteria coleoptrata | GO:0055085 | biological_process | transmembrane transport | 21 | 0 |
| Achipteria coleoptrata | GO:0043171 | biological_process | peptide catabolic process | 20 | 0 |
| Achipteria coleoptrata | GO:0051384 | biological_process | response to glucocorticoid | 19 | 0,000264795358 |
| Achipteria coleoptrata | GO:0004867 | molecular_function | serine-type endopeptidase inhibitor activity | 19 | 0,0244640084 |
| Achipteria coleoptrata | GO:0051480 | biological_process | cytosolic calcium ion homeostasis | 18 | 0,007439554246 |
| Achipteria coleoptrata | GO:0003677 | molecular_function | DNA binding | 17 | 0,003986822744 |
| Achipteria coleoptrata | GO:0016021 | cellular_component | integral component of membrane | 16 | 0 |
| Achipteria coleoptrata | GO:0046373 | biological_process | L-arabinose metabolic process | 16 | 0,000000000293 |
| Achipteria coleoptrata | GO:0006468 | biological_process | protein phosphorylation | 16 | 0,000000267624 |
| Achipteria coleoptrata | GO:0002376 | biological_process | immune system process | 16 | 0,000377978737 |
| Achipteria coleoptrata | GO:0015267 | molecular_function | channel activity | 16 | 0,000488116639 |
| Achipteria coleoptrata | GO:0016042 | biological_process | lipid catabolic process | 15 | 0,000099687592 |
| Achipteria coleoptrata | GO:0016614 | molecular_function | oxidoreductase activity, acting on CH-OH group of donors | 15 | 0,02090833368 |
| Achipteria coleoptrata | GO:0045087 | biological_process | innate immune response | 15 | 0,02113781106 |
| Achipteria coleoptrata | GO:0006511 | biological_process | ubiquitin-dependent protein catabolic process | 14 | 0 |
| Achipteria coleoptrata | GO:0016055 | biological_process | Wnt signaling pathway | 14 | 0 |

| | | | | | |
|---|---|---|---|---|---|
| Achipteria coleoptrata | GO:0010139 | biological_process | pyrimidine deoxyribonucleotide salvage | 14 | 0 |
| Achipteria coleoptrata | GO:0043181 | biological_process | vacuolar sequestering | 14 | 0 |
| Achipteria coleoptrata | GO:0007605 | biological_process | sensory perception of sound | 14 | 0,01590621918 |
| Achipteria coleoptrata | GO:0071555 | biological_process | cell wall organization | 14 | 0,02467375645 |
| Achipteria coleoptrata | GO:0001649 | biological_process | osteoblast differentiation | 13 | 0,000000643738 |
| Achipteria coleoptrata | GO:0005980 | biological_process | glycogen catabolic process | 13 | 0,000024467843 |
| Achipteria coleoptrata | GO:0009253 | biological_process | peptidoglycan catabolic process | 13 | 0,001406064411 |
| Achipteria coleoptrata | GO:0004869 | molecular_function | cysteine-type endopeptidase inhibitor activity | 12 | 0,000015725490 |
| Achipteria coleoptrata | GO:0051603 | biological_process | proteolysis involved in cellular protein catabolic process | 12 | 0,01748761338 |
| Achipteria coleoptrata | GO:0007283 | biological_process | spermatogenesis | 11 | 0 |
| Achipteria coleoptrata | GO:0070633 | biological_process | transepithelial transport | 11 | 0,000471104960 |
| Achipteria coleoptrata | GO:0003309 | biological_process | type B pancreatic cell differentiation | 11 | 0,001109825234 |
| Achipteria coleoptrata | GO:0043097 | biological_process | pyrimidine nucleoside salvage | 11 | 0,02038223955 |
| Achipteria coleoptrata | GO:0006706 | biological_process | steroid catabolic process | 11 | 0,02134254079 |
| Achipteria coleoptrata | GO:0007601 | biological_process | visual perception | 10 | 0 |
| Achipteria coleoptrata | GO:0031098 | biological_process | stress-activated protein kinase signaling cascade | 10 | 0,000024841454 |
| Achipteria coleoptrata | GO:0000922 | cellular_component | spindle pole | 10 | 0,000598602514 |
| Achipteria coleoptrata | GO:0016488 | biological_process | farnesol catabolic process | 9 | 0,000021799665 |
| Achipteria coleoptrata | GO:0009263 | biological_process | deoxyribonucleotide biosynthetic process | 9 | 0,000596754722 |
| Achipteria coleoptrata | GO:0019882 | biological_process | antigen processing and presentation | 9 | 0,001121361729 |
| Achipteria coleoptrata | GO:0034314 | biological_process | Arp2/3 complex-mediated actin nucleation | 9 | 0,009227445559 |
| Achipteria coleoptrata | GO:0051598 | biological_process | meiotic recombination checkpoint | 8 | 0,000000001056 |
| Achipteria coleoptrata | GO:0021762 | biological_process | substantia nigra development | 8 | 0,000696539193 |
| Achipteria coleoptrata | GO:0010508 | biological_process | positive regulation of autophagy | 8 | 0,005855040059 |
| Achipteria coleoptrata | GO:0044179 | biological_process | hemolysis in other organism | 8 | 0,006769696039 |
| Achipteria coleoptrata | GO:0046872 | molecular_function | metal ion binding | 7 | 0 |
| Achipteria coleoptrata | GO:0004559 | molecular_function | alpha-mannosidase activity | 7 | 0,000001345414 |
| Achipteria coleoptrata | GO:0042060 | biological_process | wound healing | 7 | 0,000003991693 |
| Achipteria coleoptrata | GO:0008202 | biological_process | steroid metabolic process | 7 | 0,01500218211 |
| Achipteria coleoptrata | GO:0046693 | biological_process | sperm storage | 7 | 0,01925859115 |
| Achipteria coleoptrata | GO:0004252 | molecular_function | serine-type endopeptidase activity | 6 | 0 |
| Achipteria coleoptrata | GO:0006313 | biological_process | transposition, DNA-mediated | 6 | 0,000000037441 |
| Achipteria coleoptrata | GO:0030433 | biological_process | ER-associated ubiquitin-dependent protein catabolic process | 6 | 0,000000070574 |
| Achipteria coleoptrata | GO:0048794 | biological_process | swim bladder development | 6 | 0,000000188932 |
| Achipteria coleoptrata | GO:0016020 | cellular_component | membrane | 6 | 0,003863743592 |
| Achipteria coleoptrata | GO:0047936 | molecular_function | glucose 1-dehydrogenase [NAD(P)] activity | 6 | 0,01489340579 |
| Achipteria coleoptrata | GO:0006412 | biological_process | translation | 5 | 0 |

| | | | | | |
|---|---|---|---|---|---|
| Achipteria coleoptrata | GO:0006633 | biological_process | fatty acid biosynthetic process | 5 | 0,000000000330 |
| Achipteria coleoptrata | GO:0042761 | biological_process | very long-chain fatty acid biosynthetic process | 5 | 0,000000003938 |
| Achipteria coleoptrata | GO:0018394 | biological_process | peptidyl-lysine acetylation | 5 | 0,000013951908 |
| Achipteria coleoptrata | GO:0002175 | biological_process | protein localization to paranode region of axon | 5 | 0,001731083962 |
| Achipteria coleoptrata | GO:0006355 | biological_process | regulation of transcription, DNA-templated | 4 | 0 |
| Achipteria coleoptrata | GO:0015031 | biological_process | protein transport | 4 | 0 |
| Achipteria coleoptrata | GO:0016567 | biological_process | protein ubiquitination | 4 | 0 |
| Achipteria coleoptrata | GO:0007275 | biological_process | multicellular organismal development | 4 | 0 |
| Achipteria coleoptrata | GO:0005576 | cellular_component | extracellular region | 4 | 0,000001241940 |
| Achipteria coleoptrata | GO:0055114 | biological_process | oxidation-reduction process | 4 | 0,000049649558 |
| Achipteria coleoptrata | GO:0031149 | biological_process | sorocarp stalk cell differentiation | 4 | 0,000154183986 |
| Achipteria coleoptrata | GO:0009742 | biological_process | brassinosteroid mediated signaling pathway | 4 | 0,000154183986 |
| Achipteria coleoptrata | GO:0000209 | biological_process | protein polyubiquitination | 4 | 0,000264946588 |
| Achipteria coleoptrata | GO:0051958 | biological_process | methotrexate transport | 4 | 0,000427672188 |
| Achipteria coleoptrata | GO:0004674 | molecular_function | protein serine/threonine kinase activity | 4 | 0,000908183838 |
| Achipteria coleoptrata | GO:0075732 | biological_process | viral penetration into host nucleus | 4 | 0,000922632967 |
| Achipteria coleoptrata | GO:0009213 | biological_process | pyrimidine deoxyribonucleoside triphosphate catabolic process | 4 | 0,001706046343 |
| Achipteria coleoptrata | GO:0006629 | biological_process | lipid metabolic process | 4 | 0,002202565605 |
| Achipteria coleoptrata | GO:0043129 | biological_process | surfactant homeostasis | 4 | 0,01919938141 |
| Achipteria coleoptrata | GO:0001836 | biological_process | release of cytochrome c from mitochondria | 4 | 0,01919938141 |
| Achipteria coleoptrata | GO:0048364 | biological_process | root development | 4 | 0,0236637265 |
| Achipteria coleoptrata | GO:0016032 | biological_process | viral process | 3 | 0 |
| Achipteria coleoptrata | GO:0006805 | biological_process | xenobiotic metabolic process | 3 | 0 |
| Achipteria coleoptrata | GO:0005634 | cellular_component | nucleus | 3 | 0,000000001199 |
| Achipteria coleoptrata | GO:0009636 | biological_process | response to toxic substance | 3 | 0,000000002826 |
| Achipteria coleoptrata | GO:0006325 | biological_process | chromatin organization | 3 | 0,000000186051 |
| Achipteria coleoptrata | GO:0001669 | cellular_component | acrosomal vesicle | 3 | 0,000001174179 |
| Achipteria coleoptrata | GO:0015074 | biological_process | DNA integration | 3 | 0,000026904060 |
| Achipteria coleoptrata | GO:0006694 | biological_process | steroid biosynthetic process | 3 | 0,000086294980 |
| Achipteria coleoptrata | GO:0048511 | biological_process | rhythmic process | 3 | 0,000483771201 |
| Achipteria coleoptrata | GO:0006641 | biological_process | triglyceride metabolic process | 3 | 0,001518539858 |
| Achipteria coleoptrata | GO:0008168 | molecular_function | methyltransferase activity | 3 | 0,005736752715 |
| Achipteria coleoptrata | GO:0050727 | biological_process | regulation of inflammatory response | 3 | 0,01126701439 |
| Achipteria coleoptrata | GO:0046547 | molecular_function | trans-aconitate 3-methyltransferase activity | 3 | 0,01666238595 |
| Achipteria coleoptrata | GO:0003281 | biological_process | ventricular septum development | 3 | 0,02183513667 |
| Achipteria coleoptrata | GO:0016192 | biological_process | vesicle-mediated transport | 2 | 0 |
| Achipteria coleoptrata | GO:0006814 | biological_process | sodium ion transport | 2 | 0 |

| | | | | | |
|---|---|---|---|---|---|
| Achipteria coleoptrata | GO:0051260 | biological_process | protein homooligomerization | 2 | 0,000000000451 |
| Achipteria coleoptrata | GO:0004497 | molecular_function | monooxygenase activity | 2 | 0,000048758836 |
| Achipteria coleoptrata | GO:0010224 | biological_process | response to UV-B | 2 | 0,000092775498 |
| Achipteria coleoptrata | GO:0007608 | biological_process | sensory perception of smell | 2 | 0,000248604450 |
| Achipteria coleoptrata | GO:0004222 | molecular_function | metalloendopeptidase activity | 2 | 0,000496486490 |
| Achipteria coleoptrata | GO:0006915 | biological_process | apoptotic process | 2 | 0,000916205010 |
| Achipteria coleoptrata | GO:0023021 | biological_process | termination of signal transduction | 2 | 0,00146509648 |
| Achipteria coleoptrata | GO:0071622 | biological_process | regulation of granulocyte chemotaxis | 2 | 0,005812019496 |
| Achipteria coleoptrata | GO:0070445 | biological_process | regulation of oligodendrocyte progenitor proliferation | 2 | 0,005812019496 |
| Achipteria coleoptrata | GO:0030163 | biological_process | protein catabolic process | 2 | 0,007848447567 |
| Achipteria coleoptrata | GO:0060625 | biological_process | regulation of protein deneddylation | 2 | 0,0161131336 |

| | | | | | |
|---|---|---|---|---|---|
| Platynothrus peltifer | GO:0055085 | biological_process | transmembrane transport | 356 | 0 |
| Platynothrus peltifer | GO:0006517 | biological_process | protein deglycosylation | 97 | 0 |
| Platynothrus peltifer | GO:0042908 | biological_process | xenobiotic transport | 96 | 0 |
| Platynothrus peltifer | GO:0016020 | cellular_component | membrane | 74 | 0 |
| Platynothrus peltifer | GO:0008218 | biological_process | bioluminescence | 71 | 0 |
| Platynothrus peltifer | GO:0003677 | molecular_function | DNA binding | 69 | 0,000006232143 |
| Platynothrus peltifer | GO:0003081 | biological_process | regulation of systemic arterial blood pressure by renin-angiotensin | 62 | 0 |
| Platynothrus peltifer | GO:0006680 | biological_process | glucosylceramide catabolic process | 56 | 0 |
| Platynothrus peltifer | GO:0006511 | biological_process | ubiquitin-dependent protein catabolic process | 53 | 0 |
| Platynothrus peltifer | GO:0008270 | molecular_function | zinc ion binding | 51 | 0,000114372170 |
| Platynothrus peltifer | GO:0042302 | molecular_function | structural constituent of cuticle | 50 | 0,01971298025 |
| Platynothrus peltifer | GO:0009617 | biological_process | response to bacterium | 47 | 0,000909979529 |
| Platynothrus peltifer | GO:0030245 | biological_process | cellulose catabolic process | 43 | 0,000185125395 |
| Platynothrus peltifer | GO:0006013 | biological_process | mannose metabolic process | 41 | 0 |
| Platynothrus peltifer | GO:0016266 | biological_process | O-glycan processing | 38 | 0 |
| Platynothrus peltifer | GO:0006508 | biological_process | proteolysis | 37 | 0 |
| Platynothrus peltifer | GO:0045493 | biological_process | xylan catabolic process | 37 | 0 |
| Platynothrus peltifer | GO:1902358 | biological_process | sulfate transmembrane transport | 36 | 0 |
| Platynothrus peltifer | GO:0004222 | molecular_function | metalloendopeptidase activity | 36 | 0,000002648715 |
| Platynothrus peltifer | GO:0004252 | molecular_function | serine-type endopeptidase activity | 36 | 0,000003654541 |
| Platynothrus peltifer | GO:0010224 | biological_process | response to UV-B | 36 | 0,000076476323 |
| Platynothrus peltifer | GO:0016042 | biological_process | lipid catabolic process | 32 | 0,001614728013 |
| Platynothrus peltifer | GO:0006313 | biological_process | transposition, DNA-mediated | 32 | 0,01198886847 |
| Platynothrus peltifer | GO:0055129 | biological_process | L-proline biosynthetic process | 31 | 0,000000027677 |
| Platynothrus peltifer | GO:0046872 | molecular_function | metal ion binding | 30 | 0 |
| Platynothrus peltifer | GO:0009405 | biological_process | pathogenesis | 29 | 0,002677776761 |
| Platynothrus peltifer | GO:0046548 | biological_process | retinal rod cell development | 28 | 0 |
| Platynothrus peltifer | GO:0004518 | molecular_function | nuclease activity | 27 | 0,000000005304 |
| Platynothrus peltifer | GO:0002376 | biological_process | immune system process | 27 | 0,000000197921 |
| Platynothrus peltifer | GO:0007601 | biological_process | visual perception | 26 | 0 |
| Platynothrus peltifer | GO:0006915 | biological_process | apoptotic process | 26 | 0,003009335471 |
| Platynothrus peltifer | GO:0010992 | biological_process | ubiquitin homeostasis | 25 | 0 |
| Platynothrus peltifer | GO:0007275 | biological_process | multicellular organismal development | 25 | 0 |
| Platynothrus peltifer | GO:0031048 | biological_process | chromatin silencing by small RNA | 25 | 0 |
| Platynothrus peltifer | GO:0007283 | biological_process | spermatogenesis | 24 | 0 |
| Platynothrus peltifer | GO:0045616 | biological_process | regulation of keratinocyte differentiation | 24 | 0,000002845724 |
| Platynothrus peltifer | GO:0001516 | biological_process | prostaglandin biosynthetic process | 24 | 0,000008803760 |

| | | | | | |
|---|---|---|---|---|---|
| Platynothrus peltifer | GO:0019882 | biological_process | antigen processing and presentation | 23 | 0 |
| Platynothrus peltifer | GO:0045599 | biological_process | negative regulation of fat cell differentiation | 23 | 0,000022706999 |
| Platynothrus peltifer | GO:1901998 | biological_process | toxin transport | 23 | 0,01696649243 |
| Platynothrus peltifer | GO:0010383 | biological_process | cell wall polysaccharide metabolic process | 22 | 0 |
| Platynothrus peltifer | GO:0045087 | biological_process | innate immune response | 22 | 0,00809660447 |
| Platynothrus peltifer | GO:0016192 | biological_process | vesicle-mediated transport | 21 | 0 |
| Platynothrus peltifer | GO:0051384 | biological_process | response to glucocorticoid | 21 | 0,000000013762 |
| Platynothrus peltifer | GO:0007169 | biological_process | transmembrane receptor protein tyrosine kinase signaling pathway | 21 | 0,008182908292 |
| Platynothrus peltifer | GO:0006623 | biological_process | protein targeting to vacuole | 20 | 0 |
| Platynothrus peltifer | GO:0043171 | biological_process | peptide catabolic process | 20 | 0 |
| Platynothrus peltifer | GO:0035093 | biological_process | spermatogenesis, exchange of chromosomal proteins | 20 | 0,000003266061 |
| Platynothrus peltifer | GO:0008210 | biological_process | estrogen metabolic process | 19 | 0 |
| Platynothrus peltifer | GO:0009636 | biological_process | response to toxic substance | 19 | 0,000081800010 |
| Platynothrus peltifer | GO:0019216 | biological_process | regulation of lipid metabolic process | 19 | 0,01506770755 |
| Platynothrus peltifer | GO:0030162 | biological_process | regulation of proteolysis | 19 | 0,01621830043 |
| Platynothrus peltifer | GO:0006355 | biological_process | regulation of transcription, DNA-templated | 18 | 0 |
| Platynothrus peltifer | GO:0009698 | biological_process | phenylpropanoid metabolic process | 18 | 0,000141908823 |
| Platynothrus peltifer | GO:0006805 | biological_process | xenobiotic metabolic process | 17 | 0,000000002575 |
| Platynothrus peltifer | GO:0008146 | molecular_function | sulfotransferase activity | 17 | 0,000000034400 |
| Platynothrus peltifer | GO:0051923 | biological_process | sulfation | 17 | 0,002106769963 |
| Platynothrus peltifer | GO:0008234 | molecular_function | cysteine-type peptidase activity | 17 | 0,02317202479 |
| Platynothrus peltifer | GO:0097310 | biological_process | cap2 mRNA methylation | 16 | 0 |
| Platynothrus peltifer | GO:0046373 | biological_process | L-arabinose metabolic process | 16 | 0,000000090443 |
| Platynothrus peltifer | GO:0045332 | biological_process | phospholipid translocation | 16 | 0,00014103628 |
| Platynothrus peltifer | GO:0034976 | biological_process | response to endoplasmic reticulum stress | 16 | 0,000153469675 |
| Platynothrus peltifer | GO:0043181 | biological_process | vacuolar sequestering | 14 | 0,000000000893 |
| Platynothrus peltifer | GO:0001507 | biological_process | acetylcholine catabolic process in synaptic cleft | 14 | 0,000725351246 |
| Platynothrus peltifer | GO:0007021 | biological_process | tubulin complex assembly | 14 | 0,02592491341 |
| Platynothrus peltifer | GO:0010718 | biological_process | positive regulation of epithelial to mesenchymal transition | 14 | 0,02798901282 |
| Platynothrus peltifer | GO:0045944 | biological_process | positive regulation of transcription from RNA polymerase II promoter | 13 | 0 |
| Platynothrus peltifer | GO:0033089 | biological_process | positive regulation of T cell differentiation in thymus | 13 | 0,000000005671 |
| Platynothrus peltifer | GO:0042060 | biological_process | wound healing | 13 | 0,000000384498 |
| Platynothrus peltifer | GO:0034612 | biological_process | response to tumor necrosis factor | 13 | 0,000033526271 |
| Platynothrus peltifer | GO:0019233 | biological_process | sensory perception of pain | 13 | 0,000897962864 |
| Platynothrus peltifer | GO:0005980 | biological_process | glycogen catabolic process | 13 | 0,001103489105 |
| Platynothrus peltifer | GO:0007586 | biological_process | digestion | 13 | 0,002735063515 |
| Platynothrus peltifer | GO:0006814 | biological_process | sodium ion transport | 12 | 0 |

| | | | | | |
|---|---|---|---|---|---|
| Platynothrus peltifer | GO:0006941 | biological_process | striated muscle contraction | 12 | 0,0001155839744 |
| Platynothrus peltifer | GO:0016021 | cellular_component | integral component of membrane | 11 | 0 |
| Platynothrus peltifer | GO:0016567 | biological_process | protein ubiquitination | 11 | 0 |
| Platynothrus peltifer | GO:0014003 | biological_process | oligodendrocyte development | 11 | 0,001389966342 |
| Platynothrus peltifer | GO:0003309 | biological_process | type B pancreatic cell differentiation | 11 | 0,01667270801 |
| Platynothrus peltifer | GO:0016055 | biological_process | Wnt signaling pathway | 10 | 0 |
| Platynothrus peltifer | GO:0008380 | biological_process | RNA splicing | 10 | 0 |
| Platynothrus peltifer | GO:0055069 | biological_process | zinc ion homeostasis | 10 | 0,000030177467 |
| Platynothrus peltifer | GO:0016488 | biological_process | farnesol catabolic process | 10 | 0,000077342461 |
| Platynothrus peltifer | GO:0014883 | biological_process | transition between fast and slow fiber | 10 | 0,000174582977 |
| Platynothrus peltifer | GO:0031222 | biological_process | arabinan catabolic process | 10 | 0,003011899065 |
| Platynothrus peltifer | GO:0006412 | biological_process | translation | 9 | 0 |
| Platynothrus peltifer | GO:0006979 | biological_process | response to oxidative stress | 9 | 0 |
| Platynothrus peltifer | GO:0043890 | molecular_function | N-acetylgalactosamine-6-sulfatase activity | 9 | 0,000000002808 |
| Platynothrus peltifer | GO:0019062 | biological_process | virion attachment to host cell | 9 | 0,000000024971 |
| Platynothrus peltifer | GO:0007605 | biological_process | sensory perception of sound | 9 | 0,000000122354 |
| Platynothrus peltifer | GO:0004674 | molecular_function | protein serine/threonine kinase activity | 9 | 0,001211642525 |
| Platynothrus peltifer | GO:0047936 | molecular_function | glucose 1-dehydrogenase [NAD(P)] activity | 9 | 0,002658017048 |
| Platynothrus peltifer | GO:0008484 | molecular_function | sulfuric ester hydrolase activity | 9 | 0,003373474836 |
| Platynothrus peltifer | GO:0031098 | biological_process | stress-activated protein kinase signaling cascade | 8 | 0 |
| Platynothrus peltifer | GO:0016614 | molecular_function | oxidoreductase activity, acting on CH-OH group of donors | 8 | 0,000000015731 |
| Platynothrus peltifer | GO:0042135 | biological_process | neurotransmitter catabolic process | 8 | 0,000001264578 |
| Platynothrus peltifer | GO:0008033 | biological_process | tRNA processing | 8 | 0,000009576096 |
| Platynothrus peltifer | GO:0008241 | molecular_function | peptidyl-dipeptidase activity | 8 | 0,000771128466 |
| Platynothrus peltifer | GO:0006749 | biological_process | glutathione metabolic process | 8 | 0,000956640665 |
| Platynothrus peltifer | GO:0047935 | molecular_function | glucose 1-dehydrogenase (NADP+) activity | 8 | 0,0211459997 |
| Platynothrus peltifer | GO:0007286 | biological_process | spermatid development | 7 | 0,000000037767 |
| Platynothrus peltifer | GO:0007399 | biological_process | nervous system development | 7 | 0,000026078147 |
| Platynothrus peltifer | GO:0008202 | biological_process | steroid metabolic process | 7 | 0,000164238874 |
| Platynothrus peltifer | GO:0030321 | biological_process | transepithelial chloride transport | 7 | 0,01881115554 |
| Platynothrus peltifer | GO:0005212 | molecular_function | structural constituent of eye lens | 7 | 0,01881115554 |
| Platynothrus peltifer | GO:0005525 | molecular_function | GTP binding | 7 | 0,02076639492 |
| Platynothrus peltifer | GO:0055114 | biological_process | oxidation-reduction process | 6 | 0,000000605498 |
| Platynothrus peltifer | GO:0048794 | biological_process | swim bladder development | 6 | 0,000002028433 |
| Platynothrus peltifer | GO:0007179 | biological_process | transforming growth factor beta receptor signaling pathway | 6 | 0,000009234203 |
| Platynothrus peltifer | GO:0051725 | biological_process | protein de-ADP-ribosylation | 6 | 0,000265180783 |
| Platynothrus peltifer | GO:0015939 | biological_process | pantothenate metabolic process | 6 | 0,000265180783 |

| | | | | | |
|---|---|---|---|---|---|
| Platynothrus peltifer | GO:0018279 | biological_process | protein N-linked glycosylation via asparagine | 6 | 0,002109778433 |
| Platynothrus peltifer | GO:0050728 | biological_process | negative regulation of inflammatory response | 6 | 0,009070423208 |
| Platynothrus peltifer | GO:0001659 | biological_process | temperature homeostasis | 6 | 0,01732291304 |
| Platynothrus peltifer | GO:0016311 | biological_process | dephosphorylation | 6 | 0,02265899831 |
| Platynothrus peltifer | GO:0035556 | biological_process | intracellular signal transduction | 6 | 0,02702136941 |
| Platynothrus peltifer | GO:0007165 | biological_process | signal transduction | 5 | 0 |
| Platynothrus peltifer | GO:0015031 | biological_process | protein transport | 5 | 0 |
| Platynothrus peltifer | GO:0006468 | biological_process | protein phosphorylation | 5 | 0 |
| Platynothrus peltifer | GO:0015893 | biological_process | drug transport | 5 | 0,000018122576 |
| Platynothrus peltifer | GO:0010044 | biological_process | response to aluminum ion | 5 | 0,000018122576 |
| Platynothrus peltifer | GO:0046693 | biological_process | sperm storage | 5 | 0,000022407153 |
| Platynothrus peltifer | GO:0031647 | biological_process | regulation of protein stability | 5 | 0,000052668629 |
| Platynothrus peltifer | GO:0050951 | biological_process | sensory perception of temperature stimulus | 5 | 0,000096564859 |
| Platynothrus peltifer | GO:0018394 | biological_process | peptidyl-lysine acetylation | 5 | 0,000096564859 |
| Platynothrus peltifer | GO:0007416 | biological_process | synapse assembly | 5 | 0,001560273277 |
| Platynothrus peltifer | GO:0030497 | biological_process | fatty acid elongation | 5 | 0,003826681021 |
| Platynothrus peltifer | GO:0001894 | biological_process | tissue homeostasis | 5 | 0,007687863762 |
| Platynothrus peltifer | GO:0001666 | biological_process | response to hypoxia | 5 | 0,01964663164 |
| Platynothrus peltifer | GO:0042770 | biological_process | signal transduction in response to DNA damage | 5 | 0,02250720944 |
| Platynothrus peltifer | GO:0005576 | cellular_component | extracellular region | 4 | 0 |
| Platynothrus peltifer | GO:0042572 | biological_process | retinol metabolic process | 4 | 0,000001422379 |
| Platynothrus peltifer | GO:0008542 | biological_process | visual learning | 4 | 0,000029839290 |
| Platynothrus peltifer | GO:0051301 | biological_process | cell division | 4 | 0,000226227332 |
| Platynothrus peltifer | GO:0007584 | biological_process | response to nutrient | 4 | 0,000750667027 |
| Platynothrus peltifer | GO:0008654 | biological_process | phospholipid biosynthetic process | 4 | 0,003328734386 |
| Platynothrus peltifer | GO:0030163 | biological_process | protein catabolic process | 4 | 0,003621694824 |
| Platynothrus peltifer | GO:0009213 | biological_process | pyrimidine deoxyribonucleoside triphosphate catabolic process | 4 | 0,007026220225 |
| Platynothrus peltifer | GO:0001932 | biological_process | regulation of protein phosphorylation | 4 | 0,01122717199 |
| Platynothrus peltifer | GO:1900426 | biological_process | positive regulation of defense response to bacterium | 4 | 0,01661042673 |
| Platynothrus peltifer | GO:0048741 | biological_process | skeletal muscle fiber development | 4 | 0,02383993336 |
| Platynothrus peltifer | GO:0006633 | biological_process | fatty acid biosynthetic process | 3 | 0 |
| Platynothrus peltifer | GO:0006470 | biological_process | protein dephosphorylation | 3 | 0 |
| Platynothrus peltifer | GO:0007608 | biological_process | sensory perception of smell | 3 | 0,000005804133 |
| Platynothrus peltifer | GO:0006641 | biological_process | triglyceride metabolic process | 3 | 0,000009815927 |
| Platynothrus peltifer | GO:0006401 | biological_process | RNA catabolic process | 3 | 0,000553205439 |
| Platynothrus peltifer | GO:0003281 | biological_process | ventricular septum development | 3 | 0,000980253647 |
| Platynothrus peltifer | GO:0001505 | biological_process | regulation of neurotransmitter levels | 3 | 0,000980253647 |

| | | | | | |
|---|---|---|---|---|---|
| Platynothrus peltifer | GO:0060333 | biological_process | interferon-gamma-mediated signaling pathway | 3 | 0,001437567086 |
| Platynothrus peltifer | GO:0048489 | biological_process | synaptic vesicle transport | 3 | 0,002481167748 |
| Platynothrus peltifer | GO:0006904 | biological_process | vesicle docking involved in exocytosis | 3 | 0,004258827582 |
| Platynothrus peltifer | GO:0006310 | biological_process | DNA recombination | 3 | 0,008552882061 |
| Platynothrus peltifer | GO:0031175 | biological_process | neuron projection development | 3 | 0,008552882061 |
| Platynothrus peltifer | GO:0006614 | biological_process | SRP-dependent cotranslational protein targeting to membrane | 3 | 0,00931260617 |
| Platynothrus peltifer | GO:0051225 | biological_process | spindle assembly | 3 | 0,0119851875 |
| Platynothrus peltifer | GO:0006357 | biological_process | regulation of transcription from RNA polymerase II promoter | 2 | 0 |
| Platynothrus peltifer | GO:0016032 | biological_process | viral process | 2 | 0 |
| Platynothrus peltifer | GO:0006457 | biological_process | protein folding | 2 | 0 |
| Platynothrus peltifer | GO:0000122 | biological_process | negative regulation of transcription from RNA polymerase II promoter | 2 | 0 |
| Platynothrus peltifer | GO:0042626 | molecular_function | ATPase activity, coupled to transmembrane movement of substances | 2 | 0 |
| Platynothrus peltifer | GO:0005506 | molecular_function | iron ion binding | 2 | 0,000000000287 |
| Platynothrus peltifer | GO:0007155 | biological_process | cell adhesion | 2 | 0,000000009732 |
| Platynothrus peltifer | GO:0006694 | biological_process | steroid biosynthetic process | 2 | 0,000000009732 |
| Platynothrus peltifer | GO:0007166 | biological_process | cell surface receptor signaling pathway | 2 | 0,000000049049 |
| Platynothrus peltifer | GO:0004497 | molecular_function | monooxygenase activity | 2 | 0,000000061705 |
| Platynothrus peltifer | GO:0048511 | biological_process | rhythmic process | 2 | 0,000000193262 |
| Platynothrus peltifer | GO:0006644 | biological_process | phospholipid metabolic process | 2 | 0,000000340691 |
| Platynothrus peltifer | GO:0009408 | biological_process | response to heat | 2 | 0,000005552819 |
| Platynothrus peltifer | GO:0071625 | biological_process | vocalization behavior | 2 | 0,000006198414 |
| Platynothrus peltifer | GO:0023021 | biological_process | termination of signal transduction | 2 | 0,000014865904 |
| Platynothrus peltifer | GO:0004930 | molecular_function | G-protein coupled receptor activity | 2 | 0,000102543944 |
| Platynothrus peltifer | GO:0001501 | biological_process | skeletal system development | 2 | 0,000126635739 |
| Platynothrus peltifer | GO:0050896 | biological_process | response to stimulus | 2 | 0,000156264901 |
| Platynothrus peltifer | GO:0042981 | biological_process | regulation of apoptotic process | 2 | 0,000359299947 |
| Platynothrus peltifer | GO:0032880 | biological_process | regulation of protein localization | 2 | 0,00110159978 |
| Platynothrus peltifer | GO:0008360 | biological_process | regulation of cell shape | 2 | 0,001642350394 |
| Platynothrus peltifer | GO:0042311 | biological_process | vasodilation | 2 | 0,003264528253 |
| Platynothrus peltifer | GO:0001558 | biological_process | regulation of cell growth | 2 | 0,003960405854 |
| Platynothrus peltifer | GO:0023052 | biological_process | signaling | 2 | 0,01114232344 |
| Platynothrus peltifer | GO:0097090 | biological_process | presynaptic membrane organization | 2 | 0,01276385322 |

| | | | | | |
|---|---|---|---|---|---|
| Nothrus palustris | GO:0032100 | biological_process | positive regulation of appetite | 218 | 0 |
| Nothrus palustris | GO:0005975 | biological_process | carbohydrate metabolic process | 120 | 0 |
| Nothrus palustris | GO:0003677 | molecular_function | DNA binding | 99 | 0 |
| Nothrus palustris | GO:0006334 | biological_process | nucleosome assembly | 92 | 0 |
| Nothrus palustris | GO:0006517 | biological_process | protein deglycosylation | 84 | 0 |
| Nothrus palustris | GO:0016301 | molecular_function | kinase activity | 77 | 0 |
| Nothrus palustris | GO:0006633 | biological_process | fatty acid biosynthetic process | 77 | 0,000000097155 |
| Nothrus palustris | GO:0055085 | biological_process | transmembrane transport | 62 | 0 |
| Nothrus palustris | GO:0003964 | molecular_function | RNA-directed DNA polymerase activity | 62 | 0 |
| Nothrus palustris | GO:0004364 | molecular_function | glutathione transferase activity | 61 | 0 |
| Nothrus palustris | GO:0030223 | biological_process | neutrophil differentiation | 58 | 0 |
| Nothrus palustris | GO:0006805 | biological_process | xenobiotic metabolic process | 58 | 0,0107620803 |
| Nothrus palustris | GO:0030245 | biological_process | cellulose catabolic process | 56 | 0,000000000187 |
| Nothrus palustris | GO:0006706 | biological_process | steroid catabolic process | 51 | 0,000000100919 |
| Nothrus palustris | GO:0055129 | biological_process | L-proline biosynthetic process | 44 | 0 |
| Nothrus palustris | GO:0046982 | molecular_function | protein heterodimerization activity | 42 | 0 |
| Nothrus palustris | GO:0007283 | biological_process | spermatogenesis | 41 | 0 |
| Nothrus palustris | GO:0006508 | biological_process | proteolysis | 40 | 0 |
| Nothrus palustris | GO:0006589 | biological_process | octopamine biosynthetic process | 37 | 0 |
| Nothrus palustris | GO:0007601 | biological_process | visual perception | 37 | 0 |
| Nothrus palustris | GO:0016266 | biological_process | O-glycan processing | 36 | 0 |
| Nothrus palustris | GO:0050661 | molecular_function | NADP binding | 35 | 0 |
| Nothrus palustris | GO:0052697 | biological_process | xenobiotic glucuronidation | 35 | 0 |
| Nothrus palustris | GO:0010383 | biological_process | cell wall polysaccharide metabolic process | 33 | 0 |
| Nothrus palustris | GO:0008270 | molecular_function | zinc ion binding | 33 | 0,000000006399 |
| Nothrus palustris | GO:0005524 | molecular_function | ATP binding | 33 | 0,00024629438 |
| Nothrus palustris | GO:0042178 | biological_process | xenobiotic catabolic process | 32 | 0,000000000346 |
| Nothrus palustris | GO:0005525 | molecular_function | GTP binding | 32 | 0,000000126769 |
| Nothrus palustris | GO:0005506 | molecular_function | iron ion binding | 32 | 0,0175078807 |
| Nothrus palustris | GO:0006313 | biological_process | transposition, DNA-mediated | 31 | 0,02102724731 |
| Nothrus palustris | GO:0030162 | biological_process | regulation of proteolysis | 30 | 0,000000270128 |
| Nothrus palustris | GO:0016491 | molecular_function | oxidoreductase activity | 30 | 0,000290136082 |
| Nothrus palustris | GO:0008202 | biological_process | steroid metabolic process | 30 | 0,003499141757 |
| Nothrus palustris | GO:0007275 | biological_process | multicellular organismal development | 29 | 0 |
| Nothrus palustris | GO:0071357 | biological_process | cellular response to type I interferon | 29 | 0 |
| Nothrus palustris | GO:0004252 | molecular_function | serine-type endopeptidase activity | 29 | 0,000000223257 |
| Nothrus palustris | GO:0009888 | biological_process | tissue development | 27 | 0,000201096991 |

| | | | | | |
|---|---|---|---|---|---|
| Nothrus palustris | GO:0006749 | biological_process | glutathione metabolic process | 27 | 0,009392543342 |
| Nothrus palustris | GO:0004497 | molecular_function | monooxygenase activity | 27 | 0,01475391819 |
| Nothrus palustris | GO:0009636 | biological_process | response to toxic substance | 27 | 0,02129061838 |
| Nothrus palustris | GO:0010951 | biological_process | negative regulation of endopeptidase activity | 26 | 0,000002825781 |
| Nothrus palustris | GO:0031048 | biological_process | chromatin silencing by small RNA | 25 | 0 |
| Nothrus palustris | GO:0010842 | biological_process | retina layer formation | 25 | 0,000031437902 |
| Nothrus palustris | GO:0009058 | biological_process | biosynthetic process | 25 | 0,000093933623 |
| Nothrus palustris | GO:0044245 | biological_process | polysaccharide digestion | 24 | 0 |
| Nothrus palustris | GO:0045616 | biological_process | regulation of keratinocyte differentiation | 24 | 0,000000789310 |
| Nothrus palustris | GO:0042761 | biological_process | very long-chain fatty acid biosynthetic process | 23 | 0,000378294346 |
| Nothrus palustris | GO:0010811 | biological_process | positive regulation of cell-substrate adhesion | 22 | 0,000000230473 |
| Nothrus palustris | GO:0016042 | biological_process | lipid catabolic process | 22 | 0,000014888530 |
| Nothrus palustris | GO:0006623 | biological_process | protein targeting to vacuole | 20 | 0 |
| Nothrus palustris | GO:0010718 | biological_process | positive regulation of epithelial to mesenchymal transition | 20 | 0,000087495661 |
| Nothrus palustris | GO:0090502 | biological_process | RNA phosphodiester bond hydrolysis, endonucleolytic | 20 | 0,000445133343 |
| Nothrus palustris | GO:0030512 | biological_process | negative regulation of transforming growth factor beta receptor signaling pathway | 18 | 0,000000001307 |
| Nothrus palustris | GO:0060049 | biological_process | regulation of protein glycosylation | 18 | 0,000000004110 |
| Nothrus palustris | GO:0006013 | biological_process | mannose metabolic process | 18 | 0,007258322968 |
| Nothrus palustris | GO:0001669 | cellular_component | acrosomal vesicle | 18 | 0,01274984309 |
| Nothrus palustris | GO:0019216 | biological_process | regulation of lipid metabolic process | 18 | 0,01506867735 |
| Nothrus palustris | GO:0070189 | biological_process | kynurenine metabolic process | 17 | 0,000000959534 |
| Nothrus palustris | GO:0044179 | biological_process | hemolysis in other organism | 16 | 0,000000823132 |
| Nothrus palustris | GO:0042302 | molecular_function | structural constituent of cuticle | 15 | 0,000002361116 |
| Nothrus palustris | GO:0006090 | biological_process | pyruvate metabolic process | 15 | 0,000009133486 |
| Nothrus palustris | GO:0007130 | biological_process | synaptonemal complex assembly | 14 | 0,01373796861 |
| Nothrus palustris | GO:0007165 | biological_process | signal transduction | 13 | 0 |
| Nothrus palustris | GO:0006152 | biological_process | purine nucleoside catabolic process | 13 | 0,000000005921 |
| Nothrus palustris | GO:0051260 | biological_process | protein homooligomerization | 13 | 0,000002864049 |
| Nothrus palustris | GO:0001649 | biological_process | osteoblast differentiation | 13 | 0,000022256478 |
| Nothrus palustris | GO:0001507 | biological_process | acetylcholine catabolic process in synaptic cleft | 13 | 0,001079710574 |
| Nothrus palustris | GO:0009617 | biological_process | response to bacterium | 12 | 0 |
| Nothrus palustris | GO:0006979 | biological_process | response to oxidative stress | 12 | 0 |
| Nothrus palustris | GO:0005983 | biological_process | starch catabolic process | 12 | 0,000000652889 |
| Nothrus palustris | GO:0034976 | biological_process | response to endoplasmic reticulum stress | 12 | 0,000018462253 |
| Nothrus palustris | GO:0006694 | biological_process | steroid biosynthetic process | 12 | 0,009916169143 |
| Nothrus palustris | GO:0016055 | biological_process | Wnt signaling pathway | 11 | 0 |
| Nothrus palustris | GO:0016192 | biological_process | vesicle-mediated transport | 11 | 0 |

| | | | | | |
|---|---|---|---|---|---|
| Nothrus palustris | GO:0015020 | molecular_function | glucuronosyltransferase activity | 11 | 0,01368365052 |
| Nothrus palustris | GO:0031098 | biological_process | stress-activated protein kinase signaling cascade | 10 | 0,000000004887 |
| Nothrus palustris | GO:0006470 | biological_process | protein dephosphorylation | 10 | 0,000000300201 |
| Nothrus palustris | GO:0035249 | biological_process | synaptic transmission, glutamatergic | 10 | 0,000195799887 |
| Nothrus palustris | GO:0000723 | biological_process | telomere maintenance | 10 | 0,000280395120 |
| Nothrus palustris | GO:0016747 | molecular_function | transferase activity, transferring acyl groups other than amino-acyl groups | 10 | 0,01555818192 |
| Nothrus palustris | GO:0006355 | biological_process | regulation of transcription, DNA-templated | 9 | 0 |
| Nothrus palustris | GO:0042060 | biological_process | wound healing | 9 | 0,000000018024 |
| Nothrus palustris | GO:0042428 | biological_process | serotonin metabolic process | 9 | 0,003322287399 |
| Nothrus palustris | GO:0009405 | biological_process | pathogenesis | 9 | 0,01193621205 |
| Nothrus palustris | GO:0006511 | biological_process | ubiquitin-dependent protein catabolic process | 8 | 0 |
| Nothrus palustris | GO:0006468 | biological_process | protein phosphorylation | 8 | 0 |
| Nothrus palustris | GO:0051598 | biological_process | meiotic recombination checkpoint | 8 | 0,000000014109 |
| Nothrus palustris | GO:0000209 | biological_process | protein polyubiquitination | 8 | 0,000302058707 |
| Nothrus palustris | GO:0006954 | biological_process | inflammatory response | 8 | 0,000977125764 |
| Nothrus palustris | GO:0021762 | biological_process | substantia nigra development | 8 | 0,004921162649 |
| Nothrus palustris | GO:0032197 | biological_process | transposition, RNA-mediated | 8 | 0,02348560077 |
| Nothrus palustris | GO:0030321 | biological_process | transepithelial chloride transport | 7 | 0,01350375596 |
| Nothrus palustris | GO:0016567 | biological_process | protein ubiquitination | 6 | 0 |
| Nothrus palustris | GO:0009699 | biological_process | phenylpropanoid biosynthetic process | 6 | 0,000008232282 |
| Nothrus palustris | GO:0015074 | biological_process | DNA integration | 6 | 0,000010364237 |
| Nothrus palustris | GO:0007179 | biological_process | transforming growth factor beta receptor signaling pathway | 6 | 0,000034895813 |
| Nothrus palustris | GO:0007155 | biological_process | cell adhesion | 6 | 0,000045053767 |
| Nothrus palustris | GO:0042572 | biological_process | retinol metabolic process | 6 | 0,000095942092 |
| Nothrus palustris | GO:0007221 | biological_process | positive regulation of transcription of Notch receptor target | 6 | 0,000350373110 |
| Nothrus palustris | GO:0090129 | biological_process | positive regulation of synapse maturation | 6 | 0,003507096891 |
| Nothrus palustris | GO:0048010 | biological_process | vascular endothelial growth factor receptor signaling pathway | 6 | 0,007226881766 |
| Nothrus palustris | GO:0001501 | biological_process | skeletal system development | 6 | 0,02377827694 |
| Nothrus palustris | GO:0006412 | biological_process | translation | 5 | 0 |
| Nothrus palustris | GO:0006325 | biological_process | chromatin organization | 5 | 0,000000002900 |
| Nothrus palustris | GO:0007586 | biological_process | digestion | 5 | 0,000001996799 |
| Nothrus palustris | GO:0007399 | biological_process | nervous system development | 5 | 0,000006634112 |
| Nothrus palustris | GO:0010224 | biological_process | response to UV-B | 5 | 0,000103736350 |
| Nothrus palustris | GO:0051131 | biological_process | chaperone-mediated protein complex assembly | 5 | 0,003012677884 |
| Nothrus palustris | GO:0036323 | biological_process | vascular endothelial growth factor receptor-1 signaling pathway | 5 | 0,004626385947 |
| Nothrus palustris | GO:0046872 | molecular_function | metal ion binding | 4 | 0 |
| Nothrus palustris | GO:0004674 | molecular_function | protein serine/threonine kinase activity | 4 | 0,000009963550 |

| | | | | | |
|---|---|---|---|---|---|
| Nothrus palustris | GO:0006260 | biological_process | DNA replication | 4 | 0,000017428904 |
| Nothrus palustris | GO:0005412 | molecular_function | glucose:sodium symporter activity | 4 | 0,000120945160 |
| Nothrus palustris | GO:0006986 | biological_process | response to unfolded protein | 4 | 0,000515786714 |
| Nothrus palustris | GO:0007584 | biological_process | response to nutrient | 4 | 0,001653666675 |
| Nothrus palustris | GO:0006667 | biological_process | sphinganine metabolic process | 4 | 0,01311104721 |
| Nothrus palustris | GO:0000070 | biological_process | mitotic sister chromatid segregation | 4 | 0,01311104721 |
| Nothrus palustris | GO:0007223 | biological_process | Wnt signaling pathway, calcium modulating pathway | 4 | 0,01731081043 |
| Nothrus palustris | GO:0006364 | biological_process | rRNA processing | 3 | 0 |
| Nothrus palustris | GO:0030433 | biological_process | ER-associated ubiquitin-dependent protein catabolic process | 3 | 0 |
| Nothrus palustris | GO:0006890 | biological_process | retrograde vesicle-mediated transport, Golgi to ER | 3 | 0 |
| Nothrus palustris | GO:0051603 | biological_process | proteolysis involved in cellular protein catabolic process | 3 | 0,000000000851 |
| Nothrus palustris | GO:0007264 | biological_process | small GTPase mediated signal transduction | 3 | 0,000000012017 |
| Nothrus palustris | GO:0008542 | biological_process | visual learning | 3 | 0,000018978725 |
| Nothrus palustris | GO:0018279 | biological_process | protein N-linked glycosylation via asparagine | 3 | 0,000125793208 |
| Nothrus palustris | GO:0045087 | biological_process | innate immune response | 3 | 0,000540568865[illegible] |
| Nothrus palustris | GO:0004930 | molecular_function | G-protein coupled receptor activity | 3 | 0,001096125722 |
| Nothrus palustris | GO:0042770 | biological_process | signal transduction in response to DNA damage | 3 | 0,005921556064 |
| Nothrus palustris | GO:0030246 | molecular_function | carbohydrate binding | 3 | 0,006419787989 |
| Nothrus palustris | GO:0060074 | biological_process | synapse maturation | 3 | 0,009279058788 |
| Nothrus palustris | GO:0016032 | biological_process | viral process | 2 | 0 |
| Nothrus palustris | GO:0007605 | biological_process | sensory perception of sound | 2 | 0 |
| Nothrus palustris | GO:0010025 | biological_process | wax biosynthetic process | 2 | 0 |
| Nothrus palustris | GO:0045892 | biological_process | negative regulation of transcription, DNA-templated | 2 | 0,000000000154 |
| Nothrus palustris | GO:0000122 | biological_process | negative regulation of transcription from RNA polymerase II promoter | 2 | 0,000000000269 |
| Nothrus palustris | GO:0055114 | biological_process | oxidation-reduction process | 2 | 0,000000001743 |
| Nothrus palustris | GO:0007166 | biological_process | cell surface receptor signaling pathway | 2 | 0,000000219588 |
| Nothrus palustris | GO:0031638 | biological_process | zymogen activation | 2 | 0,000000458584 |
| Nothrus palustris | GO:0035220 | biological_process | wing disc development | 2 | 0,000000858758 |
| Nothrus palustris | GO:0006417 | biological_process | regulation of translation | 2 | 0,000003300098 |
| Nothrus palustris | GO:1902600 | biological_process | hydrogen ion transmembrane transport | 2 | 0,000016840517 |
| Nothrus palustris | GO:0008168 | molecular_function | methyltransferase activity | 2 | 0,000045844350 |
| Nothrus palustris | GO:0050896 | biological_process | response to stimulus | 2 | 0,000356666325 |
| Nothrus palustris | GO:0001894 | biological_process | tissue homeostasis | 2 | 0,000392483152 |
| Nothrus palustris | GO:2000377 | biological_process | regulation of reactive oxygen species metabolic process | 2 | 0,00047497811[illegible] |
| Nothrus palustris | GO:0035556 | biological_process | intracellular signal transduction | 2 | 0,000762414179 |
| Nothrus palustris | GO:0008039 | biological_process | synaptic target recognition | 2 | 0,001108770241 |
| Nothrus palustris | GO:0030148 | biological_process | sphingolipid biosynthetic process | 2 | 0,001761025871 |

| | | | | | |
|---|---|---|---|---|---|
| Nothrus palustris | GO:0004104 | molecular_function | cholinesterase activity | 2 | 0,002115149267 |
| Nothrus palustris | GO:0035725 | biological_process | sodium ion transmembrane transport | 2 | 0,002317126144 |
| Nothrus palustris | GO:0051290 | biological_process | protein heterotetramerization | 2 | 0,002778415126 |
| Nothrus palustris | GO:0042311 | biological_process | vasodilation | 2 | 0,005673684495 |
| Nothrus palustris | GO:0097264 | biological_process | self proteolysis | 2 | 0,009552825949 |
| Nothrus palustris | GO:0005154 | molecular_function | epidermal growth factor receptor binding | 2 | 0,01104690043 |
| Nothrus palustris | GO:0018105 | biological_process | peptidyl-serine phosphorylation | 2 | 0,01458515204 |
| Nothrus palustris | GO:0042381 | biological_process | hemolymph coagulation | 2 | 0,01458515204 |
| Nothrus palustris | GO:0051124 | biological_process | synaptic growth at neuromuscular junction | 2 | 0,02201420198 |

| | | | | | |
|---|---|---|---|---|---|
| Hermannia gibba | GO:0007283 | biological_process | spermatogenesis | 141 | 0,008916600348 |
| Hermannia gibba | GO:0005975 | biological_process | carbohydrate metabolic process | 124 | 0 |
| Hermannia gibba | GO:0016301 | molecular_function | kinase activity | 106 | 0 |
| Hermannia gibba | GO:0016020 | cellular_component | membrane | 74 | 0 |
| Hermannia gibba | GO:0051384 | biological_process | response to glucocorticoid | 73 | 0,002647787715 |
| Hermannia gibba | GO:0045214 | biological_process | sarcomere organization | 68 | 0 |
| Hermannia gibba | GO:0032197 | biological_process | transposition, RNA-mediated | 67 | 0 |
| Hermannia gibba | GO:0008218 | biological_process | bioluminescence | 63 | 0 |
| Hermannia gibba | GO:0007165 | biological_process | signal transduction | 62 | 0 |
| Hermannia gibba | GO:0006260 | biological_process | DNA replication | 54 | 0 |
| Hermannia gibba | GO:0006508 | biological_process | proteolysis | 51 | 0 |
| Hermannia gibba | GO:0009313 | biological_process | oligosaccharide catabolic process | 47 | 0 |
| Hermannia gibba | GO:0003964 | molecular_function | RNA-directed DNA polymerase activity | 46 | 0,000000015348 |
| Hermannia gibba | GO:0007286 | biological_process | spermatid development | 42 | 0,01259690361 |
| Hermannia gibba | GO:0010025 | biological_process | wax biosynthetic process | 41 | 0,007951103668 |
| Hermannia gibba | GO:0015074 | biological_process | DNA integration | 38 | 0,002572496891 |
| Hermannia gibba | GO:0007275 | biological_process | multicellular organismal development | 37 | 0 |
| Hermannia gibba | GO:0009311 | biological_process | oligosaccharide metabolic process | 36 | 0 |
| Hermannia gibba | GO:0016266 | biological_process | O-glycan processing | 36 | 0,000000000261 |
| Hermannia gibba | GO:0055085 | biological_process | transmembrane transport | 32 | 0 |
| Hermannia gibba | GO:0050661 | molecular_function | NADP binding | 32 | 0,000000009651 |
| Hermannia gibba | GO:0006468 | biological_process | protein phosphorylation | 30 | 0,000000017874 |
| Hermannia gibba | GO:0030162 | biological_process | regulation of proteolysis | 30 | 0,000003342623 |
| Hermannia gibba | GO:0004252 | molecular_function | serine-type endopeptidase activity | 29 | 0,000000001010 |
| Hermannia gibba | GO:0006417 | biological_process | regulation of translation | 28 | 0,007131296005 |
| Hermannia gibba | GO:0019284 | biological_process | L-methionine biosynthetic process from S-adenosylmethionine | 28 | 0,02021856094 |
| Hermannia gibba | GO:0071555 | biological_process | cell wall organization | 26 | 0,000360545883 |
| Hermannia gibba | GO:0005525 | molecular_function | GTP binding | 26 | 0,000534657527 |
| Hermannia gibba | GO:0006915 | biological_process | apoptotic process | 26 | 0,005253305462 |
| Hermannia gibba | GO:0014832 | biological_process | urinary bladder smooth muscle contraction | 25 | 0,000000001294 |
| Hermannia gibba | GO:0016139 | biological_process | glycoside catabolic process | 25 | 0,000000862524 |
| Hermannia gibba | GO:0005634 | cellular_component | nucleus | 25 | 0,000870217325 |
| Hermannia gibba | GO:0006470 | biological_process | protein dephosphorylation | 25 | 0,004975916577 |
| Hermannia gibba | GO:0044245 | biological_process | polysaccharide digestion | 24 | 0 |
| Hermannia gibba | GO:0016491 | molecular_function | oxidoreductase activity | 24 | 0,000000053804 |
| Hermannia gibba | GO:0001944 | biological_process | vasculature development | 24 | 0,002916610816 |
| Hermannia gibba | GO:0004721 | molecular_function | phosphoprotein phosphatase activity | 23 | 0,000004098223 |

| | | | | | |
|---|---|---|---|---|---|
| Hermannia gibba | GO:0045599 | biological_process | negative regulation of fat cell differentiation | 23 | 0,000047537151 |
| Hermannia gibba | GO:0009058 | biological_process | biosynthetic process | 23 | 0,002516563239 |
| Hermannia gibba | GO:0016614 | molecular_function | oxidoreductase activity, acting on CH-OH group of donors | 23 | 0,005368854524 |
| Hermannia gibba | GO:0007130 | biological_process | synaptonemal complex assembly | 22 | 0,000027934990 |
| Hermannia gibba | GO:0016055 | biological_process | Wnt signaling pathway | 21 | 0 |
| Hermannia gibba | GO:0006805 | biological_process | xenobiotic metabolic process | 21 | 0,000000036679 |
| Hermannia gibba | GO:0051603 | biological_process | proteolysis involved in cellular protein catabolic process | 21 | 0,01348642188 |
| Hermannia gibba | GO:0006511 | biological_process | ubiquitin-dependent protein catabolic process | 20 | 0 |
| Hermannia gibba | GO:0090502 | biological_process | RNA phosphodiester bond hydrolysis, endonucleolytic | 20 | 0,001803700858 |
| Hermannia gibba | GO:0007601 | biological_process | visual perception | 19 | 0 |
| Hermannia gibba | GO:0000272 | biological_process | polysaccharide catabolic process | 19 | 0,000001607886 |
| Hermannia gibba | GO:0006062 | biological_process | sorbitol catabolic process | 19 | 0,00211788434 |
| Hermannia gibba | GO:0031030 | biological_process | negative regulation of septation initiation signaling | 18 | 0,000000002660 |
| Hermannia gibba | GO:0007016 | biological_process | cytoskeletal anchoring at plasma membrane | 18 | 0,000000027822 |
| Hermannia gibba | GO:0000723 | biological_process | telomere maintenance | 18 | 0,01197081139 |
| Hermannia gibba | GO:0016192 | biological_process | vesicle-mediated transport | 17 | 0 |
| Hermannia gibba | GO:0004722 | molecular_function | protein serine/threonine phosphatase activity | 17 | 0,000114938309 |
| Hermannia gibba | GO:0006310 | biological_process | DNA recombination | 17 | 0,008498174325 |
| Hermannia gibba | GO:0006728 | biological_process | pteridine biosynthetic process | 16 | 0,000000004111[illegible] |
| Hermannia gibba | GO:0006090 | biological_process | pyruvate metabolic process | 16 | 0,000008340412 |
| Hermannia gibba | GO:0006646 | biological_process | phosphatidylethanolamine biosynthetic process | 16 | 0,000554801951 |
| Hermannia gibba | GO:0021591 | biological_process | ventricular system development | 16 | 0,004473261704 |
| Hermannia gibba | GO:1902473 | biological_process | regulation of protein localization to synapse | 15 | 0,000000080015 |
| Hermannia gibba | GO:0009263 | biological_process | deoxyribonucleotide biosynthetic process | 15 | 0,000001898008 |
| Hermannia gibba | GO:0034976 | biological_process | response to endoplasmic reticulum stress | 15 | 0,000022073882 |
| Hermannia gibba | GO:0006355 | biological_process | regulation of transcription, DNA-templated | 14 | 0 |
| Hermannia gibba | GO:0010139 | biological_process | pyrimidine deoxyribonucleotide salvage | 14 | 0 |
| Hermannia gibba | GO:0043181 | biological_process | vacuolar sequestering | 14 | 0,000000001691 |
| Hermannia gibba | GO:0005983 | biological_process | starch catabolic process | 14 | 0,000000024669 |
| Hermannia gibba | GO:0006941 | biological_process | striated muscle contraction | 14 | 0,000007222679 |
| Hermannia gibba | GO:0007605 | biological_process | sensory perception of sound | 14 | 0,000012639964 |
| Hermannia gibba | GO:0034612 | biological_process | response to tumor necrosis factor | 13 | 0,000055041777 |
| Hermannia gibba | GO:0001669 | cellular_component | acrosomal vesicle | 13 | 0,000060548859 |
| Hermannia gibba | GO:0001649 | biological_process | osteoblast differentiation | 13 | 0,00007738863 |
| Hermannia gibba | GO:0005980 | biological_process | glycogen catabolic process | 13 | 0,001669567205 |
| Hermannia gibba | GO:0016114 | biological_process | terpenoid biosynthetic process | 13 | 0,004052490274 |
| Hermannia gibba | GO:0006412 | biological_process | translation | 12 | 0 |

| | | | | | |
|---|---|---|---|---|---|
| Hermannia gibba | GO:0016021 | cellular_component | integral component of membrane | 12 | 0 |
| Hermannia gibba | GO:0009617 | biological_process | response to bacterium | 12 | 0 |
| Hermannia gibba | GO:0042135 | biological_process | neurotransmitter catabolic process | 12 | 0,000044245112[illegible] |
| Hermannia gibba | GO:0046693 | biological_process | sperm storage | 12 | 0,009083172394 |
| Hermannia gibba | GO:0006644 | biological_process | phospholipid metabolic process | 12 | 0,01594191446 |
| Hermannia gibba | GO:0008380 | biological_process | RNA splicing | 11 | 0 |
| Hermannia gibba | GO:0009636 | biological_process | response to toxic substance | 11 | 0,000000003490 |
| Hermannia gibba | GO:0010869 | biological_process | regulation of receptor biosynthetic process | 11 | 0,000001650126 |
| Hermannia gibba | GO:0055114 | biological_process | oxidation-reduction process | 11 | 0,000101341794 |
| Hermannia gibba | GO:0014003 | biological_process | oligodendrocyte development | 11 | 0,002005658656 |
| Hermannia gibba | GO:0004497 | molecular_function | monooxygenase activity | 11 | 0,003440982152 |
| Hermannia gibba | GO:0070633 | biological_process | transepithelial transport | 11 | 0,01177387329 |
| Hermannia gibba | GO:0032482 | biological_process | Rab protein signal transduction | 11 | 0,01604963596 |
| Hermannia gibba | GO:0055069 | biological_process | zinc ion homeostasis | 10 | 0,000045563169 |
| Hermannia gibba | GO:0008361 | biological_process | regulation of cell size | 10 | 0,003369791404 |
| Hermannia gibba | GO:0000922 | cellular_component | spindle pole | 10 | 0,01204683045 |
| Hermannia gibba | GO:0006986 | biological_process | response to unfolded protein | 10 | 0,02508330814 |
| Hermannia gibba | GO:0016567 | biological_process | protein ubiquitination | 9 | 0 |
| Hermannia gibba | GO:0016261 | biological_process | selenocysteine catabolic process | 9 | 0,000000004327 |
| Hermannia gibba | GO:0001507 | biological_process | acetylcholine catabolic process in synaptic cleft | 9 | 0,000001800698 |
| Hermannia gibba | GO:0004674 | molecular_function | protein serine/threonine kinase activity | 9 | 0,000617160479 |
| Hermannia gibba | GO:0006811 | biological_process | ion transport | 9 | 0,001819591897 |
| Hermannia gibba | GO:0009405 | biological_process | pathogenesis | 9 | 0,00351033191 |
| Hermannia gibba | GO:0031098 | biological_process | stress-activated protein kinase signaling cascade | 8 | 0 |
| Hermannia gibba | GO:0042302 | molecular_function | structural constituent of cuticle | 8 | 0 |
| Hermannia gibba | GO:0006099 | biological_process | tricarboxylic acid cycle | 8 | 0,000000000561 |
| Hermannia gibba | GO:0006367 | biological_process | transcription initiation from RNA polymerase II promoter | 8 | 0,000000211457[illegible] |
| Hermannia gibba | GO:0005506 | molecular_function | iron ion binding | 8 | 0,000003749915[illegible] |
| Hermannia gibba | GO:0005886 | cellular_component | plasma membrane | 8 | 0,000010699092 |
| Hermannia gibba | GO:0008202 | biological_process | steroid metabolic process | 8 | 0,000229062591 |
| Hermannia gibba | GO:0031638 | biological_process | zymogen activation | 8 | 0,000380939697 |
| Hermannia gibba | GO:0008270 | molecular_function | zinc ion binding | 7 | 0 |
| Hermannia gibba | GO:0016042 | biological_process | lipid catabolic process | 7 | 0 |
| Hermannia gibba | GO:0007586 | biological_process | digestion | 7 | 0,000002899873 |
| Hermannia gibba | GO:0030435 | biological_process | sporulation resulting in formation of a cellular spore | 7 | 0,000008537178 |
| Hermannia gibba | GO:0045490 | biological_process | pectin catabolic process | 7 | 0,004860162235 |
| Hermannia gibba | GO:0006325 | biological_process | chromatin organization | 6 | 0,000000000449 |

| | | | | | |
|---|---|---|---|---|---|
| Hermannia gibba | GO:1901639 | biological_process | XDP catabolic process | 6 | 0,01121422234 |
| Hermannia gibba | GO:0006898 | biological_process | receptor-mediated endocytosis | 6 | 0,02244453776 |
| Hermannia gibba | GO:0016032 | biological_process | viral process | 5 | 0 |
| Hermannia gibba | GO:0007399 | biological_process | nervous system development | 5 | 0,000000559693 |
| Hermannia gibba | GO:0000209 | biological_process | protein polyubiquitination | 5 | 0,000000623080 |
| Hermannia gibba | GO:0050951 | biological_process | sensory perception of temperature stimulus | 5 | 0,000121867977 |
| Hermannia gibba | GO:0001505 | biological_process | regulation of neurotransmitter levels | 5 | 0,005948899409 |
| Hermannia gibba | GO:0048364 | biological_process | root development | 5 | 0,02538612223 |
| Hermannia gibba | GO:0016740 | molecular_function | transferase activity | 5 | 0,02538612223 |
| Hermannia gibba | GO:0046872 | molecular_function | metal ion binding | 4 | 0 |
| Hermannia gibba | GO:0015031 | biological_process | protein transport | 4 | 0 |
| Hermannia gibba | GO:0007179 | biological_process | transforming growth factor beta receptor signaling pathway | 4 | 0,000000139056 |
| Hermannia gibba | GO:0008542 | biological_process | visual learning | 4 | 0,000013524555 |
| Hermannia gibba | GO:0004222 | molecular_function | metalloendopeptidase activity | 4 | 0,000038345476 |
| Hermannia gibba | GO:0008080 | molecular_function | N-acetyltransferase activity | 4 | 0,000195569891 |
| Hermannia gibba | GO:0007416 | biological_process | synapse assembly | 4 | 0,000260449709 |
| Hermannia gibba | GO:0001570 | biological_process | vasculogenesis | 4 | 0,000467413662 |
| Hermannia gibba | GO:0031149 | biological_process | sorocarp stalk cell differentiation | 4 | 0,000862778126 |
| Hermannia gibba | GO:0009742 | biological_process | brassinosteroid mediated signaling pathway | 4 | 0,000862778126 |
| Hermannia gibba | GO:2000377 | biological_process | regulation of reactive oxygen species metabolic process | 4 | 0,002099467469 |
| Hermannia gibba | GO:0034405 | biological_process | response to fluid shear stress | 4 | 0,002283664436 |
| Hermannia gibba | GO:2000504 | biological_process | positive regulation of blood vessel remodeling | 4 | 0,004701167813 |
| Hermannia gibba | GO:0001666 | biological_process | response to hypoxia | 4 | 0,005566631851 |
| Hermannia gibba | GO:0009213 | biological_process | pyrimidine deoxyribonucleoside triphosphate catabolic process | 4 | 0,008295015341 |
| Hermannia gibba | GO:0070858 | biological_process | negative regulation of bile acid biosynthetic process | 4 | 0,01317212665 |
| Hermannia gibba | GO:0006890 | biological_process | retrograde vesicle-mediated transport, Golgi to ER | 3 | 0 |
| Hermannia gibba | GO:0030245 | biological_process | cellulose catabolic process | 3 | 0,000000000706 |
| Hermannia gibba | GO:0007608 | biological_process | sensory perception of smell | 3 | 0,000002491339 |
| Hermannia gibba | GO:0009408 | biological_process | response to heat | 3 | 0,000016324693 |
| Hermannia gibba | GO:0001501 | biological_process | skeletal system development | 3 | 0,000352262813 |
| Hermannia gibba | GO:0035556 | biological_process | intracellular signal transduction | 3 | 0,000965993128 |
| Hermannia gibba | GO:0046680 | biological_process | response to DDT | 3 | 0,005852287441 |
| Hermannia gibba | GO:0000011 | biological_process | vacuole inheritance | 3 | 0,01290516235 |
| Hermannia gibba | GO:0050807 | biological_process | regulation of synapse organization | 3 | 0,02218102344 |
| Hermannia gibba | GO:0006357 | biological_process | regulation of transcription from RNA polymerase II promoter | 2 | 0 |
| Hermannia gibba | GO:0006636 | biological_process | unsaturated fatty acid biosynthetic process | 2 | 0 |
| Hermannia gibba | GO:0005576 | cellular_component | extracellular region | 2 | 0 |

| | | | | | |
|---|---|---|---|---|---|
| Hermannia gibba | GO:0007169 | biological_process | transmembrane receptor protein tyrosine kinase signaling pathway | 2 | 0 |
| Hermannia gibba | GO:0006457 | biological_process | protein folding | 2 | 0 |
| Hermannia gibba | GO:0045892 | biological_process | negative regulation of transcription, DNA-templated | 2 | 0 |
| Hermannia gibba | GO:0007264 | biological_process | small GTPase mediated signal transduction | 2 | 0 |
| Hermannia gibba | GO:0010224 | biological_process | response to UV-B | 2 | 0,000000063981 |
| Hermannia gibba | GO:0035220 | biological_process | wing disc development | 2 | 0,000000081371 |
| Hermannia gibba | GO:0006641 | biological_process | triglyceride metabolic process | 2 | 0,000000616738 |
| Hermannia gibba | GO:0051301 | biological_process | cell division | 2 | 0,000004015295 |
| Hermannia gibba | GO:0008168 | molecular_function | methyltransferase activity | 2 | 0,000008031402 |
| Hermannia gibba | GO:0006383 | biological_process | transcription from RNA polymerase III promoter | 2 | 0,000014250740 |
| Hermannia gibba | GO:0007584 | biological_process | response to nutrient | 2 | 0,000017904447 |
| Hermannia gibba | GO:0045087 | biological_process | innate immune response | 2 | 0,000022479730 |
| Hermannia gibba | GO:0010468 | biological_process | regulation of gene expression | 2 | 0,000028204634 |
| Hermannia gibba | GO:0004930 | molecular_function | G-protein coupled receptor activity | 2 | 0,000055465509 |
| Hermannia gibba | GO:0042981 | biological_process | regulation of apoptotic process | 2 | 0,000209942506 |
| Hermannia gibba | GO:0030246 | molecular_function | carbohydrate binding | 2 | 0,000557539486 |
| Hermannia gibba | GO:0051290 | biological_process | protein heterotetramerization | 2 | 0,000950391307 |
| Hermannia gibba | GO:0031175 | biological_process | neuron projection development | 2 | 0,001608004681 |
| Hermannia gibba | GO:0097264 | biological_process | self proteolysis | 2 | 0,004055404534 |
| Hermannia gibba | GO:0042381 | biological_process | hemolymph coagulation | 2 | 0,006686766724 |
| Hermannia gibba | GO:0004970 | molecular_function | ionotropic glutamate receptor activity | 2 | 0,008141740367 |
| Hermannia gibba | GO:0051124 | biological_process | synaptic growth at neuromuscular junction | 2 | 0,01089835005 |
| Hermannia gibba | GO:0071622 | biological_process | regulation of granulocyte chemotaxis | 2 | 0,01403948096 |
| Hermannia gibba | GO:1902542 | biological_process | regulation of protein localization to mitotic spindle pole body | 2 | 0,01403948096 |
| Hermannia gibba | GO:0051177 | biological_process | meiotic sister chromatid cohesion | 2 | 0,01403948096 |
| Hermannia gibba | GO:0051438 | biological_process | regulation of ubiquitin-protein transferase activity | 2 | 0,01595674906 |
| Hermannia gibba | GO:0048512 | biological_process | circadian behavior | 2 | 0,0192397646 |
| Hermannia gibba | GO:1990641 | biological_process | response to iron ion starvation | 2 | 0,0192397646 |

| | | | | | |
|---|---|---|---|---|---|
| Atropacarus striculus | GO:0055085 | biological_process | transmembrane transport | 301 | 0 |
| Atropacarus striculus | GO:0032100 | biological_process | positive regulation of appetite | 218 | 0 |
| Atropacarus striculus | GO:0042908 | biological_process | xenobiotic transport | 97 | 0 |
| Atropacarus striculus | GO:0005975 | biological_process | carbohydrate metabolic process | 93 | 0,000170124334 |
| Atropacarus striculus | GO:0006805 | biological_process | xenobiotic metabolic process | 90 | 0,000000106945 |
| Atropacarus striculus | GO:0016020 | cellular_component | membrane | 81 | 0 |
| Atropacarus striculus | GO:0016301 | molecular_function | kinase activity | 78 | 0 |
| Atropacarus striculus | GO:0007601 | biological_process | visual perception | 71 | 0,000015792346 |
| Atropacarus striculus | GO:0008270 | molecular_function | zinc ion binding | 70 | 0,01152144798 |
| Atropacarus striculus | GO:0003677 | molecular_function | DNA binding | 68 | 0,000176919317 |
| Atropacarus striculus | GO:0007283 | biological_process | spermatogenesis | 67 | 0 |
| Atropacarus striculus | GO:0006706 | biological_process | steroid catabolic process | 63 | 0,000000000255 |
| Atropacarus striculus | GO:0003081 | biological_process | regulation of systemic arterial blood pressure by renin-angiotensin | 62 | 0 |
| Atropacarus striculus | GO:0004364 | molecular_function | glutathione transferase activity | 61 | 0 |
| Atropacarus striculus | GO:0042302 | molecular_function | structural constituent of cuticle | 61 | 0,001931849504 |
| Atropacarus striculus | GO:0032197 | biological_process | transposition, RNA-mediated | 59 | 0 |
| Atropacarus striculus | GO:0030223 | biological_process | neutrophil differentiation | 58 | 0 |
| Atropacarus striculus | GO:0015074 | biological_process | DNA integration | 58 | 0,000000000884 |
| Atropacarus striculus | GO:0000723 | biological_process | telomere maintenance | 50 | 0,000014084835 |
| Atropacarus striculus | GO:0009313 | biological_process | oligosaccharide catabolic process | 47 | 0 |
| Atropacarus striculus | GO:0052697 | biological_process | xenobiotic glucuronidation | 46 | 0 |
| Atropacarus striculus | GO:0009617 | biological_process | response to bacterium | 46 | 0,000035978170 |
| Atropacarus striculus | GO:0006486 | biological_process | protein glycosylation | 46 | 0,000077794393 |
| Atropacarus striculus | GO:0005524 | molecular_function | ATP binding | 45 | 0,00000311194[illegible] |
| Atropacarus striculus | GO:0006334 | biological_process | nucleosome assembly | 45 | 0,000137964686 |
| Atropacarus striculus | GO:0007275 | biological_process | multicellular organismal development | 44 | 0 |
| Atropacarus striculus | GO:0004252 | molecular_function | serine-type endopeptidase activity | 40 | 0,000002021027 |
| Atropacarus striculus | GO:0007165 | biological_process | signal transduction | 39 | 0 |
| Atropacarus striculus | GO:0010842 | biological_process | retina layer formation | 39 | 0 |
| Atropacarus striculus | GO:0010224 | biological_process | response to UV-B | 38 | 0,000097820318 |
| Atropacarus striculus | GO:0004497 | molecular_function | monooxygenase activity | 38 | 0,000275147637 |
| Atropacarus striculus | GO:0030435 | biological_process | sporulation resulting in formation of a cellular spore | 38 | 0,002475252794 |
| Atropacarus striculus | GO:0006260 | biological_process | DNA replication | 37 | 0,000317418531 |
| Atropacarus striculus | GO:0005506 | molecular_function | iron ion binding | 36 | 0,02101503132 |
| Atropacarus striculus | GO:0006814 | biological_process | sodium ion transport | 34 | 0,000013172335 |
| Atropacarus striculus | GO:0006517 | biological_process | protein deglycosylation | 34 | 0,000245580975 |
| Atropacarus striculus | GO:0016139 | biological_process | glycoside catabolic process | 33 | 0 |

| | | | | |
|---|---|---|---|---|
| Atropacarus striculus | GO:0042178 | biological_process | xenobiotic catabolic process | 32 0,000000019793 |
| Atropacarus striculus | GO:0009636 | biological_process | response to toxic substance | 32 0,01660320715 |
| Atropacarus striculus | GO:0016042 | biological_process | lipid catabolic process | 31 0,0001113911709 |
| Atropacarus striculus | GO:0006013 | biological_process | mannose metabolic process | 30 0,0000009121166 |
| Atropacarus striculus | GO:0004867 | molecular_function | serine-type endopeptidase inhibitor activity | 29 0,01596146867 |
| Atropacarus striculus | GO:0019284 | biological_process | L-methionine biosynthetic process from S-adenosylmethionine | 29 0,01985697896 |
| Atropacarus striculus | GO:0006915 | biological_process | apoptotic process | 28 0,002611159423 |
| Atropacarus striculus | GO:0009888 | biological_process | tissue development | 27 0,002027017539 |
| Atropacarus striculus | GO:0048864 | biological_process | stem cell development | 25 0 |
| Atropacarus striculus | GO:0033609 | biological_process | oxalate metabolic process | 25 0 |
| Atropacarus striculus | GO:0006468 | biological_process | protein phosphorylation | 25 0 |
| Atropacarus striculus | GO:0016266 | biological_process | O-glycan processing | 25 0,000279077317 |
| Atropacarus striculus | GO:0044245 | biological_process | polysaccharide digestion | 24 0 |
| Atropacarus striculus | GO:0006313 | biological_process | transposition, DNA-mediated | 24 0,000022552729 |
| Atropacarus striculus | GO:0071555 | biological_process | cell wall organization | 24 0,002812776526 |
| Atropacarus striculus | GO:0016192 | biological_process | vesicle-mediated transport | 23 0 |
| Atropacarus striculus | GO:0046872 | molecular_function | metal ion binding | 23 0 |
| Atropacarus striculus | GO:0015020 | molecular_function | glucuronosyltransferase activity | 23 0,000000017435 |
| Atropacarus striculus | GO:0045599 | biological_process | negative regulation of fat cell differentiation | 23 0,000088476721 |
| Atropacarus striculus | GO:0010951 | biological_process | negative regulation of endopeptidase activity | 23 0,000743469542 |
| Atropacarus striculus | GO:0030497 | biological_process | fatty acid elongation | 22 0,01339493595 |
| Atropacarus striculus | GO:0016567 | biological_process | protein ubiquitination | 21 0 |
| Atropacarus striculus | GO:0006412 | biological_process | translation | 20 0 |
| Atropacarus striculus | GO:0019216 | biological_process | regulation of lipid metabolic process | 20 0,01780272879 |
| Atropacarus striculus | GO:0008210 | biological_process | estrogen metabolic process | 19 0,000000000124 |
| Atropacarus striculus | GO:0042135 | biological_process | neurotransmitter catabolic process | 19 0,004119816951 |
| Atropacarus striculus | GO:0006069 | biological_process | ethanol oxidation | 19 0,004651752195 |
| Atropacarus striculus | GO:0051384 | biological_process | response to glucocorticoid | 18 0 |
| Atropacarus striculus | GO:0006310 | biological_process | DNA recombination | 18 0,006034650807 |
| Atropacarus striculus | GO:0008146 | molecular_function | sulfotransferase activity | 17 0,000000128955 |
| Atropacarus striculus | GO:0031098 | biological_process | stress-activated protein kinase signaling cascade | 17 0,000000141204 |
| Atropacarus striculus | GO:0042742 | biological_process | defense response to bacterium | 17 0,002132229611 |
| Atropacarus striculus | GO:0051923 | biological_process | sulfation | 17 0,00487913716 |
| Atropacarus striculus | GO:0005983 | biological_process | starch catabolic process | 16 0,000000000236 |
| Atropacarus striculus | GO:0031030 | biological_process | negative regulation of septation initiation signaling | 16 0,000000312955 |
| Atropacarus striculus | GO:0008194 | molecular_function | UDP-glycosyltransferase activity | 16 0,01826903434 |
| Atropacarus striculus | GO:0043181 | biological_process | vacuolar sequestering | 15 0,000000000160 |

| | | | | | |
|---|---|---|---|---|---|
| Atropacarus striculus | GO:0047935 | molecular_function | glucose 1-dehydrogenase (NADP+) activity | 15 | 0,000003189766 |
| Atropacarus striculus | GO:0016614 | molecular_function | oxidoreductase activity, acting on CH-OH group of donors | 15 | 0,000007262579 |
| Atropacarus striculus | GO:0007605 | biological_process | sensory perception of sound | 15 | 0,000012175513 |
| Atropacarus striculus | GO:0006090 | biological_process | pyruvate metabolic process | 15 | 0,000060522491 |
| Atropacarus striculus | GO:0006511 | biological_process | ubiquitin-dependent protein catabolic process | 14 | 0 |
| Atropacarus striculus | GO:0006355 | biological_process | regulation of transcription, DNA-templated | 14 | 0 |
| Atropacarus striculus | GO:0001507 | biological_process | acetylcholine catabolic process in synaptic cleft | 14 | 0,000142649843 |
| Atropacarus striculus | GO:0006694 | biological_process | steroid biosynthetic process | 14 | 0,005348621138 |
| Atropacarus striculus | GO:0035092 | biological_process | sperm chromatin condensation | 13 | 0,000000428308 |
| Atropacarus striculus | GO:0051603 | biological_process | proteolysis involved in cellular protein catabolic process | 13 | 0,000029227548 |
| Atropacarus striculus | GO:0001649 | biological_process | osteoblast differentiation | 13 | 0,000117451845 |
| Atropacarus striculus | GO:0005980 | biological_process | glycogen catabolic process | 13 | 0,002365807428 |
| Atropacarus striculus | GO:0016021 | cellular_component | integral component of membrane | 12 | 0 |
| Atropacarus striculus | GO:0007419 | biological_process | ventral cord development | 12 | 0,000000846436 |
| Atropacarus striculus | GO:0006941 | biological_process | striated muscle contraction | 12 | 0,000264403936 |
| Atropacarus striculus | GO:0004869 | molecular_function | cysteine-type endopeptidase inhibitor activity | 12 | 0,001338941215 |
| Atropacarus striculus | GO:0019233 | biological_process | sensory perception of pain | 12 | 0,005367563368 |
| Atropacarus striculus | GO:0006417 | biological_process | regulation of translation | 12 | 0,01802213387 |
| Atropacarus striculus | GO:0042761 | biological_process | very long-chain fatty acid biosynthetic process | 11 | 0 |
| Atropacarus striculus | GO:0006470 | biological_process | protein dephosphorylation | 11 | 0,000000008324 |
| Atropacarus striculus | GO:0047936 | molecular_function | glucose 1-dehydrogenase [NAD(P)] activity | 11 | 0,000312563068 |
| Atropacarus striculus | GO:0006644 | biological_process | phospholipid metabolic process | 11 | 0,005764283125 |
| Atropacarus striculus | GO:0004222 | molecular_function | metalloendopeptidase activity | 11 | 0,01761019077 |
| Atropacarus striculus | GO:0007286 | biological_process | spermatid development | 10 | 0,000000230073 |
| Atropacarus striculus | GO:0097066 | biological_process | response to thyroid hormone | 10 | 0,000002815502 |
| Atropacarus striculus | GO:0035249 | biological_process | synaptic transmission, glutamatergic | 10 | 0,000007956903 |
| Atropacarus striculus | GO:0055069 | biological_process | zinc ion homeostasis | 10 | 0,000064873555 |
| Atropacarus striculus | GO:0006030 | biological_process | chitin metabolic process | 10 | 0,000064873555 |
| Atropacarus striculus | GO:0042572 | biological_process | retinol metabolic process | 10 | 0,000509649887 |
| Atropacarus striculus | GO:0008361 | biological_process | regulation of cell size | 10 | 0,004455202011 |
| Atropacarus striculus | GO:0070212 | biological_process | protein poly-ADP-ribosylation | 10 | 0,01126243815 |
| Atropacarus striculus | GO:0008380 | biological_process | RNA splicing | 9 | 0 |
| Atropacarus striculus | GO:0045892 | biological_process | negative regulation of transcription, DNA-templated | 9 | 0,000000522766 |
| Atropacarus striculus | GO:0007586 | biological_process | digestion | 9 | 0,000015066414 |
| Atropacarus striculus | GO:0007224 | biological_process | smoothened signaling pathway | 9 | 0,00023197838 |
| Atropacarus striculus | GO:0008218 | biological_process | bioluminescence | 9 | 0,000356716964 |
| Atropacarus striculus | GO:0016488 | biological_process | farnesol catabolic process | 9 | 0,000835044671 |

| | | | | | |
|---|---|---|---|---|---|
| Atropacarus striculus | GO:0009408 | biological_process | response to heat | 9 | 0,007969577254 |
| Atropacarus striculus | GO:0046693 | biological_process | sperm storage | 8 | 0,000199739513 |
| Atropacarus striculus | GO:0050728 | biological_process | negative regulation of inflammatory response | 8 | 0,000616080924 |
| Atropacarus striculus | GO:0007608 | biological_process | sensory perception of smell | 8 | 0,000954145087 |
| Atropacarus striculus | GO:0008241 | molecular_function | peptidyl-dipeptidase activity | 8 | 0,001366955212 |
| Atropacarus striculus | GO:0016117 | biological_process | carotenoid biosynthetic process | 8 | 0,02038639919 |
| Atropacarus striculus | GO:0006487 | biological_process | protein N-linked glycosylation | 7 | 0,000053308550 |
| Atropacarus striculus | GO:1901639 | biological_process | XDP catabolic process | 7 | 0,003215594796 |
| Atropacarus striculus | GO:0001944 | biological_process | vasculature development | 7 | 0,008853341894 |
| Atropacarus striculus | GO:0031638 | biological_process | zymogen activation | 6 | 0,000018149926 |
| Atropacarus striculus | GO:0051725 | biological_process | protein de-ADP-ribosylation | 6 | 0,000431924533 |
| Atropacarus striculus | GO:0001501 | biological_process | skeletal system development | 6 | 0,006727671555 |
| Atropacarus striculus | GO:0001659 | biological_process | temperature homeostasis | 6 | 0,008519053081 |
| Atropacarus striculus | GO:0008234 | molecular_function | cysteine-type peptidase activity | 6 | 0,01800514553 |
| Atropacarus striculus | GO:0006357 | biological_process | regulation of transcription from RNA polymerase II promoter | 5 | 0 |
| Atropacarus striculus | GO:0042060 | biological_process | wound healing | 5 | 0 |
| Atropacarus striculus | GO:0055114 | biological_process | oxidation-reduction process | 5 | 0,000000012098 |
| Atropacarus striculus | GO:0010044 | biological_process | response to aluminum ion | 5 | 0,000028300859 |
| Atropacarus striculus | GO:0036372 | biological_process | opsin transport | 5 | 0,000028300859 |
| Atropacarus striculus | GO:0050951 | biological_process | sensory perception of temperature stimulus | 5 | 0,000149014510 |
| Atropacarus striculus | GO:0007416 | biological_process | synapse assembly | 5 | 0,000541552806 |
| Atropacarus striculus | GO:0001894 | biological_process | tissue homeostasis | 5 | 0,003414019663 |
| Atropacarus striculus | GO:0042981 | biological_process | regulation of apoptotic process | 5 | 0,006235088302 |
| Atropacarus striculus | GO:0035556 | biological_process | intracellular signal transduction | 5 | 0,006235088302 |
| Atropacarus striculus | GO:0007223 | biological_process | Wnt signaling pathway, calcium modulating pathway | 5 | 0,01300282378 |
| Atropacarus striculus | GO:0043129 | biological_process | surfactant homeostasis | 5 | 0,02298339024 |
| Atropacarus striculus | GO:0006351 | biological_process | transcription, DNA-templated | 4 | 0 |
| Atropacarus striculus | GO:0000209 | biological_process | protein polyubiquitination | 4 | 0,000000041170[illegible] |
| Atropacarus striculus | GO:0048670 | biological_process | regulation of collateral sprouting | 4 | 0,000230709577 |
| Atropacarus striculus | GO:0005412 | molecular_function | glucose:sodium symporter activity | 4 | 0,000230709577 |
| Atropacarus striculus | GO:0006267 | biological_process | pre-replicative complex assembly involved in nuclear cell cycle DNA replication | 4 | 0,000230709577 |
| Atropacarus striculus | GO:0008240 | molecular_function | tripeptidyl-peptidase activity | 4 | 0,000230709577 |
| Atropacarus striculus | GO:0045087 | biological_process | innate immune response | 4 | 0,000307213320 |
| Atropacarus striculus | GO:0005525 | molecular_function | GTP binding | 4 | 0,000509886006 |
| Atropacarus striculus | GO:0060074 | biological_process | synapse maturation | 4 | 0,00101204359 |
| Atropacarus striculus | GO:0034405 | biological_process | response to fluid shear stress | 4 | 0,002663594492 |

| | | | | | |
|---|---|---|---|---|---|
| Atropacarus striculus | GO:0042770 | biological_process | signal transduction in response to DNA damage | 4 | 0,004645626166 |
| Atropacarus striculus | GO:0016185 | biological_process | synaptic vesicle budding from presynaptic membrane | 4 | 0,005452260088 |
| Atropacarus striculus | GO:0004104 | molecular_function | cholinesterase activity | 4 | 0,006089585771 |
| Atropacarus striculus | GO:0070858 | biological_process | negative regulation of bile acid biosynthetic process | 4 | 0,01510408707 |
| Atropacarus striculus | GO:0000070 | biological_process | mitotic sister chromatid segregation | 4 | 0,02208141784 |
| Atropacarus striculus | GO:0015031 | biological_process | protein transport | 3 | 0 |
| Atropacarus striculus | GO:0005634 | cellular_component | nucleus | 3 | 0 |
| Atropacarus striculus | GO:0006457 | biological_process | protein folding | 3 | 0 |
| Atropacarus striculus | GO:0045214 | biological_process | sarcomere organization | 3 | 0,000000000187 |
| Atropacarus striculus | GO:0006413 | biological_process | translational initiation | 3 | 0,000000000311 |
| Atropacarus striculus | GO:0035220 | biological_process | wing disc development | 3 | 0,000000275179 |
| Atropacarus striculus | GO:0006641 | biological_process | triglyceride metabolic process | 3 | 0,000002063294 |
| Atropacarus striculus | GO:0003281 | biological_process | ventricular septum development | 3 | 0,000364005308 |
| Atropacarus striculus | GO:0001505 | biological_process | regulation of neurotransmitter levels | 3 | 0,000364005308 |
| Atropacarus striculus | GO:0050661 | molecular_function | NADP binding | 3 | 0,001279950853 |
| Atropacarus striculus | GO:0010758 | biological_process | regulation of macrophage chemotaxis | 3 | 0,001877226648 |
| Atropacarus striculus | GO:0072164 | biological_process | mesonephric tubule development | 3 | 0,001877226648 |
| Atropacarus striculus | GO:0031175 | biological_process | neuron projection development | 3 | 0,004260314966 |
| Atropacarus striculus | GO:0090073 | biological_process | positive regulation of protein homodimerization activity | 3 | 0,006586068283 |
| Atropacarus striculus | GO:0060675 | biological_process | ureteric bud morphogenesis | 3 | 0,006586068283 |
| Atropacarus striculus | GO:0004725 | molecular_function | protein tyrosine phosphatase activity | 3 | 0,02453121266 |
| Atropacarus striculus | GO:0044826 | biological_process | viral genome integration into host DNA | 3 | 0,02533070774 |
| Atropacarus striculus | GO:0016055 | biological_process | Wnt signaling pathway | 2 | 0 |
| Atropacarus striculus | GO:0016032 | biological_process | viral process | 2 | 0 |
| Atropacarus striculus | GO:0006979 | biological_process | response to oxidative stress | 2 | 0 |
| Atropacarus striculus | GO:0000122 | biological_process | negative regulation of transcription from RNA polymerase II promoter | 2 | 0 |
| Atropacarus striculus | GO:0042626 | molecular_function | ATPase activity, coupled to transmembrane movement of substances | 2 | 0 |
| Atropacarus striculus | GO:0007264 | biological_process | small GTPase mediated signal transduction | 2 | 0 |
| Atropacarus striculus | GO:0007399 | biological_process | nervous system development | 2 | 0,000000000490 |
| Atropacarus striculus | GO:1902600 | biological_process | hydrogen ion transmembrane transport | 2 | 0,000001224184 |
| Atropacarus striculus | GO:0051301 | biological_process | cell division | 2 | 0,000001995756 |
| Atropacarus striculus | GO:0007584 | biological_process | response to nutrient | 2 | 0,000009608365 |
| Atropacarus striculus | GO:0010468 | biological_process | regulation of gene expression | 2 | 0,000015496292 |
| Atropacarus striculus | GO:0016485 | biological_process | protein processing | 2 | 0,000017455405 |
| Atropacarus striculus | GO:0032465 | biological_process | regulation of cytokinesis | 2 | 0,000039955181 |
| Atropacarus striculus | GO:0030163 | biological_process | protein catabolic process | 2 | 0,000080623298 |
| Atropacarus striculus | GO:0008039 | biological_process | synaptic target recognition | 2 | 0,000203044024 |

| | | | | | |
|---|---|---|---|---|---|
| Atropacarus striculus | GO:0015693 | biological_process | magnesium ion transport | 2 | 0,000227647333 |
| Atropacarus striculus | GO:0030148 | biological_process | sphingolipid biosynthetic process | 2 | 0,000358781721 |
| Atropacarus striculus | GO:0032880 | biological_process | regulation of protein localization | 2 | 0,000449688957 |
| Atropacarus striculus | GO:0051290 | biological_process | protein heterotetramerization | 2 | 0,000629661548 |
| Atropacarus striculus | GO:0001516 | biological_process | prostaglandin biosynthetic process | 2 | 0,000879283881 |
| Atropacarus striculus | GO:0097264 | biological_process | self proteolysis | 2 | 0,002914447139 |
| Atropacarus striculus | GO:0071357 | biological_process | cellular response to type I interferon | 2 | 0,004946692843 |
| Atropacarus striculus | GO:0042802 | molecular_function | identical protein binding | 2 | 0,005491383929 |
| Atropacarus striculus | GO:0009611 | biological_process | response to wounding | 2 | 0,01243463902 |
| Atropacarus striculus | GO:0005154 | molecular_function | epidermal growth factor receptor binding | 2 | 0,01524596227 |
| Atropacarus striculus | GO:0009651 | biological_process | response to salt stress | 2 | 0,02238738948 |

| | | | | | |
|---|---|---|---|---|---|
| Steganacarus magnus | GO:0008218 | biological_process | bioluminescence | 69 | 0 |
| Steganacarus magnus | GO:0051384 | biological_process | response to glucocorticoid | 67 | 0,000095450331 |
| Steganacarus magnus | GO:0003964 | molecular_function | RNA-directed DNA polymerase activity | 63 | 0 |
| Steganacarus magnus | GO:0016567 | biological_process | protein ubiquitination | 61 | 0,01159708026 |
| Steganacarus magnus | GO:0007283 | biological_process | spermatogenesis | 55 | 0 |
| Steganacarus magnus | GO:0055085 | biological_process | transmembrane transport | 53 | 0 |
| Steganacarus magnus | GO:0046982 | molecular_function | protein heterodimerization activity | 49 | 0 |
| Steganacarus magnus | GO:0006334 | biological_process | nucleosome assembly | 48 | 0,000000004793 |
| Steganacarus magnus | GO:0015074 | biological_process | DNA integration | 41 | 0,000002669683 |
| Steganacarus magnus | GO:0006646 | biological_process | phosphatidylethanolamine biosynthetic process | 40 | 0,000515124800 |
| Steganacarus magnus | GO:0005506 | molecular_function | iron ion binding | 36 | 0,000477509633 |
| Steganacarus magnus | GO:0006511 | biological_process | ubiquitin-dependent protein catabolic process | 33 | 0 |
| Steganacarus magnus | GO:0045214 | biological_process | sarcomere organization | 32 | 0,005484549291 |
| Steganacarus magnus | GO:0046693 | biological_process | sperm storage | 28 | 0,001687924874 |
| Steganacarus magnus | GO:0016042 | biological_process | lipid catabolic process | 28 | 0,006753915951 |
| Steganacarus magnus | GO:0031647 | biological_process | regulation of protein stability | 27 | 0,00160578254 |
| Steganacarus magnus | GO:0006749 | biological_process | glutathione metabolic process | 27 | 0,002550674629 |
| Steganacarus magnus | GO:0005525 | molecular_function | GTP binding | 26 | 0,000013038545 |
| Steganacarus magnus | GO:0010992 | biological_process | ubiquitin homeostasis | 25 | 0 |
| Steganacarus magnus | GO:0009617 | biological_process | response to bacterium | 25 | 0,000000177442 |
| Steganacarus magnus | GO:0030162 | biological_process | regulation of proteolysis | 25 | 0,000010801868 |
| Steganacarus magnus | GO:0009263 | biological_process | deoxyribonucleotide biosynthetic process | 24 | 0 |
| Steganacarus magnus | GO:0071357 | biological_process | cellular response to type I interferon | 24 | 0,000000003842 |
| Steganacarus magnus | GO:0045616 | biological_process | regulation of keratinocyte differentiation | 24 | 0,000000096548 |
| Steganacarus magnus | GO:0001944 | biological_process | vasculature development | 23 | 0,000328075455 |
| Steganacarus magnus | GO:0010383 | biological_process | cell wall polysaccharide metabolic process | 22 | 0 |
| Steganacarus magnus | GO:0007165 | biological_process | signal transduction | 21 | 0 |
| Steganacarus magnus | GO:0006355 | biological_process | regulation of transcription, DNA-templated | 21 | 0 |
| Steganacarus magnus | GO:0003677 | molecular_function | DNA binding | 21 | 0,001689818926 |
| Steganacarus magnus | GO:0031098 | biological_process | stress-activated protein kinase signaling cascade | 21 | 0,003009918087 |
| Steganacarus magnus | GO:0030512 | biological_process | negative regulation of transforming growth factor beta receptor signaling pathway | 18 | 0,000000000200 |
| Steganacarus magnus | GO:0016021 | cellular_component | integral component of membrane | 16 | 0 |
| Steganacarus magnus | GO:0007601 | biological_process | visual perception | 16 | 0 |
| Steganacarus magnus | GO:0006728 | biological_process | pteridine biosynthetic process | 16 | 0,000000000124 |
| Steganacarus magnus | GO:0051260 | biological_process | protein homooligomerization | 16 | 0,000536470327 |
| Steganacarus magnus | GO:0007275 | biological_process | multicellular organismal development | 15 | 0 |

| | | | | | |
|---|---|---|---|---|---|
| Steganacarus magnus | GO:0004252 | molecular_function | serine-type endopeptidase activity | 15 | 0 |
| Steganacarus magnus | GO:0009168 | biological_process | purine ribonucleoside monophosphate biosynthetic process | 15 | 0,000000000166 |
| Steganacarus magnus | GO:0046685 | biological_process | response to arsenic-containing substance | 14 | 0,000000019214 |
| Steganacarus magnus | GO:0016114 | biological_process | terpenoid biosynthetic process | 14 | 0,000155157376 |
| Steganacarus magnus | GO:0051603 | biological_process | proteolysis involved in cellular protein catabolic process | 14 | 0,006416046605 |
| Steganacarus magnus | GO:0007021 | biological_process | tubulin complex assembly | 14 | 0,007340333082 |
| Steganacarus magnus | GO:0042135 | biological_process | neurotransmitter catabolic process | 14 | 0,008397442262 |
| Steganacarus magnus | GO:0006412 | biological_process | translation | 11 | 0 |
| Steganacarus magnus | GO:0006814 | biological_process | sodium ion transport | 11 | 0 |
| Steganacarus magnus | GO:0042110 | biological_process | T cell activation | 11 | 0,000026846515 |
| Steganacarus magnus | GO:0007286 | biological_process | spermatid development | 11 | 0,000311534981[illegible] |
| Steganacarus magnus | GO:0006508 | biological_process | proteolysis | 10 | 0 |
| Steganacarus magnus | GO:0006468 | biological_process | protein phosphorylation | 10 | 0 |
| Steganacarus magnus | GO:0009636 | biological_process | response to toxic substance | 10 | 0,000001032718 |
| Steganacarus magnus | GO:0042060 | biological_process | wound healing | 10 | 0,000001621687 |
| Steganacarus magnus | GO:0000209 | biological_process | protein polyubiquitination | 10 | 0,007053338438 |
| Steganacarus magnus | GO:0004674 | molecular_function | protein serine/threonine kinase activity | 10 | 0,01830768864 |
| Steganacarus magnus | GO:0008380 | biological_process | RNA splicing | 9 | 0 |
| Steganacarus magnus | GO:0044179 | biological_process | hemolysis in other organism | 9 | 0,007093494446 |
| Steganacarus magnus | GO:0006811 | biological_process | ion transport | 9 | 0,01954159721 |
| Steganacarus magnus | GO:0007586 | biological_process | digestion | 8 | 0,000608633096 |
| Steganacarus magnus | GO:0016055 | biological_process | Wnt signaling pathway | 7 | 0 |
| Steganacarus magnus | GO:0006470 | biological_process | protein dephosphorylation | 7 | 0,000000094544 |
| Steganacarus magnus | GO:0016705 | molecular_function | oxidoreductase activity, acting on paired donors, with incorporation or reduction of molecular oxygen | 7 | 0,000016837774 |
| Steganacarus magnus | GO:0055114 | biological_process | oxidation-reduction process | 7 | 0,000091951738 |
| Steganacarus magnus | GO:0071928 | biological_process | tyramine signaling pathway | 7 | 0,001830220266 |
| Steganacarus magnus | GO:0030321 | biological_process | transepithelial chloride transport | 7 | 0,007798447594 |
| Steganacarus magnus | GO:0005212 | molecular_function | structural constituent of eye lens | 7 | 0,007798447594 |
| Steganacarus magnus | GO:0046872 | molecular_function | metal ion binding | 6 | 0 |
| Steganacarus magnus | GO:0006979 | biological_process | response to oxidative stress | 6 | 0 |
| Steganacarus magnus | GO:0016491 | molecular_function | oxidoreductase activity | 6 | 0 |
| Steganacarus magnus | GO:0016614 | molecular_function | oxidoreductase activity, acting on CH-OH group of donors | 6 | 0,000000116037[illegible] |
| Steganacarus magnus | GO:0001507 | biological_process | acetylcholine catabolic process in synaptic cleft | 6 | 0,000005571337 |
| Steganacarus magnus | GO:0005524 | molecular_function | ATP binding | 6 | 0,001311034577 |

| | | | | | |
|---|---|---|---|---|---|
| Steganacarus magnus | GO:0048511 | biological_process | rhythmic process | 6 | 0,001600188888 |
| Steganacarus magnus | GO:0009405 | biological_process | pathogenesis | 6 | 0,003472782241 |
| Steganacarus magnus | GO:0006914 | biological_process | autophagy | 6 | 0,02174030608 |
| Steganacarus magnus | GO:0006805 | biological_process | xenobiotic metabolic process | 5 | 0 |
| Steganacarus magnus | GO:0045893 | biological_process | positive regulation of transcription, DNA-templated | 5 | 0,000000000633 |
| Steganacarus magnus | GO:0030435 | biological_process | sporulation resulting in formation of a cellular spore | 5 | 0,000041299937 |
| Steganacarus magnus | GO:0031638 | biological_process | zymogen activation | 5 | 0,000374155299 |
| Steganacarus magnus | GO:0006641 | biological_process | triglyceride metabolic process | 5 | 0,001914478463 |
| Steganacarus magnus | GO:0045087 | biological_process | innate immune response | 5 | 0,01389965135 |
| Steganacarus magnus | GO:0016192 | biological_process | vesicle-mediated transport | 4 | 0 |
| Steganacarus magnus | GO:0001669 | cellular_component | acrosomal vesicle | 4 | 0,000000095602 |
| Steganacarus magnus | GO:0007179 | biological_process | transforming growth factor beta receptor signaling pathway | 4 | 0,000014320780 |
| Steganacarus magnus | GO:0007166 | biological_process | cell surface receptor signaling pathway | 4 | 0,000057570224 |
| Steganacarus magnus | GO:0006629 | biological_process | lipid metabolic process | 4 | 0,000204898934 |
| Steganacarus magnus | GO:0031149 | biological_process | sorocarp stalk cell differentiation | 4 | 0,000347809518 |
| Steganacarus magnus | GO:0009742 | biological_process | brassinosteroid mediated signaling pathway | 4 | 0,000347809518 |
| Steganacarus magnus | GO:2000504 | biological_process | positive regulation of blood vessel remodeling | 4 | 0,002002773351 |
| Steganacarus magnus | GO:0075732 | biological_process | viral penetration into host nucleus | 4 | 0,002002773351 |
| Steganacarus magnus | GO:0009213 | biological_process | pyrimidine deoxyribonucleoside triphosphate catabolic process | 4 | 0,003632790779 |
| Steganacarus magnus | GO:0007584 | biological_process | response to nutrient | 4 | 0,004844792032 |
| Steganacarus magnus | GO:0001932 | biological_process | regulation of protein phosphorylation | 4 | 0,005930342287 |
| Steganacarus magnus | GO:0010468 | biological_process | regulation of gene expression | 4 | 0,00635075728 |
| Steganacarus magnus | GO:0008654 | biological_process | phospholipid biosynthetic process | 4 | 0,01475878678 |
| Steganacarus magnus | GO:0006357 | biological_process | regulation of transcription from RNA polymerase II promoter | 3 | 0 |
| Steganacarus magnus | GO:0000723 | biological_process | telomere maintenance | 3 | 0,000000179916 |
| Steganacarus magnus | GO:0005886 | cellular_component | plasma membrane | 3 | 0,000000640088 |
| Steganacarus magnus | GO:0060012 | biological_process | synaptic transmission, glycinergic | 3 | 0,000005885036 |
| Steganacarus magnus | GO:0001756 | biological_process | somitogenesis | 3 | 0,001288912673 |
| Steganacarus magnus | GO:0016485 | biological_process | protein processing | 3 | 0,002201595857 |
| Steganacarus magnus | GO:0006401 | biological_process | RNA catabolic process | 3 | 0,003450743346 |
| Steganacarus magnus | GO:0072431 | biological_process | signal transduction involved in mitotic G1 DNA damage checkpoint | 3 | 0,01226648306 |
| Steganacarus magnus | GO:0042770 | biological_process | signal transduction in response to DNA damage | 3 | 0,01336937545 |
| Steganacarus magnus | GO:0032880 | biological_process | regulation of protein localization | 3 | 0,01637182069 |
| Steganacarus magnus | GO:0009107 | biological_process | lipoate biosynthetic process | 3 | 0,01946357201 |

| | | | | | |
|---|---|---|---|---|---|
| Steganacarus magnus | GO:0016032 | biological_process | viral process | 2 | 0 |
| Steganacarus magnus | GO:0045944 | biological_process | positive regulation of transcription from RNA polymerase II promoter | 2 | 0 |
| Steganacarus magnus | GO:0006351 | biological_process | transcription, DNA-templated | 2 | 0 |
| Steganacarus magnus | GO:0006325 | biological_process | chromatin organization | 2 | 0,000000000121 |
| Steganacarus magnus | GO:0000122 | biological_process | negative regulation of transcription from RNA polymerase II promoter | 2 | 0,000000005045 |
| Steganacarus magnus | GO:0007186 | biological_process | G-protein coupled receptor signaling pathway | 2 | 0,000000028601 |
| Steganacarus magnus | GO:0034765 | biological_process | regulation of ion transmembrane transport | 2 | 0,000000046128 |
| Steganacarus magnus | GO:0006694 | biological_process | steroid biosynthetic process | 2 | 0,000000489587 |
| Steganacarus magnus | GO:0010224 | biological_process | response to UV-B | 2 | 0,000004935289 |
| Steganacarus magnus | GO:0006644 | biological_process | phospholipid metabolic process | 2 | 0,000008518746 |
| Steganacarus magnus | GO:0007217 | biological_process | tachykinin receptor signaling pathway | 2 | 0,000016029239 |
| Steganacarus magnus | GO:0006417 | biological_process | regulation of translation | 2 | 0,000019183664 |
| Steganacarus magnus | GO:0051301 | biological_process | cell division | 2 | 0,000112678479 |
| Steganacarus magnus | GO:0008168 | molecular_function | methyltransferase activity | 2 | 0,000189717224 |
| Steganacarus magnus | GO:0004930 | molecular_function | G-protein coupled receptor activity | 2 | 0,000805607834 |
| Steganacarus magnus | GO:0001659 | biological_process | temperature homeostasis | 2 | 0,001220865867 |
| Steganacarus magnus | GO:2000377 | biological_process | regulation of reactive oxygen species metabolic process | 2 | 0,00143976254 |
| Steganacarus magnus | GO:0010506 | biological_process | regulation of autophagy | 2 | 0,002349493664 |

| | | | | | |
|---|---|---|---|---|---|
| Hypochthonius rufulus | GO:0032100 | biological_process | positive regulation of appetite | 218 | 0 |
| Hypochthonius rufulus | GO:0055085 | biological_process | transmembrane transport | 207 | 0,000001794930 |
| Hypochthonius rufulus | GO:0006508 | biological_process | proteolysis | 143 | 0,000004518460 |
| Hypochthonius rufulus | GO:0004252 | molecular_function | serine-type endopeptidase activity | 136 | 0,000248196744 |
| Hypochthonius rufulus | GO:0005975 | biological_process | carbohydrate metabolic process | 130 | 0,000316774058 |
| Hypochthonius rufulus | GO:0007283 | biological_process | spermatogenesis | 127 | 0 |
| Hypochthonius rufulus | GO:0015074 | biological_process | DNA integration | 103 | 0 |
| Hypochthonius rufulus | GO:0030245 | biological_process | cellulose catabolic process | 102 | 0 |
| Hypochthonius rufulus | GO:0003677 | molecular_function | DNA binding | 100 | 0,000016236945 |
| Hypochthonius rufulus | GO:0042908 | biological_process | xenobiotic transport | 97 | 0 |
| Hypochthonius rufulus | GO:0006334 | biological_process | nucleosome assembly | 93 | 0 |
| Hypochthonius rufulus | GO:0006511 | biological_process | ubiquitin-dependent protein catabolic process | 86 | 0 |
| Hypochthonius rufulus | GO:0006517 | biological_process | protein deglycosylation | 84 | 0 |
| Hypochthonius rufulus | GO:0016567 | biological_process | protein ubiquitination | 79 | 0 |
| Hypochthonius rufulus | GO:0032197 | biological_process | transposition, RNA-mediated | 73 | 0 |
| Hypochthonius rufulus | GO:0006486 | biological_process | protein glycosylation | 69 | 0,000000757647 |
| Hypochthonius rufulus | GO:0045214 | biological_process | sarcomere organization | 67 | 0,000019764153 |
| Hypochthonius rufulus | GO:0016491 | molecular_function | oxidoreductase activity | 65 | 0,000582268964 |
| Hypochthonius rufulus | GO:0007165 | biological_process | signal transduction | 63 | 0 |
| Hypochthonius rufulus | GO:0000723 | biological_process | telomere maintenance | 63 | 0,000236557299 |
| Hypochthonius rufulus | GO:0007275 | biological_process | multicellular organismal development | 62 | 0 |
| Hypochthonius rufulus | GO:0030223 | biological_process | neutrophil differentiation | 58 | 0 |
| Hypochthonius rufulus | GO:0005506 | molecular_function | iron ion binding | 57 | 0,002996403086 |
| Hypochthonius rufulus | GO:0005524 | molecular_function | ATP binding | 56 | 0,000004839900 |
| Hypochthonius rufulus | GO:0006633 | biological_process | fatty acid biosynthetic process | 56 | 0,000536481348 |
| Hypochthonius rufulus | GO:0007586 | biological_process | digestion | 55 | 0,002607314557 |
| Hypochthonius rufulus | GO:0007601 | biological_process | visual perception | 53 | 0 |
| Hypochthonius rufulus | GO:0046982 | molecular_function | protein heterodimerization activity | 50 | 0,000000123837 |
| Hypochthonius rufulus | GO:0031098 | biological_process | stress-activated protein kinase signaling cascade | 47 | 0,000626705218 |
| Hypochthonius rufulus | GO:0006313 | biological_process | transposition, DNA-mediated | 46 | 0,000095506057 |
| Hypochthonius rufulus | GO:0006355 | biological_process | regulation of transcription, DNA-templated | 43 | 0 |
| Hypochthonius rufulus | GO:0009617 | biological_process | response to bacterium | 43 | 0 |
| Hypochthonius rufulus | GO:0051260 | biological_process | protein homooligomerization | 42 | 0,000586765833 |
| Hypochthonius rufulus | GO:0006749 | biological_process | glutathione metabolic process | 42 | 0,01263883342 |
| Hypochthonius rufulus | GO:0019542 | biological_process | propionate biosynthetic process | 41 | 0 |
| Hypochthonius rufulus | GO:0006470 | biological_process | protein dephosphorylation | 41 | 0,00124658296 |
| Hypochthonius rufulus | GO:0046693 | biological_process | sperm storage | 41 | 0,02148903943 |

| | | | | | |
|---|---|---|---|---|---|
| Hypochthonius rufulus | GO:1902358 | biological_process | sulfate transmembrane transport | 39 | 0 |
| Hypochthonius rufulus | GO:0003081 | biological_process | regulation of systemic arterial blood pressure by renin-angiotensin | 39 | 0,000000000198 |
| Hypochthonius rufulus | GO:0006468 | biological_process | protein phosphorylation | 38 | 0 |
| Hypochthonius rufulus | GO:0006412 | biological_process | translation | 37 | 0 |
| Hypochthonius rufulus | GO:0007605 | biological_process | sensory perception of sound | 37 | 0,001494281595 |
| Hypochthonius rufulus | GO:0006805 | biological_process | xenobiotic metabolic process | 35 | 0 |
| Hypochthonius rufulus | GO:0050661 | molecular_function | NADP binding | 35 | 0,000012178112[illegible] |
| Hypochthonius rufulus | GO:0016055 | biological_process | Wnt signaling pathway | 34 | 0 |
| Hypochthonius rufulus | GO:0006814 | biological_process | sodium ion transport | 34 | 0 |
| Hypochthonius rufulus | GO:0016042 | biological_process | lipid catabolic process | 34 | 0 |
| Hypochthonius rufulus | GO:0016614 | molecular_function | oxidoreductase activity, acting on CH-OH group of donors | 34 | 0,000183592208 |
| Hypochthonius rufulus | GO:0006325 | biological_process | chromatin organization | 34 | 0,000533088338 |
| Hypochthonius rufulus | GO:0008270 | molecular_function | zinc ion binding | 32 | 0 |
| Hypochthonius rufulus | GO:0016192 | biological_process | vesicle-mediated transport | 31 | 0 |
| Hypochthonius rufulus | GO:0006310 | biological_process | DNA recombination | 31 | 0,000027201688 |
| Hypochthonius rufulus | GO:0055129 | biological_process | L-proline biosynthetic process | 31 | 0,000693051574 |
| Hypochthonius rufulus | GO:0001507 | biological_process | acetylcholine catabolic process in synaptic cleft | 31 | 0,002015734097 |
| Hypochthonius rufulus | GO:0000270 | biological_process | peptidoglycan metabolic process | 30 | 0,000000003105 |
| Hypochthonius rufulus | GO:0010842 | biological_process | retina layer formation | 30 | 0,00340148547 |
| Hypochthonius rufulus | GO:0016705 | molecular_function | oxidoreductase activity, acting on paired donors, with incorporation or reduction of molecular oxygen | 29 | 0,000534978240 |
| Hypochthonius rufulus | GO:0005886 | cellular_component | plasma membrane | 29 | 0,008562578381 |
| Hypochthonius rufulus | GO:0034976 | biological_process | response to endoplasmic reticulum stress | 27 | 0,000001312763 |
| Hypochthonius rufulus | GO:0010992 | biological_process | ubiquitin homeostasis | 25 | 0 |
| Hypochthonius rufulus | GO:0031048 | biological_process | chromatin silencing by small RNA | 25 | 0,000000001794 |
| Hypochthonius rufulus | GO:0055114 | biological_process | oxidation-reduction process | 25 | 0,000401897887 |
| Hypochthonius rufulus | GO:0002376 | biological_process | immune system process | 25 | 0,005141161801 |
| Hypochthonius rufulus | GO:0044245 | biological_process | polysaccharide digestion | 24 | 0 |
| Hypochthonius rufulus | GO:0009263 | biological_process | deoxyribonucleotide biosynthetic process | 24 | 0 |
| Hypochthonius rufulus | GO:0070328 | biological_process | triglyceride homeostasis | 24 | 0,000080752819 |
| Hypochthonius rufulus | GO:0016139 | biological_process | glycoside catabolic process | 24 | 0,002909372055 |
| Hypochthonius rufulus | GO:0045616 | biological_process | regulation of keratinocyte differentiation | 24 | 0,004680091815 |
| Hypochthonius rufulus | GO:0003964 | molecular_function | RNA-directed DNA polymerase activity | 24 | 0,0241950966 |
| Hypochthonius rufulus | GO:0023052 | biological_process | signaling | 23 | 0,000955663003 |
| Hypochthonius rufulus | GO:0008202 | biological_process | steroid metabolic process | 23 | 0,007269911525 |

| | | | | | |
|---|---|---|---|---|---|
| Hypochthonius rufulus | GO:0045493 | biological_process | xylan catabolic process | 23 | 0,01767509068 |
| Hypochthonius rufulus | GO:0046872 | molecular_function | metal ion binding | 22 | 0 |
| Hypochthonius rufulus | GO:0051384 | biological_process | response to glucocorticoid | 22 | 0 |
| Hypochthonius rufulus | GO:0010383 | biological_process | cell wall polysaccharide metabolic process | 22 | 0,000000003511 |
| Hypochthonius rufulus | GO:0000209 | biological_process | protein polyubiquitination | 22 | 0,000735745030 |
| Hypochthonius rufulus | GO:0007169 | biological_process | transmembrane receptor protein tyrosine kinase signaling pathway | 21 | 0,000000025251 |
| Hypochthonius rufulus | GO:0006680 | biological_process | glucosylceramide catabolic process | 21 | 0,01599783892 |
| Hypochthonius rufulus | GO:0019882 | biological_process | antigen processing and presentation | 20 | 0,000001105477 |
| Hypochthonius rufulus | GO:0004721 | molecular_function | phosphoprotein phosphatase activity | 20 | 0,01958801509 |
| Hypochthonius rufulus | GO:0006728 | biological_process | pteridine biosynthetic process | 19 | 0,000000005201 |
| Hypochthonius rufulus | GO:0006457 | biological_process | protein folding | 19 | 0,000000096158 |
| Hypochthonius rufulus | GO:0008210 | biological_process | estrogen metabolic process | 19 | 0,000000147168 |
| Hypochthonius rufulus | GO:0007179 | biological_process | transforming growth factor beta receptor signaling pathway | 19 | 0,000106224504 |
| Hypochthonius rufulus | GO:0070633 | biological_process | transepithelial transport | 19 | 0,000282662918 |
| Hypochthonius rufulus | GO:0008380 | biological_process | RNA splicing | 18 | 0 |
| Hypochthonius rufulus | GO:0009636 | biological_process | response to toxic substance | 18 | 0 |
| Hypochthonius rufulus | GO:0042135 | biological_process | neurotransmitter catabolic process | 18 | 0,000000065043 |
| Hypochthonius rufulus | GO:0007016 | biological_process | cytoskeletal anchoring at plasma membrane | 18 | 0,000024251527 |
| Hypochthonius rufulus | GO:0060049 | biological_process | regulation of protein glycosylation | 18 | 0,000024251527 |
| Hypochthonius rufulus | GO:0004497 | molecular_function | monooxygenase activity | 18 | 0,000404432464 |
| Hypochthonius rufulus | GO:0031638 | biological_process | zymogen activation | 18 | 0,000687105426 |
| Hypochthonius rufulus | GO:0045944 | biological_process | positive regulation of transcription from RNA polymerase II promoter | 17 | 0 |
| Hypochthonius rufulus | GO:0042761 | biological_process | very long-chain fatty acid biosynthetic process | 17 | 0 |
| Hypochthonius rufulus | GO:0008146 | molecular_function | sulfotransferase activity | 17 | 0,000041450848 |
| Hypochthonius rufulus | GO:0031647 | biological_process | regulation of protein stability | 17 | 0,000868323535 |
| Hypochthonius rufulus | GO:0070189 | biological_process | kynurenine metabolic process | 17 | 0,001314438515 |
| Hypochthonius rufulus | GO:0004222 | molecular_function | metalloendopeptidase activity | 17 | 0,005482039665 |
| Hypochthonius rufulus | GO:0016021 | cellular_component | integral component of membrane | 16 | 0 |
| Hypochthonius rufulus | GO:0005983 | biological_process | starch catabolic process | 16 | 0,000000113760 |
| Hypochthonius rufulus | GO:0046373 | biological_process | L-arabinose metabolic process | 16 | 0,000070904688 |
| Hypochthonius rufulus | GO:0031030 | biological_process | negative regulation of septation initiation signaling | 16 | 0,000070904688 |
| Hypochthonius rufulus | GO:0043181 | biological_process | vacuolar sequestering | 15 | 0,000000058718 |
| Hypochthonius rufulus | GO:0008218 | biological_process | bioluminescence | 15 | 0,000033055899 |
| Hypochthonius rufulus | GO:0032434 | biological_process | regulation of proteasomal ubiquitin-dependent protein catabolic process | 15 | 0,001985386176 |

| | | | | | |
|---|---|---|---|---|---|
| Hypochthonius rufulus | GO:0006810 | biological_process | transport | 15 | 0,01155891491 |
| Hypochthonius rufulus | GO:0030497 | biological_process | fatty acid elongation | 15 | 0,02316325143 |
| Hypochthonius rufulus | GO:0051603 | biological_process | proteolysis involved in cellular protein catabolic process | 14 | 0,000000000171 |
| Hypochthonius rufulus | GO:0046685 | biological_process | response to arsenic-containing substance | 14 | 0,000080933566 |
| Hypochthonius rufulus | GO:0006941 | biological_process | striated muscle contraction | 14 | 0,000971172154 |
| Hypochthonius rufulus | GO:0006986 | biological_process | response to unfolded protein | 14 | 0,002167193694 |
| Hypochthonius rufulus | GO:0005634 | cellular_component | nucleus | 13 | 0 |
| Hypochthonius rufulus | GO:0030435 | biological_process | sporulation resulting in formation of a cellular spore | 13 | 0,000000134332 |
| Hypochthonius rufulus | GO:0010193 | biological_process | response to ozone | 13 | 0,001622608543 |
| Hypochthonius rufulus | GO:0001649 | biological_process | osteoblast differentiation | 13 | 0,005124389759 |
| Hypochthonius rufulus | GO:0031146 | biological_process | SCF-dependent proteasomal ubiquitin-dependent protein catabolic process | 13 | 0,00745616997 |
| Hypochthonius rufulus | GO:0006099 | biological_process | tricarboxylic acid cycle | 12 | 0 |
| Hypochthonius rufulus | GO:0007155 | biological_process | cell adhesion | 12 | 0,000000128930 |
| Hypochthonius rufulus | GO:0006417 | biological_process | regulation of translation | 12 | 0,000040294491 |
| Hypochthonius rufulus | GO:0045490 | biological_process | pectin catabolic process | 12 | 0,000041333913 |
| Hypochthonius rufulus | GO:0036324 | biological_process | vascular endothelial growth factor receptor-2 signaling pathway | 12 | 0,000141049151 |
| Hypochthonius rufulus | GO:0071625 | biological_process | vocalization behavior | 12 | 0,000355132595 |
| Hypochthonius rufulus | GO:0006970 | biological_process | response to osmotic stress | 12 | 0,01026990455 |
| Hypochthonius rufulus | GO:0042060 | biological_process | wound healing | 11 | 0 |
| Hypochthonius rufulus | GO:0005509 | molecular_function | calcium ion binding | 11 | 0,000000224860 |
| Hypochthonius rufulus | GO:0007608 | biological_process | sensory perception of smell | 11 | 0,000012122751 |
| Hypochthonius rufulus | GO:0039648 | biological_process | modulation by virus of host protein ubiquitination | 11 | 0,01598930877 |
| Hypochthonius rufulus | GO:0006979 | biological_process | response to oxidative stress | 10 | 0 |
| Hypochthonius rufulus | GO:0042572 | biological_process | retinol metabolic process | 10 | 0,000000036410 |
| Hypochthonius rufulus | GO:0016790 | molecular_function | thiolester hydrolase activity | 10 | 0,000000049200 |
| Hypochthonius rufulus | GO:0097066 | biological_process | response to thyroid hormone | 10 | 0,000115624322 |
| Hypochthonius rufulus | GO:0055069 | biological_process | zinc ion homeostasis | 10 | 0,001830336886 |
| Hypochthonius rufulus | GO:0010569 | biological_process | regulation of double-strand break repair via homologous recombination | 10 | 0,0127965953 |
| Hypochthonius rufulus | GO:0008360 | biological_process | regulation of cell shape | 10 | 0,02449945301 |
| Hypochthonius rufulus | GO:0005576 | cellular_component | extracellular region | 9 | 0 |
| Hypochthonius rufulus | GO:0006811 | biological_process | ion transport | 9 | 0,000000221186 |
| Hypochthonius rufulus | GO:0019062 | biological_process | virion attachment to host cell | 9 | 0,000002168891 |
| Hypochthonius rufulus | GO:0007584 | biological_process | response to nutrient | 9 | 0,000155714510 |
| Hypochthonius rufulus | GO:1902531 | biological_process | regulation of intracellular signal transduction | 9 | 0,000390211528 |

| | | | | | |
|---|---|---|---|---|---|
| Hypochthonius rufulus | GO:0001501 | biological_process | skeletal system development | 9 | 0,000759682282 |
| Hypochthonius rufulus | GO:0030162 | biological_process | regulation of proteolysis | 9 | 0,001108969157 |
| Hypochthonius rufulus | GO:0016488 | biological_process | farnesol catabolic process | 9 | 0,01235718696 |
| Hypochthonius rufulus | GO:0006357 | biological_process | regulation of transcription from RNA polymerase II promoter | 8 | 0 |
| Hypochthonius rufulus | GO:0008033 | biological_process | tRNA processing | 8 | 0 |
| Hypochthonius rufulus | GO:0051598 | biological_process | meiotic recombination checkpoint | 8 | 0,000001437414 |
| Hypochthonius rufulus | GO:0008241 | molecular_function | peptidyl-dipeptidase activity | 8 | 0,01546237111 |
| Hypochthonius rufulus | GO:0001558 | biological_process | regulation of cell growth | 8 | 0,01741087803 |
| Hypochthonius rufulus | GO:0071928 | biological_process | tyramine signaling pathway | 8 | 0,02033924745 |
| Hypochthonius rufulus | GO:0045892 | biological_process | negative regulation of transcription, DNA-templated | 7 | 0 |
| Hypochthonius rufulus | GO:0007399 | biological_process | nervous system development | 7 | 0 |
| Hypochthonius rufulus | GO:0006641 | biological_process | triglyceride metabolic process | 7 | 0,000000137982 |
| Hypochthonius rufulus | GO:0007416 | biological_process | synapse assembly | 7 | 0,000006803467 |
| Hypochthonius rufulus | GO:0030163 | biological_process | protein catabolic process | 7 | 0,000216345469 |
| Hypochthonius rufulus | GO:0035556 | biological_process | intracellular signal transduction | 7 | 0,000381135085[illegible] |
| Hypochthonius rufulus | GO:0004559 | molecular_function | alpha-mannosidase activity | 7 | 0,000503129621 |
| Hypochthonius rufulus | GO:0006898 | biological_process | receptor-mediated endocytosis | 7 | 0,000503547280 |
| Hypochthonius rufulus | GO:0004104 | molecular_function | cholinesterase activity | 7 | 0,00169107477 |
| Hypochthonius rufulus | GO:0032880 | biological_process | regulation of protein localization | 7 | 0,00169107477 |
| Hypochthonius rufulus | GO:0005997 | biological_process | xylulose metabolic process | 7 | 0,00363348243 |
| Hypochthonius rufulus | GO:0008299 | biological_process | isoprenoid biosynthetic process | 7 | 0,02036229821 |
| Hypochthonius rufulus | GO:0006351 | biological_process | transcription, DNA-templated | 6 | 0 |
| Hypochthonius rufulus | GO:0007166 | biological_process | cell surface receptor signaling pathway | 6 | 0,000000000127 |
| Hypochthonius rufulus | GO:0016485 | biological_process | protein processing | 6 | 0,000008341422 |
| Hypochthonius rufulus | GO:0001659 | biological_process | temperature homeostasis | 6 | 0,000040506021 |
| Hypochthonius rufulus | GO:0048794 | biological_process | swim bladder development | 6 | 0,000041800680 |
| Hypochthonius rufulus | GO:0042594 | biological_process | response to starvation | 6 | 0,000074996447 |
| Hypochthonius rufulus | GO:0035269 | biological_process | protein O-linked mannosylation | 6 | 0,000185328222 |
| Hypochthonius rufulus | GO:0000165 | biological_process | MAPK cascade | 6 | 0,000238313314 |
| Hypochthonius rufulus | GO:0006069 | biological_process | ethanol oxidation | 6 | 0,001044921497 |
| Hypochthonius rufulus | GO:0045599 | biological_process | negative regulation of fat cell differentiation | 6 | 0,001579635446 |
| Hypochthonius rufulus | GO:0048812 | biological_process | neuron projection morphogenesis | 6 | 0,001809479348 |
| Hypochthonius rufulus | GO:0007160 | biological_process | cell-matrix adhesion | 6 | 0,003084166697 |
| Hypochthonius rufulus | GO:0042742 | biological_process | defense response to bacterium | 6 | 0,01079259489 |
| Hypochthonius rufulus | GO:0004970 | molecular_function | ionotropic glutamate receptor activity | 6 | 0,01364656088 |
| Hypochthonius rufulus | GO:0045893 | biological_process | positive regulation of transcription, DNA-templated | 5 | 0 |

| | | | | | |
|---|---|---|---|---|---|
| Hypochthonius rufulus | GO:0006694 | biological_process | steroid biosynthetic process | 5 | 0 |
| Hypochthonius rufulus | GO:0006644 | biological_process | phospholipid metabolic process | 5 | 0,000000000428 |
| Hypochthonius rufulus | GO:0010468 | biological_process | regulation of gene expression | 5 | 0,000001564310 |
| Hypochthonius rufulus | GO:2000377 | biological_process | regulation of reactive oxygen species metabolic process | 5 | 0,000013765004 |
| Hypochthonius rufulus | GO:0042981 | biological_process | regulation of apoptotic process | 5 | 0,000031068221 |
| Hypochthonius rufulus | GO:0050951 | biological_process | sensory perception of temperature stimulus | 5 | 0,001099351985 |
| Hypochthonius rufulus | GO:0018394 | biological_process | peptidyl-lysine acetylation | 5 | 0,001099351985 |
| Hypochthonius rufulus | GO:0003993 | molecular_function | acid phosphatase activity | 5 | 0,002885492065 |
| Hypochthonius rufulus | GO:0006730 | biological_process | one-carbon metabolic process | 5 | 0,006473394475 |
| Hypochthonius rufulus | GO:0010811 | biological_process | positive regulation of cell-substrate adhesion | 5 | 0,007374881479 |
| Hypochthonius rufulus | GO:0070131 | biological_process | positive regulation of mitochondrial translation | 5 | 0,01386356786 |
| Hypochthonius rufulus | GO:0042359 | biological_process | vitamin D metabolic process | 5 | 0,01988026129 |
| Hypochthonius rufulus | GO:0016032 | biological_process | viral process | 4 | 0 |
| Hypochthonius rufulus | GO:0000122 | biological_process | negative regulation of transcription from RNA polymerase II promoter | 4 | 0 |
| Hypochthonius rufulus | GO:0051301 | biological_process | cell division | 4 | 0,000000012892 |
| Hypochthonius rufulus | GO:0023021 | biological_process | termination of signal transduction | 4 | 0,000000032318 |
| Hypochthonius rufulus | GO:0008654 | biological_process | phospholipid biosynthetic process | 4 | 0,000002828541 |
| Hypochthonius rufulus | GO:0048489 | biological_process | synaptic vesicle transport | 4 | 0,000015787585 |
| Hypochthonius rufulus | GO:0001666 | biological_process | response to hypoxia | 4 | 0,000018700461 |
| Hypochthonius rufulus | GO:0048741 | biological_process | skeletal muscle fiber development | 4 | 0,000188771472 |
| Hypochthonius rufulus | GO:0048670 | biological_process | regulation of collateral sprouting | 4 | 0,001209991561 |
| Hypochthonius rufulus | GO:0008080 | molecular_function | N-acetyltransferase activity | 4 | 0,001209991561 |
| Hypochthonius rufulus | GO:0005412 | molecular_function | glucose:sodium symporter activity | 4 | 0,001209991561 |
| Hypochthonius rufulus | GO:0008240 | molecular_function | tripeptidyl-peptidase activity | 4 | 0,001209991561 |
| Hypochthonius rufulus | GO:0018105 | biological_process | peptidyl-serine phosphorylation | 4 | 0,001432720882 |
| Hypochthonius rufulus | GO:0071357 | biological_process | cellular response to type I interferon | 4 | 0,001432720882 |
| Hypochthonius rufulus | GO:0042802 | molecular_function | identical protein binding | 4 | 0,001664512222 |
| Hypochthonius rufulus | GO:0007585 | biological_process | respiratory gaseous exchange | 4 | 0,003002355596 |
| Hypochthonius rufulus | GO:0048042 | biological_process | regulation of post-mating oviposition | 4 | 0,003002355596 |
| Hypochthonius rufulus | GO:0051124 | biological_process | synaptic growth at neuromuscular junction | 4 | 0,003002355596 |
| Hypochthonius rufulus | GO:0031149 | biological_process | sorocarp stalk cell differentiation | 4 | 0,004924827749 |
| Hypochthonius rufulus | GO:0009742 | biological_process | brassinosteroid mediated signaling pathway | 4 | 0,004924827749 |
| Hypochthonius rufulus | GO:0000055 | biological_process | ribosomal large subunit export from nucleus | 4 | 0,006122423095 |
| Hypochthonius rufulus | GO:0019226 | biological_process | transmission of nerve impulse | 4 | 0,006122423095 |
| Hypochthonius rufulus | GO:0044826 | biological_process | viral genome integration into host DNA | 4 | 0,01202610328 |
| Hypochthonius rufulus | GO:0032571 | biological_process | response to vitamin K | 4 | 0,01202610328 |

| | | | | | |
|---|---|---|---|---|---|
| Hypochthonius rufulus | GO:0016185 | biological_process | synaptic vesicle budding from presynaptic membrane | 4 | 0,02283958766 |
| Hypochthonius rufulus | GO:0075732 | biological_process | viral penetration into host nucleus | 4 | 0,02283958766 |
| Hypochthonius rufulus | GO:0060012 | biological_process | synaptic transmission, glycinergic | 3 | 0 |
| Hypochthonius rufulus | GO:0031297 | biological_process | replication fork processing | 3 | 0 |
| Hypochthonius rufulus | GO:0048511 | biological_process | rhythmic process | 3 | 0 |
| Hypochthonius rufulus | GO:0008542 | biological_process | visual learning | 3 | 0 |
| Hypochthonius rufulus | GO:0032482 | biological_process | Rab protein signal transduction | 3 | 0 |
| Hypochthonius rufulus | GO:0009408 | biological_process | response to heat | 3 | 0,000000000744 |
| Hypochthonius rufulus | GO:0018279 | biological_process | protein N-linked glycosylation via asparagine | 3 | 0,000000001328 |
| Hypochthonius rufulus | GO:0008168 | molecular_function | methyltransferase activity | 3 | 0,000000005083 |
| Hypochthonius rufulus | GO:0001756 | biological_process | somitogenesis | 3 | 0,000000013162 |
| Hypochthonius rufulus | GO:0004930 | molecular_function | G-protein coupled receptor activity | 3 | 0,000000125691 |
| Hypochthonius rufulus | GO:0003281 | biological_process | ventricular septum development | 3 | 0,000000459710 |
| Hypochthonius rufulus | GO:0010506 | biological_process | regulation of autophagy | 3 | 0,000001379193 |
| Hypochthonius rufulus | GO:0030246 | molecular_function | carbohydrate binding | 3 | 0,000005845884 |
| Hypochthonius rufulus | GO:0031175 | biological_process | neuron projection development | 3 | 0,000034219717 |
| Hypochthonius rufulus | GO:0005737 | cellular_component | cytoplasm | 3 | 0,000040726187 |
| Hypochthonius rufulus | GO:0002532 | biological_process | production of molecular mediator involved in inflammatory response | 3 | 0,000114427569 |
| Hypochthonius rufulus | GO:0000027 | biological_process | ribosomal large subunit assembly | 3 | 0,000517044201 |
| Hypochthonius rufulus | GO:0007338 | biological_process | single fertilization | 3 | 0,000609232268 |
| Hypochthonius rufulus | GO:0009611 | biological_process | response to wounding | 3 | 0,00160291035 |
| Hypochthonius rufulus | GO:0051438 | biological_process | regulation of ubiquitin-protein transferase activity | 3 | 0,00160291035 |
| Hypochthonius rufulus | GO:0004842 | molecular_function | ubiquitin-protein transferase activity | 3 | 0,002999882615 |
| Hypochthonius rufulus | GO:1901796 | biological_process | regulation of signal transduction by p53 class mediator | 3 | 0,004078609436 |
| Hypochthonius rufulus | GO:0035206 | biological_process | regulation of hemocyte proliferation | 3 | 0,004078609436 |
| Hypochthonius rufulus | GO:0004712 | molecular_function | protein serine/threonine/tyrosine kinase activity | 3 | 0,006498798117 |
| Hypochthonius rufulus | GO:0010758 | biological_process | regulation of macrophage chemotaxis | 3 | 0,006498798117 |
| Hypochthonius rufulus | GO:0072164 | biological_process | mesonephric tubule development | 3 | 0,006498798117 |
| Hypochthonius rufulus | GO:0014070 | biological_process | response to organic cyclic compound | 3 | 0,009961609779 |
| Hypochthonius rufulus | GO:0090073 | biological_process | positive regulation of protein homodimerization activity | 3 | 0,02115522622 |
| Hypochthonius rufulus | GO:0015031 | biological_process | protein transport | 2 | 0 |
| Hypochthonius rufulus | GO:0030433 | biological_process | ER-associated ubiquitin-dependent protein catabolic process | 2 | 0 |
| Hypochthonius rufulus | GO:0006636 | biological_process | unsaturated fatty acid biosynthetic process | 2 | 0 |

| | | | | | |
|---|---|---|---|---|---|
| Hypochthonius rufulus | GO:0016787 | molecular_function | hydrolase activity | 2 | 0 |
| Hypochthonius rufulus | GO:0042626 | molecular_function | ATPase activity, coupled to transmembrane movement of substances | 2 | 0 |
| Hypochthonius rufulus | GO:0035249 | biological_process | synaptic transmission, glutamatergic | 2 | 0 |
| Hypochthonius rufulus | GO:0007264 | biological_process | small GTPase mediated signal transduction | 2 | 0 |
| Hypochthonius rufulus | GO:0007186 | biological_process | G-protein coupled receptor signaling pathway | 2 | 0 |
| Hypochthonius rufulus | GO:0006413 | biological_process | translational initiation | 2 | 0 |
| Hypochthonius rufulus | GO:0034765 | biological_process | regulation of ion transmembrane transport | 2 | 0 |
| Hypochthonius rufulus | GO:0042246 | biological_process | tissue regeneration | 2 | 0 |
| Hypochthonius rufulus | GO:0015918 | biological_process | sterol transport | 2 | 0,000000001094 |
| Hypochthonius rufulus | GO:0032465 | biological_process | regulation of cytokinesis | 2 | 0,000000020942 |
| Hypochthonius rufulus | GO:0050896 | biological_process | response to stimulus | 2 | 0,000000030927 |
| Hypochthonius rufulus | GO:0032496 | biological_process | response to lipopolysaccharide | 2 | 0,000000176652 |
| Hypochthonius rufulus | GO:0008039 | biological_process | synaptic target recognition | 2 | 0,000000314331 |
| Hypochthonius rufulus | GO:0030148 | biological_process | sphingolipid biosynthetic process | 2 | 0,000000816759 |
| Hypochthonius rufulus | GO:0035725 | biological_process | sodium ion transmembrane transport | 2 | 0,000001443394 |
| Hypochthonius rufulus | GO:0051290 | biological_process | protein heterotetramerization | 2 | 0,000002106540 |
| Hypochthonius rufulus | GO:0004983 | molecular_function | neuropeptide Y receptor activity | 2 | 0,000006496113[illegible] |
| Hypochthonius rufulus | GO:0051225 | biological_process | spindle assembly | 2 | 0,000011352722 |
| Hypochthonius rufulus | GO:0090023 | biological_process | positive regulation of neutrophil chemotaxis | 2 | 0,000016439742 |
| Hypochthonius rufulus | GO:0042381 | biological_process | hemolymph coagulation | 2 | 0,000071073467 |
| Hypochthonius rufulus | GO:0009698 | biological_process | phenylpropanoid metabolic process | 2 | 0,000146106535 |
| Hypochthonius rufulus | GO:0017148 | biological_process | negative regulation of translation | 2 | 0,000249421157 |
| Hypochthonius rufulus | GO:0001967 | biological_process | suckling behavior | 2 | 0,000714995496 |
| Hypochthonius rufulus | GO:0009612 | biological_process | response to mechanical stimulus | 2 | 0,000714995496 |
| Hypochthonius rufulus | GO:0015020 | molecular_function | glucuronosyltransferase activity | 2 | 0,001010151806 |
| Hypochthonius rufulus | GO:0060071 | biological_process | Wnt signaling pathway, planar cell polarity pathway | 2 | 0,00119932819 |
| Hypochthonius rufulus | GO:0005355 | molecular_function | glucose transmembrane transporter activity | 2 | 0,002363711349 |
| Hypochthonius rufulus | GO:0033209 | biological_process | tumor necrosis factor-mediated signaling pathway | 2 | 0,002794496901 |
| Hypochthonius rufulus | GO:0006887 | biological_process | exocytosis | 2 | 0,003894657133 |
| Hypochthonius rufulus | GO:0050773 | biological_process | regulation of dendrite development | 2 | 0,005405640558 |
| Hypochthonius rufulus | GO:0019430 | biological_process | removal of superoxide radicals | 2 | 0,00746928882 |
| Hypochthonius rufulus | GO:0000278 | biological_process | mitotic cell cycle | 2 | 0,01027031658 |
| Hypochthonius rufulus | GO:0050975 | biological_process | sensory perception of touch | 2 | 0,01027031658 |
| Hypochthonius rufulus | GO:0006766 | biological_process | vitamin metabolic process | 2 | 0,02221119612 |
| Hypochthonius rufulus | GO:0030798 | molecular_function | trans-aconitate 2-methyltransferase activity | 2 | 0,02221119612 |

| | | | | |
|---|---|---|---|---|
| Oppiella subpectinata | GO:0003677 | molecular_function | DNA binding | 115 |
| Oppiella subpectinata | GO:0006517 | biological_process | protein deglycosylation | 63 |
| Oppiella subpectinata | GO:0055129 | biological_process | L-proline biosynthetic process | 44 |
| Oppiella subpectinata | GO:0016042 | biological_process | lipid catabolic process | 44 |
| Oppiella subpectinata | GO:0007601 | biological_process | visual perception | 39 |
| Oppiella subpectinata | GO:0055085 | biological_process | transmembrane transport | 38 |
| Oppiella subpectinata | GO:0009311 | biological_process | oligosaccharide metabolic process | 36 |
| Oppiella subpectinata | GO:0003964 | molecular_function | RNA-directed DNA polymerase activity | 36 |
| Oppiella subpectinata | GO:0052697 | biological_process | xenobiotic glucuronidation | 35 |
| Oppiella subpectinata | GO:0001516 | biological_process | prostaglandin biosynthetic process | 32 |
| Oppiella subpectinata | GO:0007283 | biological_process | spermatogenesis | 30 |
| Oppiella subpectinata | GO:0005975 | biological_process | carbohydrate metabolic process | 26 |
| Oppiella subpectinata | GO:0071357 | biological_process | cellular response to type I interferon | 25 |
| Oppiella subpectinata | GO:0004252 | molecular_function | serine-type endopeptidase activity | 25 |
| Oppiella subpectinata | GO:0003081 | biological_process | regulation of systemic arterial blood pressure by renin-angiotensin | 23 |
| Oppiella subpectinata | GO:0010383 | biological_process | cell wall polysaccharide metabolic process | 22 |
| Oppiella subpectinata | GO:0009268 | biological_process | response to pH | 22 |
| Oppiella subpectinata | GO:0010811 | biological_process | positive regulation of cell-substrate adhesion | 21 |
| Oppiella subpectinata | GO:0006680 | biological_process | glucosylceramide catabolic process | 21 |
| Oppiella subpectinata | GO:0008202 | biological_process | steroid metabolic process | 21 |
| Oppiella subpectinata | GO:0006633 | biological_process | fatty acid biosynthetic process | 21 |
| Oppiella subpectinata | GO:0008210 | biological_process | estrogen metabolic process | 19 |
| Oppiella subpectinata | GO:0051923 | biological_process | sulfation | 18 |
| Oppiella subpectinata | GO:0007165 | biological_process | signal transduction | 17 |
| Oppiella subpectinata | GO:0008146 | molecular_function | sulfotransferase activity | 17 |
| Oppiella subpectinata | GO:0051384 | biological_process | response to glucocorticoid | 17 |
| Oppiella subpectinata | GO:0010842 | biological_process | retina layer formation | 16 |
| Oppiella subpectinata | GO:0006810 | biological_process | transport | 15 |
| Oppiella subpectinata | GO:0010951 | biological_process | negative regulation of endopeptidase activity | 15 |
| Oppiella subpectinata | GO:0007021 | biological_process | tubulin complex assembly | 14 |
| Oppiella subpectinata | GO:0071555 | biological_process | cell wall organization | 14 |
| Oppiella subpectinata | GO:0006511 | biological_process | ubiquitin-dependent protein catabolic process | 13 |
| Oppiella subpectinata | GO:0001649 | biological_process | osteoblast differentiation | 13 |
| Oppiella subpectinata | GO:0006355 | biological_process | regulation of transcription, DNA-templated | 11 |
| Oppiella subpectinata | GO:0009617 | biological_process | response to bacterium | 11 |
| Oppiella subpectinata | GO:0014003 | biological_process | oligodendrocyte development | 11 |
| Oppiella subpectinata | GO:0015020 | molecular_function | glucuronosyltransferase activity | 11 |

| | | | | |
|---|---|---|---|---|
| Oppiella subpectinata | GO:0008194 | molecular_function | UDP-glycosyltransferase activity | 11 |
| Oppiella subpectinata | GO:0072089 | biological_process | stem cell proliferation | 10 |
| Oppiella subpectinata | GO:0016488 | biological_process | farnesol catabolic process | 10 |
| Oppiella subpectinata | GO:0014883 | biological_process | transition between fast and slow fiber | 10 |
| Oppiella subpectinata | GO:0007417 | biological_process | central nervous system development | 10 |
| Oppiella subpectinata | GO:0055114 | biological_process | oxidation-reduction process | 9 |
| Oppiella subpectinata | GO:0005506 | molecular_function | iron ion binding | 9 |
| Oppiella subpectinata | GO:0046872 | molecular_function | metal ion binding | 8 |
| Oppiella subpectinata | GO:0006979 | biological_process | response to oxidative stress | 8 |
| Oppiella subpectinata | GO:0051260 | biological_process | protein homooligomerization | 8 |
| Oppiella subpectinata | GO:0016021 | cellular_component | integral component of membrane | 7 |
| Oppiella subpectinata | GO:1901639 | biological_process | XDP catabolic process | 7 |
| Oppiella subpectinata | GO:0016117 | biological_process | carotenoid biosynthetic process | 7 |
| Oppiella subpectinata | GO:0044179 | biological_process | hemolysis in other organism | 7 |
| Oppiella subpectinata | GO:0051770 | biological_process | positive regulation of nitric-oxide synthase biosynthetic process | 7 |
| Oppiella subpectinata | GO:0048794 | biological_process | swim bladder development | 6 |
| Oppiella subpectinata | GO:0000103 | biological_process | sulfate assimilation | 6 |
| Oppiella subpectinata | GO:0007155 | biological_process | cell adhesion | 6 |
| Oppiella subpectinata | GO:0006508 | biological_process | proteolysis | 5 |
| Oppiella subpectinata | GO:0018394 | biological_process | peptidyl-lysine acetylation | 5 |
| Oppiella subpectinata | GO:0006468 | biological_process | protein phosphorylation | 4 |
| Oppiella subpectinata | GO:0031098 | biological_process | stress-activated protein kinase signaling cascade | 4 |
| Oppiella subpectinata | GO:0001669 | cellular_component | acrosomal vesicle | 4 |
| Oppiella subpectinata | GO:0051958 | biological_process | methotrexate transport | 4 |
| Oppiella subpectinata | GO:0001932 | biological_process | regulation of protein phosphorylation | 4 |
| Oppiella subpectinata | GO:0097479 | biological_process | synaptic vesicle localization | 4 |
| Oppiella subpectinata | GO:0090129 | biological_process | positive regulation of synapse maturation | 4 |
| Oppiella subpectinata | GO:0016055 | biological_process | Wnt signaling pathway | 3 |
| Oppiella subpectinata | GO:0016567 | biological_process | protein ubiquitination | 3 |
| Oppiella subpectinata | GO:0045944 | biological_process | positive regulation of transcription from RNA polymerase II promoter | 3 |
| Oppiella subpectinata | GO:0009636 | biological_process | response to toxic substance | 3 |
| Oppiella subpectinata | GO:0006325 | biological_process | chromatin organization | 3 |
| Oppiella subpectinata | GO:0060012 | biological_process | synaptic transmission, glycinergic | 3 |
| Oppiella subpectinata | GO:0060074 | biological_process | synapse maturation | 3 |
| Oppiella subpectinata | GO:1901261 | biological_process | regulation of sorocarp spore cell differentiation | 3 |
| Oppiella subpectinata | GO:0046547 | molecular_function | trans-aconitate 3-methyltransferase activity | 3 |
| Oppiella subpectinata | GO:0001756 | biological_process | somitogenesis | 3 |

| | | | | |
|---|---|---|---|---|
| Oppiella subpectinata | GO:0007275 | biological_process | multicellular organismal development | 2 |
| Oppiella subpectinata | GO:0008270 | molecular_function | zinc ion binding | 2 |
| Oppiella subpectinata | GO:0005576 | cellular_component | extracellular region | 2 |
| Oppiella subpectinata | GO:0000122 | biological_process | negative regulation of transcription from RNA polymerase II promoter | 2 |
| Oppiella subpectinata | GO:0007179 | biological_process | transforming growth factor beta receptor signaling pathway | 2 |
| Oppiella subpectinata | GO:0004674 | molecular_function | protein serine/threonine kinase activity | 2 |
| Oppiella subpectinata | GO:0008168 | molecular_function | methyltransferase activity | 2 |
| Oppiella subpectinata | GO:0097090 | biological_process | presynaptic membrane organization | 2 |
| Oppiella subpectinata | GO:0010468 | biological_process | regulation of gene expression | 2 |
| Oppiella subpectinata | GO:0030497 | biological_process | fatty acid elongation | 2 |

0
0
0
0,001766327324
0,000520700552
0
0
0,000000000433
0
0
0
0,00531982329
0
0,001448573005
0,000000000900
0
0,000000023306
0,000000000778
0,001704936031
0,007030024478
0,02010904778
0
0,000001757429
0
0
0,00036119525
0,00126883483
0,000000260749
0,001663711767
0,000572624277
0,01395180599
0
0,000000228087
0
0,000000001001
0,000022002219
0,000747958989

| |
|---|
| 0,01410486032 |
| 0,000000000134 |
| 0,000001079798 |
| 0,000002682178 |
| 0,002913366102 |
| 0,01857355084 |
| 0,0192184964 |
| 0 |
| 0 |
| 0,000077076958 |
| 0 |
| 0,000109686288 |
| 0,003608404752 |
| 0,01349652096 |
| 0,02086727952 |
| 0,000000109674 |
| 0,007313756469 |
| 0,007723021684 |
| 0 |
| 0,000008937837 |
| 0 |
| 0,000000035566 |
| 0,000029699966 |
| 0,000302249019 |
| 0,002049596295 |
| 0,01446273375 |
| 0,01795268742 |
| 0 |
| 0 |
| 0 |
| 0,000000026027[illegible] |
| 0,000001118584[illegible] |
| 0,000387157502 |
| 0,002915994339 |
| 0,008844045842 |
| 0,01317214146 |
| 0,0141791264 |

| |
|---|
| 0 |
| 0 |
| 0,000000158508 |
| 0,000002089975 |
| 0,000043318237 |
| 0,000140091952 |
| 0,003314714254 |
| 0,004854713287 |
| 0,006294895423 |
| 0,007059287716 |

| | | | | | |
|---|---|---|---|---|---|
| Oppiella nova | GO:0032100 | biological_process | positive regulation of appetite | 218 | 0 |
| Oppiella nova | GO:0007283 | biological_process | spermatogenesis | 96 | 0,002647423709 |
| Oppiella nova | GO:0016020 | cellular_component | membrane | 74 | 0 |
| Oppiella nova | GO:0006508 | biological_process | proteolysis | 71 | 0,005286769856 |
| Oppiella nova | GO:0032197 | biological_process | transposition, RNA-mediated | 63 | 0 |
| Oppiella nova | GO:0004364 | molecular_function | glutathione transferase activity | 63 | 0 |
| Oppiella nova | GO:0010025 | biological_process | wax biosynthetic process | 59 | 0 |
| Oppiella nova | GO:0030223 | biological_process | neutrophil differentiation | 58 | 0 |
| Oppiella nova | GO:0006633 | biological_process | fatty acid biosynthetic process | 56 | 0,000210432105 |
| Oppiella nova | GO:0015074 | biological_process | DNA integration | 55 | 0 |
| Oppiella nova | GO:0000723 | biological_process | telomere maintenance | 45 | 0,000000052599 |
| Oppiella nova | GO:0003964 | molecular_function | RNA-directed DNA polymerase activity | 40 | 0,000000001364 |
| Oppiella nova | GO:0009617 | biological_process | response to bacterium | 38 | 0,006064474142 |
| Oppiella nova | GO:0005506 | molecular_function | iron ion binding | 37 | 0,000055318516 |
| Oppiella nova | GO:0016192 | biological_process | vesicle-mediated transport | 35 | 0,000000000671 |
| Oppiella nova | GO:0006814 | biological_process | sodium ion transport | 34 | 0,0235797923 |
| Oppiella nova | GO:0006260 | biological_process | DNA replication | 33 | 0,000008270606 |
| Oppiella nova | GO:0042178 | biological_process | xenobiotic catabolic process | 32 | 0 |
| Oppiella nova | GO:0016491 | molecular_function | oxidoreductase activity | 32 | 0,01996888133 |
| Oppiella nova | GO:0045214 | biological_process | sarcomere organization | 31 | 0,00353031113 |
| Oppiella nova | GO:0004222 | molecular_function | metalloendopeptidase activity | 30 | 0,000004395116[illegible] |
| Oppiella nova | GO:0006749 | biological_process | glutathione metabolic process | 30 | 0,000089124722 |
| Oppiella nova | GO:0046693 | biological_process | sperm storage | 30 | 0,000120801523 |
| Oppiella nova | GO:0004252 | molecular_function | serine-type endopeptidase activity | 29 | 0,000113666714 |
| Oppiella nova | GO:0009888 | biological_process | tissue development | 28 | 0,000002736874 |
| Oppiella nova | GO:0019284 | biological_process | L-methionine biosynthetic process from S-adenosylmethionine | 28 | 0,000489882482 |
| Oppiella nova | GO:0048864 | biological_process | stem cell development | 25 | 0 |
| Oppiella nova | GO:0044245 | biological_process | polysaccharide digestion | 24 | 0 |
| Oppiella nova | GO:0045616 | biological_process | regulation of keratinocyte differentiation | 24 | 0,000000022661 |
| Oppiella nova | GO:0004867 | molecular_function | serine-type endopeptidase inhibitor activity | 24 | 0,005405845243 |
| Oppiella nova | GO:0005524 | molecular_function | ATP binding | 24 | 0,006910656388 |
| Oppiella nova | GO:0000272 | biological_process | polysaccharide catabolic process | 24 | 0,02029622129 |
| Oppiella nova | GO:0006355 | biological_process | regulation of transcription, DNA-templated | 23 | 0 |
| Oppiella nova | GO:0045599 | biological_process | negative regulation of fat cell differentiation | 23 | 0,000000276520 |
| Oppiella nova | GO:0006511 | biological_process | ubiquitin-dependent protein catabolic process | 22 | 0 |
| Oppiella nova | GO:0006811 | biological_process | ion transport | 21 | 0,0230741797 |
| Oppiella nova | GO:0006468 | biological_process | protein phosphorylation | 20 | 0,000000230288 |

| | | | | | |
|---|---|---|---|---|---|
| Oppiella nova | GO:0023052 | biological_process | signaling | 20 | 0,000000370681 |
| Oppiella nova | GO:0090502 | biological_process | RNA phosphodiester bond hydrolysis, endonucleolytic | 20 | 0,000037180203 |
| Oppiella nova | GO:0006805 | biological_process | xenobiotic metabolic process | 20 | 0,000156526301 |
| Oppiella nova | GO:0006310 | biological_process | DNA recombination | 19 | 0,000044210436 |
| Oppiella nova | GO:0006069 | biological_process | ethanol oxidation | 19 | 0,000085073602 |
| Oppiella nova | GO:0052697 | biological_process | xenobiotic glucuronidation | 19 | 0,000731939922 |
| Oppiella nova | GO:0010842 | biological_process | retina layer formation | 19 | 0,000819954781 |
| Oppiella nova | GO:0006470 | biological_process | protein dephosphorylation | 19 | 0,01704193034 |
| Oppiella nova | GO:0060049 | biological_process | regulation of protein glycosylation | 18 | 0,000000000185 |
| Oppiella nova | GO:0046872 | molecular_function | metal ion binding | 17 | 0 |
| Oppiella nova | GO:0008270 | molecular_function | zinc ion binding | 17 | 0 |
| Oppiella nova | GO:0006486 | biological_process | protein glycosylation | 16 | 0,000000000422 |
| Oppiella nova | GO:0015020 | molecular_function | glucuronosyltransferase activity | 16 | 0,000005995768 |
| Oppiella nova | GO:0007601 | biological_process | visual perception | 15 | 0 |
| Oppiella nova | GO:0006979 | biological_process | response to oxidative stress | 15 | 0,000000000374 |
| Oppiella nova | GO:1902477 | biological_process | regulation of defense response to bacterium, incompatible interaction | 14 | 0 |
| Oppiella nova | GO:0007016 | biological_process | cytoskeletal anchoring at plasma membrane | 14 | 0,000000913231 |
| Oppiella nova | GO:0005997 | biological_process | xylulose metabolic process | 14 | 0,008081211439 |
| Oppiella nova | GO:0042770 | biological_process | signal transduction in response to DNA damage | 14 | 0,01659218209 |
| Oppiella nova | GO:0042135 | biological_process | neurotransmitter catabolic process | 14 | 0,01790613328 |
| Oppiella nova | GO:0007165 | biological_process | signal transduction | 13 | 0 |
| Oppiella nova | GO:0007275 | biological_process | multicellular organismal development | 13 | 0 |
| Oppiella nova | GO:0033089 | biological_process | positive regulation of T cell differentiation in thymus | 13 | 0,000000000214 |
| Oppiella nova | GO:0035092 | biological_process | sperm chromatin condensation | 13 | 0,000000006365 |
| Oppiella nova | GO:0036324 | biological_process | vascular endothelial growth factor receptor-2 signaling pathway | 12 | 0,000000042600 |
| Oppiella nova | GO:0005983 | biological_process | starch catabolic process | 12 | 0,000000081550 |
| Oppiella nova | GO:0090023 | biological_process | positive regulation of neutrophil chemotaxis | 12 | 0,01821869405 |
| Oppiella nova | GO:0006412 | biological_process | translation | 11 | 0 |
| Oppiella nova | GO:0007605 | biological_process | sensory perception of sound | 11 | 0,000376901464 |
| Oppiella nova | GO:0003309 | biological_process | type B pancreatic cell differentiation | 11 | 0,002988842009 |
| Oppiella nova | GO:0008380 | biological_process | RNA splicing | 10 | 0 |
| Oppiella nova | GO:0016790 | molecular_function | thiolester hydrolase activity | 10 | 0 |
| Oppiella nova | GO:0031222 | biological_process | arabinan catabolic process | 10 | 0,000442832946 |
| Oppiella nova | GO:0000922 | cellular_component | spindle pole | 10 | 0,001583584344 |
| Oppiella nova | GO:0005886 | cellular_component | plasma membrane | 10 | 0,007730187726 |
| Oppiella nova | GO:0008237 | molecular_function | metallopeptidase activity | 10 | 0,01610162429 |
| Oppiella nova | GO:0003677 | molecular_function | DNA binding | 9 | 0,000000128857 |

| | | | | | |
|---|---|---|---|---|---|
| Oppiella nova | GO:1902531 | biological_process | regulation of intracellular signal transduction | 9 | 0,000000768063 |
| Oppiella nova | GO:0097066 | biological_process | response to thyroid hormone | 9 | 0,000001606193 |
| Oppiella nova | GO:0047936 | molecular_function | glucose 1-dehydrogenase [NAD(P)] activity | 9 | 0,000441543760 |
| Oppiella nova | GO:0009263 | biological_process | deoxyribonucleotide biosynthetic process | 9 | 0,001479504678 |
| Oppiella nova | GO:0051598 | biological_process | meiotic recombination checkpoint | 8 | 0,000000003069 |
| Oppiella nova | GO:0042302 | molecular_function | structural constituent of cuticle | 8 | 0,000000082516 |
| Oppiella nova | GO:0042060 | biological_process | wound healing | 8 | 0,00000078011[illegible] |
| Oppiella nova | GO:0006099 | biological_process | tricarboxylic acid cycle | 8 | 0,000003313782 |
| Oppiella nova | GO:0001669 | cellular_component | acrosomal vesicle | 8 | 0,000129403247 |
| Oppiella nova | GO:0021762 | biological_process | substantia nigra development | 8 | 0,00159996173 |
| Oppiella nova | GO:0010193 | biological_process | response to ozone | 8 | 0,003104067215 |
| Oppiella nova | GO:0047935 | molecular_function | glucose 1-dehydrogenase (NADP+) activity | 8 | 0,005459625168 |
| Oppiella nova | GO:0007586 | biological_process | digestion | 7 | 0,000637017732 |
| Oppiella nova | GO:0006954 | biological_process | inflammatory response | 7 | 0,001600882149 |
| Oppiella nova | GO:0007179 | biological_process | transforming growth factor beta receptor signaling pathway | 7 | 0,00167311511 |
| Oppiella nova | GO:0031297 | biological_process | replication fork processing | 7 | 0,007530025925 |
| Oppiella nova | GO:0005576 | cellular_component | extracellular region | 6 | 0,000001902745 |
| Oppiella nova | GO:0015939 | biological_process | pantothenate metabolic process | 6 | 0,000061535844 |
| Oppiella nova | GO:0042572 | biological_process | retinol metabolic process | 6 | 0,001297467888 |
| Oppiella nova | GO:0031638 | biological_process | zymogen activation | 6 | 0,002687104557 |
| Oppiella nova | GO:0033384 | biological_process | geranyl diphosphate biosynthetic process | 6 | 0,02522687389 |
| Oppiella nova | GO:0055085 | biological_process | transmembrane transport | 5 | 0 |
| Oppiella nova | GO:0015031 | biological_process | protein transport | 5 | 0 |
| Oppiella nova | GO:0001507 | biological_process | acetylcholine catabolic process in synaptic cleft | 5 | 0,000005666864 |
| Oppiella nova | GO:0002175 | biological_process | protein localization to paranode region of axon | 5 | 0,003059553339 |
| Oppiella nova | GO:0030449 | biological_process | regulation of complement activation | 5 | 0,003059553339 |
| Oppiella nova | GO:0016021 | cellular_component | integral component of membrane | 4 | 0 |
| Oppiella nova | GO:0016567 | biological_process | protein ubiquitination | 4 | 0 |
| Oppiella nova | GO:0005412 | molecular_function | glucose:sodium symporter activity | 4 | 0,000056650543 |
| Oppiella nova | GO:0007155 | biological_process | cell adhesion | 4 | 0,000060139734 |
| Oppiella nova | GO:0008202 | biological_process | steroid metabolic process | 4 | 0,000181489751 |
| Oppiella nova | GO:0004497 | molecular_function | monooxygenase activity | 4 | 0,000195132799 |
| Oppiella nova | GO:0009213 | biological_process | pyrimidine deoxyribonucleoside triphosphate catabolic process | 4 | 0,002765182204 |
| Oppiella nova | GO:0046982 | molecular_function | protein heterodimerization activity | 4 | 0,004816293558 |
| Oppiella nova | GO:0000070 | biological_process | mitotic sister chromatid segregation | 4 | 0,00692613884 |
| Oppiella nova | GO:0016032 | biological_process | viral process | 3 | 0 |
| Oppiella nova | GO:0016042 | biological_process | lipid catabolic process | 3 | 0 |

| | | | | | |
|---|---|---|---|---|---|
| Oppiella nova | GO:0005634 | cellular_component | nucleus | 3 | 0 |
| Oppiella nova | GO:0006325 | biological_process | chromatin organization | 3 | 0,000000009061 |
| Oppiella nova | GO:0016705 | molecular_function | oxidoreductase activity, acting on paired donors, with incorporation or reduction of molecular oxygen | 3 | 0,000000142591 |
| Oppiella nova | GO:0006694 | biological_process | steroid biosynthetic process | 3 | 0,000012384869 |
| Oppiella nova | GO:0010224 | biological_process | response to UV-B | 3 | 0,000088620685 |
| Oppiella nova | GO:0010758 | biological_process | regulation of macrophage chemotaxis | 3 | 0,000655699904 |
| Oppiella nova | GO:0050661 | molecular_function | NADP binding | 3 | 0,02002895803 |
| Oppiella nova | GO:0030246 | molecular_function | carbohydrate binding | 3 | 0,02260076259 |
| Oppiella nova | GO:0030433 | biological_process | ER-associated ubiquitin-dependent protein catabolic process | 2 | 0 |
| Oppiella nova | GO:0006890 | biological_process | retrograde vesicle-mediated transport, Golgi to ER | 2 | 0,000000001027 |
| Oppiella nova | GO:0016787 | molecular_function | hydrolase activity | 2 | 0,000000040949 |
| Oppiella nova | GO:0035249 | biological_process | synaptic transmission, glutamatergic | 2 | 0,000000069627 |
| Oppiella nova | GO:0030245 | biological_process | cellulose catabolic process | 2 | 0,000000128959 |
| Oppiella nova | GO:0007399 | biological_process | nervous system development | 2 | 0,000001133530 |
| Oppiella nova | GO:0048511 | biological_process | rhythmic process | 2 | 0,000017049645 |
| Oppiella nova | GO:0009405 | biological_process | pathogenesis | 2 | 0,000045991716 |
| Oppiella nova | GO:0007608 | biological_process | sensory perception of smell | 2 | 0,000049923092 |
| Oppiella nova | GO:0051301 | biological_process | cell division | 2 | 0,000272141052 |
| Oppiella nova | GO:0007584 | biological_process | response to nutrient | 2 | 0,000755353155 |
| Oppiella nova | GO:0016485 | biological_process | protein processing | 2 | 0,001110932563 |
| Oppiella nova | GO:0032465 | biological_process | regulation of cytokinesis | 2 | 0,001892887388 |
| Oppiella nova | GO:0001659 | biological_process | temperature homeostasis | 2 | 0,002372001906 |
| Oppiella nova | GO:0019216 | biological_process | regulation of lipid metabolic process | 2 | 0,002556287429 |
| Oppiella nova | GO:0030163 | biological_process | protein catabolic process | 2 | 0,002967137349 |
| Oppiella nova | GO:0042981 | biological_process | regulation of apoptotic process | 2 | 0,003987560229 |
| Oppiella nova | GO:0005154 | molecular_function | epidermal growth factor receptor binding | 2 | 0,007568366471 |
| Oppiella nova | GO:0002376 | biological_process | immune system process | 2 | 0,01341859223 |
| Oppiella nova | GO:0010182 | biological_process | sugar mediated signaling pathway | 2 | 0,0207379997 |
| Oppiella nova | GO:0010977 | biological_process | negative regulation of neuron projection development | 2 | 0,0207379997 |
| Oppiella nova | GO:0072091 | biological_process | regulation of stem cell proliferation | 2 | 0,0207379997 |

| | | | |
|---|---|---|---|
| Hydrozetes thienemanni | GO:0005975 | biological_process | carbohydrate metabolic process |
| Hydrozetes thienemanni | GO:0009313 | biological_process | oligosaccharide catabolic process |
| Hydrozetes thienemanni | GO:0016042 | biological_process | lipid catabolic process |
| Hydrozetes thienemanni | GO:0003964 | molecular_function | RNA-directed DNA polymerase activity |
| Hydrozetes thienemanni | GO:0055085 | biological_process | transmembrane transport |
| Hydrozetes thienemanni | GO:0006680 | biological_process | glucosylceramide catabolic process |
| Hydrozetes thienemanni | GO:0006313 | biological_process | transposition, DNA-mediated |
| Hydrozetes thienemanni | GO:0031647 | biological_process | regulation of protein stability |
| Hydrozetes thienemanni | GO:0030162 | biological_process | regulation of proteolysis |
| Hydrozetes thienemanni | GO:0048864 | biological_process | stem cell development |
| Hydrozetes thienemanni | GO:0001944 | biological_process | vasculature development |
| Hydrozetes thienemanni | GO:1901998 | biological_process | toxin transport |
| Hydrozetes thienemanni | GO:0030245 | biological_process | cellulose catabolic process |
| Hydrozetes thienemanni | GO:0023052 | biological_process | signaling |
| Hydrozetes thienemanni | GO:0030512 | biological_process | negative regulation of transforming growth factor beta receptor signaling pathway |
| Hydrozetes thienemanni | GO:0006805 | biological_process | xenobiotic metabolic process |
| Hydrozetes thienemanni | GO:0007021 | biological_process | tubulin complex assembly |
| Hydrozetes thienemanni | GO:0045087 | biological_process | innate immune response |
| Hydrozetes thienemanni | GO:0006511 | biological_process | ubiquitin-dependent protein catabolic process |
| Hydrozetes thienemanni | GO:0046685 | biological_process | response to arsenic-containing substance |
| Hydrozetes thienemanni | GO:0000272 | biological_process | polysaccharide catabolic process |
| Hydrozetes thienemanni | GO:0006412 | biological_process | translation |
| Hydrozetes thienemanni | GO:0010383 | biological_process | cell wall polysaccharide metabolic process |
| Hydrozetes thienemanni | GO:0070633 | biological_process | transepithelial transport |
| Hydrozetes thienemanni | GO:0046355 | biological_process | mannan catabolic process |
| Hydrozetes thienemanni | GO:0016567 | biological_process | protein ubiquitination |
| Hydrozetes thienemanni | GO:0000922 | cellular_component | spindle pole |
| Hydrozetes thienemanni | GO:0005634 | cellular_component | nucleus |
| Hydrozetes thienemanni | GO:0090023 | biological_process | positive regulation of neutrophil chemotaxis |
| Hydrozetes thienemanni | GO:0010718 | biological_process | positive regulation of epithelial to mesenchymal transition |
| Hydrozetes thienemanni | GO:0010869 | biological_process | regulation of receptor biosynthetic process |
| Hydrozetes thienemanni | GO:0042060 | biological_process | wound healing |
| Hydrozetes thienemanni | GO:0007601 | biological_process | visual perception |
| Hydrozetes thienemanni | GO:0006486 | biological_process | protein glycosylation |
| Hydrozetes thienemanni | GO:0003677 | molecular_function | DNA binding |
| Hydrozetes thienemanni | GO:0008138 | molecular_function | protein tyrosine/serine/threonine phosphatase activity |
| Hydrozetes thienemanni | GO:0006357 | biological_process | regulation of transcription from RNA polymerase II promoter |

| | | | |
|---|---|---|---|
| Hydrozetes thienemanni | GO:0006468 | biological_process | protein phosphorylation |
| Hydrozetes thienemanni | GO:0010193 | biological_process | response to ozone |
| Hydrozetes thienemanni | GO:0046470 | biological_process | phosphatidylcholine metabolic process |
| Hydrozetes thienemanni | GO:0043066 | biological_process | negative regulation of apoptotic process |
| Hydrozetes thienemanni | GO:0007165 | biological_process | signal transduction |
| Hydrozetes thienemanni | GO:0007283 | biological_process | spermatogenesis |
| Hydrozetes thienemanni | GO:0004252 | molecular_function | serine-type endopeptidase activity |
| Hydrozetes thienemanni | GO:0015893 | biological_process | drug transport |
| Hydrozetes thienemanni | GO:0036323 | biological_process | vascular endothelial growth factor receptor-1 signaling pathway |
| Hydrozetes thienemanni | GO:0016055 | biological_process | Wnt signaling pathway |
| Hydrozetes thienemanni | GO:0045944 | biological_process | positive regulation of transcription from RNA polymerase II promoter |
| Hydrozetes thienemanni | GO:0051260 | biological_process | protein homooligomerization |
| Hydrozetes thienemanni | GO:0031149 | biological_process | sorocarp stalk cell differentiation |
| Hydrozetes thienemanni | GO:0009742 | biological_process | brassinosteroid mediated signaling pathway |
| Hydrozetes thienemanni | GO:0001932 | biological_process | regulation of protein phosphorylation |
| Hydrozetes thienemanni | GO:0007586 | biological_process | digestion |
| Hydrozetes thienemanni | GO:0000209 | biological_process | protein polyubiquitination |
| Hydrozetes thienemanni | GO:0006694 | biological_process | steroid biosynthetic process |
| Hydrozetes thienemanni | GO:0060012 | biological_process | synaptic transmission, glycinergic |
| Hydrozetes thienemanni | GO:0006260 | biological_process | DNA replication |
| Hydrozetes thienemanni | GO:0007608 | biological_process | sensory perception of smell |
| Hydrozetes thienemanni | GO:0006355 | biological_process | regulation of transcription, DNA-templated |
| Hydrozetes thienemanni | GO:0006508 | biological_process | proteolysis |
| Hydrozetes thienemanni | GO:0046872 | molecular_function | metal ion binding |
| Hydrozetes thienemanni | GO:0008270 | molecular_function | zinc ion binding |
| Hydrozetes thienemanni | GO:0051384 | biological_process | response to glucocorticoid |
| Hydrozetes thienemanni | GO:0031098 | biological_process | stress-activated protein kinase signaling cascade |
| Hydrozetes thienemanni | GO:0051603 | biological_process | proteolysis involved in cellular protein catabolic process |
| Hydrozetes thienemanni | GO:0042135 | biological_process | neurotransmitter catabolic process |
| Hydrozetes thienemanni | GO:0090073 | biological_process | positive regulation of protein homodimerization activity |
| Hydrozetes thienemanni | GO:0004497 | molecular_function | monooxygenase activity |
| Hydrozetes thienemanni | GO:0032482 | biological_process | Rab protein signal transduction |
| Hydrozetes thienemanni | GO:0070292 | biological_process | N-acylphosphatidylethanolamine metabolic process |
| Hydrozetes thienemanni | GO:0016192 | biological_process | vesicle-mediated transport |
| Hydrozetes thienemanni | GO:0030433 | biological_process | ER-associated ubiquitin-dependent protein catabolic process |
| Hydrozetes thienemanni | GO:0016705 | molecular_function | oxidoreductase activity, acting on paired donors, with incorporation or reduction of molecular oxygen |
| Hydrozetes thienemanni | GO:0034765 | biological_process | regulation of ion transmembrane transport |

| | | | |
|---|---|---|---|
| Hydrozetes thienemanni | GO:0042572 | biological_process | retinol metabolic process |
| Hydrozetes thienemanni | GO:0006644 | biological_process | phospholipid metabolic process |
| Hydrozetes thienemanni | GO:0070445 | biological_process | regulation of oligodendrocyte progenitor proliferation |
| Hydrozetes thienemanni | GO:0032197 | biological_process | transposition, RNA-mediated |
| Hydrozetes thienemanni | GO:0030194 | biological_process | positive regulation of blood coagulation |
| Hydrozetes thienemanni | GO:1901557 | biological_process | response to fenofibrate |
| Hydrozetes thienemanni | GO:0061077 | biological_process | chaperone-mediated protein folding |
| Hydrozetes thienemanni | GO:0071555 | biological_process | cell wall organization |
| Hydrozetes thienemanni | GO:0032465 | biological_process | regulation of cytokinesis |

| | |
|---|---|
| 61 | 0,000000109692 |
| 47 | 0 |
| 43 | 0,000103030125 |
| 42 | 0 |
| 42 | 0,000000002117[illegible] |
| 37 | 0 |
| 32 | 0,008326090537 |
| 27 | 0,000000626645 |
| 26 | 0,000000000240 |
| 25 | 0 |
| 23 | 0,000000176712 |
| 23 | 0,000003279246 |
| 23 | 0,003472557394 |
| 20 | 0,000000000450 |
| 18 | 0 |
| 16 | 0,007967404623 |
| 14 | 0,000103362007 |
| 14 | 0,006655784478 |
| 13 | 0 |
| 13 | 0,000000000643 |
| 13 | 0,009976027962 |
| 11 | 0 |
| 11 | 0,000001729105 |
| 11 | 0,000049434003 |
| 11 | 0,000394423827 |
| 10 | 0 |
| 10 | 0,000074991459 |
| 10 | 0,002972156164 |
| 10 | 0,00886156028 |
| 10 | 0,01034197571 |
| 9 | 0,000000370460 |
| 9 | 0,002485112557 |
| 8 | 0 |
| 8 | 0,000000007089 |
| 7 | 0,000056215053 |
| 7 | 0,006704529419 |
| 6 | 0 |

| | |
|---|---|
| 6 | 0,000000000960 |
| 6 | 0,006486533136 |
| 6 | 0,007523005626 |
| 6 | 0,0086643631 |
| 5 | 0 |
| 5 | 0 |
| 5 | 0 |
| 5 | 0,000000672459 |
| 5 | 0,000351627598 |
| 4 | 0 |
| 4 | 0 |
| 4 | 0,000007656196 |
| 4 | 0,000054820727 |
| 4 | 0,000054820727 |
| 4 | 0,001088602465 |
| 4 | 0,002333737767 |
| 4 | 0,004304513318 |
| 4 | 0,005327355863 |
| 4 | 0,006852580976 |
| 4 | 0,01259243224 |
| 4 | 0,02498686402 |
| 3 | 0 |
| 3 | 0 |
| 3 | 0 |
| 3 | 0 |
| 3 | 0,000000000293 |
| 3 | 0,000000246491[illegible] |
| 3 | 0,000087006170 |
| 3 | 0,000119014929[illegible] |
| 3 | 0,000755484201 |
| 3 | 0,003577207993 |
| 3 | 0,01091940781 |
| 3 | 0,02574009967 |
| 2 | 0 |
| 2 | 0,000000018315[illegible] |
| 2 | 0,000020473086 |
| 2 | 0,000102742908 |

| | |
|---|---|
| 2 | 0,000638442340 |
| 2 | 0,001990217325 |
| 2 | 0,003440241709 |
| 2 | 0,009140007179 |
| 2 | 0,009719202162 |
| 2 | 0,009719202162 |
| 2 | 0,01830517802 |
| 2 | 0,02398063965 |
| 2 | 0,0261930004 |

| | | | |
|---|---|---|---|
| Hydrozetes lacustris | GO:0032100 | biological_process | positive regulation of appetite |
| Hydrozetes lacustris | GO:0055085 | biological_process | transmembrane transport |
| Hydrozetes lacustris | GO:0007283 | biological_process | spermatogenesis |
| Hydrozetes lacustris | GO:0042908 | biological_process | xenobiotic transport |
| Hydrozetes lacustris | GO:0016020 | cellular_component | membrane |
| Hydrozetes lacustris | GO:0016301 | molecular_function | kinase activity |
| Hydrozetes lacustris | GO:0003677 | molecular_function | DNA binding |
| Hydrozetes lacustris | GO:0006508 | biological_process | proteolysis |
| Hydrozetes lacustris | GO:0007165 | biological_process | signal transduction |
| Hydrozetes lacustris | GO:0030223 | biological_process | neutrophil differentiation |
| Hydrozetes lacustris | GO:0006260 | biological_process | DNA replication |
| Hydrozetes lacustris | GO:0045214 | biological_process | sarcomere organization |
| Hydrozetes lacustris | GO:0016567 | biological_process | protein ubiquitination |
| Hydrozetes lacustris | GO:0055129 | biological_process | L-proline biosynthetic process |
| Hydrozetes lacustris | GO:0007601 | biological_process | visual perception |
| Hydrozetes lacustris | GO:0004364 | molecular_function | glutathione transferase activity |
| Hydrozetes lacustris | GO:0019542 | biological_process | propionate biosynthetic process |
| Hydrozetes lacustris | GO:0008194 | molecular_function | UDP-glycosyltransferase activity |
| Hydrozetes lacustris | GO:0004252 | molecular_function | serine-type endopeptidase activity |
| Hydrozetes lacustris | GO:0004867 | molecular_function | serine-type endopeptidase inhibitor activity |
| Hydrozetes lacustris | GO:0009311 | biological_process | oligosaccharide metabolic process |
| Hydrozetes lacustris | GO:0005506 | molecular_function | iron ion binding |
| Hydrozetes lacustris | GO:0052697 | biological_process | xenobiotic glucuronidation |
| Hydrozetes lacustris | GO:0016192 | biological_process | vesicle-mediated transport |
| Hydrozetes lacustris | GO:0042178 | biological_process | xenobiotic catabolic process |
| Hydrozetes lacustris | GO:0009617 | biological_process | response to bacterium |
| Hydrozetes lacustris | GO:0008270 | molecular_function | zinc ion binding |
| Hydrozetes lacustris | GO:0006486 | biological_process | protein glycosylation |
| Hydrozetes lacustris | GO:0000270 | biological_process | peptidoglycan metabolic process |
| Hydrozetes lacustris | GO:0000272 | biological_process | polysaccharide catabolic process |
| Hydrozetes lacustris | GO:0009888 | biological_process | tissue development |
| Hydrozetes lacustris | GO:0006915 | biological_process | apoptotic process |
| Hydrozetes lacustris | GO:0031098 | biological_process | stress-activated protein kinase signaling cascade |
| Hydrozetes lacustris | GO:0006511 | biological_process | ubiquitin-dependent protein catabolic process |
| Hydrozetes lacustris | GO:0010992 | biological_process | ubiquitin homeostasis |
| Hydrozetes lacustris | GO:0071357 | biological_process | cellular response to type I interferon |
| Hydrozetes lacustris | GO:0005975 | biological_process | carbohydrate metabolic process |

| | | | |
|---|---|---|---|
| Hydrozetes lacustris | GO:0045599 | biological_process | negative regulation of fat cell differentiation |
| Hydrozetes lacustris | GO:0051260 | biological_process | protein homooligomerization |
| Hydrozetes lacustris | GO:0016491 | molecular_function | oxidoreductase activity |
| Hydrozetes lacustris | GO:0006468 | biological_process | protein phosphorylation |
| Hydrozetes lacustris | GO:0006355 | biological_process | regulation of transcription, DNA-templated |
| Hydrozetes lacustris | GO:0043171 | biological_process | peptide catabolic process |
| Hydrozetes lacustris | GO:0016042 | biological_process | lipid catabolic process |
| Hydrozetes lacustris | GO:0051384 | biological_process | response to glucocorticoid |
| Hydrozetes lacustris | GO:0006069 | biological_process | ethanol oxidation |
| Hydrozetes lacustris | GO:0007016 | biological_process | cytoskeletal anchoring at plasma membrane |
| Hydrozetes lacustris | GO:0060049 | biological_process | regulation of protein glycosylation |
| Hydrozetes lacustris | GO:0010951 | biological_process | negative regulation of endopeptidase activity |
| Hydrozetes lacustris | GO:0010842 | biological_process | retina layer formation |
| Hydrozetes lacustris | GO:0070189 | biological_process | kynurenine metabolic process |
| Hydrozetes lacustris | GO:0004518 | molecular_function | nuclease activity |
| Hydrozetes lacustris | GO:0004983 | molecular_function | neuropeptide Y receptor activity |
| Hydrozetes lacustris | GO:0006805 | biological_process | xenobiotic metabolic process |
| Hydrozetes lacustris | GO:0031030 | biological_process | negative regulation of septation initiation signaling |
| Hydrozetes lacustris | GO:0006470 | biological_process | protein dephosphorylation |
| Hydrozetes lacustris | GO:0004722 | molecular_function | protein serine/threonine phosphatase activity |
| Hydrozetes lacustris | GO:0006605 | biological_process | protein targeting |
| Hydrozetes lacustris | GO:0002376 | biological_process | immune system process |
| Hydrozetes lacustris | GO:0015267 | molecular_function | channel activity |
| Hydrozetes lacustris | GO:0009168 | biological_process | purine ribonucleoside monophosphate biosynthetic process |
| Hydrozetes lacustris | GO:0032434 | biological_process | regulation of proteasomal ubiquitin-dependent protein catabolic process |
| Hydrozetes lacustris | GO:0006090 | biological_process | pyruvate metabolic process |
| Hydrozetes lacustris | GO:0006412 | biological_process | translation |
| Hydrozetes lacustris | GO:0010139 | biological_process | pyrimidine deoxyribonucleotide salvage |
| Hydrozetes lacustris | GO:0001963 | biological_process | synaptic transmission, dopaminergic |
| Hydrozetes lacustris | GO:0042135 | biological_process | neurotransmitter catabolic process |
| Hydrozetes lacustris | GO:0006152 | biological_process | purine nucleoside catabolic process |
| Hydrozetes lacustris | GO:0000209 | biological_process | protein polyubiquitination |
| Hydrozetes lacustris | GO:0007605 | biological_process | sensory perception of sound |
| Hydrozetes lacustris | GO:0004497 | molecular_function | monooxygenase activity |
| Hydrozetes lacustris | GO:0042060 | biological_process | wound healing |
| Hydrozetes lacustris | GO:0016031 | biological_process | tRNA import into mitochondrion |
| Hydrozetes lacustris | GO:0001669 | cellular_component | acrosomal vesicle |

| | | | |
|---|---|---|---|
| Hydrozetes lacustris | GO:0042110 | biological_process | T cell activation |
| Hydrozetes lacustris | GO:0003309 | biological_process | type B pancreatic cell differentiation |
| Hydrozetes lacustris | GO:0006979 | biological_process | response to oxidative stress |
| Hydrozetes lacustris | GO:0097066 | biological_process | response to thyroid hormone |
| Hydrozetes lacustris | GO:0055069 | biological_process | zinc ion homeostasis |
| Hydrozetes lacustris | GO:0014883 | biological_process | transition between fast and slow fiber |
| Hydrozetes lacustris | GO:0031222 | biological_process | arabinan catabolic process |
| Hydrozetes lacustris | GO:0007275 | biological_process | multicellular organismal development |
| Hydrozetes lacustris | GO:0008380 | biological_process | RNA splicing |
| Hydrozetes lacustris | GO:0006099 | biological_process | tricarboxylic acid cycle |
| Hydrozetes lacustris | GO:0008218 | biological_process | bioluminescence |
| Hydrozetes lacustris | GO:0009263 | biological_process | deoxyribonucleotide biosynthetic process |
| Hydrozetes lacustris | GO:0019882 | biological_process | antigen processing and presentation |
| Hydrozetes lacustris | GO:0051598 | biological_process | meiotic recombination checkpoint |
| Hydrozetes lacustris | GO:0021762 | biological_process | substantia nigra development |
| Hydrozetes lacustris | GO:0016117 | biological_process | carotenoid biosynthetic process |
| Hydrozetes lacustris | GO:0039648 | biological_process | modulation by virus of host protein ubiquitination |
| Hydrozetes lacustris | GO:0042572 | biological_process | retinol metabolic process |
| Hydrozetes lacustris | GO:0016021 | cellular_component | integral component of membrane |
| Hydrozetes lacustris | GO:0007399 | biological_process | nervous system development |
| Hydrozetes lacustris | GO:0007179 | biological_process | transforming growth factor beta receptor signaling pathway |
| Hydrozetes lacustris | GO:0000070 | biological_process | mitotic sister chromatid segregation |
| Hydrozetes lacustris | GO:0005739 | cellular_component | mitochondrion |
| Hydrozetes lacustris | GO:0016887 | molecular_function | ATPase activity |
| Hydrozetes lacustris | GO:0004222 | molecular_function | metalloendopeptidase activity |
| Hydrozetes lacustris | GO:1901639 | biological_process | XDP catabolic process |
| Hydrozetes lacustris | GO:0016055 | biological_process | Wnt signaling pathway |
| Hydrozetes lacustris | GO:0046872 | molecular_function | metal ion binding |
| Hydrozetes lacustris | GO:0042761 | biological_process | very long-chain fatty acid biosynthetic process |
| Hydrozetes lacustris | GO:0007155 | biological_process | cell adhesion |
| Hydrozetes lacustris | GO:0050951 | biological_process | sensory perception of temperature stimulus |
| Hydrozetes lacustris | GO:0006629 | biological_process | lipid metabolic process |
| Hydrozetes lacustris | GO:0045893 | biological_process | positive regulation of transcription, DNA-templated |
| Hydrozetes lacustris | GO:0006325 | biological_process | chromatin organization |
| Hydrozetes lacustris | GO:0006694 | biological_process | steroid biosynthetic process |
| Hydrozetes lacustris | GO:0031638 | biological_process | zymogen activation |
| Hydrozetes lacustris | GO:0010224 | biological_process | response to UV-B |

| | | | |
|---|---|---|---|
| Hydrozetes lacustris | GO:0008240 | molecular_function | tripeptidyl-peptidase activity |
| Hydrozetes lacustris | GO:0060074 | biological_process | synapse maturation |
| Hydrozetes lacustris | GO:0016485 | biological_process | protein processing |
| Hydrozetes lacustris | GO:0075732 | biological_process | viral penetration into host nucleus |
| Hydrozetes lacustris | GO:0009213 | biological_process | pyrimidine deoxyribonucleoside triphosphate catabolic process |
| Hydrozetes lacustris | GO:0030246 | molecular_function | carbohydrate binding |
| Hydrozetes lacustris | GO:0015031 | biological_process | protein transport |
| Hydrozetes lacustris | GO:0016032 | biological_process | viral process |
| Hydrozetes lacustris | GO:0005634 | cellular_component | nucleus |
| Hydrozetes lacustris | GO:0005576 | cellular_component | extracellular region |
| Hydrozetes lacustris | GO:0016705 | molecular_function | oxidoreductase activity, acting on paired donors, with incorporation or reduction of molecular oxygen |
| Hydrozetes lacustris | GO:0008202 | biological_process | steroid metabolic process |
| Hydrozetes lacustris | GO:0006641 | biological_process | triglyceride metabolic process |
| Hydrozetes lacustris | GO:0007584 | biological_process | response to nutrient |
| Hydrozetes lacustris | GO:0045087 | biological_process | innate immune response |
| Hydrozetes lacustris | GO:0030162 | biological_process | regulation of proteolysis |
| Hydrozetes lacustris | GO:0001505 | biological_process | regulation of neurotransmitter levels |
| Hydrozetes lacustris | GO:0005737 | cellular_component | cytoplasm |
| Hydrozetes lacustris | GO:0045944 | biological_process | positive regulation of transcription from RNA polymerase II promoter |
| Hydrozetes lacustris | GO:0006351 | biological_process | transcription, DNA-templated |
| Hydrozetes lacustris | GO:0006890 | biological_process | retrograde vesicle-mediated transport, Golgi to ER |
| Hydrozetes lacustris | GO:0007186 | biological_process | G-protein coupled receptor signaling pathway |
| Hydrozetes lacustris | GO:0035220 | biological_process | wing disc development |
| Hydrozetes lacustris | GO:0008168 | molecular_function | methyltransferase activity |
| Hydrozetes lacustris | GO:0010468 | biological_process | regulation of gene expression |
| Hydrozetes lacustris | GO:0030163 | biological_process | protein catabolic process |
| Hydrozetes lacustris | GO:0035556 | biological_process | intracellular signal transduction |
| Hydrozetes lacustris | GO:0010506 | biological_process | regulation of autophagy |
| Hydrozetes lacustris | GO:0032496 | biological_process | response to lipopolysaccharide |
| Hydrozetes lacustris | GO:0004104 | molecular_function | cholinesterase activity |
| Hydrozetes lacustris | GO:0032880 | biological_process | regulation of protein localization |
| Hydrozetes lacustris | GO:0042311 | biological_process | vasodilation |
| Hydrozetes lacustris | GO:0097264 | biological_process | self proteolysis |
| Hydrozetes lacustris | GO:0097090 | biological_process | presynaptic membrane organization |
| Hydrozetes lacustris | GO:0004970 | molecular_function | ionotropic glutamate receptor activity |
| Hydrozetes lacustris | GO:0007585 | biological_process | respiratory gaseous exchange |
| Hydrozetes lacustris | GO:0017148 | biological_process | negative regulation of translation |

| | |
|---|---|
| 218 | 0 |
| 162 | 0,01823730816 |
| 125 | 0,004470838693 |
| 96 | 0 |
| 81 | 0 |
| 78 | 0 |
| 70 | 0,000000735472 |
| 65 | 0,000000021141[illegible] |
| 64 | 0 |
| 58 | 0 |
| 52 | 0 |
| 52 | 0,000000046129 |
| 45 | 0,000000013455 |
| 44 | 0 |
| 44 | 0,000000000104 |
| 44 | 0,000000001858 |
| 41 | 0 |
| 40 | 0 |
| 40 | 0,000177421388 |
| 38 | 0,000004455116[illegible] |
| 36 | 0 |
| 35 | 0,007351649326 |
| 33 | 0,000000002474 |
| 32 | 0 |
| 32 | 0,000000000789 |
| 32 | 0,000000280683 |
| 31 | 0,000000000167[illegible] |
| 31 | 0,000000302609 |
| 30 | 0 |
| 30 | 0,01261661752 |
| 28 | 0,000141925367 |
| 26 | 0,001821490902 |
| 26 | 0,003768550233 |
| 25 | 0 |
| 25 | 0 |
| 24 | 0,000000066846 |
| 23 | 0,000000001080 |

| | |
|---|---|
| 23 | 0,000011951855[illegible] |
| 23 | 0,003419480765 |
| 22 | 0,000000212917 |
| 21 | 0 |
| 20 | 0 |
| 20 | 0 |
| 20 | 0,000000952942 |
| 19 | 0,000000007514 |
| 19 | 0,001218048937 |
| 18 | 0,000000006872 |
| 18 | 0,000000006872 |
| 18 | 0,007985273084 |
| 18 | 0,01432938251 |
| 17 | 0,000001498764 |
| 17 | 0,001048886759 |
| 17 | 0,003496324567 |
| 16 | 0,000000003717 |
| 16 | 0,000000051142 |
| 16 | 0,000078763010 |
| 16 | 0,000139962425 |
| 16 | 0,000173957780 |
| 16 | 0,01012947945 |
| 16 | 0,01216509301 |
| 15 | 0,000000001323 |
| 15 | 0,000004776001 |
| 15 | 0,000013402122 |
| 14 | 0 |
| 14 | 0 |
| 14 | 0,000538304574 |
| 14 | 0,001221696565 |
| 13 | 0,000000008783 |
| 13 | 0,009780647322 |
| 12 | 0,000013146597 |
| 12 | 0,01628969239 |
| 11 | 0,000000107796 |
| 11 | 0,000012791562 |
| 11 | 0,000052322306 |

| | |
|---|---|
| 11 | 0,000100979818 |
| 11 | 0,01310116735 |
| 10 | 0 |
| 10 | 0,000000846605 |
| 10 | 0,000021233710 |
| 10 | 0,000125340857 |
| 10 | 0,002274289657 |
| 9 | 0 |
| 9 | 0 |
| 9 | 0,000000044114 |
| 9 | 0,002191637667 |
| 9 | 0,006059959107 |
| 9 | 0,01022650211 |
| 8 | 0,000000018252 |
| 8 | 0,005896712352 |
| 8 | 0,01064544475 |
| 8 | 0,01266939351 |
| 7 | 0,000187821209 |
| 6 | 0 |
| 6 | 0,000015071402 |
| 6 | 0,000019729893 |
| 6 | 0,000212213804 |
| 6 | 0,001040076109 |
| 6 | 0,001313962593 |
| 6 | 0,001533171441 |
| 6 | 0,007556637333 |
| 5 | 0 |
| 5 | 0 |
| 5 | 0 |
| 5 | 0,000006316067 |
| 5 | 0,000079286014 |
| 5 | 0,000098943141 |
| 4 | 0 |
| 4 | 0,000000000153 |
| 4 | 0,000001279097 |
| 4 | 0,000010096299 |
| 4 | 0,000014846475 |

| | |
|---|---|
| 4 | 0,000137473058 |
| 4 | 0,000613338558 |
| 4 | 0,001798113318 |
| 4 | 0,003418080789 |
| 4 | 0,006099351313 |
| 4 | 0,01394338025 |
| 3 | 0 |
| 3 | 0 |
| 3 | 0 |
| 3 | 0 |
| 3 | 0,000000000597 |
| 3 | 0,000000986055 |
| 3 | 0,000018776179 |
| 3 | 0,000310715430 |
| 3 | 0,000374642306 |
| 3 | 0,001233334341 |
| 3 | 0,001475362932 |
| 3 | 0,01232131262 |
| 2 | 0 |
| 2 | 0 |
| 2 | 0 |
| 2 | 0,000000000851 |
| 2 | 0,000000474092 |
| 2 | 0,000029574292 |
| 2 | 0,000091342230 |
| 2 | 0,000371986638 |
| 2 | 0,000551560451 |
| 2 | 0,000608297742 |
| 2 | 0,000608297742 |
| 2 | 0,001597749247 |
| 2 | 0,001597749247 |
| 2 | 0,004474412955 |
| 2 | 0,007712203524 |
| 2 | 0,01177569282 |
| 2 | 0,01428762839 |
| 2 | 0,01847857478 |
| 2 | 0,02187772943 |

| | | | | |
|---|---|---|---|---|
| Achipteria coleoptrata | GO:0055085 | biological_process | transmembrane transport | 347 |
| Achipteria coleoptrata | GO:0032100 | biological_process | positive regulation of appetite | 218 |
| Achipteria coleoptrata | GO:0042908 | biological_process | xenobiotic transport | 96 |
| Achipteria coleoptrata | GO:0003677 | molecular_function | DNA binding | 89 |
| Achipteria coleoptrata | GO:0015074 | biological_process | DNA integration | 84 |
| Achipteria coleoptrata | GO:0003964 | molecular_function | RNA-directed DNA polymerase activity | 79 |
| Achipteria coleoptrata | GO:0016301 | molecular_function | kinase activity | 77 |
| Achipteria coleoptrata | GO:0055129 | biological_process | L-proline biosynthetic process | 75 |
| Achipteria coleoptrata | GO:0032197 | biological_process | transposition, RNA-mediated | 73 |
| Achipteria coleoptrata | GO:0008218 | biological_process | bioluminescence | 72 |
| Achipteria coleoptrata | GO:0007283 | biological_process | spermatogenesis | 69 |
| Achipteria coleoptrata | GO:0006313 | biological_process | transposition, DNA-mediated | 60 |
| Achipteria coleoptrata | GO:0030223 | biological_process | neutrophil differentiation | 58 |
| Achipteria coleoptrata | GO:0006633 | biological_process | fatty acid biosynthetic process | 58 |
| Achipteria coleoptrata | GO:0006511 | biological_process | ubiquitin-dependent protein catabolic process | 57 |
| Achipteria coleoptrata | GO:0007601 | biological_process | visual perception | 57 |
| Achipteria coleoptrata | GO:0008194 | molecular_function | UDP-glycosyltransferase activity | 49 |
| Achipteria coleoptrata | GO:0009313 | biological_process | oligosaccharide catabolic process | 47 |
| Achipteria coleoptrata | GO:0006508 | biological_process | proteolysis | 47 |
| Achipteria coleoptrata | GO:0006334 | biological_process | nucleosome assembly | 46 |
| Achipteria coleoptrata | GO:0006646 | biological_process | phosphatidylethanolamine biosynthetic process | 44 |
| Achipteria coleoptrata | GO:0010224 | biological_process | response to UV-B | 42 |
| Achipteria coleoptrata | GO:0006706 | biological_process | steroid catabolic process | 40 |
| Achipteria coleoptrata | GO:0003081 | biological_process | regulation of systemic arterial blood pressure by renin-angiotensin | 39 |
| Achipteria coleoptrata | GO:0007275 | biological_process | multicellular organismal development | 39 |
| Achipteria coleoptrata | GO:0006749 | biological_process | glutathione metabolic process | 38 |
| Achipteria coleoptrata | GO:0006517 | biological_process | protein deglycosylation | 34 |
| Achipteria coleoptrata | GO:0016042 | biological_process | lipid catabolic process | 33 |
| Achipteria coleoptrata | GO:0001516 | biological_process | prostaglandin biosynthetic process | 32 |
| Achipteria coleoptrata | GO:0006805 | biological_process | xenobiotic metabolic process | 32 |
| Achipteria coleoptrata | GO:0006013 | biological_process | mannose metabolic process | 30 |
| Achipteria coleoptrata | GO:0016567 | biological_process | protein ubiquitination | 29 |
| Achipteria coleoptrata | GO:0004222 | molecular_function | metalloendopeptidase activity | 29 |
| Achipteria coleoptrata | GO:0046548 | biological_process | retinal rod cell development | 28 |
| Achipteria coleoptrata | GO:0009617 | biological_process | response to bacterium | 28 |
| Achipteria coleoptrata | GO:0006310 | biological_process | DNA recombination | 28 |
| Achipteria coleoptrata | GO:1902358 | biological_process | sulfate transmembrane transport | 27 |

| | | | | |
|---|---|---|---|---|
| Achipteria coleoptrata | GO:0043252 | biological_process | sodium-independent organic anion transport | 27 |
| Achipteria coleoptrata | GO:0009888 | biological_process | tissue development | 27 |
| Achipteria coleoptrata | GO:0031098 | biological_process | stress-activated protein kinase signaling cascade | 26 |
| Achipteria coleoptrata | GO:0006915 | biological_process | apoptotic process | 26 |
| Achipteria coleoptrata | GO:0010992 | biological_process | ubiquitin homeostasis | 25 |
| Achipteria coleoptrata | GO:0044245 | biological_process | polysaccharide digestion | 25 |
| Achipteria coleoptrata | GO:0071357 | biological_process | cellular response to type I interferon | 24 |
| Achipteria coleoptrata | GO:0045616 | biological_process | regulation of keratinocyte differentiation | 24 |
| Achipteria coleoptrata | GO:0004252 | molecular_function | serine-type endopeptidase activity | 23 |
| Achipteria coleoptrata | GO:0016491 | molecular_function | oxidoreductase activity | 23 |
| Achipteria coleoptrata | GO:0007165 | biological_process | signal transduction | 22 |
| Achipteria coleoptrata | GO:0016192 | biological_process | vesicle-mediated transport | 22 |
| Achipteria coleoptrata | GO:0006412 | biological_process | translation | 21 |
| Achipteria coleoptrata | GO:0007169 | biological_process | transmembrane receptor protein tyrosine kinase signaling pathway | 20 |
| Achipteria coleoptrata | GO:0010842 | biological_process | retina layer formation | 20 |
| Achipteria coleoptrata | GO:0006468 | biological_process | protein phosphorylation | 19 |
| Achipteria coleoptrata | GO:0008210 | biological_process | estrogen metabolic process | 19 |
| Achipteria coleoptrata | GO:0006069 | biological_process | ethanol oxidation | 19 |
| Achipteria coleoptrata | GO:0005087 | molecular_function | Ran guanyl-nucleotide exchange factor activity | 19 |
| Achipteria coleoptrata | GO:0009636 | biological_process | response to toxic substance | 18 |
| Achipteria coleoptrata | GO:0010951 | biological_process | negative regulation of endopeptidase activity | 18 |
| Achipteria coleoptrata | GO:0006979 | biological_process | response to oxidative stress | 17 |
| Achipteria coleoptrata | GO:0008146 | molecular_function | sulfotransferase activity | 17 |
| Achipteria coleoptrata | GO:0051923 | biological_process | sulfation | 17 |
| Achipteria coleoptrata | GO:0034976 | biological_process | response to endoplasmic reticulum stress | 16 |
| Achipteria coleoptrata | GO:0007286 | biological_process | spermatid development | 16 |
| Achipteria coleoptrata | GO:0021591 | biological_process | ventricular system development | 16 |
| Achipteria coleoptrata | GO:0008270 | molecular_function | zinc ion binding | 15 |
| Achipteria coleoptrata | GO:0009263 | biological_process | deoxyribonucleotide biosynthetic process | 15 |
| Achipteria coleoptrata | GO:0032434 | biological_process | regulation of proteasomal ubiquitin-dependent protein catabolic process | 15 |
| Achipteria coleoptrata | GO:0006090 | biological_process | pyruvate metabolic process | 15 |
| Achipteria coleoptrata | GO:0006810 | biological_process | transport | 15 |
| Achipteria coleoptrata | GO:0016055 | biological_process | Wnt signaling pathway | 14 |
| Achipteria coleoptrata | GO:0005576 | cellular_component | extracellular region | 14 |
| Achipteria coleoptrata | GO:0004674 | molecular_function | protein serine/threonine kinase activity | 14 |
| Achipteria coleoptrata | GO:0033089 | biological_process | positive regulation of T cell differentiation in thymus | 13 |

| | | | | |
|---|---|---|---|---|
| Achipteria coleoptrata | GO:0016117 | biological_process | carotenoid biosynthetic process | 13 |
| Achipteria coleoptrata | GO:0042135 | biological_process | neurotransmitter catabolic process | 13 |
| Achipteria coleoptrata | GO:0008202 | biological_process | steroid metabolic process | 13 |
| Achipteria coleoptrata | GO:0036324 | biological_process | vascular endothelial growth factor receptor-2 signaling pathway | 12 |
| Achipteria coleoptrata | GO:0005983 | biological_process | starch catabolic process | 12 |
| Achipteria coleoptrata | GO:0007419 | biological_process | ventral cord development | 12 |
| Achipteria coleoptrata | GO:0006941 | biological_process | striated muscle contraction | 12 |
| Achipteria coleoptrata | GO:0005506 | molecular_function | iron ion binding | 12 |
| Achipteria coleoptrata | GO:0019233 | biological_process | sensory perception of pain | 12 |
| Achipteria coleoptrata | GO:0006486 | biological_process | protein glycosylation | 11 |
| Achipteria coleoptrata | GO:0010383 | biological_process | cell wall polysaccharide metabolic process | 11 |
| Achipteria coleoptrata | GO:0005509 | molecular_function | calcium ion binding | 11 |
| Achipteria coleoptrata | GO:0006355 | biological_process | regulation of transcription, DNA-templated | 10 |
| Achipteria coleoptrata | GO:0008380 | biological_process | RNA splicing | 10 |
| Achipteria coleoptrata | GO:0042060 | biological_process | wound healing | 10 |
| Achipteria coleoptrata | GO:0055069 | biological_process | zinc ion homeostasis | 10 |
| Achipteria coleoptrata | GO:0014883 | biological_process | transition between fast and slow fiber | 10 |
| Achipteria coleoptrata | GO:0007224 | biological_process | smoothened signaling pathway | 10 |
| Achipteria coleoptrata | GO:0039648 | biological_process | modulation by virus of host protein ubiquitination | 10 |
| Achipteria coleoptrata | GO:0042572 | biological_process | retinol metabolic process | 9 |
| Achipteria coleoptrata | GO:0006811 | biological_process | ion transport | 9 |
| Achipteria coleoptrata | GO:0006629 | biological_process | lipid metabolic process | 9 |
| Achipteria coleoptrata | GO:0042302 | molecular_function | structural constituent of cuticle | 8 |
| Achipteria coleoptrata | GO:0006470 | biological_process | protein dephosphorylation | 8 |
| Achipteria coleoptrata | GO:0006694 | biological_process | steroid biosynthetic process | 8 |
| Achipteria coleoptrata | GO:0008241 | molecular_function | peptidyl-dipeptidase activity | 8 |
| Achipteria coleoptrata | GO:0016021 | cellular_component | integral component of membrane | 7 |
| Achipteria coleoptrata | GO:0046872 | molecular_function | metal ion binding | 7 |
| Achipteria coleoptrata | GO:0031638 | biological_process | zymogen activation | 7 |
| Achipteria coleoptrata | GO:0005739 | cellular_component | mitochondrion | 7 |
| Achipteria coleoptrata | GO:1901639 | biological_process | XDP catabolic process | 7 |
| Achipteria coleoptrata | GO:0006357 | biological_process | regulation of transcription from RNA polymerase II promoter | 6 |
| Achipteria coleoptrata | GO:0007399 | biological_process | nervous system development | 6 |
| Achipteria coleoptrata | GO:0007155 | biological_process | cell adhesion | 6 |
| Achipteria coleoptrata | GO:0015939 | biological_process | pantothenate metabolic process | 6 |
| Achipteria coleoptrata | GO:0001501 | biological_process | skeletal system development | 6 |
| Achipteria coleoptrata | GO:0042594 | biological_process | response to starvation | 6 |

| | | | | |
|---|---|---|---|---|
| Achipteria coleoptrata | GO:0006351 | biological_process | transcription, DNA-templated | 5 |
| Achipteria coleoptrata | GO:0045893 | biological_process | positive regulation of transcription, DNA-templated | 5 |
| Achipteria coleoptrata | GO:0036372 | biological_process | opsin transport | 5 |
| Achipteria coleoptrata | GO:0050951 | biological_process | sensory perception of temperature stimulus | 5 |
| Achipteria coleoptrata | GO:0000272 | biological_process | polysaccharide catabolic process | 4 |
| Achipteria coleoptrata | GO:0007605 | biological_process | sensory perception of sound | 4 |
| Achipteria coleoptrata | GO:0055114 | biological_process | oxidation-reduction process | 4 |
| Achipteria coleoptrata | GO:0060012 | biological_process | synaptic transmission, glycinergic | 4 |
| Achipteria coleoptrata | GO:0006641 | biological_process | triglyceride metabolic process | 4 |
| Achipteria coleoptrata | GO:0046982 | molecular_function | protein heterodimerization activity | 4 |
| Achipteria coleoptrata | GO:0008240 | molecular_function | tripeptidyl-peptidase activity | 4 |
| Achipteria coleoptrata | GO:0007416 | biological_process | synapse assembly | 4 |
| Achipteria coleoptrata | GO:0030162 | biological_process | regulation of proteolysis | 4 |
| Achipteria coleoptrata | GO:0008654 | biological_process | phospholipid biosynthetic process | 4 |
| Achipteria coleoptrata | GO:0052697 | biological_process | xenobiotic glucuronidation | 4 |
| Achipteria coleoptrata | GO:0031175 | biological_process | neuron projection development | 4 |
| Achipteria coleoptrata | GO:0048741 | biological_process | skeletal muscle fiber development | 4 |
| Achipteria coleoptrata | GO:0051384 | biological_process | response to glucocorticoid | 3 |
| Achipteria coleoptrata | GO:0030433 | biological_process | ER-associated ubiquitin-dependent protein catabolic process | 3 |
| Achipteria coleoptrata | GO:0009405 | biological_process | pathogenesis | 3 |
| Achipteria coleoptrata | GO:0032482 | biological_process | Rab protein signal transduction | 3 |
| Achipteria coleoptrata | GO:0007584 | biological_process | response to nutrient | 3 |
| Achipteria coleoptrata | GO:0045087 | biological_process | innate immune response | 3 |
| Achipteria coleoptrata | GO:0008360 | biological_process | regulation of cell shape | 3 |
| Achipteria coleoptrata | GO:0090073 | biological_process | positive regulation of protein homodimerization activity | 3 |
| Achipteria coleoptrata | GO:0004983 | molecular_function | neuropeptide Y receptor activity | 3 |
| Achipteria coleoptrata | GO:0005737 | cellular_component | cytoplasm | 3 |
| Achipteria coleoptrata | GO:0000011 | biological_process | vacuole inheritance | 3 |
| Achipteria coleoptrata | GO:0097264 | biological_process | self proteolysis | 3 |
| Achipteria coleoptrata | GO:0004518 | molecular_function | nuclease activity | 3 |
| Achipteria coleoptrata | GO:0045944 | biological_process | positive regulation of transcription from RNA polymerase II promoter | 2 |
| Achipteria coleoptrata | GO:0006457 | biological_process | protein folding | 2 |
| Achipteria coleoptrata | GO:0042626 | molecular_function | ATPase activity, coupled to transmembrane movement of substances | 2 |
| Achipteria coleoptrata | GO:0035249 | biological_process | synaptic transmission, glutamatergic | 2 |
| Achipteria coleoptrata | GO:0007186 | biological_process | G-protein coupled receptor signaling pathway | 2 |
| Achipteria coleoptrata | GO:0034765 | biological_process | regulation of ion transmembrane transport | 2 |

| | | | | |
|---|---|---|---|---|
| Achipteria coleoptrata | GO:0030435 | biological_process | sporulation resulting in formation of a cellular spore | 2 |
| Achipteria coleoptrata | GO:0035220 | biological_process | wing disc development | 2 |
| Achipteria coleoptrata | GO:0006644 | biological_process | phospholipid metabolic process | 2 |
| Achipteria coleoptrata | GO:0051301 | biological_process | cell division | 2 |
| Achipteria coleoptrata | GO:0050896 | biological_process | response to stimulus | 2 |
| Achipteria coleoptrata | GO:0001659 | biological_process | temperature homeostasis | 2 |
| Achipteria coleoptrata | GO:0035556 | biological_process | intracellular signal transduction | 2 |
| Achipteria coleoptrata | GO:0042981 | biological_process | regulation of apoptotic process | 2 |
| Achipteria coleoptrata | GO:0004104 | molecular_function | cholinesterase activity | 2 |
| Achipteria coleoptrata | GO:0032880 | biological_process | regulation of protein localization | 2 |
| Achipteria coleoptrata | GO:0002376 | biological_process | immune system process | 2 |
| Achipteria coleoptrata | GO:0042311 | biological_process | vasodilation | 2 |
| Achipteria coleoptrata | GO:0010718 | biological_process | positive regulation of epithelial to mesenchymal transition | 2 |
| Achipteria coleoptrata | GO:0008289 | molecular_function | lipid binding | 2 |
| Achipteria coleoptrata | GO:0008063 | biological_process | Toll signaling pathway | 2 |
| Achipteria coleoptrata | GO:0007338 | biological_process | single fertilization | 2 |
| Achipteria coleoptrata | GO:0043434 | biological_process | response to peptide hormone | 2 |
| Achipteria coleoptrata | GO:1902542 | biological_process | regulation of protein localization to mitotic spindle pole body | 2 |
| Achipteria coleoptrata | GO:0005154 | molecular_function | epidermal growth factor receptor binding | 2 |
| Achipteria coleoptrata | GO:0017148 | biological_process | negative regulation of translation | 2 |
| Achipteria coleoptrata | GO:0004842 | molecular_function | ubiquitin-protein transferase activity | 2 |
| Achipteria coleoptrata | GO:0033197 | biological_process | response to vitamin E | 2 |

| |
|---|
| 0 |
| 0 |
| 0 |
| 0 |
| 0 |
| 0 |
| 0 |
| 0 |
| 0 |
| 0 |
| 0 |
| 0,00255351576 |
| 0 |
| 0,0227993762 |
| 0 |
| 0,000000043734 |
| 0 |
| 0 |
| 0 |
| 0,000019899602 |
| 0,002952808757 |
| 0,000000949856 |
| 0,003422309163 |
| 0 |
| 0 |
| 0,000037290385 |
| 0,000093732016 |
| 0,001233763724 |
| 0,000000000302 |
| 0,000343491288 |
| 0,000000297102 |
| 0 |
| 0,001627256227 |
| 0 |
| 0,000000000162 |
| 0,000000061742 |
| 0 |

| |
|---|
| 0,000069147880 |
| 0,001001785599 |
| 0,000929499801 |
| 0,004628475991 |
| 0 |
| 0 |
| 0,000000284842 |
| 0,000005346780 |
| 0 |
| 0,000000030741 |
| 0 |
| 0 |
| 0 |
| 0,002929481945 |
| 0,008224173249 |
| 0 |
| 0 |
| 0,002760267961 |
| 0,008124803999 |
| 0,000014260751 |
| 0,01496751388 |
| 0 |
| 0,000000059364 |
| 0,003000681983 |
| 0,000070138211[illegible] |
| 0,000367627086 |
| 0,004053471817 |
| 0 |
| 0,000001648116 |
| 0,000012158900[illegible] |
| 0,000033127599 |
| 0,000137243377 |
| 0 |
| 0,000029117196[illegible] |
| 0,02095326274 |
| 0,000000008911[illegible] |

| |
|---|
| 0,000015719725 |
| 0,000142102304 |
| 0,01278233016 |
| 0,000001137464[illegible] |
| 0,000002105287 |
| 0,000003390574 |
| 0,000162884484 |
| 0,000337580925 |
| 0,00363195997 |
| 0 |
| 0,000890807250 |
| 0,003024288874 |
| 0 |
| 0 |
| 0,000000002238 |
| 0,000041393357 |
| 0,000234950424 |
| 0,001289775581 |
| 0,001963642261 |
| 0,000452317242 |
| 0,002095597164 |
| 0,002254293688 |
| 0 |
| 0,000000000534 |
| 0,000075007681 |
| 0,000977285214 |
| 0 |
| 0 |
| 0,000154730526 |
| 0,001096670907 |
| 0,002406987895 |
| 0 |
| 0,000003302033 |
| 0,000006011269[illegible] |
| 0,000324207176 |
| 0,01061007514 |
| 0,01733886103 |

0
0
0,000021763782
0,000115417809
0
0
0,000000007682
0,000000483563
0,000027391691
0,000184889628
0,000187030969
0,000301030480
0,001960012136
0,00234578085
0,003343917069
0,01756994231
0,01897772542
0
0
0,000002731456
0,000003803714
0,000117544814
0,000144626313
0,004514054766
0,005668217864
0,007111507096
0,007111507096
0,01251787278
0,01433343145
0,01560603243

0
0

0
0
0,000000000102
0,000000000189

| |
|---|
| 0,000000001526 |
| 0,000000102828 |
| 0,000000165066 |
| 0,000004819907[illegible] |
| 0,000099860352 |
| 0,000111440441[illegible] |
| 0,000238779235 |
| 0,000238779235 |
| 0,000772469136 |
| 0,000772469136 |
| 0,001445281844 |
| 0,00241500277 |
| 0,002673401454 |
| 0,004419474697 |
| 0,009665290411 |
| 0,009665290411 |
| 0,01063587427 |
| 0,01373025039 |
| 0,01373025039 |
| 0,01412779626 |
| 0,02453741347 |
| 0,02453741347 |

| | | | |
|---|---|---|---|
| Platynothrus peltifer | GO:0032100 | biological_process | positive regulation of appetite |
| Platynothrus peltifer | GO:0007283 | biological_process | spermatogenesis |
| Platynothrus peltifer | GO:0016301 | molecular_function | kinase activity |
| Platynothrus peltifer | GO:0005975 | biological_process | carbohydrate metabolic process |
| Platynothrus peltifer | GO:0006633 | biological_process | fatty acid biosynthetic process |
| Platynothrus peltifer | GO:0051384 | biological_process | response to glucocorticoid |
| Platynothrus peltifer | GO:0004364 | molecular_function | glutathione transferase activity |
| Platynothrus peltifer | GO:0030223 | biological_process | neutrophil differentiation |
| Platynothrus peltifer | GO:0006313 | biological_process | transposition, DNA-mediated |
| Platynothrus peltifer | GO:0055085 | biological_process | transmembrane transport |
| Platynothrus peltifer | GO:0009313 | biological_process | oligosaccharide catabolic process |
| Platynothrus peltifer | GO:0006508 | biological_process | proteolysis |
| Platynothrus peltifer | GO:0007601 | biological_process | visual perception |
| Platynothrus peltifer | GO:0010025 | biological_process | wax biosynthetic process |
| Platynothrus peltifer | GO:0019542 | biological_process | propionate biosynthetic process |
| Platynothrus peltifer | GO:0006589 | biological_process | octopamine biosynthetic process |
| Platynothrus peltifer | GO:0007275 | biological_process | multicellular organismal development |
| Platynothrus peltifer | GO:0016266 | biological_process | O-glycan processing |
| Platynothrus peltifer | GO:0052697 | biological_process | xenobiotic glucuronidation |
| Platynothrus peltifer | GO:0007165 | biological_process | signal transduction |
| Platynothrus peltifer | GO:0050661 | molecular_function | NADP binding |
| Platynothrus peltifer | GO:0005524 | molecular_function | ATP binding |
| Platynothrus peltifer | GO:0006706 | biological_process | steroid catabolic process |
| Platynothrus peltifer | GO:0042178 | biological_process | xenobiotic catabolic process |
| Platynothrus peltifer | GO:0045214 | biological_process | sarcomere organization |
| Platynothrus peltifer | GO:0006811 | biological_process | ion transport |
| Platynothrus peltifer | GO:0004497 | molecular_function | monooxygenase activity |
| Platynothrus peltifer | GO:0006511 | biological_process | ubiquitin-dependent protein catabolic process |
| Platynothrus peltifer | GO:0009617 | biological_process | response to bacterium |
| Platynothrus peltifer | GO:0009888 | biological_process | tissue development |
| Platynothrus peltifer | GO:0006417 | biological_process | regulation of translation |
| Platynothrus peltifer | GO:0005525 | molecular_function | GTP binding |
| Platynothrus peltifer | GO:0006749 | biological_process | glutathione metabolic process |
| Platynothrus peltifer | GO:0014832 | biological_process | urinary bladder smooth muscle contraction |
| Platynothrus peltifer | GO:0016139 | biological_process | glycoside catabolic process |
| Platynothrus peltifer | GO:0030162 | biological_process | regulation of proteolysis |
| Platynothrus peltifer | GO:0009058 | biological_process | biosynthetic process |

| | | | |
|---|---|---|---|
| Platynothrus peltifer | GO:0016042 | biological_process | lipid catabolic process |
| Platynothrus peltifer | GO:0010951 | biological_process | negative regulation of endopeptidase activity |
| Platynothrus peltifer | GO:0042302 | molecular_function | structural constituent of cuticle |
| Platynothrus peltifer | GO:0006468 | biological_process | protein phosphorylation |
| Platynothrus peltifer | GO:0010811 | biological_process | positive regulation of cell-substrate adhesion |
| Platynothrus peltifer | GO:0016614 | molecular_function | oxidoreductase activity, acting on CH-OH group of donors |
| Platynothrus peltifer | GO:0006069 | biological_process | ethanol oxidation |
| Platynothrus peltifer | GO:0010842 | biological_process | retina layer formation |
| Platynothrus peltifer | GO:0006486 | biological_process | protein glycosylation |
| Platynothrus peltifer | GO:0030512 | biological_process | negative regulation of transforming growth factor beta receptor signaling pathway |
| Platynothrus peltifer | GO:0051260 | biological_process | protein homooligomerization |
| Platynothrus peltifer | GO:0043252 | biological_process | sodium-independent organic anion transport |
| Platynothrus peltifer | GO:0016055 | biological_process | Wnt signaling pathway |
| Platynothrus peltifer | GO:0016567 | biological_process | protein ubiquitination |
| Platynothrus peltifer | GO:0042135 | biological_process | neurotransmitter catabolic process |
| Platynothrus peltifer | GO:0016491 | molecular_function | oxidoreductase activity |
| Platynothrus peltifer | GO:0000272 | biological_process | polysaccharide catabolic process |
| Platynothrus peltifer | GO:0008270 | molecular_function | zinc ion binding |
| Platynothrus peltifer | GO:0032434 | biological_process | regulation of proteasomal ubiquitin-dependent protein catabolic process |
| Platynothrus peltifer | GO:0006090 | biological_process | pyruvate metabolic process |
| Platynothrus peltifer | GO:0007016 | biological_process | cytoskeletal anchoring at plasma membrane |
| Platynothrus peltifer | GO:0001669 | cellular_component | acrosomal vesicle |
| Platynothrus peltifer | GO:0000723 | biological_process | telomere maintenance |
| Platynothrus peltifer | GO:0006152 | biological_process | purine nucleoside catabolic process |
| Platynothrus peltifer | GO:0001649 | biological_process | osteoblast differentiation |
| Platynothrus peltifer | GO:0015020 | molecular_function | glucuronosyltransferase activity |
| Platynothrus peltifer | GO:0006979 | biological_process | response to oxidative stress |
| Platynothrus peltifer | GO:0031098 | biological_process | stress-activated protein kinase signaling cascade |
| Platynothrus peltifer | GO:0006470 | biological_process | protein dephosphorylation |
| Platynothrus peltifer | GO:0006099 | biological_process | tricarboxylic acid cycle |
| Platynothrus peltifer | GO:0034976 | biological_process | response to endoplasmic reticulum stress |
| Platynothrus peltifer | GO:0046355 | biological_process | mannan catabolic process |
| Platynothrus peltifer | GO:0016192 | biological_process | vesicle-mediated transport |
| Platynothrus peltifer | GO:0004674 | molecular_function | protein serine/threonine kinase activity |
| Platynothrus peltifer | GO:1902531 | biological_process | regulation of intracellular signal transduction |
| Platynothrus peltifer | GO:0008218 | biological_process | bioluminescence |
| Platynothrus peltifer | GO:0031297 | biological_process | replication fork processing |

| | | | |
|---|---|---|---|
| Platynothrus peltifer | GO:0009405 | biological_process | pathogenesis |
| Platynothrus peltifer | GO:0006355 | biological_process | regulation of transcription, DNA-templated |
| Platynothrus peltifer | GO:0051598 | biological_process | meiotic recombination checkpoint |
| Platynothrus peltifer | GO:0004867 | molecular_function | serine-type endopeptidase inhibitor activity |
| Platynothrus peltifer | GO:0021762 | biological_process | substantia nigra development |
| Platynothrus peltifer | GO:0032482 | biological_process | Rab protein signal transduction |
| Platynothrus peltifer | GO:0001507 | biological_process | acetylcholine catabolic process in synaptic cleft |
| Platynothrus peltifer | GO:0005886 | cellular_component | plasma membrane |
| Platynothrus peltifer | GO:0007179 | biological_process | transforming growth factor beta receptor signaling pathway |
| Platynothrus peltifer | GO:0031638 | biological_process | zymogen activation |
| Platynothrus peltifer | GO:0032197 | biological_process | transposition, RNA-mediated |
| Platynothrus peltifer | GO:0046872 | molecular_function | metal ion binding |
| Platynothrus peltifer | GO:0016032 | biological_process | viral process |
| Platynothrus peltifer | GO:0042060 | biological_process | wound healing |
| Platynothrus peltifer | GO:0015074 | biological_process | DNA integration |
| Platynothrus peltifer | GO:0007155 | biological_process | cell adhesion |
| Platynothrus peltifer | GO:0016020 | cellular_component | membrane |
| Platynothrus peltifer | GO:0042572 | biological_process | retinol metabolic process |
| Platynothrus peltifer | GO:0008354 | biological_process | germ cell migration |
| Platynothrus peltifer | GO:0048010 | biological_process | vascular endothelial growth factor receptor signaling pathway |
| Platynothrus peltifer | GO:0007584 | biological_process | response to nutrient |
| Platynothrus peltifer | GO:0016021 | cellular_component | integral component of membrane |
| Platynothrus peltifer | GO:0042761 | biological_process | very long-chain fatty acid biosynthetic process |
| Platynothrus peltifer | GO:0051131 | biological_process | chaperone-mediated protein complex assembly |
| Platynothrus peltifer | GO:0036323 | biological_process | vascular endothelial growth factor receptor-1 signaling pathway |
| Platynothrus peltifer | GO:0045087 | biological_process | innate immune response |
| Platynothrus peltifer | GO:0030449 | biological_process | regulation of complement activation |
| Platynothrus peltifer | GO:0007286 | biological_process | spermatid development |
| Platynothrus peltifer | GO:0007608 | biological_process | sensory perception of smell |
| Platynothrus peltifer | GO:0006986 | biological_process | response to unfolded protein |
| Platynothrus peltifer | GO:0031149 | biological_process | sorocarp stalk cell differentiation |
| Platynothrus peltifer | GO:0009742 | biological_process | brassinosteroid mediated signaling pathway |
| Platynothrus peltifer | GO:0001505 | biological_process | regulation of neurotransmitter levels |
| Platynothrus peltifer | GO:0006667 | biological_process | sphinganine metabolic process |
| Platynothrus peltifer | GO:0006364 | biological_process | rRNA processing |
| Platynothrus peltifer | GO:0030433 | biological_process | ER-associated ubiquitin-dependent protein catabolic process |
| Platynothrus peltifer | GO:0006325 | biological_process | chromatin organization |

| | | | |
|---|---|---|---|
| Platynothrus peltifer | GO:0006890 | biological_process | retrograde vesicle-mediated transport, Golgi to ER |
| Platynothrus peltifer | GO:0045892 | biological_process | negative regulation of transcription, DNA-templated |
| Platynothrus peltifer | GO:0010224 | biological_process | response to UV-B |
| Platynothrus peltifer | GO:0006644 | biological_process | phospholipid metabolic process |
| Platynothrus peltifer | GO:0006641 | biological_process | triglyceride metabolic process |
| Platynothrus peltifer | GO:0006915 | biological_process | apoptotic process |
| Platynothrus peltifer | GO:0018279 | biological_process | protein N-linked glycosylation via asparagine |
| Platynothrus peltifer | GO:0016485 | biological_process | protein processing |
| Platynothrus peltifer | GO:0035556 | biological_process | intracellular signal transduction |
| Platynothrus peltifer | GO:0004104 | molecular_function | cholinesterase activity |
| Platynothrus peltifer | GO:0000011 | biological_process | vacuole inheritance |
| Platynothrus peltifer | GO:0005737 | cellular_component | cytoplasm |
| Platynothrus peltifer | GO:0006412 | biological_process | translation |
| Platynothrus peltifer | GO:0015031 | biological_process | protein transport |
| Platynothrus peltifer | GO:0006357 | biological_process | regulation of transcription from RNA polymerase II promoter |
| Platynothrus peltifer | GO:0005576 | cellular_component | extracellular region |
| Platynothrus peltifer | GO:0000122 | biological_process | negative regulation of transcription from RNA polymerase II promoter |
| Platynothrus peltifer | GO:0007264 | biological_process | small GTPase mediated signal transduction |
| Platynothrus peltifer | GO:0006260 | biological_process | DNA replication |
| Platynothrus peltifer | GO:0035220 | biological_process | wing disc development |
| Platynothrus peltifer | GO:0008168 | molecular_function | methyltransferase activity |
| Platynothrus peltifer | GO:0004930 | molecular_function | G-protein coupled receptor activity |
| Platynothrus peltifer | GO:0001894 | biological_process | tissue homeostasis |
| Platynothrus peltifer | GO:2000377 | biological_process | regulation of reactive oxygen species metabolic process |
| Platynothrus peltifer | GO:0042981 | biological_process | regulation of apoptotic process |
| Platynothrus peltifer | GO:0035725 | biological_process | sodium ion transmembrane transport |
| Platynothrus peltifer | GO:0051290 | biological_process | protein heterotetramerization |
| Platynothrus peltifer | GO:0001516 | biological_process | prostaglandin biosynthetic process |
| Platynothrus peltifer | GO:0008289 | molecular_function | lipid binding |
| Platynothrus peltifer | GO:0097264 | biological_process | self proteolysis |
| Platynothrus peltifer | GO:1902542 | biological_process | regulation of protein localization to mitotic spindle pole body |
| Platynothrus peltifer | GO:0071357 | biological_process | cellular response to type I interferon |
| Platynothrus peltifer | GO:0042742 | biological_process | defense response to bacterium |
| Platynothrus peltifer | GO:0051124 | biological_process | synaptic growth at neuromuscular junction |

| | |
|---|---|
| 218 | 0 |
| 99 | 0,000000702552 |
| 78 | 0 |
| 78 | 0,00088813166 |
| 73 | 0,000002100107 |
| 65 | 0,004151710965 |
| 61 | 0 |
| 58 | 0 |
| 54 | 0,003930555578 |
| 53 | 0 |
| 47 | 0 |
| 43 | 0 |
| 43 | 0,000000000223 |
| 43 | 0,000439829013 |
| 41 | 0 |
| 37 | 0 |
| 37 | 0,000000000195 |
| 36 | 0 |
| 36 | 0 |
| 35 | 0 |
| 35 | 0 |
| 33 | 0,000268579647 |
| 33 | 0,01791843976 |
| 32 | 0,000000000394 |
| 32 | 0,01995969949 |
| 30 | 0,001227723725 |
| 28 | 0,009819127696 |
| 27 | 0 |
| 27 | 0,000000009085 |
| 27 | 0,000217306754 |
| 27 | 0,003254981223 |
| 26 | 0,000093962817 |
| 26 | 0,01571339667 |
| 25 | 0,000000000109 |
| 25 | 0,000000096168 |
| 25 | 0,000075352012 |
| 24 | 0,000255570177 |

| | |
|---|---|
| 23 | 0,000028892799 |
| 23 | 0,000084695981 |
| 23 | 0,001234007684 |
| 22 | 0,000000000857 |
| 22 | 0,000000251431 |
| 20 | 0,006193044043 |
| 19 | 0,000901222606 |
| 19 | 0,006069579 |
| 18 | 0 |
| 18 | 0,000000001418 |
| 18 | 0,000243407834 |
| 18 | 0,01799699207 |
| 17 | 0 |
| 17 | 0 |
| 17 | 0,01066902919 |
| 16 | 0,000000000702 |
| 16 | 0,000002482220 |
| 15 | 0 |
| 15 | 0,000003422684 |
| 15 | 0,000009698827 |
| 14 | 0,000008990866 |
| 14 | 0,00114336931 |
| 14 | 0,005550211746 |
| 13 | 0,000000006297 |
| 13 | 0,000023461218 |
| 13 | 0,002207724313 |
| 12 | 0 |
| 11 | 0,000000017004 |
| 11 | 0,000000921567 |
| 11 | 0,000001285829 |
| 11 | 0,000005580657 |
| 11 | 0,0237560455 |
| 10 | 0 |
| 10 | 0,005996555892 |
| 9 | 0,000003958766 |
| 9 | 0,003035574537 |
| 9 | 0,005012828914 |

| | |
|---|---|
| 9 | 0,0114137783 |
| 8 | 0 |
| 8 | 0,000000014687 |
| 8 | 0,002747451217 |
| 8 | 0,005062853814 |
| 8 | 0,00676709302 |
| 7 | 0,000001836124 |
| 7 | 0,000029358686 |
| 7 | 0,000109458022 |
| 7 | 0,000708848701[illegible] |
| 7 | 0,01173433809 |
| 6 | 0 |
| 6 | 0 |
| 6 | 0,000000000107 |
| 6 | 0,000009416908 |
| 6 | 0,000041371584 |
| 6 | 0,000053421534 |
| 6 | 0,000088594284 |
| 6 | 0,000359999139 |
| 6 | 0,006911200593 |
| 6 | 0,01096994995 |
| 5 | 0 |
| 5 | 0 |
| 5 | 0,003078083097 |
| 5 | 0,004724047531 |
| 5 | 0,005289318516 |
| 5 | 0,00687234514 |
| 4 | 0,000000001247 |
| 4 | 0,000083611604[illegible] |
| 4 | 0,000485645199 |
| 4 | 0,000552268585 |
| 4 | 0,000552268585 |
| 4 | 0,006023719044 |
| 4 | 0,01332854164 |
| 3 | 0 |
| 3 | 0 |
| 3 | 0 |

| | |
|---|---|
| 3 | 0 |
| 3 | 0,000000001410 |
| 3 | 0,000004458535 |
| 3 | 0,000008049560 |
| 3 | 0,000028430993 |
| 3 | 0,000107266713 |
| 3 | 0,000117777954 |
| 3 | 0,000668695685 |
| 3 | 0,002930808427 |
| 3 | 0,00726053087 |
| 3 | 0,009407895351 |
| 3 | 0,01471279598 |
| 2 | 0 |
| 2 | 0 |
| 2 | 0 |
| 2 | 0 |
| 2 | 0,000000000234 |
| 2 | 0,000000001103 |
| 2 | 0,000000417107 |
| 2 | 0,000000783986 |
| 2 | 0,000042865831 |
| 2 | 0,000229344312 |
| 2 | 0,000371922126 |
| 2 | 0,000450640742 |
| 2 | 0,000725539532 |
| 2 | 0,002221076597 |
| 2 | 0,002666452466 |
| 2 | 0,003499660591 |
| 2 | 0,00924527341 |
| 2 | 0,00924527341 |
| 2 | 0,01115750717 |
| 2 | 0,0141579097 |
| 2 | 0,0141579097 |
| 2 | 0,02143332077 |

| | | | | |
|---|---|---|---|---|
| Nothrus palustris | GO:0055085 | biological_process | transmembrane transport | 388 |
| Nothrus palustris | GO:0042908 | biological_process | xenobiotic transport | 96 |
| Nothrus palustris | GO:0007283 | biological_process | spermatogenesis | 83 |
| Nothrus palustris | GO:0016020 | cellular_component | membrane | 74 |
| Nothrus palustris | GO:0008218 | biological_process | bioluminescence | 70 |
| Nothrus palustris | GO:0015074 | biological_process | DNA integration | 69 |
| Nothrus palustris | GO:0003677 | molecular_function | DNA binding | 66 |
| Nothrus palustris | GO:0003081 | biological_process | regulation of systemic arterial blood pressure by renin-angiotensin | 62 |
| Nothrus palustris | GO:0032197 | biological_process | transposition, RNA-mediated | 61 |
| Nothrus palustris | GO:0006313 | biological_process | transposition, DNA-mediated | 57 |
| Nothrus palustris | GO:0009617 | biological_process | response to bacterium | 51 |
| Nothrus palustris | GO:0045214 | biological_process | sarcomere organization | 50 |
| Nothrus palustris | GO:0006680 | biological_process | glucosylceramide catabolic process | 37 |
| Nothrus palustris | GO:0010224 | biological_process | response to UV-B | 37 |
| Nothrus palustris | GO:0006486 | biological_process | protein glycosylation | 37 |
| Nothrus palustris | GO:0009311 | biological_process | oligosaccharide metabolic process | 36 |
| Nothrus palustris | GO:0006805 | biological_process | xenobiotic metabolic process | 35 |
| Nothrus palustris | GO:0007601 | biological_process | visual perception | 34 |
| Nothrus palustris | GO:0006260 | biological_process | DNA replication | 34 |
| Nothrus palustris | GO:0005975 | biological_process | carbohydrate metabolic process | 32 |
| Nothrus palustris | GO:1902358 | biological_process | sulfate transmembrane transport | 30 |
| Nothrus palustris | GO:0019284 | biological_process | L-methionine biosynthetic process from S-adenosylmethionine | 29 |
| Nothrus palustris | GO:0006915 | biological_process | apoptotic process | 26 |
| Nothrus palustris | GO:0010992 | biological_process | ubiquitin homeostasis | 25 |
| Nothrus palustris | GO:0051384 | biological_process | response to glucocorticoid | 24 |
| Nothrus palustris | GO:0001516 | biological_process | prostaglandin biosynthetic process | 24 |
| Nothrus palustris | GO:0016266 | biological_process | O-glycan processing | 24 |
| Nothrus palustris | GO:0007165 | biological_process | signal transduction | 23 |
| Nothrus palustris | GO:0008270 | molecular_function | zinc ion binding | 23 |
| Nothrus palustris | GO:0006468 | biological_process | protein phosphorylation | 23 |
| Nothrus palustris | GO:0045599 | biological_process | negative regulation of fat cell differentiation | 23 |
| Nothrus palustris | GO:0009636 | biological_process | response to toxic substance | 22 |
| Nothrus palustris | GO:0016042 | biological_process | lipid catabolic process | 20 |
| Nothrus palustris | GO:0004721 | molecular_function | phosphoprotein phosphatase activity | 20 |
| Nothrus palustris | GO:0042302 | molecular_function | structural constituent of cuticle | 20 |
| Nothrus palustris | GO:0007169 | biological_process | transmembrane receptor protein tyrosine kinase signaling pathway | 20 |
| Nothrus palustris | GO:0008210 | biological_process | estrogen metabolic process | 19 |

| | | | | |
|---|---|---|---|---|
| Nothrus palustris | GO:0010025 | biological_process | wax biosynthetic process | 19 |
| Nothrus palustris | GO:0007275 | biological_process | multicellular organismal development | 18 |
| Nothrus palustris | GO:0033609 | biological_process | oxalate metabolic process | 18 |
| Nothrus palustris | GO:0009698 | biological_process | phenylpropanoid metabolic process | 18 |
| Nothrus palustris | GO:0016192 | biological_process | vesicle-mediated transport | 17 |
| Nothrus palustris | GO:0008146 | molecular_function | sulfotransferase activity | 17 |
| Nothrus palustris | GO:0051923 | biological_process | sulfation | 17 |
| Nothrus palustris | GO:0006310 | biological_process | DNA recombination | 17 |
| Nothrus palustris | GO:0010842 | biological_process | retina layer formation | 17 |
| Nothrus palustris | GO:0006728 | biological_process | pteridine biosynthetic process | 16 |
| Nothrus palustris | GO:0031030 | biological_process | negative regulation of septation initiation signaling | 16 |
| Nothrus palustris | GO:0007605 | biological_process | sensory perception of sound | 16 |
| Nothrus palustris | GO:0002376 | biological_process | immune system process | 16 |
| Nothrus palustris | GO:0004252 | molecular_function | serine-type endopeptidase activity | 15 |
| Nothrus palustris | GO:0009263 | biological_process | deoxyribonucleotide biosynthetic process | 15 |
| Nothrus palustris | GO:0006810 | biological_process | transport | 15 |
| Nothrus palustris | GO:0034976 | biological_process | response to endoplasmic reticulum stress | 15 |
| Nothrus palustris | GO:0004983 | molecular_function | neuropeptide Y receptor activity | 15 |
| Nothrus palustris | GO:0006511 | biological_process | ubiquitin-dependent protein catabolic process | 14 |
| Nothrus palustris | GO:0043181 | biological_process | vacuolar sequestering | 14 |
| Nothrus palustris | GO:0007016 | biological_process | cytoskeletal anchoring at plasma membrane | 14 |
| Nothrus palustris | GO:0042135 | biological_process | neurotransmitter catabolic process | 14 |
| Nothrus palustris | GO:0007021 | biological_process | tubulin complex assembly | 14 |
| Nothrus palustris | GO:0006633 | biological_process | fatty acid biosynthetic process | 13 |
| Nothrus palustris | GO:0033089 | biological_process | positive regulation of T cell differentiation in thymus | 13 |
| Nothrus palustris | GO:0005634 | cellular_component | nucleus | 13 |
| Nothrus palustris | GO:0006941 | biological_process | striated muscle contraction | 13 |
| Nothrus palustris | GO:0034612 | biological_process | response to tumor necrosis factor | 13 |
| Nothrus palustris | GO:0019233 | biological_process | sensory perception of pain | 13 |
| Nothrus palustris | GO:0005980 | biological_process | glycogen catabolic process | 13 |
| Nothrus palustris | GO:0046355 | biological_process | mannan catabolic process | 13 |
| Nothrus palustris | GO:0004518 | molecular_function | nuclease activity | 13 |
| Nothrus palustris | GO:0006355 | biological_process | regulation of transcription, DNA-templated | 12 |
| Nothrus palustris | GO:0045944 | biological_process | positive regulation of transcription from RNA polymerase II promoter | 12 |
| Nothrus palustris | GO:0000272 | biological_process | polysaccharide catabolic process | 12 |
| Nothrus palustris | GO:0031098 | biological_process | stress-activated protein kinase signaling cascade | 12 |
| Nothrus palustris | GO:0009099 | biological_process | valine biosynthetic process | 12 |

| | | | | |
|---|---|---|---|---|
| Nothrus palustris | GO:0006470 | biological_process | protein dephosphorylation | 12 |
| Nothrus palustris | GO:0019882 | biological_process | antigen processing and presentation | 12 |
| Nothrus palustris | GO:0016114 | biological_process | terpenoid biosynthetic process | 12 |
| Nothrus palustris | GO:0006412 | biological_process | translation | 11 |
| Nothrus palustris | GO:0016055 | biological_process | Wnt signaling pathway | 11 |
| Nothrus palustris | GO:0008277 | biological_process | regulation of G-protein coupled receptor protein signaling pathway | 11 |
| Nothrus palustris | GO:0055114 | biological_process | oxidation-reduction process | 11 |
| Nothrus palustris | GO:0070633 | biological_process | transepithelial transport | 11 |
| Nothrus palustris | GO:0004674 | molecular_function | protein serine/threonine kinase activity | 11 |
| Nothrus palustris | GO:0016021 | cellular_component | integral component of membrane | 10 |
| Nothrus palustris | GO:0006508 | biological_process | proteolysis | 10 |
| Nothrus palustris | GO:0016567 | biological_process | protein ubiquitination | 10 |
| Nothrus palustris | GO:0008380 | biological_process | RNA splicing | 10 |
| Nothrus palustris | GO:0010869 | biological_process | regulation of receptor biosynthetic process | 10 |
| Nothrus palustris | GO:0055069 | biological_process | zinc ion homeostasis | 10 |
| Nothrus palustris | GO:0016488 | biological_process | farnesol catabolic process | 10 |
| Nothrus palustris | GO:0046872 | molecular_function | metal ion binding | 9 |
| Nothrus palustris | GO:0006979 | biological_process | response to oxidative stress | 9 |
| Nothrus palustris | GO:0019062 | biological_process | virion attachment to host cell | 9 |
| Nothrus palustris | GO:0051603 | biological_process | proteolysis involved in cellular protein catabolic process | 9 |
| Nothrus palustris | GO:0042572 | biological_process | retinol metabolic process | 9 |
| Nothrus palustris | GO:0047936 | molecular_function | glucose 1-dehydrogenase [NAD(P)] activity | 9 |
| Nothrus palustris | GO:0004222 | molecular_function | metalloendopeptidase activity | 9 |
| Nothrus palustris | GO:0006099 | biological_process | tricarboxylic acid cycle | 8 |
| Nothrus palustris | GO:0016614 | molecular_function | oxidoreductase activity, acting on CH-OH group of donors | 8 |
| Nothrus palustris | GO:0008241 | molecular_function | peptidyl-dipeptidase activity | 8 |
| Nothrus palustris | GO:0003964 | molecular_function | RNA-directed DNA polymerase activity | 8 |
| Nothrus palustris | GO:0047935 | molecular_function | glucose 1-dehydrogenase (NADP+) activity | 8 |
| Nothrus palustris | GO:0016032 | biological_process | viral process | 7 |
| Nothrus palustris | GO:0008202 | biological_process | steroid metabolic process | 7 |
| Nothrus palustris | GO:0048364 | biological_process | root development | 7 |
| Nothrus palustris | GO:0005739 | cellular_component | mitochondrion | 7 |
| Nothrus palustris | GO:0048794 | biological_process | swim bladder development | 6 |
| Nothrus palustris | GO:0005886 | cellular_component | plasma membrane | 6 |
| Nothrus palustris | GO:0007586 | biological_process | digestion | 6 |
| Nothrus palustris | GO:0015939 | biological_process | pantothenate metabolic process | 6 |
| Nothrus palustris | GO:0031638 | biological_process | zymogen activation | 6 |

| | | | | |
|---|---|---|---|---|
| Nothrus palustris | GO:0009405 | biological_process | pathogenesis | 6 |
| Nothrus palustris | GO:0007608 | biological_process | sensory perception of smell | 6 |
| Nothrus palustris | GO:0006641 | biological_process | triglyceride metabolic process | 6 |
| Nothrus palustris | GO:0018279 | biological_process | protein N-linked glycosylation via asparagine | 6 |
| Nothrus palustris | GO:0015031 | biological_process | protein transport | 5 |
| Nothrus palustris | GO:0015893 | biological_process | drug transport | 5 |
| Nothrus palustris | GO:0036372 | biological_process | opsin transport | 5 |
| Nothrus palustris | GO:0046693 | biological_process | sperm storage | 5 |
| Nothrus palustris | GO:0050951 | biological_process | sensory perception of temperature stimulus | 5 |
| Nothrus palustris | GO:0018394 | biological_process | peptidyl-lysine acetylation | 5 |
| Nothrus palustris | GO:0030497 | biological_process | fatty acid elongation | 5 |
| Nothrus palustris | GO:0005525 | molecular_function | GTP binding | 5 |
| Nothrus palustris | GO:0001894 | biological_process | tissue homeostasis | 5 |
| Nothrus palustris | GO:0030162 | biological_process | regulation of proteolysis | 5 |
| Nothrus palustris | GO:0035556 | biological_process | intracellular signal transduction | 5 |
| Nothrus palustris | GO:0006357 | biological_process | regulation of transcription from RNA polymerase II promoter | 4 |
| Nothrus palustris | GO:0006694 | biological_process | steroid biosynthetic process | 4 |
| Nothrus palustris | GO:0004497 | molecular_function | monooxygenase activity | 4 |
| Nothrus palustris | GO:0006629 | biological_process | lipid metabolic process | 4 |
| Nothrus palustris | GO:0008542 | biological_process | visual learning | 4 |
| Nothrus palustris | GO:0051301 | biological_process | cell division | 4 |
| Nothrus palustris | GO:0031149 | biological_process | sorocarp stalk cell differentiation | 4 |
| Nothrus palustris | GO:0009742 | biological_process | brassinosteroid mediated signaling pathway | 4 |
| Nothrus palustris | GO:0007416 | biological_process | synapse assembly | 4 |
| Nothrus palustris | GO:2000504 | biological_process | positive regulation of blood vessel remodeling | 4 |
| Nothrus palustris | GO:0008654 | biological_process | phospholipid biosynthetic process | 4 |
| Nothrus palustris | GO:0009213 | biological_process | pyrimidine deoxyribonucleoside triphosphate catabolic process | 4 |
| Nothrus palustris | GO:0001932 | biological_process | regulation of protein phosphorylation | 4 |
| Nothrus palustris | GO:0006904 | biological_process | vesicle docking involved in exocytosis | 4 |
| Nothrus palustris | GO:0005576 | cellular_component | extracellular region | 3 |
| Nothrus palustris | GO:0005506 | molecular_function | iron ion binding | 3 |
| Nothrus palustris | GO:0071625 | biological_process | vocalization behavior | 3 |
| Nothrus palustris | GO:0015918 | biological_process | sterol transport | 3 |
| Nothrus palustris | GO:0006401 | biological_process | RNA catabolic process | 3 |
| Nothrus palustris | GO:0001505 | biological_process | regulation of neurotransmitter levels | 3 |
| Nothrus palustris | GO:0048489 | biological_process | synaptic vesicle transport | 3 |
| Nothrus palustris | GO:0031175 | biological_process | neuron projection development | 3 |

| | | | | |
|---|---|---|---|---|
| Nothrus palustris | GO:0044826 | biological_process | viral genome integration into host DNA | 3 |
| Nothrus palustris | GO:0051225 | biological_process | spindle assembly | 3 |
| Nothrus palustris | GO:0042626 | molecular_function | ATPase activity, coupled to transmembrane movement of substances | 2 |
| Nothrus palustris | GO:0070328 | biological_process | triglyceride homeostasis | 2 |
| Nothrus palustris | GO:0000723 | biological_process | telomere maintenance | 2 |
| Nothrus palustris | GO:0007186 | biological_process | G-protein coupled receptor signaling pathway | 2 |
| Nothrus palustris | GO:0007179 | biological_process | transforming growth factor beta receptor signaling pathway | 2 |
| Nothrus palustris | GO:0007155 | biological_process | cell adhesion | 2 |
| Nothrus palustris | GO:0006644 | biological_process | phospholipid metabolic process | 2 |
| Nothrus palustris | GO:0071555 | biological_process | cell wall organization | 2 |
| Nothrus palustris | GO:0004930 | molecular_function | G-protein coupled receptor activity | 2 |
| Nothrus palustris | GO:0032465 | biological_process | regulation of cytokinesis | 2 |
| Nothrus palustris | GO:0001501 | biological_process | skeletal system development | 2 |
| Nothrus palustris | GO:0042981 | biological_process | regulation of apoptotic process | 2 |
| Nothrus palustris | GO:0008234 | molecular_function | cysteine-type peptidase activity | 2 |
| Nothrus palustris | GO:0009268 | biological_process | response to pH | 2 |
| Nothrus palustris | GO:0001558 | biological_process | regulation of cell growth | 2 |
| Nothrus palustris | GO:0002532 | biological_process | production of molecular mediator involved in inflammatory response | 2 |
| Nothrus palustris | GO:0071622 | biological_process | regulation of granulocyte chemotaxis | 2 |
| Nothrus palustris | GO:0097090 | biological_process | presynaptic membrane organization | 2 |
| Nothrus palustris | GO:0008063 | biological_process | Toll signaling pathway | 2 |
| Nothrus palustris | GO:0043434 | biological_process | response to peptide hormone | 2 |

| |
|---|
| 0 |
| 0 |
| 0 |
| 0 |
| 0 |
| 0 |
| 0,000006552920 |
| 0 |
| 0 |
| 0,001346753444 |
| 0,01237070939 |
| 0,000000177363 |
| 0,000001493702 |
| 0,000009880938 |
| 0,000046011099 |
| 0 |
| 0,008854079821 |
| 0 |
| 0,000187500648 |
| 0,000008124776 |
| 0 |
| 0,004154387236 |
| 0,001460421212 |
| 0 |
| 0,000002107720 |
| 0,000003307965 |
| 0,000079715988 |
| 0 |
| 0 |
| 0,000000001684 |
| 0,000009065284 |
| 0,001972050811 |
| 0,000001640179 |
| 0,000033371808 |
| 0,000137365824 |
| 0,0105290673 |
| 0 |

| |
|---|
| 0,01958293319 |
| 0 |
| 0 |
| 0,000069211397 |
| 0 |
| 0,000000014478 |
| 0,001180453045 |
| 0,00335283137 |
| 0,02205494362 |
| 0,000000000862 |
| 0,000000040114 |
| 0,000638351287 |
| 0,008854877661 |
| 0 |
| 0,000000494116 |
| 0,000048035948 |
| 0,000214765620 |
| 0,01331814109 |
| 0 |
| 0,000000000412 |
| 0,000010070928 |
| 0,001585740316 |
| 0,01812771034 |
| 0,000000001046 |
| 0,000000002779 |
| 0,000000482437 |
| 0,000012602940 |
| 0,000018276789 |
| 0,000533258322 |
| 0,000660009755 |
| 0,005027736471 |
| 0,02260859256 |
| 0 |
| 0 |
| 0,000000017571 |
| 0,000000046600 |
| 0,000001374335 |

0,000002295136
0,000224314952
0,004744041478
0
0
0,000231941373
0,000623561929
0,005931206415
0,01034178035
0
0
0
0
0,000005973163
0,000018280601
0,000047521734
0
0
0,000000014941
0,000009730833
0,001696298497
0,001820008209
0,01908401763
0,000000016704
0,000000082672
0,000528827610
0,002448514566
0,01602033176
0
0,000390902740
0,000635971754
0,003304400468
0,000001435371
0,000006489344
0,000006489344
0,00019303789
0,000205962649

| |
|---|
| 0,000804979111[illegible] |
| 0,000870117799[illegible] |
| 0,001278439226 |
| 0,003869884053 |
| 0 |
| 0,000013589615 |
| 0,000013589615 |
| 0,000058137663 |
| 0,000072925460 |
| 0,000072925460 |
| 0,006470000362 |
| 0,006952448903 |
| 0,01217089543 |
| 0,01217089543 |
| 0,01938357595 |
| 0 |
| 0,000001797947[illegible] |
| 0,000008432637 |
| 0,000031538024 |
| 0,000072297547 |
| 0,000469364056 |
| 0,00057393363 |
| 0,00057393363 |
| 0,000930960299 |
| 0,003211681918 |
| 0,00554306206 |
| 0,005742865511 |
| 0,009241739923 |
| 0,0180493121 |
| 0 |
| 0,000000012698 |
| 0,000085383675 |
| 0,000288647378 |
| 0,001027124967 |
| 0,001741969522 |
| 0,004099309034 |
| 0,01276501288 |

| |
|---|
| 0,01728493592 |
| 0,01737699993 |
| 0,000000000436 |
| 0,000000000436 |
| 0,000000000952 |
| 0,000000001329 |
| 0,000000025923 |
| 0,000000035927 |
| 0,000000997689 |
| 0,000204645305 |
| 0,000204645305 |
| 0,000249055080 |
| 0,000249055080 |
| 0,000656887433 |
| 0,000874973705 |
| 0,003543099564 |
| 0,006082357311 |
| 0,007262202394 |
| 0,01138037357 |
| 0,01138037357 |
| 0,01719246922 |
| 0,01869242262 |

| | | | |
|---|---|---|---|
| Hermannia gibba | GO:0055085 | biological_process | transmembrane transport |
| Hermannia gibba | GO:0032100 | biological_process | positive regulation of appetite |
| Hermannia gibba | GO:0003677 | molecular_function | DNA binding |
| Hermannia gibba | GO:0006517 | biological_process | protein deglycosylation |
| Hermannia gibba | GO:0042908 | biological_process | xenobiotic transport |
| Hermannia gibba | GO:0006633 | biological_process | fatty acid biosynthetic process |
| Hermannia gibba | GO:0006313 | biological_process | transposition, DNA-mediated |
| Hermannia gibba | GO:0000272 | biological_process | polysaccharide catabolic process |
| Hermannia gibba | GO:0007601 | biological_process | visual perception |
| Hermannia gibba | GO:0030223 | biological_process | neutrophil differentiation |
| Hermannia gibba | GO:0006706 | biological_process | steroid catabolic process |
| Hermannia gibba | GO:0015074 | biological_process | DNA integration |
| Hermannia gibba | GO:0006680 | biological_process | glucosylceramide catabolic process |
| Hermannia gibba | GO:0008270 | molecular_function | zinc ion binding |
| Hermannia gibba | GO:0004364 | molecular_function | glutathione transferase activity |
| Hermannia gibba | GO:0005975 | biological_process | carbohydrate metabolic process |
| Hermannia gibba | GO:0006814 | biological_process | sodium ion transport |
| Hermannia gibba | GO:0006511 | biological_process | ubiquitin-dependent protein catabolic process |
| Hermannia gibba | GO:0004867 | molecular_function | serine-type endopeptidase inhibitor activity |
| Hermannia gibba | GO:0006486 | biological_process | protein glycosylation |
| Hermannia gibba | GO:0016491 | molecular_function | oxidoreductase activity |
| Hermannia gibba | GO:0003964 | molecular_function | RNA-directed DNA polymerase activity |
| Hermannia gibba | GO:0006334 | biological_process | nucleosome assembly |
| Hermannia gibba | GO:0055129 | biological_process | L-proline biosynthetic process |
| Hermannia gibba | GO:0052697 | biological_process | xenobiotic glucuronidation |
| Hermannia gibba | GO:0030435 | biological_process | sporulation resulting in formation of a cellular spore |
| Hermannia gibba | GO:0046982 | molecular_function | protein heterodimerization activity |
| Hermannia gibba | GO:0019542 | biological_process | propionate biosynthetic process |
| Hermannia gibba | GO:0006013 | biological_process | mannose metabolic process |
| Hermannia gibba | GO:0010224 | biological_process | response to UV-B |
| Hermannia gibba | GO:0010842 | biological_process | retina layer formation |
| Hermannia gibba | GO:0009617 | biological_process | response to bacterium |
| Hermannia gibba | GO:0016042 | biological_process | lipid catabolic process |
| Hermannia gibba | GO:1902358 | biological_process | sulfate transmembrane transport |
| Hermannia gibba | GO:0042302 | molecular_function | structural constituent of cuticle |
| Hermannia gibba | GO:0004497 | molecular_function | monooxygenase activity |
| Hermannia gibba | GO:0010383 | biological_process | cell wall polysaccharide metabolic process |

| | | | |
|---|---|---|---|
| Hermannia gibba | GO:0004252 | molecular_function | serine-type endopeptidase activity |
| Hermannia gibba | GO:0031297 | biological_process | replication fork processing |
| Hermannia gibba | GO:0071357 | biological_process | cellular response to type I interferon |
| Hermannia gibba | GO:0042178 | biological_process | xenobiotic catabolic process |
| Hermannia gibba | GO:0004222 | molecular_function | metalloendopeptidase activity |
| Hermannia gibba | GO:0051260 | biological_process | protein homooligomerization |
| Hermannia gibba | GO:0006508 | biological_process | proteolysis |
| Hermannia gibba | GO:0043171 | biological_process | peptide catabolic process |
| Hermannia gibba | GO:0010811 | biological_process | positive regulation of cell-substrate adhesion |
| Hermannia gibba | GO:0016567 | biological_process | protein ubiquitination |
| Hermannia gibba | GO:0009888 | biological_process | tissue development |
| Hermannia gibba | GO:0010992 | biological_process | ubiquitin homeostasis |
| Hermannia gibba | GO:0031048 | biological_process | chromatin silencing by small RNA |
| Hermannia gibba | GO:0006069 | biological_process | ethanol oxidation |
| Hermannia gibba | GO:0010951 | biological_process | negative regulation of endopeptidase activity |
| Hermannia gibba | GO:0016266 | biological_process | O-glycan processing |
| Hermannia gibba | GO:0045616 | biological_process | regulation of keratinocyte differentiation |
| Hermannia gibba | GO:0042135 | biological_process | neurotransmitter catabolic process |
| Hermannia gibba | GO:0006979 | biological_process | response to oxidative stress |
| Hermannia gibba | GO:0004518 | molecular_function | nuclease activity |
| Hermannia gibba | GO:0003081 | biological_process | regulation of systemic arterial blood pressure by renin-angiotensin |
| Hermannia gibba | GO:0051384 | biological_process | response to glucocorticoid |
| Hermannia gibba | GO:0007169 | biological_process | transmembrane receptor protein tyrosine kinase signaling pathway |
| Hermannia gibba | GO:0007165 | biological_process | signal transduction |
| Hermannia gibba | GO:0007275 | biological_process | multicellular organismal development |
| Hermannia gibba | GO:0010025 | biological_process | wax biosynthetic process |
| Hermannia gibba | GO:0007283 | biological_process | spermatogenesis |
| Hermannia gibba | GO:0015020 | molecular_function | glucuronosyltransferase activity |
| Hermannia gibba | GO:0045214 | biological_process | sarcomere organization |
| Hermannia gibba | GO:0033609 | biological_process | oxalate metabolic process |
| Hermannia gibba | GO:0030512 | biological_process | negative regulation of transforming growth factor beta receptor signaling pathway |
| Hermannia gibba | GO:0023052 | biological_process | signaling |
| Hermannia gibba | GO:0030245 | biological_process | cellulose catabolic process |
| Hermannia gibba | GO:0006355 | biological_process | regulation of transcription, DNA-templated |
| Hermannia gibba | GO:0044179 | biological_process | hemolysis in other organism |
| Hermannia gibba | GO:0070189 | biological_process | kynurenine metabolic process |
| Hermannia gibba | GO:0006310 | biological_process | DNA recombination |

| | | | |
|---|---|---|---|
| Hermannia gibba | GO:0016055 | biological_process | Wnt signaling pathway |
| Hermannia gibba | GO:0046373 | biological_process | L-arabinose metabolic process |
| Hermannia gibba | GO:0006694 | biological_process | steroid biosynthetic process |
| Hermannia gibba | GO:0006470 | biological_process | protein dephosphorylation |
| Hermannia gibba | GO:0006810 | biological_process | transport |
| Hermannia gibba | GO:0007179 | biological_process | transforming growth factor beta receptor signaling pathway |
| Hermannia gibba | GO:0042742 | biological_process | defense response to bacterium |
| Hermannia gibba | GO:0046872 | molecular_function | metal ion binding |
| Hermannia gibba | GO:0051603 | biological_process | proteolysis involved in cellular protein catabolic process |
| Hermannia gibba | GO:0042761 | biological_process | very long-chain fatty acid biosynthetic process |
| Hermannia gibba | GO:0033089 | biological_process | positive regulation of T cell differentiation in thymus |
| Hermannia gibba | GO:0010193 | biological_process | response to ozone |
| Hermannia gibba | GO:0042572 | biological_process | retinol metabolic process |
| Hermannia gibba | GO:0006412 | biological_process | translation |
| Hermannia gibba | GO:0009636 | biological_process | response to toxic substance |
| Hermannia gibba | GO:0042060 | biological_process | wound healing |
| Hermannia gibba | GO:0034976 | biological_process | response to endoplasmic reticulum stress |
| Hermannia gibba | GO:0035092 | biological_process | sperm chromatin condensation |
| Hermannia gibba | GO:0009099 | biological_process | valine biosynthetic process |
| Hermannia gibba | GO:0055114 | biological_process | oxidation-reduction process |
| Hermannia gibba | GO:0000209 | biological_process | protein polyubiquitination |
| Hermannia gibba | GO:0035249 | biological_process | synaptic transmission, glutamatergic |
| Hermannia gibba | GO:0008277 | biological_process | regulation of G-protein coupled receptor protein signaling pathway |
| Hermannia gibba | GO:0031098 | biological_process | stress-activated protein kinase signaling cascade |
| Hermannia gibba | GO:0016790 | molecular_function | thiolester hydrolase activity |
| Hermannia gibba | GO:0014883 | biological_process | transition between fast and slow fiber |
| Hermannia gibba | GO:0047936 | molecular_function | glucose 1-dehydrogenase [NAD(P)] activity |
| Hermannia gibba | GO:0047935 | molecular_function | glucose 1-dehydrogenase (NADP+) activity |
| Hermannia gibba | GO:0031222 | biological_process | arabinan catabolic process |
| Hermannia gibba | GO:0070212 | biological_process | protein poly-ADP-ribosylation |
| Hermannia gibba | GO:0016192 | biological_process | vesicle-mediated transport |
| Hermannia gibba | GO:0019062 | biological_process | virion attachment to host cell |
| Hermannia gibba | GO:0007224 | biological_process | smoothened signaling pathway |
| Hermannia gibba | GO:0004674 | molecular_function | protein serine/threonine kinase activity |
| Hermannia gibba | GO:0008218 | biological_process | bioluminescence |
| Hermannia gibba | GO:0018279 | biological_process | protein N-linked glycosylation via asparagine |
| Hermannia gibba | GO:0042428 | biological_process | serotonin metabolic process |

| | | | |
|---|---|---|---|
| Hermannia gibba | GO:0030497 | biological_process | fatty acid elongation |
| Hermannia gibba | GO:0051598 | biological_process | meiotic recombination checkpoint |
| Hermannia gibba | GO:0008033 | biological_process | tRNA processing |
| Hermannia gibba | GO:0007586 | biological_process | digestion |
| Hermannia gibba | GO:0031638 | biological_process | zymogen activation |
| Hermannia gibba | GO:0050728 | biological_process | negative regulation of inflammatory response |
| Hermannia gibba | GO:0021762 | biological_process | substantia nigra development |
| Hermannia gibba | GO:0071555 | biological_process | cell wall organization |
| Hermannia gibba | GO:0007605 | biological_process | sensory perception of sound |
| Hermannia gibba | GO:0007155 | biological_process | cell adhesion |
| Hermannia gibba | GO:0006749 | biological_process | glutathione metabolic process |
| Hermannia gibba | GO:2001242 | biological_process | regulation of intrinsic apoptotic signaling pathway |
| Hermannia gibba | GO:0043312 | biological_process | neutrophil degranulation |
| Hermannia gibba | GO:0007286 | biological_process | spermatid development |
| Hermannia gibba | GO:0045892 | biological_process | negative regulation of transcription, DNA-templated |
| Hermannia gibba | GO:0048794 | biological_process | swim bladder development |
| Hermannia gibba | GO:0046693 | biological_process | sperm storage |
| Hermannia gibba | GO:0005524 | molecular_function | ATP binding |
| Hermannia gibba | GO:0006641 | biological_process | triglyceride metabolic process |
| Hermannia gibba | GO:0051725 | biological_process | protein de-ADP-ribosylation |
| Hermannia gibba | GO:0015939 | biological_process | pantothenate metabolic process |
| Hermannia gibba | GO:0008354 | biological_process | germ cell migration |
| Hermannia gibba | GO:0007584 | biological_process | response to nutrient |
| Hermannia gibba | GO:0042594 | biological_process | response to starvation |
| Hermannia gibba | GO:0006898 | biological_process | receptor-mediated endocytosis |
| Hermannia gibba | GO:0034446 | biological_process | substrate adhesion-dependent cell spreading |
| Hermannia gibba | GO:0004104 | molecular_function | cholinesterase activity |
| Hermannia gibba | GO:0006357 | biological_process | regulation of transcription from RNA polymerase II promoter |
| Hermannia gibba | GO:0045944 | biological_process | positive regulation of transcription from RNA polymerase II promoter |
| Hermannia gibba | GO:0006468 | biological_process | protein phosphorylation |
| Hermannia gibba | GO:0016614 | molecular_function | oxidoreductase activity, acting on CH-OH group of donors |
| Hermannia gibba | GO:0005886 | cellular_component | plasma membrane |
| Hermannia gibba | GO:0007399 | biological_process | nervous system development |
| Hermannia gibba | GO:0015893 | biological_process | drug transport |
| Hermannia gibba | GO:0010044 | biological_process | response to aluminum ion |
| Hermannia gibba | GO:0036372 | biological_process | opsin transport |
| Hermannia gibba | GO:0018394 | biological_process | peptidyl-lysine acetylation |

| | | | |
|---|---|---|---|
| Hermannia gibba | GO:0030162 | biological_process | regulation of proteolysis |
| Hermannia gibba | GO:0035556 | biological_process | intracellular signal transduction |
| Hermannia gibba | GO:0006114 | biological_process | glycerol biosynthetic process |
| Hermannia gibba | GO:0051131 | biological_process | chaperone-mediated protein complex assembly |
| Hermannia gibba | GO:0016021 | cellular_component | integral component of membrane |
| Hermannia gibba | GO:0000122 | biological_process | negative regulation of transcription from RNA polymerase II promoter |
| Hermannia gibba | GO:0006629 | biological_process | lipid metabolic process |
| Hermannia gibba | GO:0008542 | biological_process | visual learning |
| Hermannia gibba | GO:0048670 | biological_process | regulation of collateral sprouting |
| Hermannia gibba | GO:0005412 | molecular_function | glucose:sodium symporter activity |
| Hermannia gibba | GO:0006401 | biological_process | RNA catabolic process |
| Hermannia gibba | GO:0001894 | biological_process | tissue homeostasis |
| Hermannia gibba | GO:0048489 | biological_process | synaptic vesicle transport |
| Hermannia gibba | GO:0006904 | biological_process | vesicle docking involved in exocytosis |
| Hermannia gibba | GO:0048741 | biological_process | skeletal muscle fiber development |
| Hermannia gibba | GO:0051225 | biological_process | spindle assembly |
| Hermannia gibba | GO:0001932 | biological_process | regulation of protein phosphorylation |
| Hermannia gibba | GO:0006667 | biological_process | sphinganine metabolic process |
| Hermannia gibba | GO:0015031 | biological_process | protein transport |
| Hermannia gibba | GO:0030433 | biological_process | ER-associated ubiquitin-dependent protein catabolic process |
| Hermannia gibba | GO:0006099 | biological_process | tricarboxylic acid cycle |
| Hermannia gibba | GO:0005576 | cellular_component | extracellular region |
| Hermannia gibba | GO:0016705 | molecular_function | oxidoreductase activity, acting on paired donors, with incorporation or reduction of molecular oxygen |
| Hermannia gibba | GO:0006413 | biological_process | translational initiation |
| Hermannia gibba | GO:0006915 | biological_process | apoptotic process |
| Hermannia gibba | GO:0051301 | biological_process | cell division |
| Hermannia gibba | GO:0004930 | molecular_function | G-protein coupled receptor activity |
| Hermannia gibba | GO:0015739 | biological_process | sialic acid transport |
| Hermannia gibba | GO:0003281 | biological_process | ventricular septum development |
| Hermannia gibba | GO:0030163 | biological_process | protein catabolic process |
| Hermannia gibba | GO:0050661 | molecular_function | NADP binding |
| Hermannia gibba | GO:0019285 | biological_process | glycine betaine biosynthetic process from choline |
| Hermannia gibba | GO:0005737 | cellular_component | cytoplasm |
| Hermannia gibba | GO:0002532 | biological_process | production of molecular mediator involved in inflammatory response |
| Hermannia gibba | GO:0042802 | molecular_function | identical protein binding |
| Hermannia gibba | GO:0000027 | biological_process | ribosomal large subunit assembly |
| Hermannia gibba | GO:0060074 | biological_process | synapse maturation |

| | | | |
|---|---|---|---|
| Hermannia gibba | GO:0016787 | molecular_function | hydrolase activity |
| Hermannia gibba | GO:0042626 | molecular_function | ATPase activity, coupled to transmembrane movement of substances |
| Hermannia gibba | GO:0007166 | biological_process | cell surface receptor signaling pathway |
| Hermannia gibba | GO:0006260 | biological_process | DNA replication |
| Hermannia gibba | GO:1902600 | biological_process | hydrogen ion transmembrane transport |
| Hermannia gibba | GO:0071625 | biological_process | vocalization behavior |
| Hermannia gibba | GO:0023021 | biological_process | termination of signal transduction |
| Hermannia gibba | GO:0031146 | biological_process | SCF-dependent proteasomal ubiquitin-dependent protein catabolic process |
| Hermannia gibba | GO:0009058 | biological_process | biosynthetic process |
| Hermannia gibba | GO:0001505 | biological_process | regulation of neurotransmitter levels |
| Hermannia gibba | GO:0042981 | biological_process | regulation of apoptotic process |
| Hermannia gibba | GO:0008039 | biological_process | synaptic target recognition |
| Hermannia gibba | GO:0030246 | molecular_function | carbohydrate binding |
| Hermannia gibba | GO:0008360 | biological_process | regulation of cell shape |
| Hermannia gibba | GO:0001516 | biological_process | prostaglandin biosynthetic process |
| Hermannia gibba | GO:0042311 | biological_process | vasodilation |
| Hermannia gibba | GO:0042381 | biological_process | hemolymph coagulation |
| Hermannia gibba | GO:0008063 | biological_process | Toll signaling pathway |
| Hermannia gibba | GO:0051124 | biological_process | synaptic growth at neuromuscular junction |
| Hermannia gibba | GO:0017148 | biological_process | negative regulation of translation |
| Hermannia gibba | GO:0009611 | biological_process | response to wounding |
| Hermannia gibba | GO:0051919 | biological_process | positive regulation of fibrinolysis |
| Hermannia gibba | GO:0009651 | biological_process | response to salt stress |
| Hermannia gibba | GO:0070445 | biological_process | regulation of oligodendrocyte progenitor proliferation |
| Hermannia gibba | GO:0097090 | biological_process | presynaptic membrane organization |

| | |
|---|---|
| 393 | 0 |
| 218 | 0 |
| 109 | 0 |
| 97 | 0 |
| 96 | 0 |
| 81 | 0,000034367491 |
| 78 | 0,000005110750 |
| 69 | 0,000423589745 |
| 61 | 0,000000001015 |
| 58 | 0 |
| 58 | 0,000000254281 |
| 57 | 0,000000019940 |
| 56 | 0 |
| 54 | 0,000002043210 |
| 53 | 0 |
| 52 | 0,00115515688 |
| 52 | 0,01332639083 |
| 51 | 0 |
| 49 | 0,000000013826 |
| 49 | 0,000059892375 |
| 47 | 0,005591452744 |
| 45 | 0,000000753731 |
| 45 | 0,000448761753 |
| 44 | 0 |
| 43 | 0 |
| 43 | 0,000384504443 |
| 42 | 0,000000011038 |
| 41 | 0 |
| 41 | 0 |
| 40 | 0,000069626576 |
| 38 | 0,000000000460 |
| 38 | 0,000000009701 |
| 37 | 0,000777424456 |
| 36 | 0 |
| 36 | 0,01159555823 |
| 34 | 0,006627196204 |
| 33 | 0 |

| | |
|---|---|
| 33 | 0,000000000355 |
| 33 | 0,005976474794 |
| 32 | 0 |
| 32 | 0,000000075256 |
| 31 | 0,002097853049 |
| 31 | 0,006910394862 |
| 30 | 0 |
| 29 | 0 |
| 28 | 0,000000001292 |
| 27 | 0 |
| 27 | 0,004073224594 |
| 25 | 0 |
| 25 | 0 |
| 25 | 0,000048247486 |
| 25 | 0,000290405615 |
| 25 | 0,000619273530 |
| 24 | 0,000032210444 |
| 24 | 0,0189512469 |
| 23 | 0 |
| 23 | 0,000020055253 |
| 23 | 0,000061664304 |
| 22 | 0 |
| 21 | 0,000824901667 |
| 20 | 0 |
| 20 | 0 |
| 20 | 0,002435563734 |
| 19 | 0 |
| 19 | 0,000018947918 |
| 19 | 0,006313250802 |
| 18 | 0,000000000733 |
| 18 | 0,000000042607 |
| 18 | 0,001112694461 |
| 18 | 0,003909999771 |
| 17 | 0 |
| 17 | 0,000002887599 |
| 17 | 0,000018137845 |
| 17 | 0,01811680497 |

| | |
|---|---|
| 16 | 0 |
| 16 | 0,000000674338 |
| 16 | 0,00847214736 |
| 15 | 0,000000202036 |
| 15 | 0,000435198990 |
| 15 | 0,00397240442 |
| 15 | 0,01585210179 |
| 14 | 0 |
| 14 | 0,000021531143 |
| 13 | 0 |
| 13 | 0,000000033729 |
| 13 | 0,000049499695 |
| 13 | 0,002450746861 |
| 12 | 0 |
| 12 | 0,000000000396 |
| 12 | 0,000000000785 |
| 12 | 0,000000052315 |
| 12 | 0,000006606316 |
| 12 | 0,000011517453 |
| 12 | 0,000038379939 |
| 12 | 0,000376376337 |
| 11 | 0,000006813929 |
| 11 | 0,001237051917 |
| 10 | 0 |
| 10 | 0,000000001341 |
| 10 | 0,000555911221 |
| 10 | 0,001961019126 |
| 10 | 0,006431860989 |
| 10 | 0,007841182713 |
| 10 | 0,01560765527 |
| 9 | 0 |
| 9 | 0,000000090876 |
| 9 | 0,000087922513 |
| 9 | 0,000128457930 |
| 9 | 0,000141115371 |
| 9 | 0,005572395709 |
| 9 | 0,01220925383 |

| | |
|---|---|
| 9 | 0,02244524873 |
| 8 | 0,000000080942 |
| 8 | 0,000000322557 |
| 8 | 0,000001209582 |
| 8 | 0,000077260343 |
| 8 | 0,000890064653 |
| 8 | 0,0158252778 |
| 8 | 0,01595818221 |
| 7 | 0 |
| 7 | 0,000002458308 |
| 7 | 0,000032023244 |
| 7 | 0,000817318696 |
| 7 | 0,02219294161 |
| 6 | 0 |
| 6 | 0,000000000978 |
| 6 | 0,000004849790 |
| 6 | 0,000006164042 |
| 6 | 0,000007689790 |
| 6 | 0,000073584943 |
| 6 | 0,000585204427 |
| 6 | 0,000585204427 |
| 6 | 0,001120802734 |
| 6 | 0,00131889239 |
| 6 | 0,007168040946 |
| 6 | 0,01081622781 |
| 6 | 0,01604855695 |
| 6 | 0,02171443967 |
| 5 | 0 |
| 5 | 0 |
| 5 | 0 |
| 5 | 0 |
| 5 | 0,000000011020 |
| 5 | 0,000000054588 |
| 5 | 0,000037440776 |
| 5 | 0,000037440776 |
| 5 | 0,000037440776 |
| 5 | 0,000195545913 |

| | |
|---|---|
| 5 | 0,001916071674 |
| 5 | 0,003710465034 |
| 5 | 0,007520518145 |
| 5 | 0,007520518145 |
| 4 | 0 |
| 4 | 0 |
| 4 | 0,000000766530 |
| 4 | 0,000002269661 |
| 4 | 0,000288536507 |
| 4 | 0,000288536507 |
| 4 | 0,000386714783 |
| 4 | 0,000592700405 |
| 4 | 0,00205214829 |
| 4 | 0,003724009679 |
| 4 | 0,008769490274 |
| 4 | 0,01153538441 |
| 4 | 0,01813818659 |
| 4 | 0,02630243786 |
| 3 | 0 |
| 3 | 0 |
| 3 | 0 |
| 3 | 0 |
| 3 | 0 |
| 3 | 0 |
| 3 | 0,000004004545 |
| 3 | 0,000005119737 |
| 3 | 0,000080968723 |
| 3 | 0,000129089216 |
| 3 | 0,000182592962 |
| 3 | 0,000204840947 |
| 3 | 0,000709818497 |
| 3 | 0,000885607280 |
| 3 | 0,002901859203 |
| 3 | 0,005409216694 |
| 3 | 0,01200830588 |
| 3 | 0,01322733384 |
| 3 | 0,01680058709 |

| | |
|---|---|
| 2 | 0 |
| 2 | 0 |
| 2 | 0,000000001579 |
| 2 | 0,000000004082 |
| 2 | 0,000000432394 |
| 2 | 0,000000492851 |
| 2 | 0,000001396624 |
| 2 | 0,000007423432 |
| 2 | 0,000020439710 |
| 2 | 0,000033752351 |
| 2 | 0,000062870568 |
| 2 | 0,000102973976 |
| 2 | 0,000189714126 |
| 2 | 0,000391356499 |
| 2 | 0,000496994541 |
| 2 | 0,000898057059 |
| 2 | 0,003193229662 |
| 2 | 0,004471867346 |
| 2 | 0,005583458327 |
| 2 | 0,006956402501 |
| 2 | 0,008647214468 |
| 2 | 0,008647214468 |
| 2 | 0,01635870746 |
| 2 | 0,01704597343 |
| 2 | 0,01704597343 |

| GO.ID | Term | Annotated |
|---|---|---|
| GO:0043130 | ubiquitin binding | 13 |
| GO:0016817 | hydrolase activity, acting on acid anhyd... | 127 |
| GO:1990756 | ubiquitin ligase-substrate adaptor activ... | 2 |
| GO:0017150 | tRNA dihydrouridine synthase activity | 2 |
| GO:0070577 | lysine-acetylated histone binding | 2 |
| GO:0000213 | tRNA-intron endonuclease activity | 2 |
| GO:0008046 | axon guidance receptor activity | 2 |
| GO:0004435 | phosphatidylinositol phospholipase C act... | 2 |
| GO:0000400 | four-way junction DNA binding | 2 |
| GO:0000340 | RNA 7-methylguanosine cap binding | 2 |
| GO:0004222 | metalloendopeptidase activity | 18 |
| GO:0007131 | reciprocal meiotic recombination | 7 |
| GO:0016477 | cell migration | 22 |
| GO:0006812 | monoatomic cation transport | 68 |
| GO:0007274 | neuromuscular synaptic transmission | 3 |
| GO:0045003 | double-strand break repair via synthesis... | 2 |
| GO:0007220 | Notch receptor processing | 2 |
| GO:0048208 | COPII vesicle coating | 2 |
| GO:0000379 | tRNA-type intron splice site recognition... | 2 |
| GO:0043328 | protein transport to vacuole involved in... | 2 |
| GO:0051209 | release of sequestered calcium ion into ... | 2 |
| GO:0000723 | telomere maintenance | 2 |
| GO:0006559 | L-phenylalanine catabolic process | 2 |
| GO:0048280 | vesicle fusion with Golgi apparatus | 2 |
| GO:0015749 | monosaccharide transmembrane transport | 2 |
| GO:0016485 | protein processing | 18 |
| GO:0019005 | SCF ubiquitin ligase complex | 9 |
| GO:0009986 | cell surface | 11 |
| GO:0030904 | retromer complex | 2 |
| GO:0033063 | Rad51B-Rad51C-Rad51D-XRCC2 complex | 2 |
| GO:0055038 | recycling endosome membrane | 2 |
| GO:0000214 | tRNA-intron endonuclease complex | 2 |
| GO:0016342 | catenin complex | 2 |

| Significant | Expected | classicFisher | FDR |
|---|---|---|---|
| 3 | 0,26 | 0,0018 | 0,36 |
| 4 | 2,51 | 0,0195 | 0,36 |
| 1 | 0,04 | 0,0392 | 0,412631578947368 |
| 1 | 0,04 | 0,0392 | 0,412631578947368 |
| 1 | 0,04 | 0,0392 | 0,412631578947368 |
| 1 | 0,04 | 0,0392 | 0,412631578947368 |
| 1 | 0,04 | 0,0392 | 0,412631578947368 |
| 1 | 0,04 | 0,0392 | 0,412631578947368 |
| 1 | 0,04 | 0,0392 | 0,412631578947368 |
| 1 | 0,04 | 0,0392 | 0,412631578947368 |
| 2 | 0,36 | 0,0478 | 0,431111111111111 |
| 2 | 0,14 | 0,0074 | 0,351818181818182 |
| 3 | 0,43 | 0,0083 | 0,351818181818182 |
| 4 | 1,33 | 0,0371 | 0,351818181818182 |
| 2 | 0,06 | 0,0378 | 0,351818181818182 |
| 1 | 0,04 | 0,0387 | 0,351818181818182 |
| 1 | 0,04 | 0,0387 | 0,351818181818182 |
| 1 | 0,04 | 0,0387 | 0,351818181818182 |
| 1 | 0,04 | 0,0387 | 0,351818181818182 |
| 1 | 0,04 | 0,0387 | 0,351818181818182 |
| 1 | 0,04 | 0,0387 | 0,351818181818182 |
| 1 | 0,04 | 0,0387 | 0,351818181818182 |
| 1 | 0,04 | 0,0387 | 0,351818181818182 |
| 1 | 0,04 | 0,0387 | 0,351818181818182 |
| 1 | 0,04 | 0,0387 | 0,351818181818182 |
| 2 | 0,35 | 0,0468 | 0,396551724137931 |
| 2 | 0,19 | 0,014 | 0,7 |
| 2 | 0,23 | 0,021 | 0,7 |
| 1 | 0,04 | 0,041 | 0,745454545454546 |
| 1 | 0,04 | 0,041 | 0,745454545454546 |
| 1 | 0,04 | 0,041 | 0,745454545454546 |
| 1 | 0,04 | 0,041 | 0,745454545454546 |
| 1 | 0,04 | 0,041 | 0,745454545454546 |

| ontology |
| --- |
| Molecular_Function |
| Molecular_Function |
| Molecular_Function |
| Molecular_Function |
| Molecular_Function |
| Molecular_Function |
| Molecular_Function |
| Molecular_Function |
| Molecular_Function |
| Molecular_Function |
| Molecular_Function |
| Biological_Process |
| Biological_Process |
| Biological_Process |
| Biological_Process |
| Biological_Process |
| Biological_Process |
| Biological_Process |
| Biological_Process |
| Biological_Process |
| Biological_Process |
| Biological_Process |
| Biological_Process |
| Biological_Process |
| Biological_Process |
| Biological_Process |
| Cellular_Component |
| Cellular_Component |
| Cellular_Component |
| Cellular_Component |
| Cellular_Component |
| Cellular_Component |
| Cellular_Component |
